# Nementin is a Nematode-Selective Small Molecule Agonist of Neurotransmitter Release

**DOI:** 10.1101/2022.03.12.484074

**Authors:** Sean Harrington, Jessica J. Knox, Andrew R. Burns, Ken-Loon Choo, Aaron Au, Megan Kitner, Cecile Haeberli, Jacob Pyche, Cassandra D’Amata, Yong-Hyun Kim, Jonathan R. Volpatti, Maximillano Guiliani, Jamie Snider, Victoria Wong, Bruna M. Palmeira, Elizabeth M. Redman, Aditya S. Vaidya, John S. Gilleard, Igor Stagljar, Sean R. Cutler, Daniel Kulke, James J. Dowling, Christopher M. Yip, Jennifer Keiser, Inga Zasada, Mark Lautens, Peter J. Roy

**Affiliations:** Department of Pharmacology and Toxicology, University of Toronto, Toronto, ON, M5S 1A8, Canada; The Donnelly Centre for Cellular and Biomolecular Research, University of Toronto, Toronto, ON, M5S 3E1, Canada; Department of Molecular Genetics, University of Toronto; Toronto, ON, M5S 1A8, Canada; The Department of Chemistry, University of Toronto, 80 St. George Street, M5S 3H6, Toronto, Canada; Institute of Biomaterials and Biomedical Engineering, University of Toronto, Toronto, Ontario, Canada; USDA-ARS Horticultural Crops Research Laboratory, Corvallis, OR 97330, USA; Department of Medical Parasitology and Infection Biology; Swiss-Tropical and Public Health Institute, (Swiss TPH), Socinstr. 57, CH–4051 Basel, Switzerland; Faculty of Science; University of Basel, Petersplatz 1, 4001 Basel; Program in Developmental and Stem Cell Biology, The Hospital for Sick Children, Toronto, Ontario, Canada M5G 0A4; Program in Genetics and Genome Biology, The Hospital for Sick Children, Toronto, Ontario, Canada M5G 0A4; Department of Comparative Biology and Experimental Medicine, Host-Parasite Interactions (HPI) Program, Faculty of Veterinary Medicine, University of Calgary, Calgary, AB, Canada; Institute for Integrative Genome Biology, University of California, Riverside, Riverside, CA 92521, USA; Department of Botany and Plant Sciences, University of California, Riverside, Riverside, CA 92521, USA; Department of Biochemistry, University of Toronto, Toronto, ON, M5S 1A8, Canada; Mediterranean Institute for Life Sciences, Meštrovićevo Šetalište 45, HR-21000, Split, Croatia; School of Medicine, University of Split, Šoltanska ul. 2, 21000, Split, Croatia; Research Parasiticides, Bayer Animal Health GmbH, Monheim, Germany

**Keywords:** chemical genetics, nematicide, anthelmintic, *C. elegans*, parasitic nematodes, *Strongyloides ratti*, *Meloidogyne hapla*, small molecule screens, early-stage drug discovery, dense core vesicles, convulsions, dgk-1, egl-30, goa-1, unc-13, unc-17, unc-31, unc-43, unc-64

## Abstract

Nematode parasites of humans, livestock and crops pose a significant burden on human health and welfare. Alarmingly, parasitic nematodes of animals have rapidly evolved resistance to anthelmintic drugs, and traditional nematicides that protect crops are facing increasing restrictions because of poor phylogenetic selectivity. Here, we present a pipeline that exploits multiple motor outputs of the model nematode *C. elegans* for nematicide discovery. This pipeline yielded multiple compounds that selectively kill and/or immobilize diverse nematode parasites. We focus on one compound that induces violent convulsions and paralysis that we call Nementin. We find that Nementin agonizes neuronal dense core vesicle release, which in turn agonizes cholinergic signaling. Consequently, Nementin synergistically enhances the potency of widely-used non-selective acetylcholinesterase inhibitors (AChEIs), but in a nematode-selective manner. Nementin therefore has the potential to reduce the environmental impact of toxic AChEI pesticides used to control nematode infections and infestations.

**Significance Statement:** Parasitic nematodes pose a considerable burden to human health and food security. Small molecules that have traditionally been used to control these parasites have either been banned because of toxicity concerns or are being rendered ineffective because of the evolution of resistance. Significant gaps in our nematicidal toolkit are therefore becoming an alarming problem. Here, we describe our discovery of Nementin, a small molecule that disrupts the nematode nervous system but is ineffective against non-targeted organisms. We find that Nementin also enhances the activity of non-selective pesticides but does so in a nematode-selective manner. Hence, Nementin is an innovative solution to combat parasitic nematodes in a safe and phylum-selective manner.

**One-Sentence Summary:** A *C. elegans*-based screening pipeline identifies a selective nematicide that also potentiates acetylcholinesterase inhibitors.

## Introduction

Parasitic nematodes are a scourge to humanity. Not only do helminths currently infect more than 1.5 billion people (WHO Soil Transmitted Helminth Fact Sheet, 2022) but nematode parasitism also destroys tens of billions of dollars (USD) worth of livestock annually^1^. Alarmingly, the rate of destruction is growing due to the evolution of anthelmintic resistance^2^. Plant-parasitic nematodes (PPNs) are even more destructive as they ruin over 125 billions of dollars (USD) worth of food crops annually^3^. Traditional nematicides that protect crops have been justifiably banned or severely restricted due to a lack of phylum selectivity^4^, but these restrictions severely impact food security^5^. Compounding these issues, global food demand is expected to increase by 70% by the year 2050^6^. Hence, the development of new and selective nematicides is essential to our collective welfare.

Here, we present a novel motor-centric screening pipeline to identify novel candidate nematicides. Disrupting a parasite’s motor control has repeatedly proven effective in mitigating nematode infection^7,8^. We therefore screened our collection of worm-active (wactive) compounds^9^ using two successive behavioural assays of the free-living nematode *Caenorhabditis elegans*. We identified four molecules that fail to elicit phenotypes in off-target systems at concentrations that kill multiple nematode species. We focused on one of these, called Nementin, that induces *C. elegans* hyperactivity within seconds of exposure, followed by whole-body convulsions minutes later. Nementin and its analogs selectively immobilize and/or kill nematode parasites of plants and mammals. Chemical-genetic analyses reveal that Nementin agonizes neuronal dense core vesicle release, which in turn, agonizes synaptic vesicle release. This insight led to the finding that Nementin enhances the paralytic effects of organophosphate and carbamate acetylcholinesterase inhibitors not only in *C. elegans*, but in nematode parasites of mammals and plants. We conclude that Nementin is a novel nematode-selective scaffold that may also improve the selectivity of broad-acting pesticides.

## Results

### 26 Compounds Disrupt Multiple *C. elegans* Motor Activities

To identify small molecules that disrupt nematode neuromuscular activity, we designed a 96-well plate imaging system that measures the egg-laying rate of adult *C. elegans* hermaphrodites (Fig. 1a-c). The regulation of *C. elegans* egg-laying relies on well-characterized cholinergic, serotonergic, GABAergic and peptidergic circuits^10^, and is therefore a paragon to assess neuromuscular perturbation by small molecules. To identify egg-laying stimulators, we screened small molecules in the background of M9 worm buffer^9^. Animals incubated in M9-vehicle control wells lay 1.1 eggs per hour on average (Fig. 1d; Supplementary Data 1). To identify egg-laying inhibitors, we screened small molecules in the background of exogenous serotonin plus nicotine that stimulates animals to lay an average of 7.4 eggs per hour (Fig. 1d; Supplementary Data 1).

**Fig. 1.**
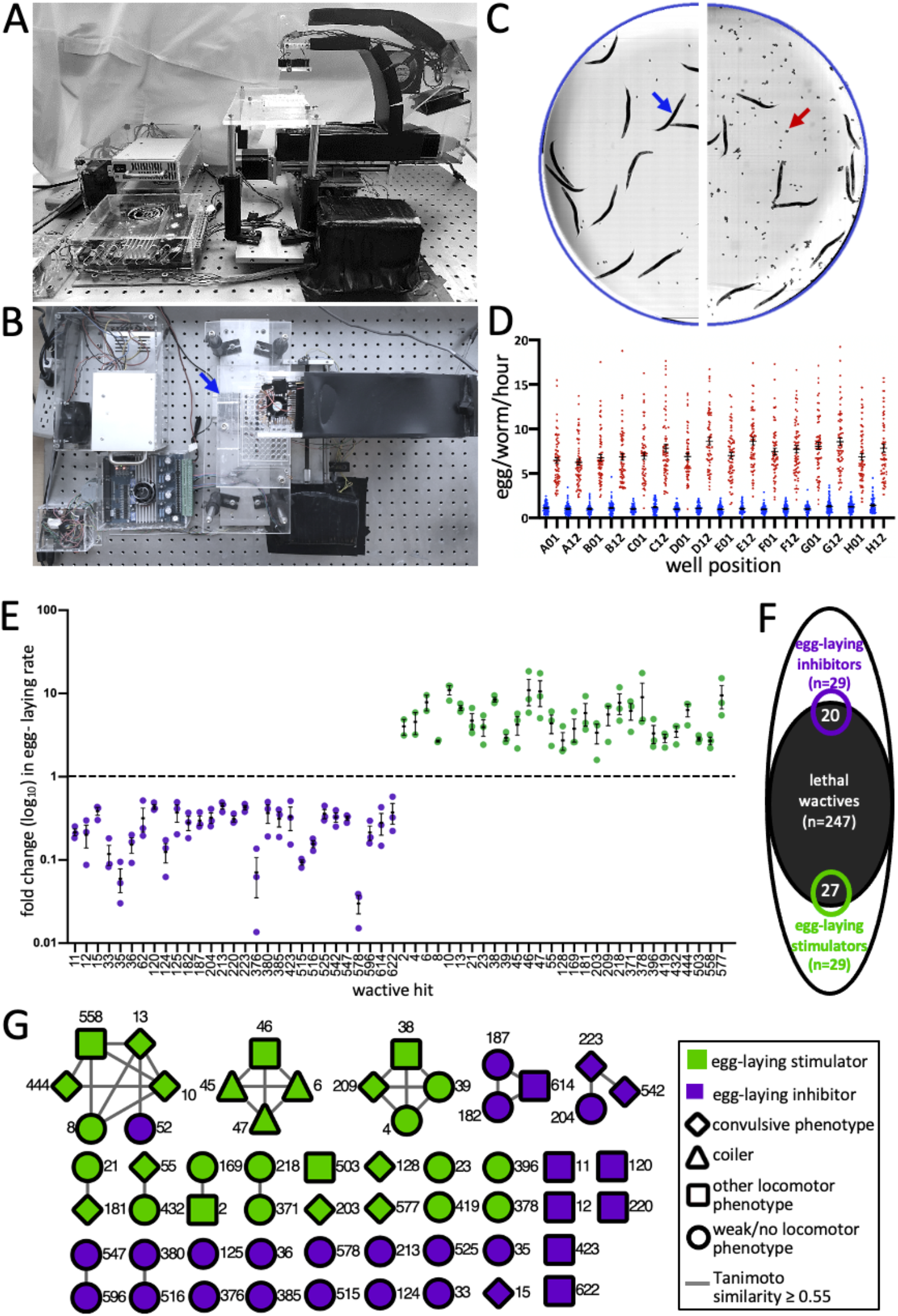
The Identification of 58 Small Molecules that Modulate the *C. elegans* Nervous System. (**a** and **b**) Two perspectives of the ‘2020 Imager’ showing one 96-well plate in the imager holder (arrow). (**c**) Representative half-well images of vehicle controls in the basal (M9-only) background (left) and the ‘stimulatory cocktail’ of 12.3 mM serotonin creatine sulfate and 7.7 mM nicotine (right) after 1 hour. A blue arrow points to an adult hermaphrodite and a red arrow points to a pair of embryos. (**d**) The distribution of counts of eggs/worm/hour for the basal background (blue) and the stimulatory background (red) from the indicated well-coordinates over 22 experimental replicates (96-well plates). The mean and SEM is shown. (**e**) A screen of the worm active (wactive) library yields 29 inhibitors (purple) and 29 stimulators (green) of the *C. elegans* egg-laying rate. Mean Egl rate and SEM is shown. (**f**) Venn diagram showing the overlap of the wactive molecules that are lethal (at 60 μM) with those that modulate egg-laying among the 486 molecules assayed in both assays. (**g**) A structure similarity network of the 58 egg-laying modulators. Each node represents a numbered wactive molecule. Connecting lines indicate shared structural similarity. Additional information is provided in Supplementary Table 1.

A screen of the 486 molecules from our worm-active library^9^ revealed 29 stimulators and 29 inhibitors of egg-laying (Fig. 1e; Supplementary Data 2). We cross-referenced the list of acute egg-laying modulators to the list of 247 wactive molecules that are lethal in six-day viability assays^9^. We found a 1.8-fold enrichment of lethality among the stimulators (*p*<0.001) (Fig. 1f), suggesting that excessive neuromuscular modulation may lead to death. A 1.4-fold enrichment of lethality among the inhibitors was also found (*p*<0.05), which is not unexpected given that egg-laying inhibition is certain to be a consequence of molecules that kill rapidly. A Tanimoto structural similarity cut-off of 0.55 reveals that the 58 egg-laying modulators can parsed into 37 distinct core scaffolds, consisting of 11 clusters and 26 singletons^11^ (Fig. 1g). 10 of the 11 clusters are composed of molecules with the identical egg-laying phenotype, which validates the phenotypic assignments.

We reasoned that molecules that disrupt multiple motor circuits are more likely to disrupt the parasitic nematode lifecycle. We therefore surveyed the 58 egg-laying modulators for those that disrupt the normal sinusoidal locomotion of wild type *C. elegans*. 26 of the 58 egg-laying modulators (45%) induce at least one locomotory phenotype such as convulsions, coiling, and paralysis (Supplementary Table 1; Fig. 1g; Supplementary Movies 1-6; Supplementary Data 3). The molecules that induce slow movement and paralysis are dominated (88%) by molecules that inhibit egg-laying (Supplementary Table 1). This is consistent with the idea that the egg-laying inhibitor class may be enriched with acutely lethal molecules. By contrast, the molecules that induce convulsions, shaking, and coiling are dominated (80%) by those that enhance egg-laying (Supplementary Table 1).

### Nementin-1 is Effective Against Multiple Parasitic Nematodes

To prioritize the 26 neuromodulatory molecules, we eliminated those from consideration that: i) are cytotoxic to human HEK293 cells; ii) induce developmental defects in zebrafish; iii) have lackluster lethality against the five nematodes previously assayed; iv) induce subtle locomotory phenotypes (reversal-defective), or v) were closely related to the commercial nematicide fluopyram^9^ (Supplementary Table 1; Supplementary Data 4). This left <u>w</u>orm-<u>act</u>ive (wact) molecules 10, 13, 15, 55, 120, 128, and 444 for further consideration (Fig. 2a; Supplementary Table 1).

**Fig. 2.**
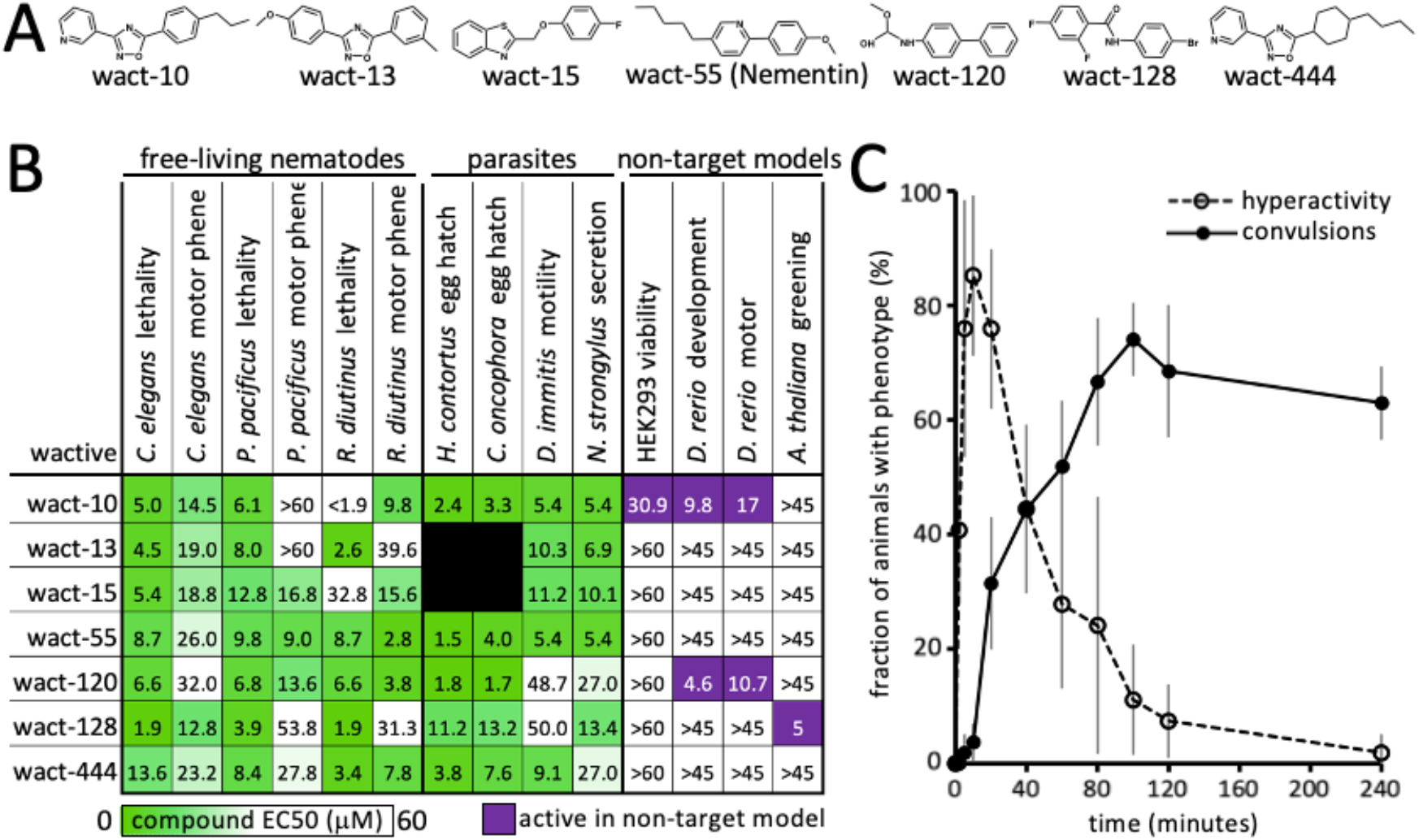
The motor-centric pipeline yields convulsing-inducing Nementin-1. (**a**) Structures of the 7 prioritized neuromodulators. (**b**) Bioactivity of the seven prioritized molecules. Data are EC50s except for the *A. thaliana* greening assay, which is the lowest concentration that yields a discernible difference from control. (**c**) A time-course analysis of 60 μM Nementin-1-induced locomotory phenotypes. Data are the mean of biological triplicate measurement of 18 animals per each time point. The standard deviation is shown.

We further assessed the activity of our seven hits against seven nematode species from three phylogenetic clades (Fig. 2b). Four of these seven species are parasites of mammals. We also analyzed (or reanalyzed) the molecules’ activity against the non-target models *D. rerio* fish, HEK293 cells, and *Arabidopsis thaliana* plants (Fig. 2b; Supplementary Fig. 1). We prioritized wact-55 because it arguably demonstrates the best combination of broad nematode activity and selectivity. SciFinder-based literature searches failed to reveal prior annotation of nematicidal activity for wact-55’s alkyl phenylpiperidine core scaffold^12^.

A detailed temporal analysis showed that within 90 seconds of exposure of 60 μM wact-55, *C. elegans* exhibits a hyperactive nose movement phenotype (Fig. 2c). The hyperactive phenotype dissipates at the expense of spastic convulsions over the course of 2 hours (Fig. 2c). By 24 hours, 100% of the adult animals incubated in wact-55 are dead (non-moving and/or disintegrating; three trials, n = 18 animals per trial). For reasons outlined below, we renamed the wact-55 molecule ‘Nementin-1’ (<u>nem</u>atode <u>e</u>nhancer of <u>n</u>eurotransmissio<u>n</u>-1).

To determine whether Nementin-1 might have a canonical mechanism-of-action (MOA), we tested whether any of the *C. elegans* mutant strains that resist the effects of seven popular anthelmintics also resist the lethal effects of Nementin-1. All drug-resistant strains remain sensitive to Nementin-1 (Supplementary Fig. 2). Furthermore, we previously showed that *C. elegans* cannot be easily mutated to resist Nementin-1’s lethality; 290,000 randomly mutated genomes failed to yield wact-55-resistant mutants^9^. These results suggest that: i) Nementin-1 does not share an MOA with canonical anthelmintics, ii) Nementin-1’s MOA may not be limited to a single protein target, and, iii) genetic resistance to Nementin-1 may be difficult to achieve in the field. All of these properties make Nementin-1 an attractive hit for further investigation.

### Nementin Convulsions are Dependent on Dense Core Vesicle Release

We reasoned that mutations in the pathway targeted by Nementin-1 might phenocopy the motor defects induced by the molecule and provide mechanistic insight. A survey of the literature yielded eight mutants that share at least some of Nementin-1’s phenotypes, including *acr-2, unc-2, unc-43, unc-58, unc-93, sup-9, sup-10*, and *twk-18*^13-21^ (Supplementary Table 2). We tested the hypothesis that Nementin-1 disrupts the pathway disrupted by each of these mutants. We did this by asking whether known suppressors of the phenocopying mutants can suppress Nementin-1’s effects (see Supplementary Table 2). Of the potential suppressing mutants tested, only the *unc-43* gain-of-function (GF) mutant resisted Nementin-1’s convulsions (Fig. 3a). We also found the converse to be true; *unc-43* reduction-of-function (RF) alleles enhance Nementin-1 convulsions (Fig. 3a). Remarkably, the interaction between the *unc-43* GF and Nementin-1 is mutual in that Nementin-1 can rescue the paralysis of *unc-43* GF (Fig. 3b). This observation argues against the idea that the *unc-43* GF mutant alters Nementin-1 absorption or metabolism. These results suggest that Nementin-1 may be disrupting UNC-43 pathway activity.

**Fig. 3.**
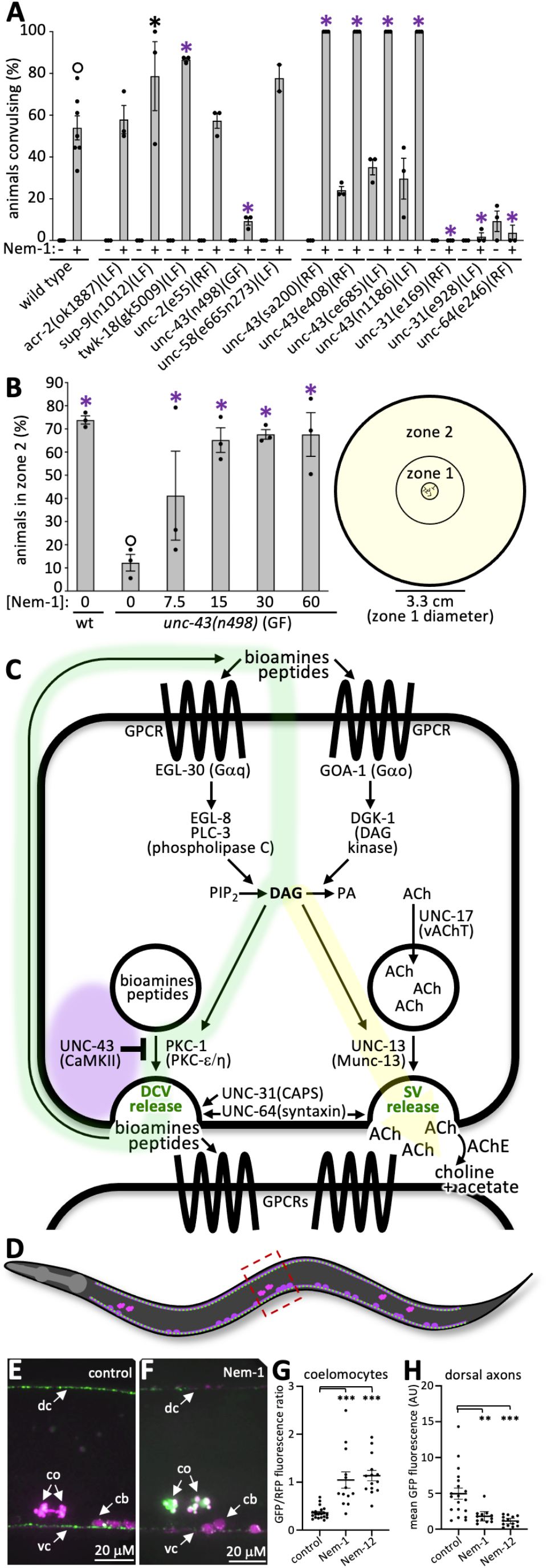
Nementin agonizes dense core vesicle release. (**a**) Convulsions induced by Nementin-1 (Nem-1) (60 μM) in the indicated genetic background after 80 minutes. The mean of three biological replicates with 18 animals per trial is shown. (**b**) The mean percentage of animals of the indicated genotype (three trials with ∼150 animals per trial) that locomote to zone 2 over 180 minutes after being placed in the centre of a plate (see schematic on right). For A and B, black and purple asterisks represent *p*<0.05 and p<0.001, Chi-square test of independence with Bonferroni correction relative to control (open circle); the standard error of the mean (SEM) is shown. (**c**) A schematic of neuronal signaling pathways relevant to Nementin activity. Purple glow indicates area of possible Nementin action; the green glow indicates how DCV neurotransmitter cargo may activate Gα_q_ and Gα_o_ signaling pathways; the yellow glow indicates how the secondary messenger diacylglycerol (DAG) can also promote synaptic vesicle release. (**d**) Schematic of adult (strain KG4247) expressing INS-22::GFP (green dots; packaged into DCVs) and secreted mCherry (fuchsia) in cholinergic motor neurons (cell bodies are oval). mCherry is constitutively secreted and taken up by the coelomocytes (three pairs of pink objects in the animal). Box indicates the area shown in E and F. Anterior is the left and dorsal is up. (**e** and **f**) Images of the midbody region of control and 60 μM Nementin-1-treated KG4247 animals after 4 hours. dc, dorsal cord; vc, ventral cord; co, coelomocytes; cb, cell bodies. (**g**) Quantification of the midbody coelomocytes (ratio of measured GFP/RFP) and dorsal cord fluorescence (relative to background tissues) of the animal. AU, arbitrary units., 1-way ANOVA with Dunnett correction for multiple comparisons; ** *p*<0.01; *** *p*<0.001; SEM is shown. Other regions of the animal are quantified in Supplementary Fig. 3.

UNC-43 is the *C. elegans* ortholog of CaMKII (calcium/calmodulin-dependent protein kinase II) and is a key negative regulator of dense core vesicle (DCV) release in *C. elegans* neurons^14,22^ (Fig. 3c). In animals with reduced UNC-43 function, neuronal DCV content is released in excess and is likely responsible for the mutant’s convulsive and constitutive egg-laying phenotypes^13,22^. In *unc-43* GF animals, DCVs accumulate, and their limited release is likely responsible for the mutant’s severely lethargic locomotion^22^.

To examine DCV behaviour in response to Nementin-1, we exploited the neuropeptide reporter (INS-22::GFP) that is packaged into DCVs in cholinergic motor neurons^22^. Like *unc-43* RF mutants^22^, worms treated with either Nementin-1 or a second analog called Nementin-12 have reduced axonal INS-22::GFP signal (*p*<0.05) (Fig. 3d-f,h; Supplementary Fig. 3c,d). Coincidentally, INS-22::GFP accumulates in the pseudocoelomic fluid-scavenging cells, called coelomocytes, of Nementin-treated animals ().

We reasoned that if Nementin disrupts motor behavior by agonizing DCV release, then disruption of UNC-31 (calcium-dependent activator protein for secretion (CAPS)), which is required for DCV release^22,23^, or UNC-64 syntaxin, which is required for all vesical fusions^24,25^, should suppress the Nementin-1-driven locomotory defects. Indeed, we find that both the weaker temperature-sensitive *unc-31(e169)*, the *unc-31(e928)* null, and the canonical *unc-64(e246)* RF mutants suppress the convulsions induced by Nementin-1 (Fig. 4a). Together, these data indicate that Nementin-induced DCV release is responsible for convulsions.

**Fig. 4.**
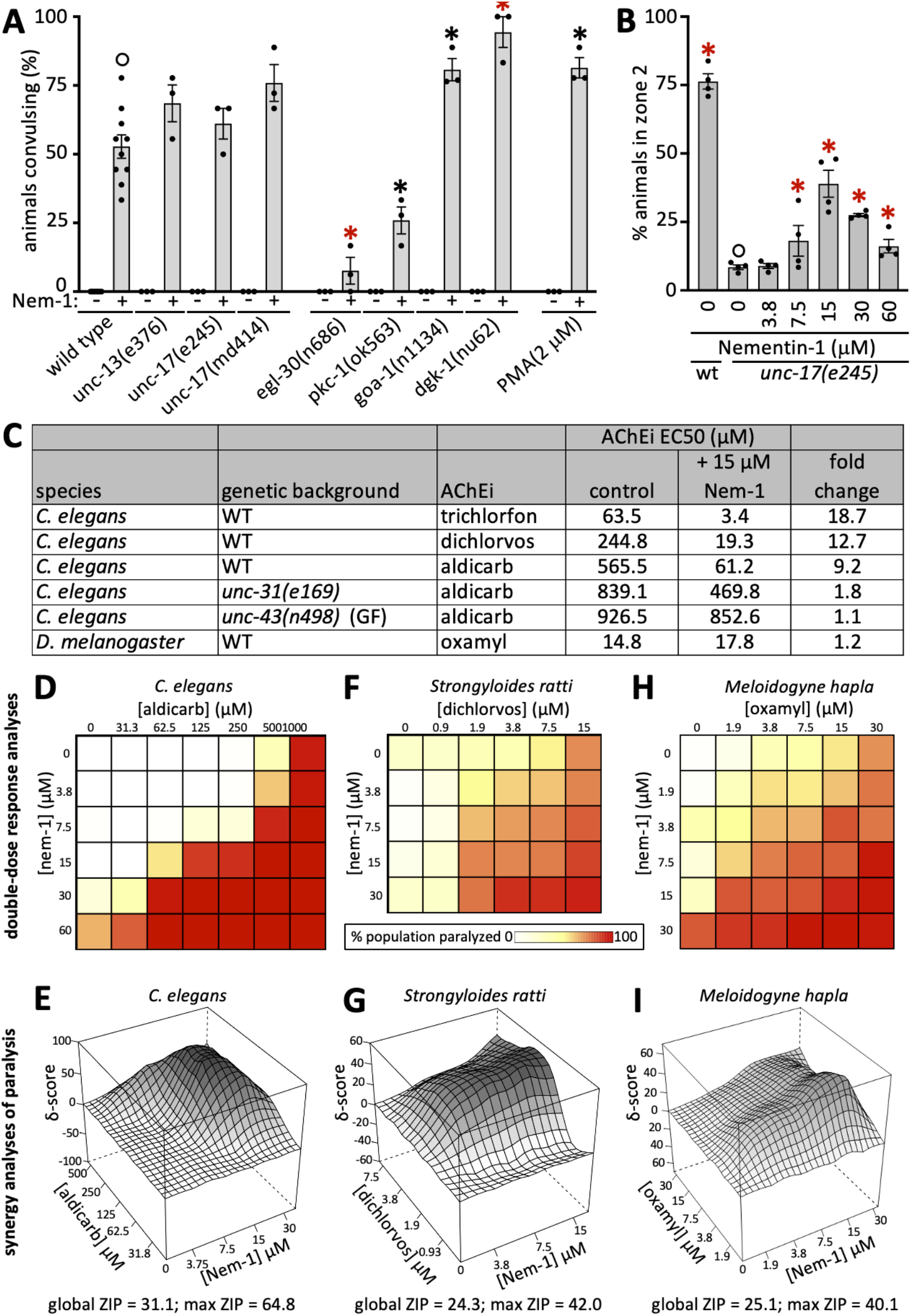
Nementin enhances AChE inhibitor activity via agonism of DCV release. (**a**) Convulsions induced after 80 minutes of exposure to Nementin-1 (Nem-1) (60 μM) or vehicle control in the background of the indicated mutant gene, or in the background of 24 hours of preincubation with the DAG memetic PMA. (**b**) The ability of wild type (WT) or *unc-17* mutant animals to productively locomote to zone 2 over 180 minutes after being placed in the centre of a plate (see schematic in Fig. 3b) in the indicated Nementin concentration. For both A and B, black asterisk *p*<0.01, red asterisk *p*<0.001, Bonferroni corrected Chi-square test of independence relative to control (open circle); SEM is shown. (**c**) Acetylcholinesterase inhibitor (AChEi) EC50s for different species and genetic backgrounds. Oxamyl is the only AChEi tested that exhibited activity against flies. Data are the mean of 3 biological replicates scoring 18 animals per condition. (**d-f**) Double dose-response analyses of Nementin-1 (y-axis) vs the indicated AChEi (x-axis) in the indicated nematode species. The scale for the percentage of the population paralyzed applies to d-f. (**g-i**) The corresponding zero interaction potency (ZIP) synergy score heatmaps for the double dose-response analyses presented in d-f. Max ZIP-score is defined as the maximal ZIP-score obtained from a combination of three concentrations of each compound in the double dose response. Shading highlights the topology of the plot and differs between g-i. N=3 trials for *elegans* with n=18 within each trial, N=6 trials for *ratti* with n≥30 within each trial, N≥3 trials for *hapla* with n≥15 within each trial.

Collectively, these results beg the question of whether UNC-43 is likely to be the physiologically relevant target of Nementin-1. Several observations argue against this. First, we note that the terminal phenotype of Nementin-1 is death, yet presumptive null alleles of *unc-43* (*ce685* and *n1186*) are viable^14,22^. Second, if Nementin-1 inhibits UNC-43, then Nementin-1 should not be able to enhance the convulsion phenotype of *unc-43* null mutants. We find that Nementin-1 does indeed enhance presumptive *unc-43* null mutants (*ce685* and *n1186*) (Fig. 3a). Third, if UNC-43 was Nementin-1’s sole physiological target, then *unc-43* GF mutations would have been identified in our genetic suppressor screens^9^, but no resistant mutants arose. Finally, the *unc-43* GF mutant (and *unc-31* and *unc-64* RF mutants) remain sensitive to the lethal effects of Nementin-1 (Supplementary Fig. 4). Together, these data argue against the idea that UNC-43 is Nementin-1’s sole physiologically relevant target. Instead, the similarities between Nementin-1-induced phenotypes and the *unc-43* RF mutants suggest that the compound targets a component(s) that acts with UNC-43 (Fig. 3c, purple glow).

### Nementin-1 Agonizes Cholinergic Signaling

We next investigated whether cholinergic signaling is also required for Nementin-1 activity. We tested whether mutants of the UNC-17 vesicular acetylcholine transporter (VAChT), which loads synaptic vesicles (SVs) with acetylcholine, or mutants of UNC-13, which is specifically required for SV release^26,27^, could suppress Nementin-1-induced convulsions. Reduction-of-function mutants of UNC-17 and UNC-13 remained sensitive to Nementin-1 (Fig. 4a). Incidentally, we found that Nementin-1 could partially rescue the lethargic locomotion of *unc-17* RF mutants (Fig. 4b). These data show that despite cholinergic signalling being dispensable for Nementin-1-induced convulsions, Nementin-1 may nevertheless agonize cholinergic signalling.

We further investigated the idea that Nementin-1 agonizes cholinergic signalling. To do so, we used the acetylcholinesterase (AChE) inhibitor aldicarb, which is a well-characterized tool used to investigate perturbations of cholinergic signaling in *C. elegans*^28^. AChE catabolizes acetylcholine at the neuromuscular junction (NMJ) and its inhibition increases acetylcholine levels that in turn paralyzes the animal^29^ (see Fig. 3c). We found that Nementin-1 sensitizes animals to the paralytic effects of aldicarb (Fig. 4c), suggesting that Nementin-1 agonizes acetylcholine release at the NMJ. The relationship between Nementin-1 and aldicarb is synergistic in nature, yielding a zero-interaction potency (ZIP) δ-score of 31.1 with a δ-score of 64.8 over the most synergistic set of concentrations (ZIP scores >10 are considered synergistic^30,31^ (Fig. 4d,e)). Nementin-1 also enhances the paralysis induced by dichlorvos and trichlorfon, which are two other AChE inhibitors that are also commercial anthelmintics, by 12.7 and 18.7-fold, respectively (Fig. 4c). By contrast, Nementin-1 fails to enhance the effects of an AChE-inhibitor on the fruit fly *D. melanogaster* (Fig. 4c). These data provide additional support for the idea that Nementin-1 agonizes cholinergic signalling and does so in a nematode-selective manner.

### Cholinergic Agonism is a Secondary Effect of DCV Release

Neuropeptides released from DCVs are known to interact with G-protein coupled-receptors (GPCRs) that signal through G-proteins^32^. In *C. elegans* neurons, the Gα_q_ protein EGL-30 stimulates DAG production via the EGL-8 and PLC-3 phospholipases^33^ (see Fig. 3c). By contrast, the Gα_o_ protein GOA-1 reduces DAG abundance by triggering its phosphorylation by diacylglycerol kinase DGK-1^34^. Neuronal DAG primes both DCV a. nd SV release via interaction with PKC-1(PKC-ε/η)^35^ and UNC-1 ^36^, respectively. The bioamines and neuropeptides that are released from the DCVs can have both paracrine and autocrine activity^37,38^. This circuitry raised the possibility that Nementin-1-induced DCV release may initiate a positive feedback loop that amplifies DCV release and agonizes SV release as a secondary consequence (see green and yellow pathways in Fig. 3c).

We tested this hypothesis in three ways. First, we examined the impact of mutant components of the ‘green’ circuit depicted in Fig. 3c. We found that *egl-30* RF, which reduces DAG production, suppresses Nementin-1 convulsions (Fig. 4a). By contrast, *goa-1* RF and *dgk-1* loss-of-function (LF) mutants, which lack regulation of membrane DAG, enhance Nementin-1 convulsions. We also found that disruption of PKC-1 reduces Nementin-1 convulsions (Fig. 4a). Second, we asked whether the cell permeable DAG mimetic phorbol myristate acetate (PMA) enhances Nementin-1 convulsions and found that it did (Fig. 4a). Consistent with the lack of convulsions in *goa-1* RF mutants (not treated with Nementin; Figure 4a), PMA treatment (without Nementin) does not induce convulsions in wild type animals (Figure 4a). This indicates that PKC-1-agonism alone cannot severely disrupt motor activity. Finally, we asked whether Nementin-1’s agonism of cholinergic signaling is dependent on the machinery needed for DCV release. Indeed, we found that the *unc-31(e169)* RF mutation and the *unc-43(n498)* GF mutation suppress Nementin-1’s agonism of cholinergic signaling (Fig. 4c). Together, these observations are consistent with a model whereby Nementin-1 agonizes DCV release whose contents may stimulate autocrine/paracrine G-protein signaling, which in turn agonizes both more DCV release and cholinergic signaling.

### Nementin Disrupts Parasitic Nematodes of Plants and Animals

We carried out a small structure-activity relationship (SAR) analysis of Nementin analogs assayed against free-living nematodes, parasitic nematodes, and non-target models (Supplementary Table 3; Supplementary Data 4). We purchased 18 commercially available analogs of Nementin-1 and found that Nementin-12 was the most potent inducer of acute motor phenotypes (Supplementary Table 3). We expanded the SAR by synthesizing 22 analogs of Nementin-12 (see Materials and Methods). Nementins 1, 12-5, 13 and 14 demonstrated broad anthelmintic activity without inducing obvious phenotype against non-target organisms. In addition, Nementin analogs were identified with improved activity in each parasitic nematode model tested. These data suggest that the Nementin scaffold may be further refined to improve selectivity and potency.

Given that Nementin-1 synergistically enhances the activity of AChE inhibitors in *C. elegans*, we tested whether it might act similarly against parasitic nematodes. We tested Nementin-1’s interaction with the organophosphate dichlorvos against *Strongyloides ratti*, a nematode parasite of mammals. Dichlorvos is approved for anthelmintic use in mammals^39,40^. Indeed, we find that Nementin-1 synergistically enhances dichlorvos’ activity against *S. ratti* (Fig. 4f,g). Nementin-1 was also found to synergistically paralyze the plant-parasitic nematode *Meloidogyne hapla* with the carbamate AChE inhibitor oxamyl, which is approved for field use^39,40^ (Fig. 4h,i). The Nementins may therefore have added benefit against parasitic nematodes when used in combination with pesticides that are AChE inhibitors.

## Discussion

Here, we have developed a *C. elegans*-based pipeline to identify novel anthelmintic small molecules with phylum-selectivity. The pipeline was designed to yield broad-acting anthelmintics that acutely affect nematode behaviour. The strategy yielded four small molecules that exhibit phylum-selectivity within the limited range of organisms tested. We focused on one of these molecules, called Nementin-1, which induces *C. elegans* hyperactivity within seconds of exposure, followed by spastic convulsions minutes later. Several Nementin analogs affect parasitic nematodes from distinct clades at concentrations that fail to elicit obvious phenotypes from non-target organisms. Despite significant effort, we were not able to generate *C. elegans* mutants that resist the lethal effects of Nementin, suggesting that resistance in the wild may be difficult to achieve.

Our data support a model whereby Nementin exposure initiates DCV release, which in turn agonizes SV release that promotes cholinergic signaling. Nementin-induced convulsions depend on DCV release but not cholinergic signaling, indicating that agonized DCV release is key to Nementin’s ability to disrupt motor behavior. How Nementin agonizes DCV release remains unclear. Despite the phenotypic similarities between Nementin exposure and *unc-43* RF mutants, our data suggests that Nementin is unlikely to inhibit UNC-43 in any canonical fashion. However, because of CaMKII’s complexity and its many isoforms^14,41^, we cannot rule out the possibility that Nementin interacts with CaMKII in a complex and perhaps tissue-specific manner.

Convulsions in *C. elegans* and other animals are thought to manifest from an imbalance of excitatory and inhibitory neurotransmitter signals, termed E/I imbalance ^42^. Gain-of-function mutations in motor neuron cation channels like the ACR-2 acetylcholine receptor or the UNC-2 voltage-gated calcium channel simultaneously increase excitatory cholinergic signals and decrease inhibitory GABAergic signals that are relayed to muscles. This in turn generates an excitation-dominated E/I imbalance that culminate in convulsions ^18,19,38^. Several parallels exist between Nementin-treated wild-type worms and either the *unc-2* and/or *acr-2* gain-of-function mutants including hyperactive locomotion, convulsions, a phenotypic dependence on UNC-31 CAPS, and aldicarb hypersensitivity ^18,19,38^. Nementin-induced convulsions are independent of ACR-2, UNC-2, and robust cholinergic signaling. We therefore deduce that the likely E/I imbalance induced by Nementin must originate from signals that are released from neuronal dense core vesicles.

Nementin has several features that make it an attractive anthelmintic for further development. First, *in vitro* tests demonstrate that it exhibits potentially broad-spectrum activity while maintaining phylogenetic selectivity. Such a feature is key to the development of an environmentally safe anthelmintic. Second, Nementin analogs exhibit an *in vitro* potency that is comparable to several commercial anthelmintics. Third, the Nementin synthesis route is relatively simple and uses inexpensive starting materials (‘Chemistry’ in Supplementary Materials)^43^. Finally, Nementin has the potential to reduce the amount of non-selective pesticides that are notoriously released into the environment.

Despite the toxicity concerns over indiscriminate activity, many AChE inhibitors such as trichlorfon, dichlorvos, coumaphos, aldicarb, fosthiazate, chlorpyrifos and oxamyl remain approved for nematicidal field use as of 2021 because of limited control options^39^ (see the FDA Approved Animal Drug Products at https://tinyurl.com/by83ks98, the US Environmental Protection Agency at https://tinyurl.com/3n3kf34n; and the US Geological Survey pesticide study at https://tinyurl.com/bp5h5em4). Because of Nementin’s selectivity and synergistic effects, using it in combination with AChE inhibitors has the potential to maintain nematode-selectivity whilst simultaneously reducing the amount of non-selective pesticide applied to the environment.

## Materials and Methods

*Materials and Methods are provided in the Supplementary Information for this initial submission.

## Acknowledgements

We are grateful for the strains given to us by the *Caenorhabditis* Genetics Center (University of Minnesota), Marie-Anne Félix (IBENS, Paris), Ken Miller (Oklahoma Medical Research Foundation), and Mei Zhen and Wesley Hung (Lunenfeld-Tanenbaum Research Institute, Toronto). Many thanks to Dr. Doug Colwell and Dawn Gray at the Lethbridge and Agri-Food Canada Research Centre for supplying faeces from infected calves for *Cooperia oncophora* egg collection.

## Ethics Statement

DK: Experiments on *D. immitis* and *N. brasiliensis* were performed in the laboratories of Bayer Animal Health GmbH (Monheim, Germany) in accordance with the local Animal Care and Use Committee and governmental authorities. JK: The generation of *N. americanis, T. muris*, S. ratti, and *H. polygyrus* and in vitro studies were carried out at the Swiss Tropical Institute (Basel, Switzerland), in accordance with both cantonal (license no. 2070) and Swiss national regulations on animal experimentation. JD: All zebrafish experiments were performed in compliance with any relevant ethical regulations, specifically following an institutionally reviewed and approved animal use protocol as well as the policies and guidelines of the Canadian Council on Animal Care and Animals for Research Act of Ontario.

## Author Contributions

Conceptualization: PJR, SH

Methodology: SH, JJK, K-L C, AA, MK, CH, JP, JRV, MG, JS, VW, BMP, EMR, ASV,

Investigation: SH, JJK, AB, MK, CH, JP, C’DA, Y-H K, JRV, MG, JS, VW, BMP, EMR, ADS

Visualization: PJR, SH

Funding acquisition: PJR

Project administration: PJR

Supervision: PJR, JG, IS, SRC, JJD, CMY, JK, IZ, ML

Writing – original draft: PJR, SH, JJK, K-LC, AA, SRC, DK, JK

Writing – review & editing: PJR, SH, AB, JJK, SC, DK, JK, IZ, ML

## Data and materials availability

Original data for all analyses presented are included in Supplementary Data 1-S4. Original images for analyses can be made available per request through the corresponding author P.J.R.

## Supplementary Materials

Materials and Methods

Supplementary Figures 1-4

Supplementary Table 1-3

Supplementary Movies 1 to 6 (description)

Supplementary Data 1 to S4 (description)

