## Supplemental Materials (Methods, Figures, Tables) for "Nementin is a Nematode-Selective Small Molecule Agonist of Neurotransmitter Release"

**This PDF file includes:**

Materials and Methods
Supplementary Figs. 1 to 4
Supplementary Tables 1 to 3
Captions for Supplementary Movies 1 to 6
Captions for Data S1 to S4
References

**Other Supplementary Materials for this manuscript include the following:**

Supplementary Movies 1 to 6

Supplementary Data 1. Egg-laying rates with and without a stimulatory cocktail.

Supplementary Data 2. The Wactive Library egg-laying screen data.

Supplementary Data 3. The locomotory survey of the egl-modulators.

Supplementary Data 4. Nematode and Counter-Screen Bioassay Data.

### Materials and Methods

Free-Living Nematode Strains and Culture

All nematode strains were cultured using standard methods at 20 °C unless otherwise indicated (1). The N2 (wild-type) strain of *Caenorhabditis elegans, Caenorhabditis briggsae* strain AF16 and *Pristionchus pacificus* strain PS312 were all obtained from the *C. elegans* Genetic Center (CGC; University of Minnesota). *Rhabditophanes sp. KR3021* was obtained from Marie-Anne Félix (Institute of Biology of the Ecole Normale Supérieure (IBENS), Paris, France). Mutant *C. elegans* strains were also obtained from the *C. elegans* Genetic Center.

*C. elegans* Small Molecule Screens and Phenotypic Analyses

A library of 486 small-molecules (Chembridge) previously found to induce phenotypes in *C. elegans* (486 worm actives, aka wactives; 26108372) were tested for their ability to modulate *C. elegans* egg-laying. Drug dilutions were prepared from stock plates using a 96-well pinner tool (FP3S200 V&P Scientific, Inc.) transferring 0.3 mL of drug stock solution prepared in DMSO.

*C. elegans Egg-laying* Assay

To screen for small molecules that modulate egg laying, ~20 young-adult wild-type (N2 Bristol) animals were pipetted in 15 mL of M9 buffer to 96-well flat-bottom polystyrene plates (2024-06 TC plate – Sarstedt) to a final volume of 50 mL containing 60 µM of test molecules in either 1) M9 buffer to identify egg-laying stimulators; or 2) a combination of 12.3 mM serotonin creatine sulfate monohydrate (H775 - Sigma-Aldrich) and 7.7 mM nicotine (N3876 - Sigma-Aldrich) in M9 buffer that induces a robust egg-laying response to identify egg-laying inhibitors (aka NS condition). Screen molecules were transferred to test wells using a 96-well pinner tool (FP3S200 V&P Scientific, Inc.) transferring 0.3 mL of drug stock solution prepared in DMSO. Plates were incubated for 1 hour at room-temperature. After 1 hour 182 mL of solution containing 50mM sodium azide (71289 – Sigma Aldrich) and 0.25% sodium dodecyl sulfate (SDS001.100 - BioShop) in M9 buffer using a multi-channel pipette followed by 182 mL of M9 buffer to raise the volume of each well such that a flat meniscus is produced. Plates were immediately imaged on the *2020 Imager* and were separated by 5 minutes to allow time for preparation of subsequent plates for imaging. After imaging egg-laying data was extracted from raw captured images using the 'Egg & Worm Counter' ImageJ plugin described above. Stimulators were considered molecules that stimulated egg-laying ≥ 2-fold greater than two proximal controls in the benign M9 buffer conditions with significance ($p < 0.05$ unpaired heteroscedastic t-test) over triplicate measurement (3 test compared against 6 control wells). Egl inhibitors were considered molecules that suppressed egg-laying ≤ 0.5 the normalized egg-laying rate of proximal controls in the 'nicotine (7.7 mM) + serotonin (12.3 mM)' condition described above with significance ($p < 0.05$ unpaired heteroscedastic t-test) over triplicate measurement (3 test compared against 6 control wells). Primary egg-laying modulators that reached the above threshold in at least 1 of 2 additional tests were considered bonafide 'Egl modulators'. As a more stringent criteria to narrow our focus on robust Egl modulators we limited our inquiry of Egl stimulators that induced Egl ≥ 2 fold that of control over 3 trials (primary screen + 2 retests) or stimulated Egl 3-fold relative to control over at least 2 of 3 trials and Egl inhibitors that suppressed Egl ≤ 2 fold control over all 3 re-tests or molecules that suppressed Egl ≤ 3 fold that of control in at least 2 of 3 trials.

Construction of the '2020 Imager'

High-content brightfield data were acquired on a custom brightfield 96 well-plate imager. The well-plate was mounted on a stationary platform, while the imaging setup travelled parallel to the bottom of the well-plate via a motorized stage. The plate was illuminated from above using a 10 W white LED. Underneath the plate, an Olympus 4x objective (Olympus, UPLFLN4XPH) was used with a 150 mm tube lens (Thorlabs, AC254-150-A-ML) to create an effective magnification of 3.33x. A mirror was used to maintain a low-profile imaging setup and minimize distortions induced by the displacement of the imaging optics. The magnified image was projected onto a 4K line scan camera (Dalsa, P2-23-04K40), resulting in an effective resolution of 3.00 $\mu$m/px. The camera had a single line of 4096 px which captured an area of 12.3mm by 3 $\mu$m with a bit depth of 10 bits. To capture a row of 12 wells, the image acquisition software (EPIX Inc, XCAP-Ltd) was setup to acquire scans at 3000 lines/s. This resulted in a 4K by 48K image that was acquired in about 15 seconds. To sequentially image each row in of a well-plate, an Arduino UNO was used to synchronize the image acquisition with the movement of the motorized stage. For additional details see Aaron Au's Master's thesis titled: 'Optical

Imaging Strategies for High-Content Studies of Development' available through the URL https://tspace.library.utoronto.ca/bitstream/1807/91538/3/Au_Aaron_K_201811_MAS_thesis.pdf.

ImageJ Analysis of *C. elegans* Egg-Laying Rate

A custom ImageJ (version 1.52i) plugin was used to quantify the number of worms and eggs present in each well (versions used available on github: github.com/seanph16/WormScanner3000/upload). Prior to object counting a threshold was applied based on the mean pixel intensity in each well image. The number of worms in a well was determined by measuring the area covered by non-egg shaped objects divided by the average area of a worm. Worm-like objects were recognized by the built-in ImageJ analyze particles function identifying objects greater than 11450 px in area with a circularity of 0.029–0.80 (the approximate minimum single adult worm size and range of shape circularities adopted by a worm) divided by the median worm area. Single eggs were counted by first creating a mask of egg and egg clump shaped objects (objects that are 300-6000 px in area with circularity of 0.15-1.00), applying the built-in ImageJ 'Watershed' function to recognize single eggs within clumps and counting objects with a circularity of 0.5–1 with a size of 300-2500 px. The number of egg objects divided by the number of worms was used as a read-out of egg-laying behaviour.

*C. elegans* Locomotory Survey

Locomotor phenotype analyses were done in 24-well plates with 1 mL of MYOB substrate (27.5 g Trizma HCl, 12 g Trizma Base, 230 g bacto tryptone, 10 g NaCl, 0.4 g cholesterol (95%)) seeded with 25 $\mu$L of OP50 *Escherichia coli* on each well. Each compound was added to the MYOB substrate before pouring to achieve the desired final concentrations of 30 $\mu$M or 60 $\mu$M after diffusion through the media. The final concentration of dimethyl sulfoxide (DMSO) in each of the wells was 1% v/v. Young adult or late fourth-staged larval worms are transferred into each well using a platinum wire pick. A Leica MZ75 stereomicroscope was used to visualize the movement of worms on the solid substrate. The specific locomotor phenotype (i.e. 'rubber-band' or 'coiler') was noted, and a qualitative assessment of the severity was made based on the degree of locomotor incapacitation and penetrance of the phenotype. Samples deemed 'Severe' indicated a strong perturbation and high penetrance, 'moderate' indicated a strong phenotype with low penetrance or a weak phenotype that is highly penetrant, and 'mild' indicated a weak phenotype that has low penetrance. Paralysis was distinguished from death by the presence of pharyngeal pumping.

*C. elegans Motor Phenotype Analyses:* The intensity of nementin-induced convulsions is tightly correlated with the degree of paralysis (i.e. animals that exhibit paralysis invariably convulse). We therefore used the degree of paralysis as a conveniently measurable proxy of convulsion intensity. In our survey of the effects of nementin analogs across *C. elegans, P. pacificus* and *R. diutinus*, animals were scored as convulsive if they failed to back at least ½ a body length after a touch on the head with a platinum wire. A more stringent scoring method was employed to compare convulsive phenotypes exhibited by *C. elegans* mutants. Animals were scored as convulsive if animals failed to demonstrate a sinusoidal wave form before or after a touch on the head with a platinum wire and failed to back ½ a body length. Like the convulsion scoring method above, animals treated with acetylcholinesterase inhibitors that failed to demonstrate a sinusoidal wave form before or after a touch on the head with a platinum wire that also fail to back ½ a body length were scored as paralyzed.

Locomotory Radiation Assay

~150 L4/young adult worms were pipetted in a 15 $\mu$L droplet onto the centre of a standard 10 cm round culture plate containing MYOB media + agar containing small-molecule with 1% DMSO. Media with drug were prepared in 50 mL falcon tube inverted 10x before pouring. Plates were dried for 90 minutes before being supplemented with a full lawn of OP50 bacteria seeded from a saturated culture of OP50 grown in LB broth. Plates were left uncovered adjacent to a flame until the (~15 minutes). 3 hours after pipetting worms onto plates the plates were flash frozen at -80°C for 3 minutes to freeze worms in place. The fraction of worms that travelled at least 1.65 cm (diameter after measuring 1cm from the edge of the droplet) from centre of the plate were recorded.

Developmental Growth Assay

*C. elegans* larval development assays were conducted in 96-well flatbottom clear flatbottom plates. ~20 L1 larvae in 10 µL of M9 buffer were pipetted into each test wells containing 40 µL of NGM media supplemented with HB101 *E. coli* with the desired test compound (+0.6% dimethylsulfoxide (DMSO; Sigma-Aldrich product ID: D8418) as the chemical solvent). Plates were wrapped in 3 layers of brown paper towels soaked with water. After either 3 or 6 days of incubation the number of *C. elegans* animals of different larval stages were recorded using a Leica MZ75 stereomicroscope.

*C. elegans* Confocal Microscopy

*C. elegans* KG4247 expressing *ceIs201* [unc-17p::ins-22::Venus + unc-17p::RFP + unc-17p::ssmCherry + myo-2p::RFP] were incubated on 6 cm MYOB media + 2% agar plates containing either 60 µM Nementin-1 or 60 µM Nementin-12 with 1% DMSO or 1% DMSO alone for 4 hours at room temperature. Wells were seeded with OP50 *E. coli* bacteria and used the day after preparation (see locomotory survey for further details on the preparation of media + drug plates). After incubation, animals were picked onto a 5% agar pad, 10 µL 10 mM tetramisole hydrochloride (prepared from 99% (–)-tetramisole hydrochloride, Sigma-Aldrich product ID: L9756) solvated in standard M9 buffer was pipetted onto the pad and a cover glass put on top. Animals were imaged using a Leica DMI 6000 B confocal microscope with a Hamatsu C9100-31 camera with a 100x oil immersion objective. A 491 nm laser was used to excite INS-22::Venus and images were captured with 25ms of exposure. A 510 nm laser was used to excite ssmCherry and RFP and images were captured with 100 ms of exposure. Images were captured after anterior and dorsal nerve cord features were brought into focus in the red channel (the RFP remained stable for the duration of imaging). Anterior, midbody and posterior regions containing respective coelomocytes (ccPR + ccAR, ccPL + ccAL & ccDL respectively) were captured. Images were captured over a 30 µm Z-stack captured with a 0.5 µm step and all images were captured within 25 minutes of slide preparation. A maximal projection containing the ventral and dorsal nerve cords and coelomocyte was generated for each captured section in ImageJ (version 1.52i). Tracings of captured axonal sections and coelomocytes were manually drawn and fluorescence signal measured in ImageJ. Due to the variability in coelomocyte endocytic/lysosomal vesicle content, coelomocytes were reported as the measurement of mean fluorescence signal of the GFP channel compared to the RFP channel. For axonal segments, the mean of two measurements of each region and representative background were collected to adjust for variability in manual measurement. Regions of interest for at least 15 animals were captured over several imaging sessions, at least 3 control animals were captured in each imaging session.

<u>Parasitic Nematode Assays</u>

*Cooperia oncophora Assay:* Fresh cattle feces containing eggs of an ivermectin-resistant strain of *C. oncophora* were kindly supplied by Dr. Doug Colwell and Dawn Gray (Lethbridge Research Station, Agriculture and Agri-Food Canada). Established methods were used to carry out the experimental cattle infections, and these methods were approved by the Lethbridge AAFC Animal Care committee and conducted under animal use license ACC1407. Cattle faeces containing C*. oncophora* eggs were stored anaerobically at room temperature for a maximum of 6 days before use. Eggs were isolated from faeces using a standard saturated salt flotation method immediately before the egg hatch assay. 80 $\mu$l of distilled and deionized water was added to each well of a 96-well culture plate, and then 1 $\mu$L of chemical at the appropriate concentration in DMSO was added to each well using a multichannel pipette. Approximately 50 eggs were added per well in 20 $\mu$L of water for a final volume of 100 $\mu$L in each well; the final DMSO concentration was 1% (v/v). The eggs were incubated in the chemicals for 2 days at room temperature, after which hatching was stopped by the addition of 1 $\mu$L iodine tincture to each well. The number of hatched larvae was counted at each concentration, and eggs that failed to hatch were scored as dead. "Relative viability" values were calculated by dividing the fraction of eggs that hatched at each concentration by the fraction of eggs that hatched in the corresponding DMSO control well. Two biological replicates were performed for each dose-response experiment, and the relative viability values were averaged across the biological replicates. The average hatch rate for the DMSO control wells was greater than 93% for both biological replicates.

*Dirofilaria immitis Assay:* Experiments on *D. immitis* microfilariae were performed in the laboratories of Bayer Animal Health GmbH (Monheim, Germany). The Missouri *D. immitis* isolate used for all assays was originally isolated from an infected dog from Missouri (USA). From 2005 onwards, the isolate was maintained and passaged in beagle dogs at the University of Georgia (Athens, GA, USA). From 2012 onward, the isolate was also maintained at the laboratories of Bayer Animal Health GmbH in Monheim, Germany. For the experiments with microfilariae, blood was sampled from beagle dogs (Marshall BioResources, North Rose, NY, USA) with patent infections, and microfilariae were purified according to the protocol described by the FR3.

Approximately 250 freshly purified microfilariae were cultured in single wells of a 96-well microtiter plate containing supplemented RPMI 1640 medium. Microfilariae exposed to medium substituted with 1% DMSO were used as negative controls. Motility of microfilariae was evaluated after 72 hours of drug exposure using an image-based approach – DiroImager, developed by Bayer Technology Services. This device is a fully automated high-throughput platform, allowing high-resolution optical imaging of an entire 96-well microtiter plate. The DiroImager integrates a high-resolution video camera (Prosilica GT6600; Allied Vision) with a telecentric lens (S5LPJ3005; Sill Optics) that prevents perspective distortion of the recorded images, ensuring high accuracy of measured values across all samples. In brief, a series of 20 high-resolution images were recorded (one per second). In a first step, image processing filters were used that discriminate larger objects to avoid the detection of crystallized or undissolved particles. In the actual calculation, pixel-wise differences between sequential images were calculated to determine worm movement between single images of a series; test compound activity was determined as the reduction of motility in comparison to the solvent control. Based on the evaluation of a wide concentration range, concentration–response curves as well as IC50 values were calculated were applicable.

*Nippostrongylus brasiliensis acetylcholine esterase secretion assay:* This assay has been previously described in detail (2). AChE is secreted by many parasitic nematodes, including *N. brasiliensis*. Assaying a small molecule's impact on AChE secretion is therefore a proxy for its ability to modulate the nematode's nervous system. Methods have been previously developed to assay AChE secretion from *Nippostrongylus* using colourimetric determination of AChE activity in the culture medium (3). Briefly, test compounds were dissolved in DMSO at a concentration of X, Y, Z and serial dilutions were performed in DMSO resulting in stock solutions of A, B, C. Stock solutions were stored at -20 °C until they were diluted 1:200 with culture medium (20 g/l Bacto Casitone, 10 g/l yeast extract, 5 g/l glucose, 0.8 g/l KH2PO4, 0.8 g/l K2HPO4, 10 µg/ml sisomycin and 1 µg/ml clotrimazole, pH 7.2). Final drug concentrations were E, F, G *µ*M in 1.0 % DMSO. Because secretion of AChE is gender and body weight specific two female and three male adult worms were placed in each well containing 1 ml of pre-warmed medium with drugs plus vehicle and incubated at 37 °C and 95% relative humidity for five days (4). All drug concentrations were performed in duplicate. From each well 25 µl medium were transferred into a 96 well plate. Then, 250 µl 5,5′-dithio-bis (2-nitrobenzoic acid) (0.25 µM) and 25 µl acetylthiocholine (4 mM) were added. AChE cleaves acetylthiocholine into acetate and thiocholine. In a consecutive reaction, thiocholine reacts with 5,5′-dithio-bis(2- nitrobenzoic acid) to thionitrobenzoate. Thionitrobenzoate is a yellow dye and its concentration can be determined by measuring the absorption at 405 nm. The A405 was measured after two and seven minutes of incubation at RT using an Expert 96 plate reader (Asys-Hitech, Salzburg,

Austria) and the software MikroWin 2000 (Mikrotek, Overrath, Germany). The difference in absorption between both time points was taken as measure of AChE activity. The arithmetic mean of 12 no drug control wells was set to 100% activity, and reduction of AChE activity in percentage relative to the negative control was calculated for each test compound concentrations. Within an assay, every drug concentration was performed in duplicate, and the software reported the mean of these duplicates.

*Strongyloides ratti L3 larvae lethality:* Data are the measurement of the % of Larval stage 3 (L3) worms (as indicated) that respond to 80°C hot water stimulus after 24 hours or 72 hours of incubation in wells containing the indicated compound. Data are the mean measurement of 30-40 larvae incubated in a dark box at room temperature for 24 or 72h over duplicate biological replicate conducted in triplicate.

*Trichuris muris L1 larvae experiments: T. muris* eggs were collected from the feces of the infected mice (as described above) using a flotation method with saturated NaCl solution in Milli-Q water. *T. muris* eggs were stored in Milli-Q water in the dark for 3 months at 23-25°C, until the eggs were embryonated. *T. muris* L1 were obtained using a hatching procedure with *E. coli (5)*. 30-40 larvae were placed in each well of a 96-well plate containing 175 μl culture medium and 25 μl of the test drug stock solutions. Larvae were kept at 37°C, 5% $CO_2$ for 24 hours. To evaluate the drug effect first the total number of L1 per well was determined. Then, 50-80 μl of hot water (≈80°C) was added to each well and the larvae that responded to this stimulus were counted. The proportion of larval death was determined. Larval survival counts were averaged over duplicate biological replicate conducted in triplicate normalized to controls.

*T. muris adult experiments:* Mice (C57BL/6NRj) were infected with 200 embryonated *T. muris* eggs. Seven weeks post-infection *T. muris* adult worms were collected from the intestines. Three worms were placed in each well of a 24-well plate containing 1980 µl culture medium and 20 µl of the test drugs (10 µM of a 1mM stock solution). After 72 hours of incubation at 37°C, 5% $CO_2$ the condition of the worms was microscopically evaluated using a viability scale from 3 (normal activity) to 0 (dead). Viability scores were averaged across replicates and normalized to the control wells. The experiment was conducted in duplicate.

*Heligmosomoides polygyrus L3 viability: H. polygyrus* infection three-week-old female NMRI mice were obtained from Charles River (Sulzfeld, Germany). Rodents were kept under environmentally-controlled conditions (temperature: 25°C, humidity: 70%, light/dark cycle 12 h /12 h) and had free access to water (municipal tap water supply) and rodent food and were allowed to acclimatize for one week. NMRI mice were infected with 88 *H. polygyrus* L3. Two weeks post-infection, mice were dissected cultivating the eggs on an agar plate for 8-10 days in the dark at 24°C. For the assays, 30-40 larvae were placed in each well of a 96-well plate containing 175 µl culture medium and 25 µl of the test drug stock solutions. *H. polygyrus* adults and stage 3 larvae (L3) were incubated in RPMI 1640 (Gibco, Waltham MA, USA) medium supplemented with 5% amphotericin B (250 µg/ml, Sigma-Aldrich, Buchs, Switzerland) and 1% penicillin 10,000 U/ml, and streptomycin 10 mg/ml solution (Sigma-Aldrich, Buchs, Switzerland). Culture plates were kept in a dark box at room temperature for up to 72 hours. To evaluate the drug effect first the total number of L3 per well was determined. Then, 50-80 $\mu$l of hot water (≈80°C) was added to each well and the larvae that responded to this stimulus were counted. The proportion of larval death was determined. Larval survival counts were averaged over duplicate biological replicate conducted in triplicate normalized to controls.

*Necator americanus L3 viability: N. americanus* larvae (L3) were obtained by filtering the feces of infected hamsters and cultivating the eggs on an agar plate for 8-10 days in the dark at 24°C. *Necator americanus* L3 were incubated in Hanks' balanced salt solution (HBSS; Gibco, Waltham MA, USA) supplemented with 10% amphotericin B and 1% penicillin (10,000 U/ml) and streptomycin (10 mg/ml) solution. For the assays, 30-40 larvae were placed in each well of a 96-well plate containing 175 $\mu$l culture medium and 25 $\mu$l of the test drug stock solutions. Treated Larvae were kept in a dark box at room temperature for up to 72 hours. To evaluate the drug effect first the total number L3 per well was determined. Then, 50-80 $\mu$l of hot water (≈80°C) was added to each well and the larvae that responded to this stimulus were counted. The proportion of larval death was determined. Larval survival counts were averaged over duplicate biological replicate conducted in triplicate normalized to controls.

*Meloidogyne incognita assays: M. incognita* infective second stage juvenile (J2) in vitro viability assays were performed in 96-well polystyrene plates. Each well contained approximately 25 J2s and compounds were added at a final concentration of 45 $\mu$M (0.5% DMSO v/v) in a total volume of 100 $\mu$L of sterile distilled water. Plates were sealed with parafilm and incubated for 72 hours at 25 °C. At the end point the fraction of viable nematodes in each drug condition and DMSO solvent controls was calculated by dividing the number of mobile nematodes by the total number of nematodes in the well. The experiment was conducted twice, with three technical replicates per treatment in each trial. *M. incognita* egg hatching assays were performed in sterile distilled water in 96-well plates similarly to the J2 viability assays described. Embryos were incubated in 45 $\mu$M (0.5% DMSO v/v) compound for 7 days at 25 °C. At the end point the number of hatched embryos was quantified in each condition and DMSO solvent controls. The fraction of hatched juveniles that were mobile was also quantified ('hatchling mobility'). The experiment was conducted twice, once with 50 embryos plated per well and once with 100 embryos plated per well, with three technical replicates per treatment in each trial. *M. incognita* 50-day soil reproduction assays were conducted in 90 grams of soil (1:1 sand:loam mix) per compartment in 6-pack planting containers. The soil was drenched with 18 mL of deionized water containing dissolved chemical or DMSO solvent alone. Approximately 1500 J2s were inoculated into the soil in 2 mL of water, for a total volume of 20 mL. The J2s were incubated in the soil and chemical for 24 hours after which a 2-3 week old tomato seedling was transplanted into the soil. Tomatoes were grown for 8 weeks in a greenhouse under long-day conditions (16 hour photoperiod) with 26/18 °C day/night temperatures. At the end point of the assay the tomato roots were harvested and eggs were extracted by rinsing in 0.6% sodium hypochlorite solution with agitation at 300 rpm for 3 minutes. Roots were rinsed with water over nested sieves and eggs present in each root system were collected and quantified. Roots were dried in a 65 °C oven and the number of eggs per milligram of dried root material was calculated. The experiment was conducted twice, with two technical replicates per treatment in each trial.

*Meloidogyne hapla motor assay: M. hapla* motor assays were conducted using J2 infective larvae isolated from ornamental tomato plant roots. J2s were isolated by isolating egg masses from the root network of infected plants and hatching in deionized water at room temperature for ~1 week. 10 µL of deionized water containing ~15 J2s (no fewer than 10 J2s) were pipetted into 96-well polystyrene plates containing the drug condition of interest with 0.6% DMSO. Addition of J2s to wells were staggered by 35 seconds for the purpose of maintaining a stringent endpoint. Drug dilutions were prepared from stock plates using a 96-well pinner tool (FP3S200 V&P Scientific, Inc.) transferring 0.3 mL of drug stock solution prepared in DMSO. Animals were incubated at room temperature with shaking for 4 hours (100 RPM; helps concentrate J2s in the middle of wells). At the 4 hour endpoint, 30 second videos of each well were captured using a Leica FLEXACAM C1 USB camera mounted to a Leica MZ75 stereomicroscope using Leica LAS EZ image capture software (V3.4.0). Videos were sped up 5x and the number of body bends generated over 30 seconds was recorded; animals were scored as paralyzed if they failed to generate more than 1 body bend over the 30 second recording. Data is reported as the mean % of paralyzed J2s over 3 or 4 independent biological replicates with ~15 animals per well (wells containing <10 animals were not scored). The dose-response matrix reporting the mean % of paralyzed J2s was used as the input for the SynergyFinder 2.0 server (https://synergyfinder.fimm.fi/).

Small-Molecule Tanimoto Coefficient Pairwise Similarity

Pairwise similarity scores were calculated as the Tanimoto coefficient of shared FP2 fingerprints using OpenBabel (http://openbabel.org). A Further description of Tanimoto pairwise similarity is provided in Burns et al. 2015 *(26)*. Network visualization for Fig. 1g was performed using Cytoscape (version 3.7.2).

Zebrafish Chemical Treatments and Phenotypic Analyses

All phenotypic analysis was performed on a stereomicroscope. At 1 dpf, 5 embryos were placed in 1 mL filter-sterilized egg water with chemicals in sterile 24-well plates (Falcon). At 3dpf, larvae were anaesthetized with ~0.6 mM tricaine methanesulfonate (tricaine), mounted in 3% methylcellulose on glass slides and bright-field images were taken with a 4x objective using a light microscope (Olympus BX43). The morphology of embryos relative to vehicle controls was assessed including their size, presence of edema, heart rate (normal, slow, or nearly absent), and presence of necrosis.

All chemicals were prepared in DMSO and added to filter-sterilized egg water at 0.1% of the final volume. Equal volumes of vehicle solvent were used in all conditions for a single assay. Note that methylene blue was not added to the egg water in any chemical assays. Culture plates were sealed with parafilm, wrapped in aluminum foil, and incubated at 28.5°C until the assay date.

A photoactivation assay was used to elicit movement and assess locomotion of zebrafish larvae as previously described (6). At 1 dpf, embryos in their chorions were aliquoted into 150 µL system water in 96-well plates (Falcon). Next, 50 µL of 4X chemical was added to each well to bring the volume to 200µL and 1X final concentration (either 3.75-60 µM). Plates were incubated until 3dpf, at which time any embryos still in their chorions were manually dechorionated in their wells. To assay locomotion, 10 µL of 210 µM optovin analog 6b8 (ChemDiv ID#2149-0111 or ChemBridge ID#5707191) was added to each well for a final concentration of 10 µM, incubated for 5 min, and movement tracked on the ZebraBox platform (ViewPoint) using a 30s lights on/off for 3m30s.

*Arabidopsis thaliana* Greening Assay

Greening experiments were performed with *Arabidopsis thaliana* seeds of wild type Col-0; seeds were surface sterilized in bleach and plated onto 0.5X MS, 0.5% sucrose agar medium supplemented with compounds of interest at 5, 15 and 45 µM concentrations (0.2% DMSO (v/v/)). After 4d of stratification at 4°C, plates were transferred to a growth chamber (16h / 8h, 150 $\mu E/m^2$) and greening recorded after 4 days. Pictures were recorded by camera (SONY a7s) with FE1.8/55 lens (FE 55 mm F1.8 ZA; SEL55F18Z). Experiments were performed in triplicate for each treatment.

*Drosophila melanogaster* Dose-Response Assay

Fly food in agar substrate was prepared by mixing 100 mL of unsulfured molasses, 100 mL of cornmeal, 41.2 g of Baker's yeast, and 14.8 g of agar into 1400 mL of distilled de-ionized water and boiling for 30 minutes. The media was allowed to cool to 56°C, at which point 5 mL was added by syringe to plastic cylindrical fly vials. 10 µL of chemical, or DMSO alone, was added to the media in each vial. The chemicals were mixed into the media by mechanical mixing using a pipette. The final DMSO concentration was 0.2% (v/v). The media was allowed to solidify at room temperature (~22°C) overnight. The following day (Day 0), eight pairs of male and female w1118 flies were added to each vial so that there were 16 flies in total per vial. The vials were stored at room temperature for 7 days, at which point the number of mobile flies was counted. Fly mobility was scored as any observable movement after the vial had been vigorously jostled. “Relative mobility” was calculated by dividing the number of mobile flies in the treatment vials by the average number of mobile flies in two DMSO control vials. On Day 8 the 16 parental flies were removed from the vials and the progeny larvae were allowed to continue to grow and hatch into adult flies. To assess larval viability, hatched flies were counted and discarded on Days 10, 12, 14, 16, 18, and 20. The counts were summed. “Relative viability” was calculated by dividing the number of hatched flies in the treatment vials by the average number of hatched flies in the two DMSO control vials. The final “relative mobility” and “relative viability” values are an average across three experimental replicates.

HEK293 Proliferation Assay

HEK293 cells were seeded into 96 well plates, at 5000 cells per well, in 100 μL total volumes of DMEM/10%FBS/1%PS media and grown overnight at 37˚C in the presence of 5% $CO_2$. Compounds (0.5 μL volumes from appropriate source plates) were then added to cells, and growth was continued for an additional 48 hours. Following growth,

10 μL of CellTiter-Blue Viability reagent (Promega) was added to each well, and plates were incubated for an additional 4 hours at 37˚C in the presence of 5% $CO_2$. Fluorescence measurements (560 nm excitation/590 nm emission) were then performed using a CLARIOstar Plate Reader (BMG Labtech) to quantify reagent reduction and estimate cell viability.

Spicule Protraction Assay

L4 males grown overnight on Nematode Growth Media (NGM) 2% agar plates + OP50. Next day adult males plated in liquid NGM liquid culture with 1% DMSO or 60 µM Nementin-1 with 1% DMSO in polystyrene 96-well plates. Adults were observed under a Leica MZ75 stereomicroscope at indicated time points. Reported '% Protracted' includes partial spicule protraction.

Statistical Analyses & Synergy Modelling

Unpaired one or two-sided t-tests or one-sided ANOVA with Dunnett's adjustment for multiple comparisons were conducted between control and treatment groups with where appropriate. Two-sided Chi-square tests with Bonferroni correction were conducted for comparison of proportional convulsion data to respective controls. Extra sum-of-squares F tests were conducted comparing EC50 curves generated for dose-response data. Statistical analyses were conducted using GraphPad Prism (version 9). Zero-Interaction

Potency (ZIP) synergy scores and heatmaps were generated using the SynergyFinder2.0 server using the default parameter set.

Chemistry

*General Considerations*

Unless otherwise stated, all reactions were set up under inert atmosphere (argon) utilizing glassware (or 2 dram vials) that were flame-dried and cooled under argon purging. Unless otherwise stated, flash column chromatography was performed on Silicycle® Siliaflash® P60, 40-63 µm silica gel. Starting materials and catalysts were purchased from commercial suppliers (Sigma Aldrich, Strem, Alfa Aesar, TCI or Combi-Blocks) and used without further purification unless otherwise stated. All solvents were distilled, purified, and dried according to standard procedures. Reactions were monitored using thin-layer chromatography (TLC) on EMD Silica Gel 60 F254 plates. Visualization of the developed plates was performed under UV light (254 nm) or by immersion in Ceric Ammonium Molybdate (CAM) or Potassium Permanganate ($KMnO_4$) stains.

**NMR** characterization data was collected at 296 K on a Varian Mercury 300, Varian Mercury 400, Bruker Avance III 400, Agilent DD2 500 (with cold probe), or an Agilent DD2 600 operating at 300, 400, 500, or 600 MHz for $^{1}H$ NMR, and 75, 100, 125, or 150 MHz for $^{13}C$ NMR. (Funded by the Canadian Foundation for Innovation, project number 19119, and the Ontario MRI). $^{1}H$ NMR spectra were internally referenced to the solvent residual signal ($CDCl_3$ = 7.26 ppm) unless otherwise stated. $^{13}C$ NMR spectra were internally referenced to the residual solvent signal (CDCl3 = 77.16 ppm) unless otherwise stated. $^{19}F$ NMR spectra were externally referenced to $CFCl_3$. Data for $^{1}H$ NMR are reported as follows: chemical shift (δ ppm), multiplicity (s = singlet, d = doublet, t = triplet, q = quartet, m = multiplet, br = broad), coupling constant (Hz), integration.

**Melting point (mp)** ranges were determined on a Fisher-Johns® Melting Point Apparatus and are reported uncorrected.

**Infrared (IR)** spectra were acquired using a Shimadzu FTIR-8400S FT-IR spectrometer as thin films ($CHCl_3$ or $CH_2Cl_2$) or neat on NaCl plates. Data is presented in wavenumbers ($v_{max}$, $cm^{-1}$).

**High Resolution Mass Spectra (HRMS)** were obtained on a micromass 70S-250 spectrometer (EI) or an AB SCIEX QSTAR® Mass Spectrometer (ESI) or a JEOL® AccuTOF medel JMS-T1000LC mass spectrometer equipped with an IONICS® Direct Analysis in real Time (DART) ion source at Advanced Instrumentation For Molecular Structure (AIMS) in the Department of Chemistry at the University of Toronto. Where ESI+ was employed, the values correspond to the ionic species of interest and the given ionic formula includes the charging agent ($H^+$ or $Na^+$); both the measured and calculated values are corrected for the mass of the electron and are reported as m/z. The DART-MS accurate mass report is generated using the *Elemental Composition Estimation* feature as implemented in the JEOL Mass Centre software package. Where DART was employed, the measured values correspond to the neutral species of interest and the given molecular formula includes the charging agent ($H^+$ or $NH_4^+$); the measured and calculated values are not corrected for the mass of the electron and are reported as neutral masses.

*General Procedures for the Synthesis*

*General Procedure A – Wittig Reaction*

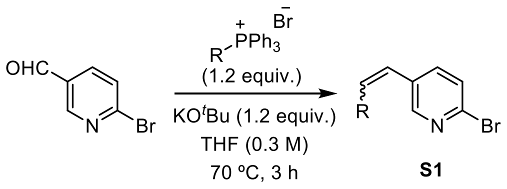


The phosphonium salt (1.2 equiv.) was first dissolved in THF (0.3 M). KO$^t$Bu (1.2 equiv.) was then added and the mixture was stirred at room temperature for 30 minutes. The pyridinecarboxaldehyde (1 equiv.) was then added in three portions and the reaction was refluxed at 70 °C for 3 hours. The mixture was then allowed to cool to room temperature and was filtered through a Celite pad eluting with pentanes. The filtrate was concentrated in vacuo and the product (mixture of isomers) was purified by flash column chromatography, eluting with a mixture of EtOAc:pentanes.

*General Procedure B – Suzuki Reaction (7)*

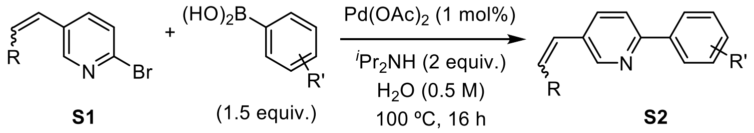


Substituted aryl boronic acid (1.5 equiv.) and $Pd(OAc)_2$ (1 mol%) were added to a mixture of the substituted bromopyridine (1 equiv.) in water (0.5 M). $^i Pr_2NH$ (2 equiv.) was then added and the reaction was refluxed at 100 °C for 16 hours. The mixture was allowed to cool to room temperature and brine was added. The aqueous phase was extracted with ethyl acetate. The combined organics was washed with brine, dried with $MgSO_4$, and concentrated in vacuo. The product was purified by flash column chromatography, eluting with a mixture of EtOAc:pentanes.

*General Procedure C – Hydrogenation*

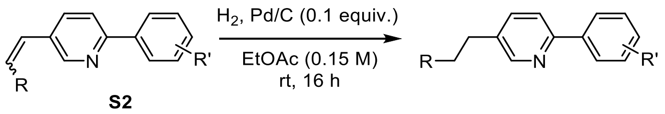


The vinylbiarene was dissolved in EtOAc (0.15 M). Pd/C (3% Pd, total 10 mol% Pd used) was added. Three cycles of evacuation and backfill with argon, followed by $H_2$ from a balloon was carried out. The reaction was stirred at room temperature under a $H_2$ atmosphere (balloon) for 16 hours. The contents of the flask were filtered over a Celite pad eluting with EtOAc and concentrated *in vacuo*. The product was purified by flash column chromatography, eluting with a mixture of EtOAc:pentanes.

**2-(4-((difluoromethyl)thio)phenyl)-5-propylpyridine [nementin-12-5]**

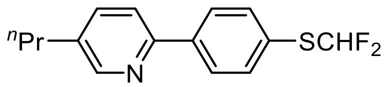


**nementin-12-5** was synthesized according to General Procedure A, B, and C using 6-bromonicotinaldehyde, ethyltriphenylphosphonium bromide, and (4-((difluoromethyl)thio)phenyl)boronic acid with an overall yield of 21% as a pale yellow oil. **$^1$H NMR** (500 MHz, $CDCl_3$) δ 8.53 (s, 1H), 8.04 – 7.95 (m, 2H), 7.69 – 7.62 (m, 3H), 7.60 – 7.53 (m, 1H), 6.86 (t, *J* = 56.9 Hz, 1H), 2.64 (d, *J* = 8.4 Hz, 2H), 1.76 – 1.64 (m, 2H), 0.98 (t, *J* = 7.3 Hz, 3H). **$^{13}$C NMR** (125 MHz, $CDCl_3$) δ 153.7, 150.0, 140.8, 137.0, 136.8, 135.4 (d, *J* = 1.2 Hz), 127.5,

126.3 (t, $J$ = 3.0 Hz), 121.0 (t, $J$ = 275.3 Hz), 120.2, 34.7, 24.2, 13.7. **$^{19}$F NMR** (375 MHz, $CDCl_3$) δ -91.2 (d, $J$ = 57.0 Hz). **IR** (thin film): 2962, 2932, 2871, 1554, 1472, 1319, 1068, 1036, 817 cm$^{-1}$. **HRMS** (DART-TOF+): mass [M+H] calc'd for $C_{15}H_{16}F_2NS$ 280.0972; found 280.0974.

**4-(difluoromethoxy)-4'-propyl-1,1'-biphenyl [nementin-12-6]**

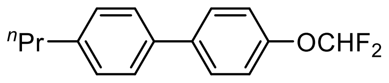


**nementin-12-6** was synthesized according to General Procedure A, B, and C using 4-bromobenzaldehyde, ethyltriphenylphosphonium bromide, and (4-(difluoromethoxy)phenyl)boronic acid with an overall yield of 48% as a white solid. **$^1$H NMR** (500 MHz, $CDCl_3$) δ 7.61 – 7.53 (m, 2H), 7.50 – 7.45 (m, 2H), 7.29 – 7.24 (m, 2H), 7.21 – 7.16 (m, 2H), 6.55 (t, $J$ = 74.0 Hz, 1H), 2.64 (t, $J$ = 8.5 Hz, 2H), 1.75 – 1.63 (m, 2H), 0.99 (t, $J$ = 7.3 Hz, 3H). **$^{13}$C NMR** (125 MHz, $CDCl_3$) δ 150.4 (t, $J$ = 2.8 Hz), 142.1, 138.6, 137.4, 129.0, 128.3, 126.8, 119.8, 116.0 (t, $J$ = 259.5 Hz), 37.7, 24.5, 13.9. **$^{19}$F NMR** (375 MHz, $CDCl_3$) δ -80.6 (d, $J$ = 74.5 Hz). **IR** (thin film): 2961, 2931, 1497, 1381, 1224, 1130, 1049, 909, 734 cm$^{-1}$. **HRMS** (DART-TOF+): mass [M] calc'd for $C_{16}H_{16}F_2O$ 262.1169; found 262.1162. **mp** 74-75 °C.

**5-(4-(difluoromethoxy)phenyl)-2-propylpyridine [nementin-12-7]**

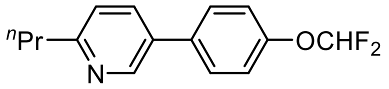


**nementin-12-7** was synthesized according to General Procedure A, B, and C using 5-bromopicolinaldehyde, ethyltriphenylphosphonium bromide, and (4-(difluoromethoxy)phenyl)boronic acid with an overall yield of 16% as a white solid. **$^1$H NMR** (500 MHz, $CDCl_3$) δ 8.75 – 8.68 (m,

1H), 7.77 – 7.72 (m, 1H), 7.59 – 7.53 (m, 2H), 7.25 – 7.19 (m, 3H), 6.56 (t, *J* = 73.7 Hz, 1H), 2.81 (t, *J* = 7.6 Hz, 2H), 1.87 – 1.72 (m, 2H), 1.00 (t, *J* = 7.4 Hz, 3H). **$^{13}$C NMR** (125 MHz, $CDCl_3$) δ 161.4, 150.9 (d, *J* = 3.2 Hz), 147.4, 135.3, 134.5, 132.8, 128.4, 122.7, 120.1, 115.8 (t, *J* = 260.3 Hz), 40.0, 23.1, 13.9. **$^{19}$F NMR** (375 MHz, $CDCl_3$) δ -80.9 (d, *J* = 73.3 Hz). **IR** (thin film): 2962, 1599, 1483, 1224, 1127,1046, 819 $cm^{-1}$. **HRMS** (DART-TOF+): mass [M+H] calc'd for $C_{15}H_{16}F_2ON$ 264.1200; found 264.1194. **mp** 47-48 °C.

**5-(difluoromethoxy)-2-(4-propylphenyl)pyridine [nementin-12-8]**

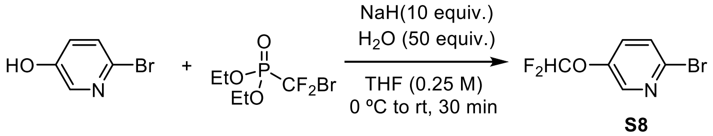


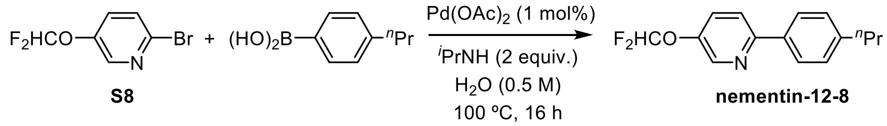


**2-bromo-5-(difluoromethoxy)pyridine (S8)**: The procedure was adapted from Wu and Wu *et al. (8)* 6-bromopyridin-3-ol (348 mg, 2 mmol, 1 equiv.) was dissolved in THF (8 mL, 0.25 M) and cooled to 0 °C. NaH (800 mg, 20 mmol, 60%, 10 equiv.) was added and the mixture was stirred at 0 °C for 30 minutes. $H_2O$ (1.80 mL, 100 mmol, 50 equiv.) was added dropwise and the mixture was stirred at 0 °C for 10 minutes. Diethyl (bromodifluoromethyl)phosphonate (0.71 mL, 4 mmol, 2 equiv.) was added and the reaction was allowed to stir from 0 °C to room temperature for 30 minutes. $H_2O$ was added and the aqueous phase was extracted with EtOAc. The combined organics was washed with brine, dried with $MgSO_4$, and concentrated in vacuo. The crude mixture was purified by flash column chromatography, eluting with 5% (v/v) EtOAc:pentanes to give S8 (161 mg, 0.72 mmol, 36%).

**5-(difluoromethoxy)-2-(4-propylphenyl)pyridine [nementin-12-8]:** Suzuki reaction was carried out to couple S8 (161 mg, 0.72 mmol, 1 equiv.) and (4-propylphenyl)boronic acid (177 mg, 1.08 mmol, 1.5 equiv.) according to General Procedure B to give **nementin-12-8** (140 mg, 0.53 mmol, 74%) as a white solid. **$^1$H NMR** (500 MHz, $CDCl_3$) δ 8.55 – 8.51 (m, 1H), 7.90 – 7.84 (m, 2H), 7.73 – 7.69 (m, 1H), 7.56 – 7.49 (m, 1H), 7.33 – 7.27 (m, 2H), 6.57 (t, *J* = 72.9 Hz, 1H), 2.65 (t, *J* = 7.3 Hz, 2H), 1.74 – 1.64 (m, 2H), 0.97 (t, *J* = 7.3 Hz, 3H). **$^{13}$C NMR** (125 MHz, $CDCl_3$) δ 154.9, 146.0 (t, *J* = 2.7 Hz), 143.9, 141.8, 135.8, 129.0, 128.2, 126.7, 120.6, 115.4 (t, *J* = 263.2 Hz), 37.8, 24.4, 13.8. **$^{19}$F NMR** (375 MHz, $CDCl_3$) δ -81.2 (d, *J* = 72.7 Hz). **IR** (thin film): 2961, 2930, 1471, 1379, 1215, 1118, 1047, 831 cm$^{-1}$. **HRMS** (DART-TOF+): mass [M+H] calc'd for $C_{15}H_{16}F_2ON$ 264.1200; found 264.1197. **mp** 25-26 °C.

**2-(difluoromethoxy)-5-(4-propylphenyl)pyridine [nementin-12-9]**

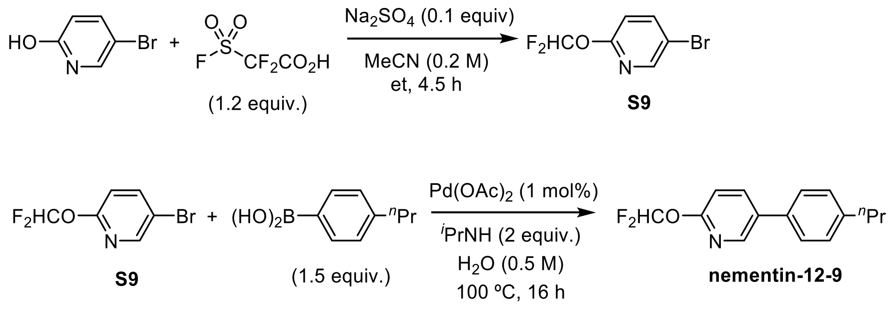

**5-bromo-2-(difluoromethoxy)pyridine (S9)**: The procedure was adapted from Ando *et al. (9).* 5-bromopyridin-2-ol (870 mg, 5 mmol, 1 equiv.) was dissolved in MeCN (25 mL, 0.2 M). 2,2-difluoro-2-(fluorosulfonyl)acetic acid (0.67 mL, 6 mmol, 1.2 equiv.) was added, followed by $Na_2SO_4$ (7 mg, 0.5 mmol, 0.1 equiv.). The reaction was quenched with saturated $NaHCO_{3(aq)}$ solution and extracted with EtOAC. The combined organics was washed with brine, dried with $MgSO_4$, and concentrated in vacuo. The crude mixture was purified by flash column chromatography, eluting with a 5 to 10% (v/v) EtOAc:pentanes gradient to give S9 (522 mg, 2.3 mmol, 47%).

**2-(difluoromethoxy)-5-(4-propylphenyl)pyridine (nementin-12-9)**: Suzuki reaction was carried out to couple S9 (112 mg, 0.5 mmol, 1 equiv.) and (4-propylphenyl)boronic acid (123 mg, 0.75 mmol, 1.5 equiv.) according to General Procedure B to give **nementin-12-9** (114 mg, 0.433 mmol, 87%) as an amorphous solid. **$^1$H NMR** (500 MHz, $CDCl_3$) δ 8.41 – 8.37 (m, 1H), 7.93 – 7.88 (m, 1H), 7.50 (t, *J* = 73.1 Hz, 1H), 7.46 – 7.42 (m, 2H), 7.31 – 7.27 (m, 2H), 6.96 (dd, *J* = 8.5, 0.7 Hz, 1H), 2.64 (t, *J* = 7.6 Hz, 2H), 1.74 – 1.62 (m, 2H), 0.98 (t, *J* = 7.3 Hz, 3H). **$^{13}$C NMR** (125 MHz, $CDCl_3$) δ 158.2 (t, *J* = 3.7 Hz), 144.9, 142.8, 138.6, 134.3, 133.6, 129.2, 126.7, 114.1 (t, *J* = 255.2 Hz), 111.6 – 111.0 (m), 37.7, 24.5, 13.8. **$^{19}$F NMR** (375 MHz, $CDCl_3$) δ -88.6 (d, *J* = 73.2 Hz). **IR** (thin film): 3362, 2962, 1601, 1478, 1347, 1258, 1129, 1103, 1072, 831 $cm^{-1}$. **HRMS** (DART-TOF+): mass [M+H] calc'd for $C_{15}H_{16}F_2ON$ 264.1200; found 264.1200.

**2-(4-(difluoromethoxy)phenyl)-5-ethylpyridine [nementin-12-10]**

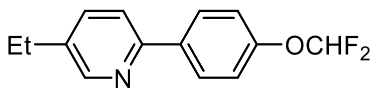

**nementin-12-10** was synthesized according to General Procedure A, B, and C using 6-bromonicotinaldehyde, methyltriphenylphosphonium bromide, and (4-(difluoromethoxy)phenyl)boronic acid with an overall yield of 4% as a white solid. **$^{1}$H NMR** (500 MHz, $CDCl_3$) δ 8.53 (s, 1H), 8.01 – 7.91 (m, 2H), 7.66 – 7.51 (m, 2H), 7.23 – 7.17 (m, 2H), 6.56 (t, $J$ = 73.9 Hz, 1H), 2.69 (q, $J$ = 7.6 Hz, 2H), 1.29 (t, $J$ = 7.6 Hz, 3H). **$^{13}$C NMR** (125 MHz, $CDCl_3$) δ 153.8, 151.6 (t, $J$ = 2.8 Hz), 149.5 – 149.3 (m), 137.9, 136.7, 136.2, 119.9, 119.5, 115.9 (td, $J$ = 259.3, 3.1 Hz), 25.8, 15.3. **$^{19}$F NMR** (375 MHz, $CDCl_3$) δ -80.8 (d, $J$ = 73.8 Hz). **IR** (thin film): 2967, 1478, 1381, 1221, 1129, 1052, 908, 733, 665 $cm^{-1}$. **HRMS** (DART-TOF+): mass [M+H] calc'd for $C_{14}H_{14}F_2ON$ 250.1044; found 250.1049. **mp** 32-33 °C.

**2-(4-(difluoromethoxy)phenyl)-5-methylpyridine [nementin-12-11]**

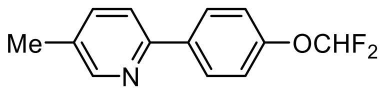

**nementin-12-11** was synthesized according to General Procedure B using 2-bromo-5-methylpyridine and (4-(difluoromethoxy)phenyl)boronic acid with a yield of 14% as a white solid. **$^{1}$H NMR** (500 MHz, $CDCl_3$) δ 8.51 (s, 1H), 8.00 – 7.95 (m, 2H), 7.64 – 7.50 (m, 2H), 7.23 – 7.18 (m, 2H), 6.56 (t, $J$ = 73.9 Hz, 1H), 2.37 (s, 3H). **$^{13}$C NMR** (125 MHz, $CDCl_3$) δ 153.6, 151.6 (t, $J$ = 2.8 Hz), 137.4, 136.7, 131.8, 128.1, 119.8, 119.5, 115.9 (t, $J$ = 259.7 Hz), 18.1. **$^{19}$F NMR** (375 MHz, $CDCl_3$) δ -80.8 (d, $J$ = 73.9 Hz). **IR** (thin film): 3004, 1605, 1477, 1220, 1123, 1-42, 823 $cm^{-1}$. **HRMS** (DART-TOF+): mass [M+H] calc'd for $C_{13}H_{12}F_2ON$ 236.0882; found 236.0884. **mp** 67-69 °C.

**2-(4-(difluoromethoxy)phenyl)pyridine [nementin-12-12]**

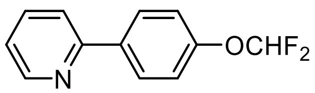

**nementin-12-12** was synthesized according to General Procedure B using 2-bromopyridine and (4-(difluoromethoxy)phenyl)boronic acid with a yield of 31% as a white solid. **$^1$H NMR** (500 MHz, $CDCl_3$) δ 8.71 – 8.63 (m, 1H), 8.04 – 7.98 (m, 2H), 7.79 – 7.73 (m, 1H), 7.72 – 7.68 (m, 1H), 7.26 – 7.19 (m, 3H), 6.57 (t, $J$ = 73.8 Hz, 1H). **$^{13}$C NMR** (125 MHz, $CDCl_3$) δ 156.3, 151.9, 149.7, 136.8, 136.6, 128.4, 122.2, 120.3, 119.5, 115.8 (td, $J$ = 260.0, 2.8 Hz). **$^{19}$F NMR** (375 MHz, $CDCl_3$) δ -80.8 (d, $J$ = 74.0 Hz). **IR** (thin film): 3368, 2924, 1592, 1468, 1437, 1382, 1222, 1178, 1043, 780 $cm^{-1}$. **HRMS** (DART-TOF+): mass [M+H] calc'd for $C_{15}H_{16}F_2ON$ 222.0731; found 222.0724. **mp** 35-36 °C.

**2-(4-(difluoromethoxy)phenyl)-5-(prop-1-yn-1-yl)pyridine [nementin-12-14]**

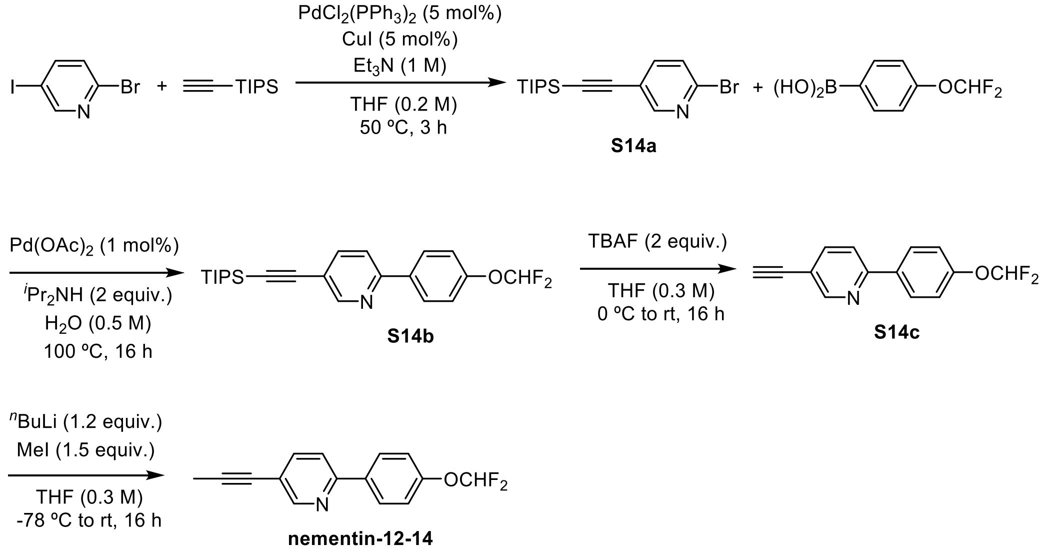

**2-bromo-5-((triisopropylsilyl)ethynyl)pyridine (S14a)**: 2-bromo-5-iodopyridine (283 mg, 1 mmol, 1 equiv.), $PdCl_2(PPh_3)_2$ (35 mg, 0.05 mmol, 5 mol%), and CuI (10 mg, 0.05 mmol, 5 mol%) were weighed into a flame-dried round bottom flask. The contents were purged under nitrogen for 5 minutes. THF (6.0 mL, 0.2 M) was then added to the flask. $Et_3N$ (1 mL, 1 M) and ethynyltriisopropylsilane (0.27 mL, 1.2 mmol, 1.2 equiv.) were added subsequently. The reaction was stirred at 50 °C for 3 hours. The mixture was cooled to room temperature and then filtered through celite, eluting with EtOAc. The filtrate was concentrated in vacuo and the crude mixture was purified by flash column chromatography, eluting with 10% (v/v) EtOAc:pentanes to give S14a (237 mg, 0.70 mmol, 70%)

**2-(4-(difluoromethoxy)phenyl)-5-((triisopropylsilyl)ethynyl)pyridine (S14b):** Suzuki reaction was carried out to couple S14a (237 mg, 0.70 mmol, 1 equiv.) and (4-(difluoromethoxy)phenyl)boronic acid (197 mg, 1.05 mmol, 1.5 equiv.) according to General Procedure B to give S14b (162 mg, 0.40 mmol, 58%).

**2-(4-(difluoromethoxy)phenyl)-5-ethynylpyridine (S14c):** S14b (162 mg, 0.4 mmol, 1 equiv.) was dissolved in THF (1.3 mL, 0.3 M) and cooled to 0 °C. TBAF (0.8 mL, 0.8 mmol, 1 M in THF, 2 equiv.) was added dropwise. The reaction was allowed to stir from 0 °C to room temperature for 16 hours. $H_2O$ was added and the aqueous phase was extracted with EtOAc. The combined organics was washed with brine, dried with $MgSO_4$, and concentrated in vacuo. The crude mixture was purified by flash column chromatography, eluting with a 5 to 10% (v/v) EtOAc:pentanes gradient to give S14c (77 mg, 0.31 mmol, 78%).

**2-(4-(difluoromethoxy)phenyl)-5-(prop-1-yn-1-yl)pyridine (nementin-12-14):** S14c (77 mg, 0.31 mmol, 1 equiv.) was dissolved in THF (1.0 mL, 0.3 M) and cooled to -78 °C. $^{n}$BuLi (0.15 mL, 0.37 mmol, 2.5 M in hexane, 1.2 equiv.) was added dropwise. The mixture was stirred at -78 °C for 1 hour. Iodomethane (0.03 mL, 0.47 mmol, 1.5 equiv.) was added dropwise. The reaction was allowed to stir from -78 °C to room temperature for 16 hours. The reaction was quenched with saturated $NH_4Cl_{(aq)}$ solution and extracted with EtOAc. The combined organics was washed with brine, dried with $MgSO_4$, and concentrated in vacuo. The crude mixture was purified by flash column chromatography, eluting with a 2.5 to 5% (v/v) EtOAc:pentanes gradient to give **nementin-12-14** (28 mg, 0.11 mmol, 35%) as a white solid. **$^{1}$H NMR** (500 MHz, $CDCl_3$) δ 8.68 – 8.67 (m, 1H), 8.02 – 7.96 (m, 2H), 7.75 – 7.70 (m, 1H), 7.65 – 7.59 (m, 1H), 7.24 – 7.17 (m, 2H), 6.56 (t, *J* = 73.8 Hz, 1H), 2.10 (s, 3H). **$^{13}$C NMR** (125 MHz, $CDCl_3$) δ 154.3, 152.2, 152.0 (t, *J* = 2.8 Hz), 139.2, 136.0, 119.6, 119.5, 119.3, 115.8 (t, *J* = 260.0 Hz), 90.0, 76.6, 4.5. **$^{19}$F NMR** (375 MHz, $CDCl_3$) δ -80.9 (d, *J* = 72.9 Hz). **IR** (thin film): 2932, 2256, 2221, 1588, 1475, 1378, 1230, 1129, 1041, 828 $cm^{-1}$. **HRMS** (DART-TOF+): mass [M+H] calc'd for $C_{15}H_{12}F_2ON$ 260.0887; found 260.0879. **mp** 68-70 °C.

### (6-(4-(difluoromethoxy)phenyl)pyridin-3-yl)methanamine [nementin-12-17]

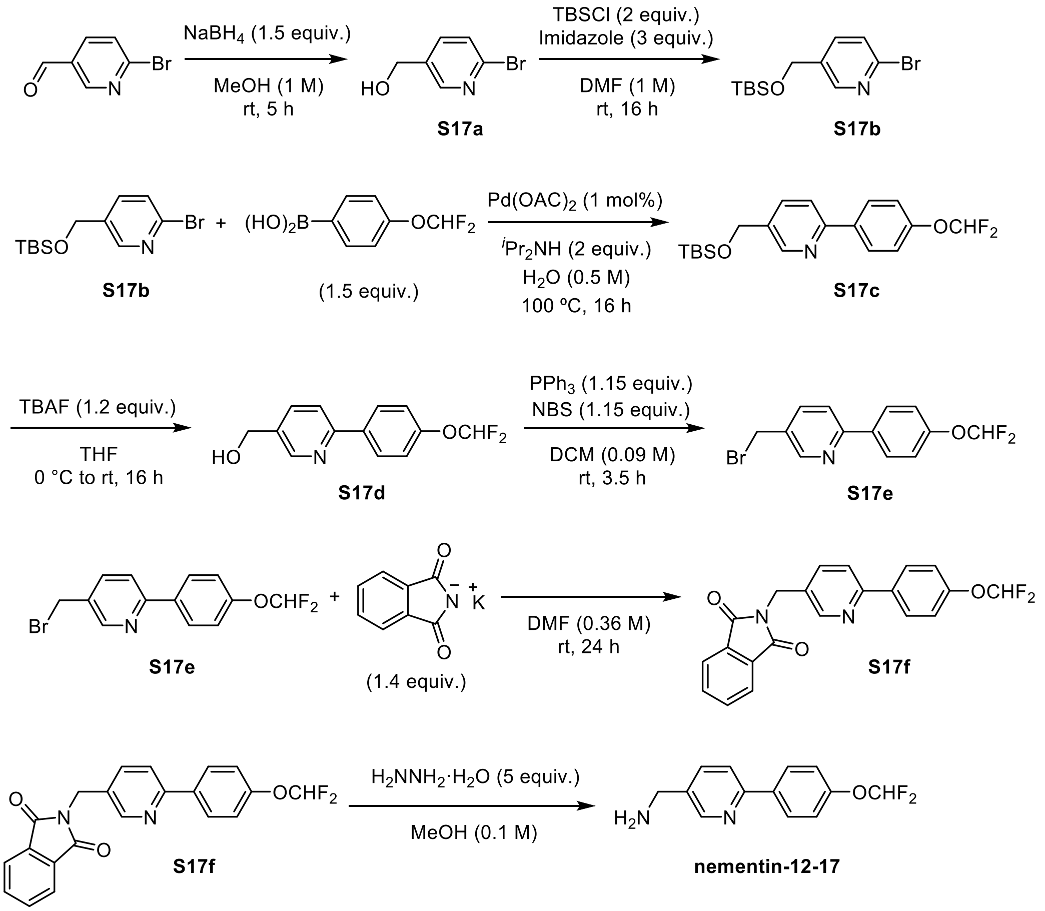


**(6-bromopyridin-3-yl)methanol (S17a)**: 6-bromonicotinaldehyde (1.302 g, 7 mmol, 1 equiv.) was dissolved in MeOH (7 mL, 1 M). $NaBH_4$ (397 mg, 10.5 mmol, 1.5 equiv.) was added in three portions. The reaction was stirred at room temperature for 5 hours. The mixture was then cooled to 0 °C and 1 M HCl was added dropwise until bubbling seized, followed with dilution of the mixture with $H_2O$. The aqueous phase was extracted with EtOAc. The combined organics was washed with brine, dried with $MgSO_4$, and concentrated in vacuo. The crude mixture was purified by flash column chromatography, eluting with a 50 to 60% (v/v) EtOAc:pentanes gradient to give S17a (1.286 g, 6.84 mmol, 98%).

**2-bromo-5-(((tert-butyldimethylsilyl)oxy)methyl)pyridine (S17b)**: S17a (884 mg, 4.7 mmol, 1 equiv.) was dissolved in DMF (4.7 mL, 1 M) and cooled to 0 °C. TBSCl (1.416 g, 9.4 mmol, 2 equiv.) was added followed by imidazole (960 mg, 14.1 mmol, 3 equiv.). The reaction was allowed to stir from 0 °C to room temperature for 16 hours. $H_2O$ was added to the mixture and the aqueous phase was extracted with EtOAc. The combined organics was washed with brine, dried with $MgSO_4$, and concentrated in vacuo. The crude mixture was purified by flash column chromatography, eluting with a 1 to 2% (v/v) EtOAc:pentanes gradient to give S17b (1.095 g, 3.63 mmol, 77%).

**5-(((tert-butyldimethylsilyl)oxy)methyl)-2-(4-(difluoromethoxy)phenyl)pyridine (S17c)**: Suzuki reaction was carried out to couple S17b (1.090 g, 3.63 mmol, 1 equiv.) and (4-(difluoromethoxy)phenyl)boronic acid (1.020 g, 5.45 mmol, 1.5 equiv.) according to General Procedure B to give S17c (752 mg, 1.98 mmol, 54%).

**(6-(4-(difluoromethoxy)phenyl)pyridin-3-yl)methanol (S17d)**: **S17c** (752 mg, 1.98 mmol, 1 equiv.) was dissolved in THF (6.6 mL, 0.3 M) and cooled to 0 °C. TBAF (2.38 mL, 2.38 mmol, 1 M in THF, 1.2 equiv.) was added dropwise. The reaction was allowed to stir from 0 °C to room temperature for 16 hours. $H_2O$ was added to the mixture and the aqueous phase was extracted with EtOAc. The combined organics was washed with brine, dried with $MgSO_4$, and concentrated in vacuo. The crude mixture was purified by flash column chromatography, eluting with a 50 to 60% (v/v) EtOAc:pentanes gradient to give S17d (363 mg, 1.45 mmol, 73%).

**5-(bromomethyl)-2-(4-(difluoromethoxy)phenyl)pyridine (S17e):** S17d (363 mg, 1.45 mmol, 1 equiv.) was dissolved in DCM (16.11 mL, 0.09 M). $PPh_3$ (438 mg, 1.67 mmol, 1.15 equiv.) followed by NBS (297 mg, 1.67 mmol, 1.15 equiv.) were added. The reaction was stirred at room temperature for 3.5 hours. $H_2O$ was added to the mixture and the aqueous phase was extracted with EtOAc. The combined organics was washed with brine, dried with $MgSO_4$, and concentrated in vacuo. The crude mixture was purified by flash column chromatography, eluting with a 15% (v/v) EtOAc:pentanes to give S17e (350 mg, 1.11 mmol, 77%).

**2-((6-(4-(difluoromethoxy)phenyl)pyridin-3-yl)methyl)isoindoline-1,3-dione (S17f):** S17e (349 mg, 1.11 mmol, 1 equiv.) was dissolved in DMF (3.1 mL, 0.36 M). Phthalimide potassium salt (438 mg, 1.67 mmol, 1.15 equiv.) was added. The reaction was stirred at room temperature for 24 hours. $H_2O$ was added to the mixture and the aqueous phase was extracted with EtOAc. The combined organics was washed with brine, dried with $MgSO_4$, and concentrated in vacuo. The crude mixture was purified by flash column chromatography, eluting with a 20 to 25% (v/v) EtOAc:pentanes gradient to give S17f (262 mg, 0.69 mmol, 62%).

**(6-(4-(difluoromethoxy)phenyl)pyridin-3-yl)methanamine (nementin-12-17)**: S17f (262 mg, 0.69 mmol, 1 equiv.) was dissolved in MeOH (6.9 mL, 0.1 M). Hydrazine hydrate (0.26 mL, 3.56 mmol, 50-60%, 5 equiv.) was added. The reaction was refluxed at 70 °C for 4 hours. The mixture was cooled to room temperature and filtered through Celite, eluting with MeOH. The filtrate was concentrated in vacuo. $H_2O$ (7 mL) was added, followed by 1 M KOH (1.75 mL). The aqueous phase was extracted with DCM. The combined organics was dried with $MgSO_4$ and then concentrated in vacuo to give **nementin-12-17** (131 mg, 0.53 mmol, 76%) as a white solid. **$^1$H NMR** (500 MHz, $CDCl_3$) δ 8.64 – 8.58 (m, 1H), 8.01 – 7.96 (m, 2H), 7.79 – 7.71 (m, 1H), 7.71 – 7.64 (m, 1H), 7.23 – 7.18 (m, 2H), 6.56 (t, $J$ = 73.8 Hz, 1H), 3.95 (s, 2H), 1.52 (br, 2H). **$^{13}$C NMR** (125 MHz, $CDCl_3$) δ 155.0, 151.8 (t, $J$ = 2.8 Hz), 148.8, 136.8, 136.5, 135.8, 128.3, 120.1, 119.5, 115.8 (t, $J$ = 259.9 Hz), 43.7. **$^{19}$F NMR** (375 MHz, $CDCl_3$) δ -80.8 (d, $J$ = 73.3 Hz). **IR** (thin film): 3265, 1591, 1478, 1381, 1231, 1125, 1037, 783 $cm^{-1}$. **HRMS** (DART-TOF+): mass [M+H] calc'd for $C_{13}H_{13}F_2ON_2$ 251.0991; found 251.0988. **mp** 41-43 °C.

**2-(4-(difluoromethoxy)phenyl)-6-propylpyridine [nementin-12-20]**

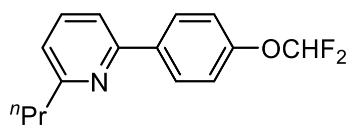


**nementin-12-20** was synthesized according to General Procedure A, B, and C using 6-bromopicolinaldehyde, ethyltriphenylphosphonium bromide, and (4-(difluoromethoxy)phenyl)boronic acid with an overall yield of 18% as a pale yellow oil. **$^1$H NMR** (500 MHz, $CDCl_3$) δ 8.04 – 7.99 (m, 2H), 7.68 – 7.62 (m, 1H), 7.51 – 7.47 (m, 1H), 7.24 – 7.15 (m, 2H), 7.11 – 7.06 (m, 1H), 6.55 (t, $J$ = 73.9 Hz, 1H), 2.83 (t, $J$ = 7.9 Hz, 2H), 1.87 – 1.77 (m, 2H), 1.01 (t, $J$ = 7.4 Hz, 3H). **$^{13}$C NMR** (125 MHz, $CDCl_3$) δ 162.3, 155.6, 151.7 (t, $J$ = 2.8 Hz), 137.2, 136.8, 128.5, 121.2, 119.5, 117.4, 115.9 (dd, $J$ = 260.1, 3.2 Hz), 40.5, 22.9, 13.9. **$^{19}$F NMR** (375 MHz, $CDCl_3$) δ -80.7 (d, $J$ = 74.3 Hz). **IR** (thin film): $cm^{-1}$. **HRMS** (DART-TOF+): mass [M+H] calc'd for $C_{15}H_{16}F_2ON$ 264.1200; found 264.1197.

**2-(4-(difluoromethoxy)phenyl)-4-propylpyridine [nementin-12-21]**

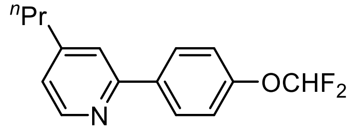


**nementin-12-21(G)** was synthesized according to General Procedure A, B, and C using 2-bromoisonicotinaldehyde, ethyltriphenylphosphonium bromide, and (4-(difluoromethoxy)phenyl)boronic acid with an overall yield of 9% as an amorphous solid. **$^1$H NMR** (500 MHz, $CDCl_3$) δ 8.55 (d, *J* = 5.0 Hz, 1H), 8.01 – 7.95 (m, 2H), 7.51 – 7.48 (m, 1H), 7.23 – 7.16 (m, 2H), 7.07 – 7.03 (m, 1H), 6.56 (t, *J* = 73.9 Hz, 1H), 2.64 (t, *J* = 7.7 Hz, 2H), 1.77 – 1.64 (m, 2H), 0.98 (t, *J* = 7.3 Hz, 3H). **$^{13}$C NMR** (125 MHz, $CDCl_3$) δ 156.2, 152.4, 151.8 (t, *J* = 2.8 Hz), 149.6 – 149.4 (m), 136.9, 128.5 (d, *J* = 1.2 Hz), 122.6, 120.6 (t, *J* = 2.1 Hz), 119.4, 115.9 (td, *J* = 260.0, 3.8 Hz), 37.5, 23.6, 13.7. **$^{19}$F NMR** (375 MHz, $CDCl_3$) δ -80.8 (d, *J* = 73.3 Hz). **IR** (thin film): 2961, 1605, 1222, 1130, 1050, 909, 733 $cm^{-1}$. **HRMS** (DART-TOF+): mass [M+H] calc'd for $C_{15}H_{16}F_2ON$ 264.1200; found 264.1205.

**2-(4-(difluoromethoxy)phenyl)-3-propylpyridine [nementin-12-22]**

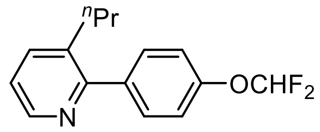


**nementin-12-22** was synthesized according to General Procedure A, B, and C using 2-bromonicotinaldehyde, ethyltriphenylphosphonium bromide, and (4-(difluoromethoxy)phenyl)boronic acid with an overall yield of 6% as a pale yellow oil. **$^1$H NMR** (500 MHz, $CDCl_3$) δ 8.53 – 8.48 (m, 1H), 7.64 – 7.58 (m, 1H), 7.51 – 7.46 (m, 2H), 7.24 – 7.16 (m, 3H), 6.56 (t, *J* = 73.9 Hz, 1H), 2.61 (t, *J* = 7.7 Hz, 2H), 1.60 – 1.46 (m, 2H), 0.86 (t, *J* = 7.3 Hz, 3H). **$^{13}$C NMR** (125 MHz, $CDCl_3$) δ 157.6, 150.9 (t, *J* = 2.8 Hz), 146.9 – 146.7 (m), 138.0, 137.4, 135.5, 130.4, 122.4, 119.1, 116.0 (td, *J* = 259.1, 2.6 Hz), 34.4, 24.0, 13.9. **$^{19}$F NMR** (375 MHz,

CDCl3) δ -80.7 (d, *J* = 73.9 Hz). **IR** (thin film): 3367, 2962, 2873, 1610, 1511, 1435, 1382, 1223, 1179, 1127, 1045, 843, 783, 665 $cm^{-1}$. **HRMS** (DART-TOF+): mass [M+H] calc'd for $C_{15}H_{16}F_2ON$ 264.1200; found 264.1193.

**2-(3-(difluoromethoxy)phenyl)-5-propylpyridine [nementin-12-23]**

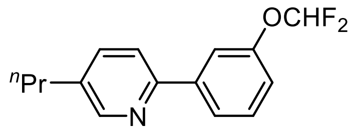


**nementin-12-23** was synthesized according to General Procedure A, B, and C using 6-bromonicotinaldehyde, ethyltriphenylphosphonium bromide, and (3-(difluoromethoxy)phenyl)boronic acid with an overall yield of 21% as a pale yellow oil. **$^1$H NMR** (500 MHz, $CDCl_3$) δ 8.53 – 8.51 (m, 1H), 7.82 – 7.79 (m, 1H), 7.79 – 7.77 (m, 1H), 7.66 – 7.61 (m, 1H), 7.59 – 7.55 (m, 1H), 7.48 – 7.42 (m, 1H), 7.16 – 7.12 (m, 1H), 6.59 (t, *J* = 74.0 Hz, 1H), 2.64 (t, *J* = 7.5 Hz, 2H), 1.75 – 1.64 (m, 2H), 0.98 (t, *J* = 7.3 Hz, 3H). **$^{13}$C NMR** (125 MHz, $CDCl_3$) δ 153.6, 151.8 (t, *J* = 2.8 Hz), 150.0 – 149.8 (m), 141.5, 136.9, 136.8, 130.0, 123.5, 120.1, 119.5, 117.7, 116.1 (td, *J* = 259.2, 3.1 Hz), 34.7, 24.2, 13.7. **$^{19}$F NMR** (375 MHz, $CDCl_3$) δ -80.5 (d, *J* = 73.9 Hz). **IR** (thin film): 2962, 1593, 1564, 1470, 1382, 1197, 1045, 792 $cm^{-1}$. **HRMS** (DART-TOF+): mass [M+H] calc'd for $C_{15}H_{16}F_2ON$ 264.1200; found 264.1197.

**2-(2-(difluoromethoxy)phenyl)-5-propylpyridine (nementin-12-24)**

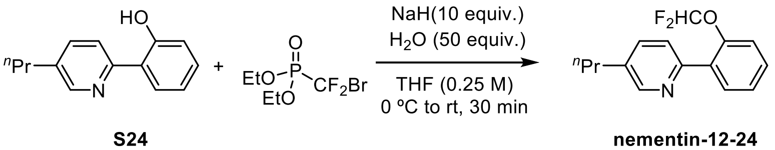


**S24** **nementin-12-24**

**2-(5-propylpyridin-2-yl)phenol (S24)**: **S24** was synthesized according to General Procedure A, B, and C using 6-bromonicotinaldehyde, ethyltriphenylphosphonium bromide, and (2-(benzyloxy)phenyl)boronic acid with an overall yield of 15%.

**2-(2-(difluoromethoxy)phenyl)-5-propylpyridine (nementin-12-24)**: **S24** (43 mg, 0.2 mmol, 1 equiv.) was dissolved in THF (0.80 mL, 0.25 M) and cooled to 0 °C. NaH (80 mg, 10 mmol, 60%, 10 equiv.) was added and the mixture was stirred at 0 °C for 30 minutes. $H_2O$ (0.18 mL, 10 mmol, 50 equiv.) was added dropwise and the mixture was stirred at 0 °C for 10 minutes. Diethyl (bromodifluoromethyl)phosphonate (0.07 mL, 0.4 mmol, 2 equiv.) was added and the reaction was allowed to stir from 0 °C to room temperature for 30 minutes. $H_2O$ was added and the aqueous phase was extracted with EtOAc. The combined organics was washed with brine, dried with $MgSO_4$, and concentrated in vacuo. The crude mixture was purified by flash column chromatography, eluting with 5% (v/v) EtOAc:pentanes to give **nementin-12-24** (9 mg, 0.032 mmol, 16%) as a pale yellow oil. **$^1$H NMR** (500 MHz, $CDCl_3$) δ 8.55 – 8.49 (m, 1H), 7.82 – 7.74 (m, 1H), 7.66 – 7.61 (m, 1H), 7.58 – 7.54 (m, 1H), 7.42 – 7.36 (m, 1H), 7.37 – 7.29 (m, 1H), 7.25 – 7.20 (m, 1H), 6.48 (t, $J$ = 74.6 Hz, 1H), 2.64 (t, $J$ = 7.5 Hz, 2H), 1.75 – 1.66 (m, 2H), 0.99 (t, $J$ = 7.3 Hz, 3H). **$^{13}$C NMR** (125 MHz, $CDCl_3$) δ 152.2, 149.7, 148.6 (t, $J$ = 2.8 Hz), 136.5, 136.0, 132.8, 131.5, 129.7, 126.0, 124.3, 120.3, 116.7 (t, $J$ = 259.1 Hz), 34.8, 24.2, 13.7. **$^{19}$F NMR** (375 MHz, $CDCl_3$) δ -80.4 (d, $J$ = 74.4 Hz). **IR** (thin film): 2966, 2933, 1600, 1472, 1383, 1213, 1127, 1047, 1027, 758 $cm^{-1}$. **HRMS** (DART-TOF+): mass [M+H] calc'd for $C_{15}H_{16}F_2ON$ 264.1195; found 264.1195.

**2-(4-(difluoromethoxy)phenyl)-5-propylpyrimidine [nementin-12-25]**

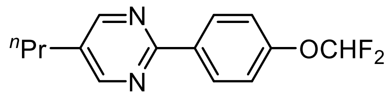

Pd(OAc)$_2$ (3 mg, 0.015 mmol, 5 mol%), PPh$_3$ (8.0 mg, 0.03 mmol, 10 mol%), K$_2$CO$_3$ (124.4 mg, 0.9 mmol, 3 equiv.) and (4-(difluoromethoxy)phenyl)boronic acid (85 mg, 0.45 mmol, 1.5 equiv.) were added to a flamed-dried flask. The mixture was purged with nitrogen for 5 minutes. Dioxane (0.55 mL, 0.55 M) and $H_2O$ (0.14 mL, 2.18 M) were added, followed by 2-chloro-5-propylpyrimidine (0.04 mL, 0.3 mmol, 1 equiv.). The reaction was then refluxed at 100 °C for 16 hours. $H_2O$ was added to the mixture and the aqueous phase was extracted with EtOAc. The combined organics was washed with brine, dried with $MgSO_4$, and concentrated in vacuo. The crude mixture was purified by flash column chromatography, eluting with a 2.5 to 5% (v/v) EtOAc:pentanes gradient to give **nementin-12-25** (69 mg, 0.26 mmol, 86%) as a white solid. **$^1$H NMR** (500 MHz, CDCl$_3$) δ 8.61 (s, 2H), 8.46 – 8.41 (m, 2H), 7.23 – 7.18 (m, 2H), 6.59 (t, *J* = 73.8 Hz, 1H), 2.61 (t, *J* = 7.5 Hz, 2H), 1.75 – 1.65 (m, 2H), 1.00 (t, *J* = 7.4 Hz, 3H). **$^{13}$C NMR** (125 MHz, CDCl$_3$) δ 161.7, 157.1, 153.0 (t, *J* = 2.7 Hz), 134.8, 132.8, 129.6, 119.0, 115.8 (t, *J* = 259.6 Hz), 32.2, 24.0, 13.6. **$^{19}$F NMR** (375 MHz, CDCl$_3$) δ -81.0 (d, *J* = 73.2 Hz). **IR** (thin film): 2963, 1588, 1431, 1383, 1220, 1126, 1048, 799 cm$^{-1}$. **HRMS** (DART-TOF+): mass [M+H] calc'd for $C_{14}H_{15}F_2ON_2$ 265.1147; found 265.1148. **mp** 37-39 °C.

**2-(4-(difluoromethoxy)phenyl)-5-isobutylpyridine [nementin-12-26]**

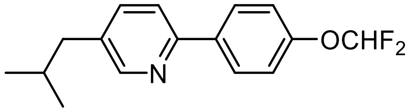

**nementin-12-26** was synthesized according to General Procedure A, B, and C using 6-bromonicotinaldehyde, isopropyltriphenylphosphonium iodide, and (4-(difluoromethoxy)phenyl)boronic acid with an overall yield of 23% as a white solid. **$^1$H NMR** (500 MHz, $CDCl_3$) δ 8.49 – 8.45 (m, 1H), 8.03 – 7.95 (m, 2H), 7.64 – 7.59 (m, 1H), 7.56 – 7.51 (m, 1H), 7.23 – 7.17 (m, 2H), 6.56 (t, $J$ = 73.9 Hz, 1H), 2.52 (d, $J$ = 7.1 Hz, 2H), 1.95 – 1.85 (m, 1H), 0.95 (d, $J$ = 6.6 Hz, 7H). **$^{13}$C NMR** (125 MHz, $CDCl_3$) δ 153.8, 151.6 (t, $J$ = 2.9 Hz), 150.4, 137.3, 136.7, 135.4, 128.2, 119.7, 119.5, 115.9 (t, $J$ = 259.7 Hz), 42.0, 30.0, 22.2. **$^{19}$F NMR** (375 MHz, $CDCl_3$) δ -80.8 (d, $J$ = 73.3 Hz). **IR** (thin film): 2959, 1598, 1511, 1476, 1383, 1227, 1125, 1045, 836, 804 cm$^{-1}$. **HRMS** (DART-TOF+): mass [M+H] calc'd for $C_{16}H_{18}F_2ON$ 278.1351; found 278.1351. **mp** 29-30 °C.

**5-butyl-2-(4-(difluoromethoxy)phenyl)pyridine [nementin-12-27]**

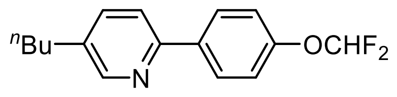


**nementin-12-27** was synthesized according to General Procedure A, B, and C using 6-bromonicotinaldehyde, propyltriphenylphosphonium bromide, and (4-(difluoromethoxy)phenyl)boronic acid with an overall yield of 32% as an amorphous solid. **$^1$H NMR** (500 MHz, $CDCl_3$) δ 8.59 – 8.45 (m, 1H), 8.03 – 7.91 (m, 2H), 7.66 – 7.47 (m, 2H), 7.24 – 7.15 (m, 2H), 6.56 (t, $J$ = 73.9 Hz, 1H), 2.65 (t, $J$ = 7.7 Hz, 2H), 1.70 – 1.56 (m, 3H), 1.46 – 1.32 (m, 2H), 0.95 (t, $J$ = 7.3 Hz, 4H). **$^{13}$C NMR** (125 MHz, $CDCl_3$) δ 153.8, 151.6 (t, $J$ = 3.0 Hz), 149.8, 136.7, 136.7, 136.6, 128.2, 119.8, 119.5, 115.9 (t, $J$ = 259.6 Hz), 33.2, 32.4, 22.2, 13.9. **$^{19}$F NMR** (375 MHz, $CDCl_3$) δ -80.7 (d, $J$ = 73.9 Hz). **IR** (thin film): 2961, 2932, 1607, 1477, 1383, 1224, 1129, 865, 827 cm$^{-1}$. **HRMS** (DART-TOF+): mass [M+H] calc'd for $C_{16}H_{18}F_2ON$ 278.1351; found 278.1351.

**5-butyl-2-(4-methoxyphenyl)pyridine [nementin-12-28]**

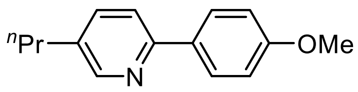

**nementin-12-28** was synthesized according to General Procedure A, B, and C using 6-bromonicotinaldehyde, ethyltriphenylphosphonium bromide, and (4-methoxyphenyl)boronic acid with an overall yield of 41% as a white solid. **$^1$H NMR** (500 MHz, $CDCl_3$) δ 8.48 – 8.46 (m, 1H), 7.95 – 7.89 (m, 2H), 7.61 – 7.57 (m, 1H), 7.55 – 7.50 (m, 1H), 7.02 – 6.95 (m, 2H), 3.86 (s, 3H), 2.61 (t, $J$ = 7.5 Hz, 2H), 1.73 – 1.59 (m, 2H), 0.97 (t, $J$ = 7.3 Hz, 3H). **$^{13}$C NMR** (125 MHz, $CDCl_3$) δ 160.1, 154.7, 149.6, 136.6, 135.4, 132.1, 127.9, 119.3, 114.0, 55.3, 34.7, 24.3, 13.7. **IR** (thin film): 2958, 2869, 1607, 1514, 1476, 1275, 1246, 1174, 1019, 818 cm$^{-1}$. **HRMS** (DART-TOF+): mass [M+H] calc'd for $C_{15}H_{18}ON$ 228.1383; found 228.1381. **mp** 39-40 °C.

**4-(5-butylpyridin-2-yl)phenol [nementin-12-29]**

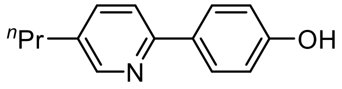

**nementin-12-29** was synthesized according to General Procedure A, B, and C using 6-bromonicotinaldehyde, ethyltriphenylphosphonium bromide, and (4-(benzyloxy)phenyl)boronic acid with an overall yield of 31% as a white solid. **$^1$H NMR** (500 MHz, $CDCl_3$) δ 8.45 – 8.39 (m, 1H), 7.73 – 7.64 (m, 2H), 7.60 – 7.53 (m, 2H), 6.78 – 6.72 (m, 2H), 2.59 (t, $J$ = 7.6 Hz, 2H), 1.71 – 1.60 (m, 2H), 0.95 (t, $J$ = 7.4 Hz, 3H). **$^{13}$C NMR** (125 MHz, $CDCl_3$) δ 158.1 (d, $J$ = 13.0 Hz), 155.3 – 155.3 (m), 148.6 (d, $J$ = 3.9 Hz), 137.8 – 137.3 (m), 135.7, 130.1 (d, $J$ = 12.8 Hz), 128.4, 120.5 (d, $J$ = 5.3 Hz), 115.9 (d, $J$ = 1.5 Hz), 34.6, 24.2, 13.7. **IR** (thin film): 3851, 2967, 1603, 1479, 1388, 1275, 1244, 1172, 824 $cm^{-1}$. **HRMS** (DART-TOF+): mass [M+H] calc'd for $C_{14}H_{16}ON$ 214.1226; found 214.1230. **mp** 133-134 °C.

**2-(4-(2,2-difluoroethoxy)phenyl)-5-propylpyridine [nementin-12-30]**

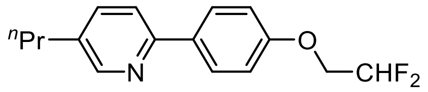

**nementin-12-29** (21 mg, 0.1 mmol) was dissolved in DMF (0.33 ml, 0.3 M) and cooled to 0 °C. NaH (6 mg, 0.15 mmol, 60%, 1.5 equiv.) was added and the mixture was stirred at 0 °C for 30 minutes. 2-bromo-1,1-difluoroethane (0.02 mL, 0.15 mmol, 1.5 equiv.) was added dropwise and the reaction mixture was stirred from 0 °C to room temperature overnight. $H_2O$ was added and the aqueous phase was extracted with EtOAc. The combined organics was washed with brine, dried with $MgSO_4$, and concentrated in vacuo. The crude mixture was purified by flash column chromatography, eluting with 5 to 10% (v/v) EtOAc:pentanes to give **nementin-12-30** (18 mg, 0.063 mmol, 63%) as a white solid. **$^1$H NMR** (500 MHz, $CDCl_3$) δ 8.50 – 8.44 (m, 1H), 7.98 – 7.89 (m, 2H), 7.59 (dd, *J* = 8.1, 0.9 Hz, 1H), 7.56 – 7.48 (m, 1H), 7.06 – 6.96 (m, 2H), 6.11 (tt, *J* = 55.2, 4.1 Hz, 1H), 4.24 (td, *J* = 13.0, 4.1 Hz, 2H), 2.61 (t, *J* = 7.4 Hz, 2H), 1.71 – 1.64 (m, 2H), 0.97 (t, *J* = 7.3 Hz, 3H). **$^{13}$C NMR** (125 MHz, $CDCl_3$) δ 158.2, 154.3, 149.7, 136.7, 135.8, 133.4, 128.1, 119.5, 114.7, 113.6 (t, *J* = 241.1 Hz), 67.3 (t, *J* = 29.6 Hz), 34.7, 24.3, 13.7. **$^{19}$F NMR** (375 MHz, $CDCl_3$) δ -125.2 (td, *J* = 55.6, 12.9 Hz). **IR** (thin film): 2958, 2931, 2872, 1606, 1514, 1475, 1274, 1240, 1135, 1065, 914, 821 $cm^{-1}$. **HRMS** (DART-TOF+): mass [M+H] calc'd for $C_{16}H_{18}F_2ON$ 278.1351; found 278.1345. **mp** 67-68 °C.

**$^{13}$C NMR (125 MHz, $CDCl_3$)**

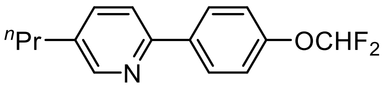


**nementin-12**

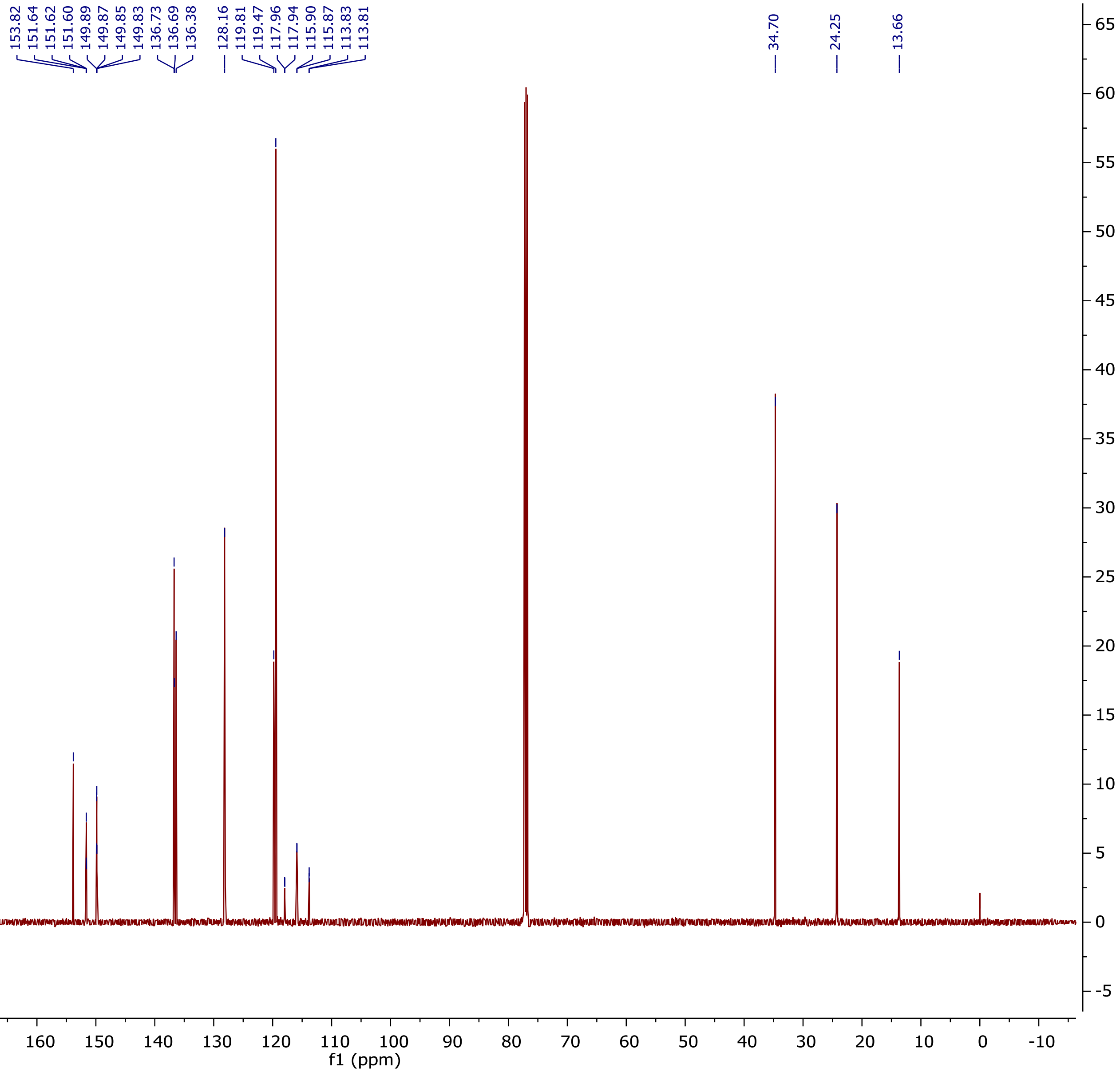

$^{19}$F NMR (375 MHz, $CDCl_3$)

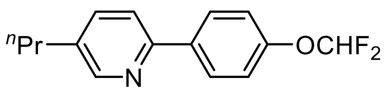


**nementin-12**

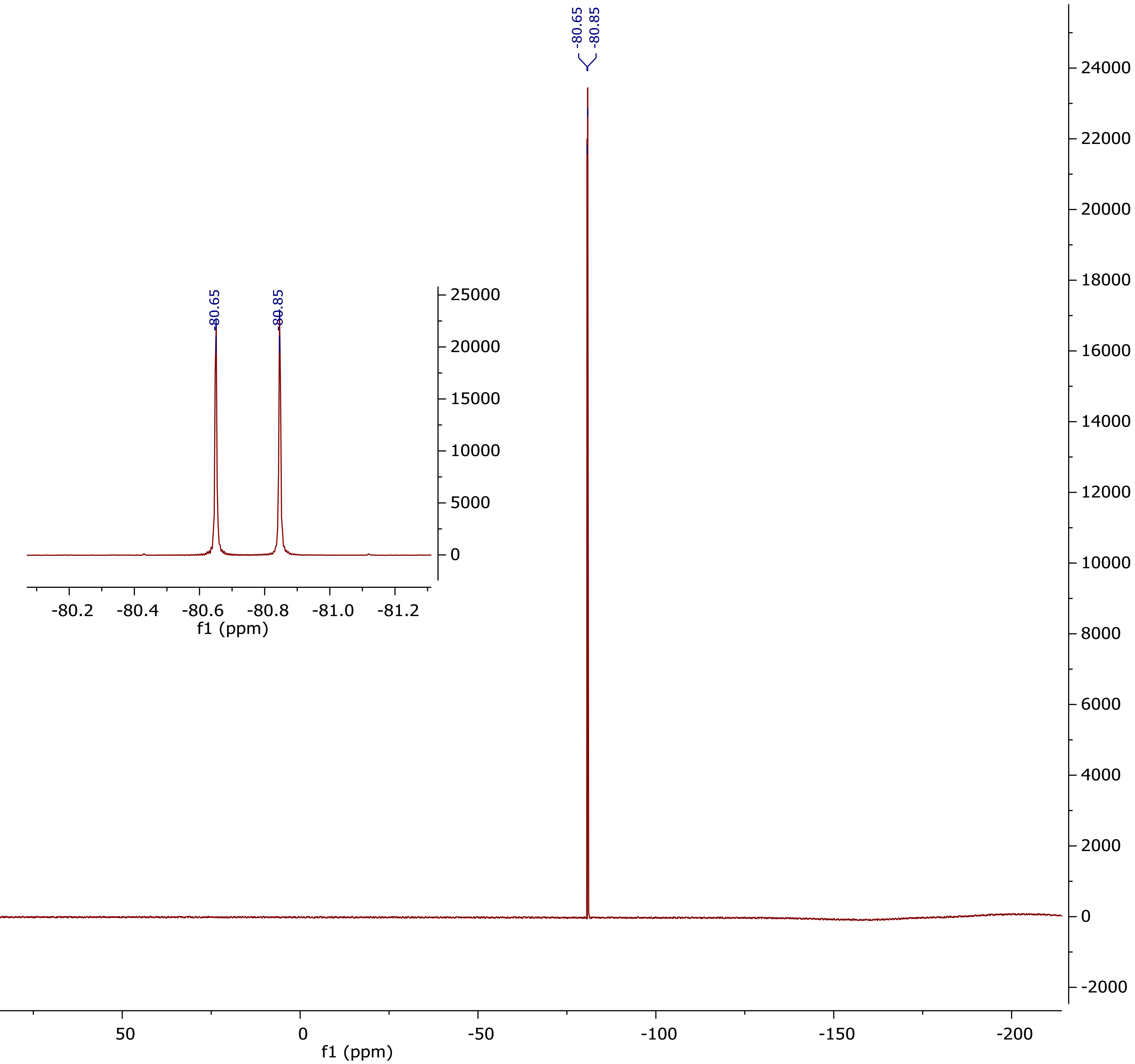

$^1$H NMR (500 MHz, $CDCl_3$) (nementin-12-5)

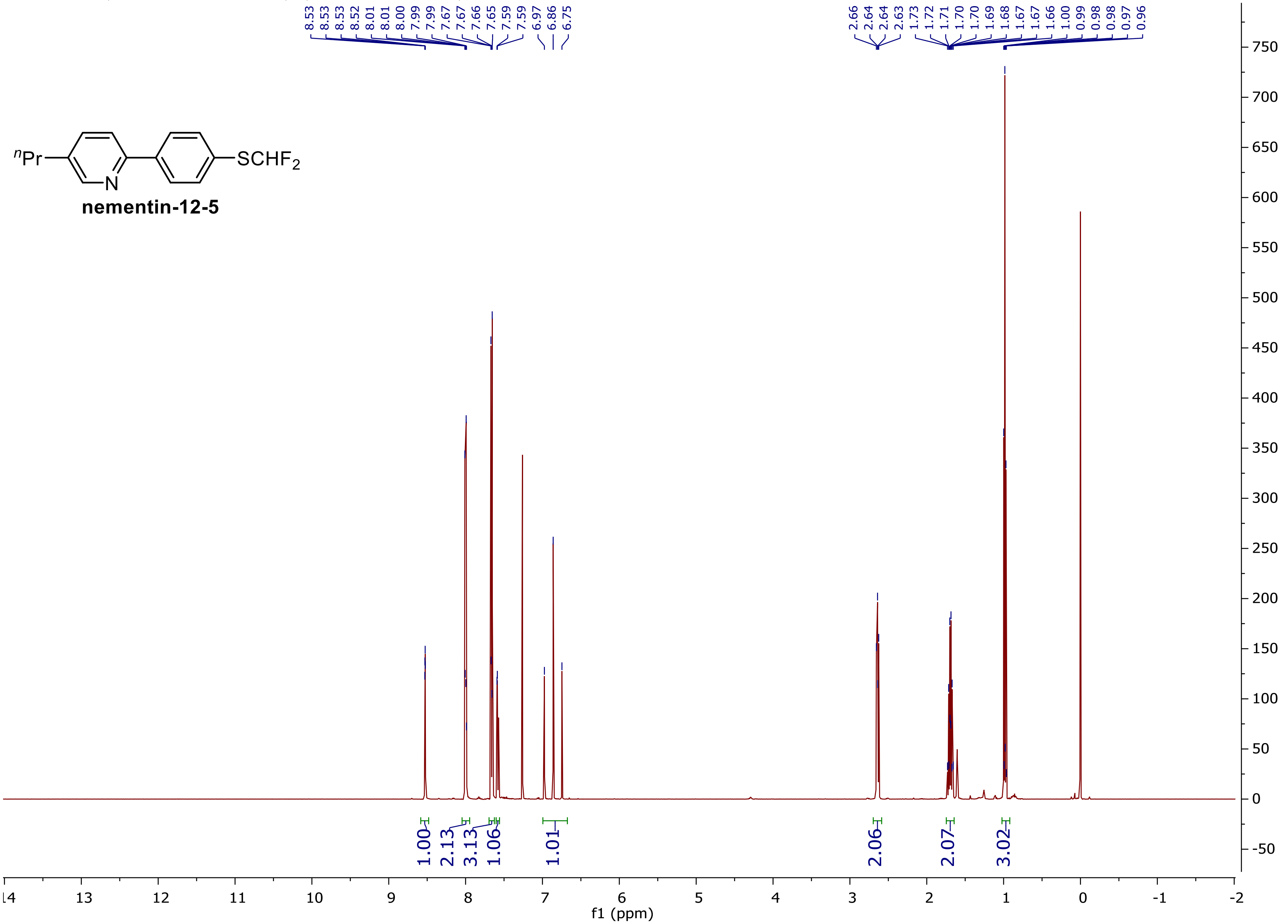

$^{13}C$ NMR (125 MHz, $CDCl_3$)

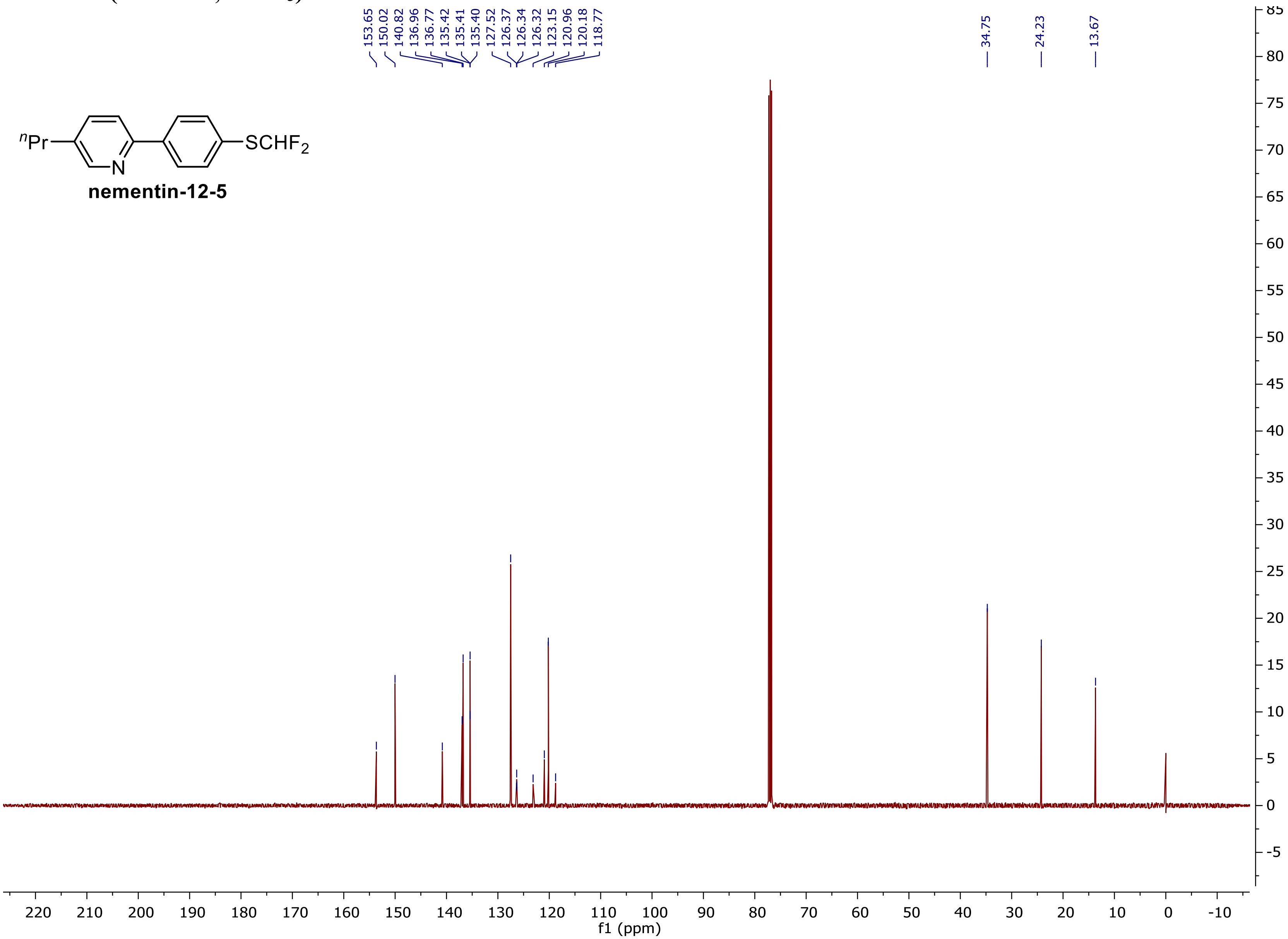

$^{19}$F NMR (375 MHz, $CDCl_3$)

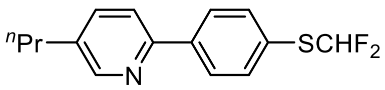


**nementin-12-5**

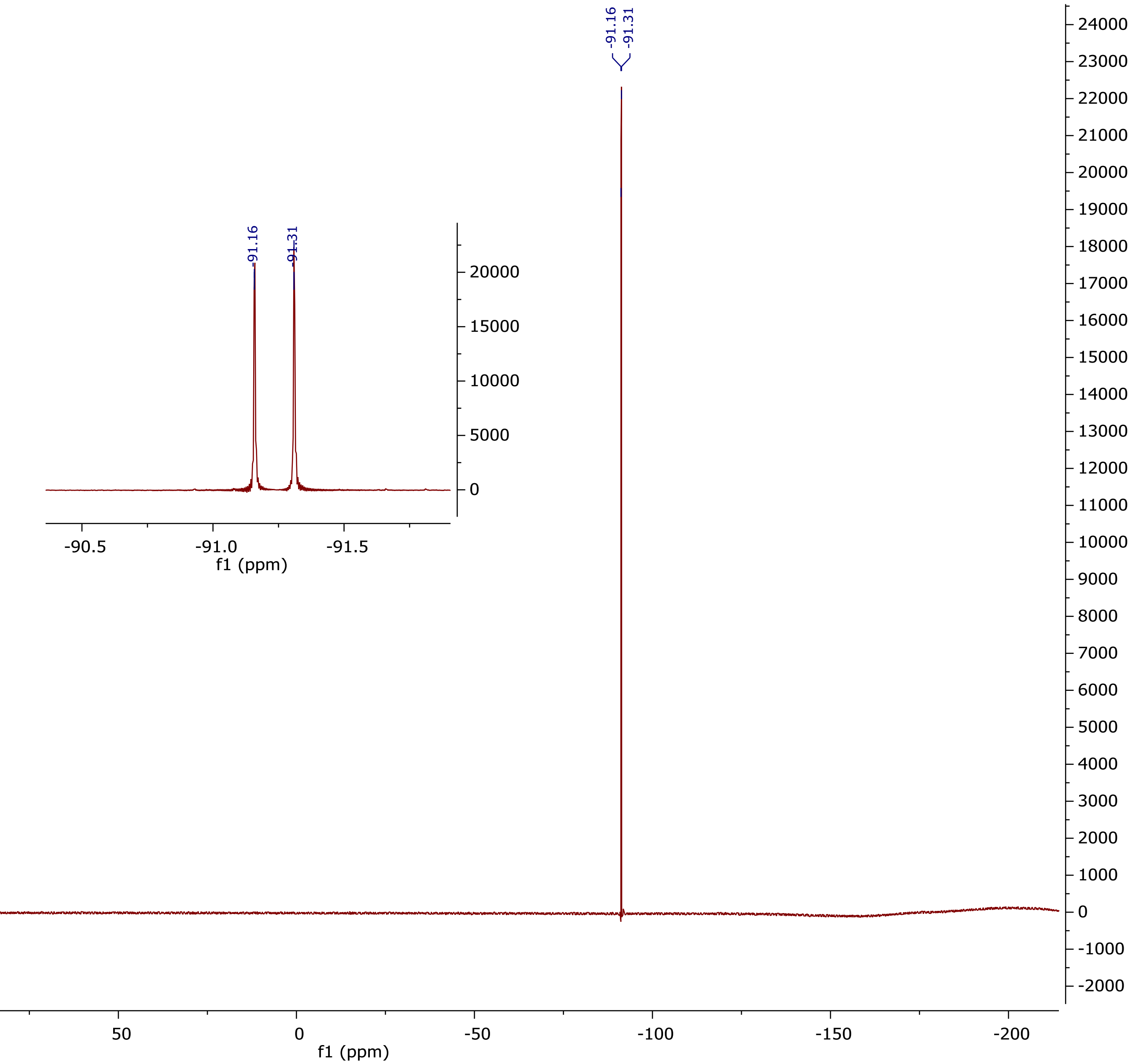

**[1]H NMR (500 MHz, CDCl3) (nementin-12-6)**

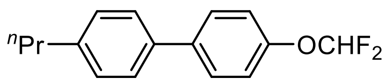


**nementin-12-6**

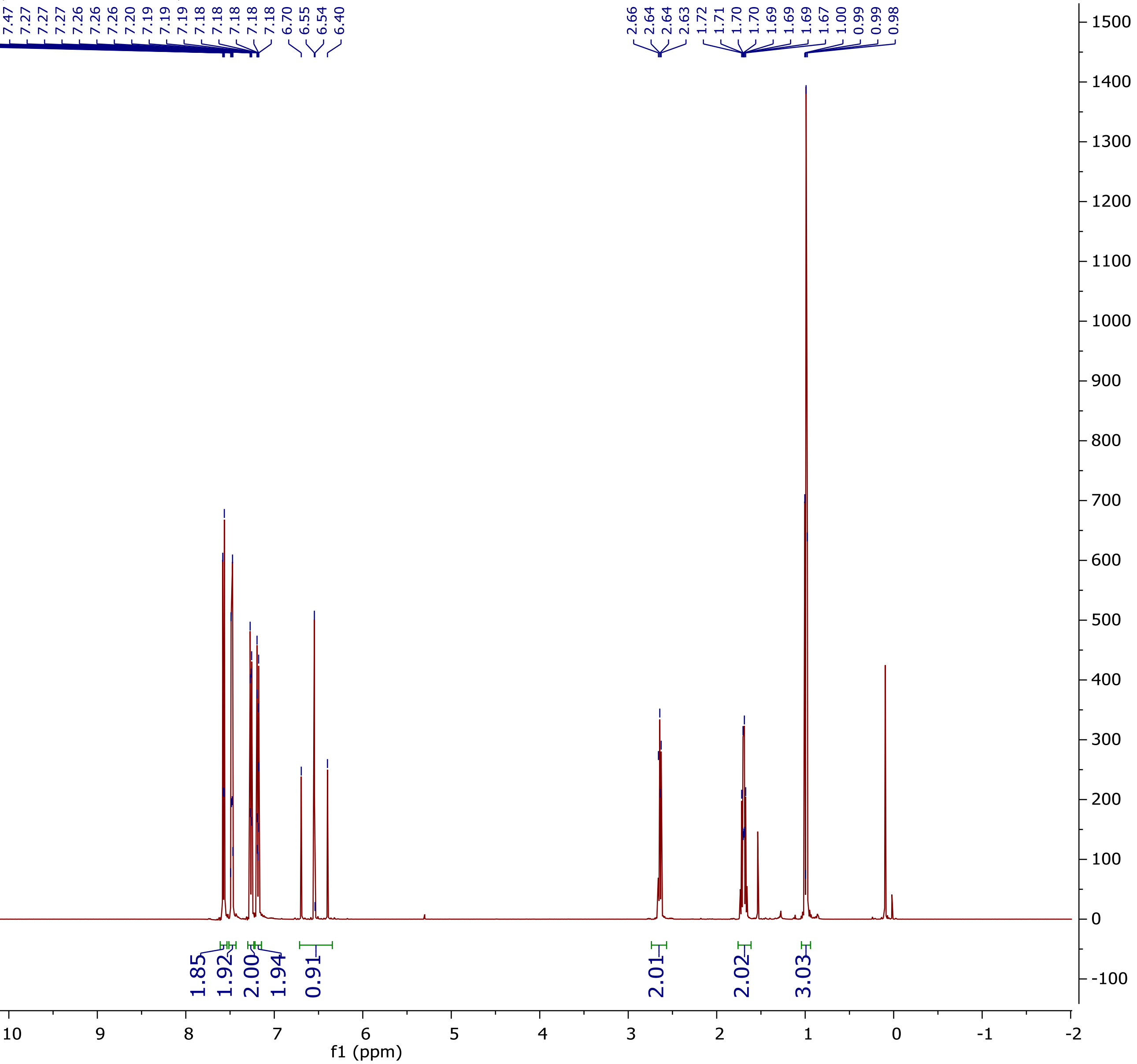

**$^{13}$C NMR (125 MHz, CDCl$_3$)**

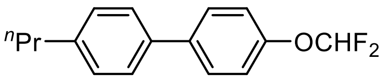

**nementin-12-6**

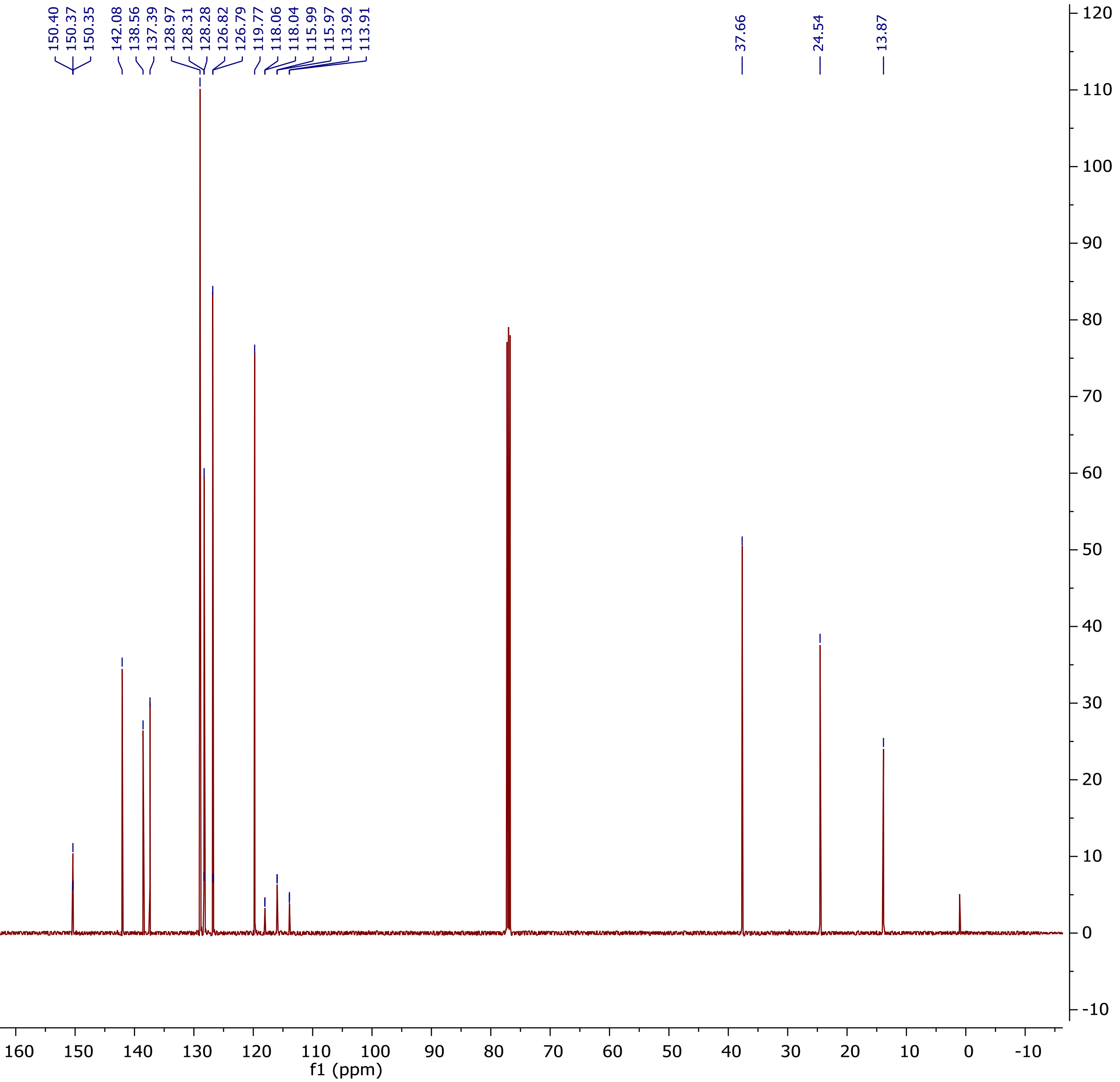

$^{19}$F NMR (375 MHz, CDCl$_3$)

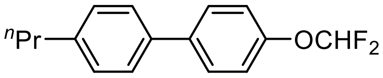


**nementin-12-6**

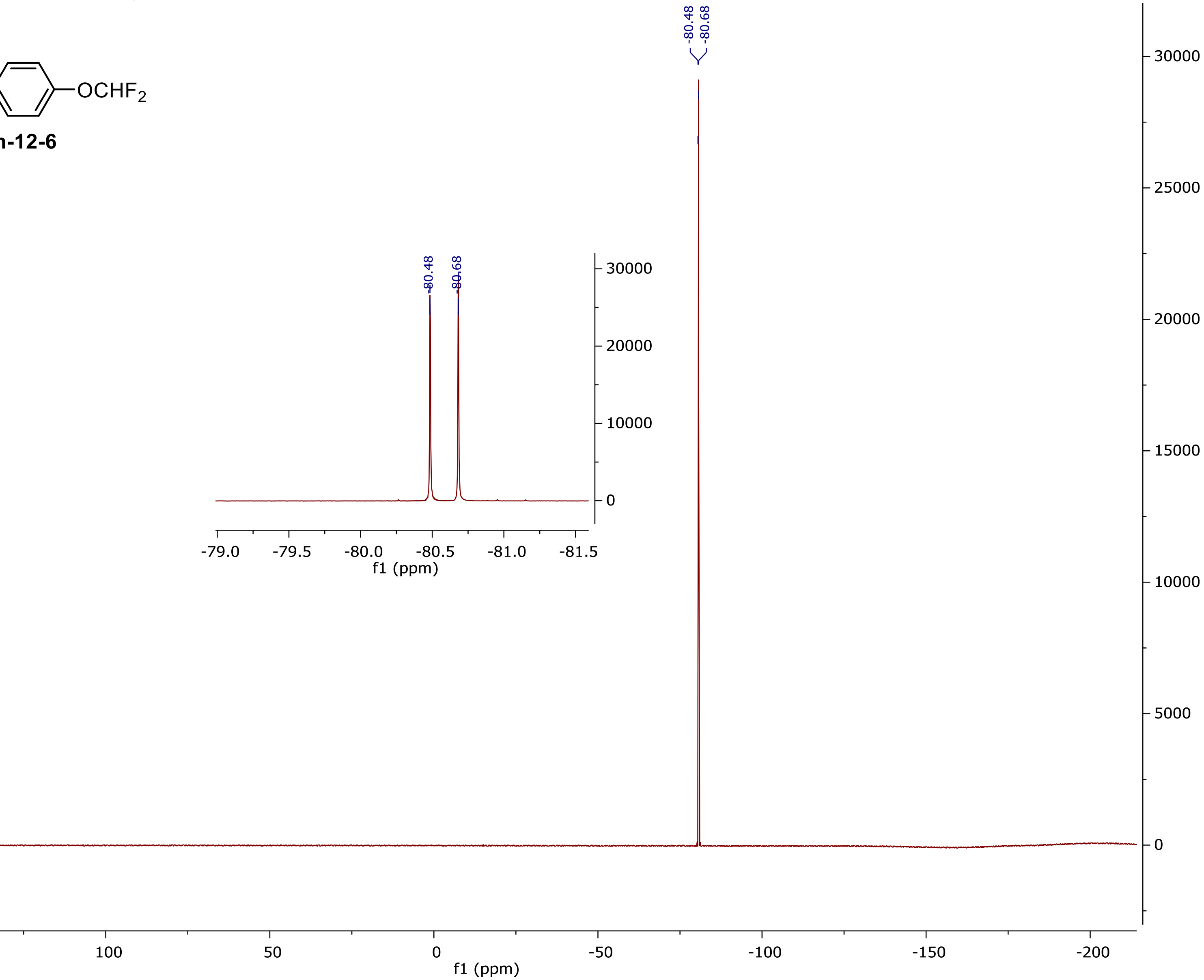

### $^{1}H$ NMR (500 MHz, $CDCl_3$) (nementin-12-8)

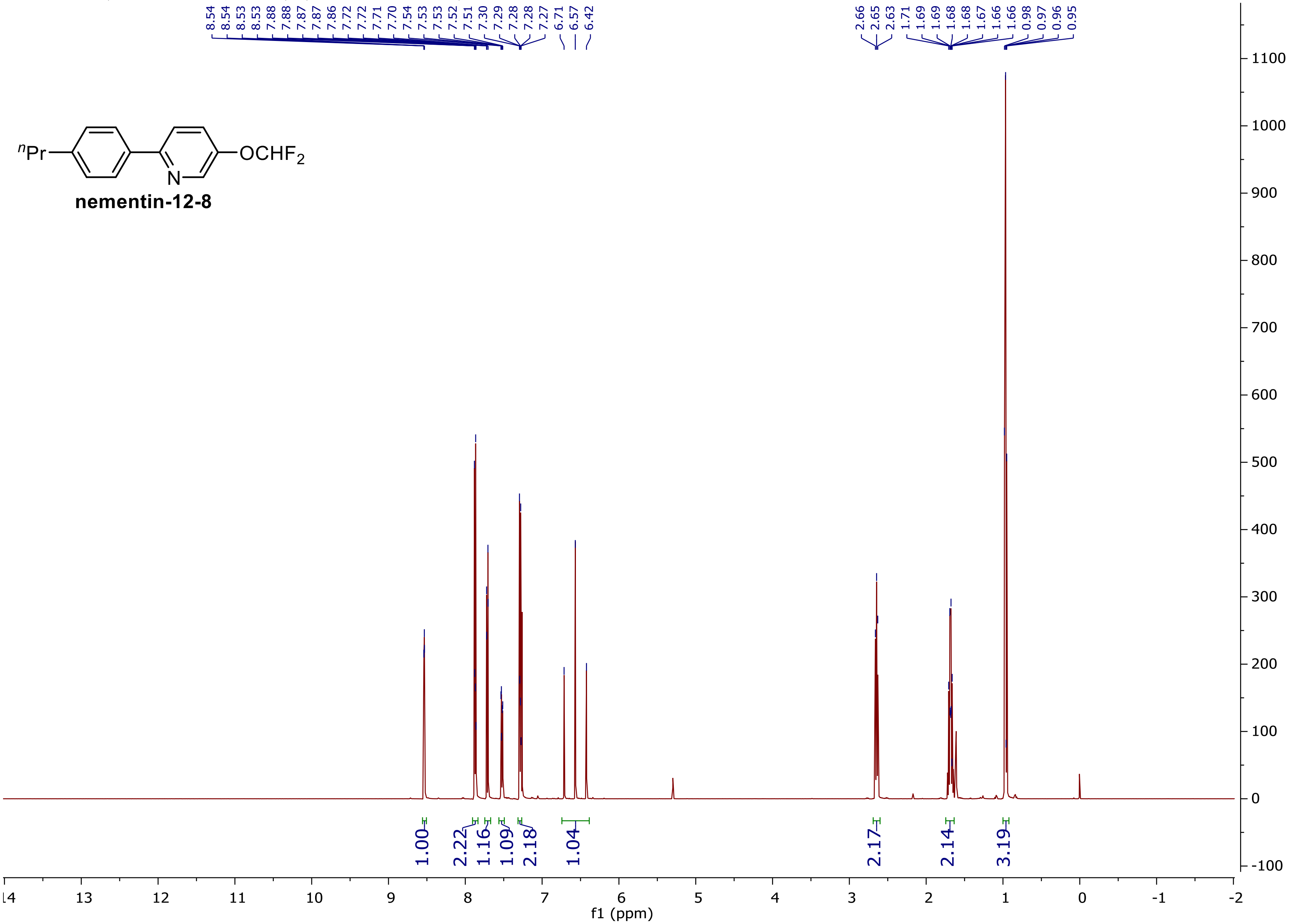

**$^{13}$C NMR (125 MHz, $CDCl_3$)**

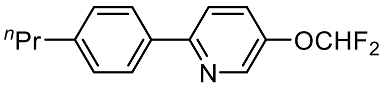


**nementin-12-8**

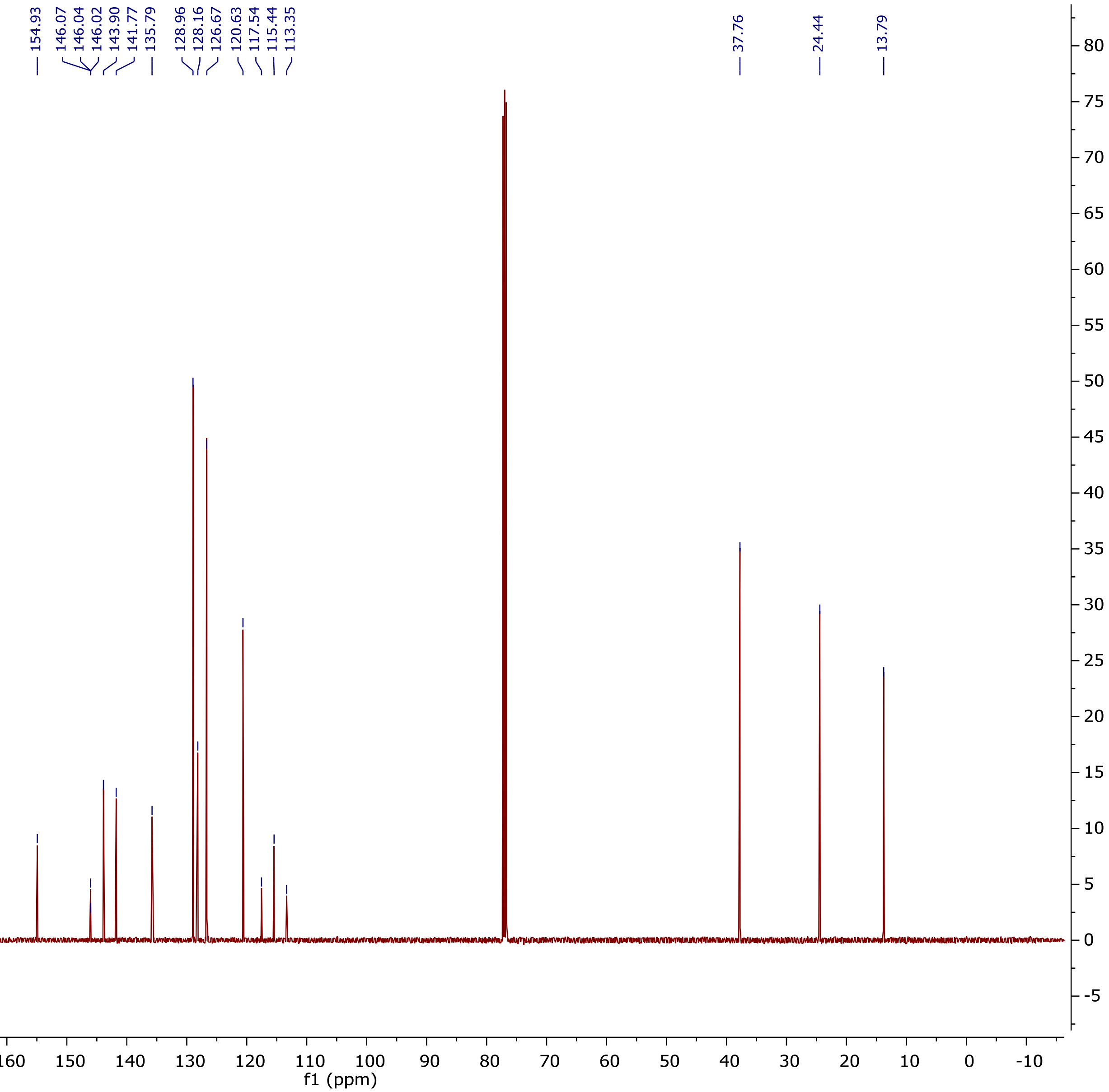

$^{19}$F NMR (375 MHz, $CDCl_3$)

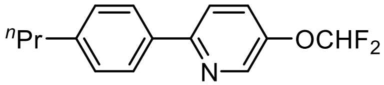


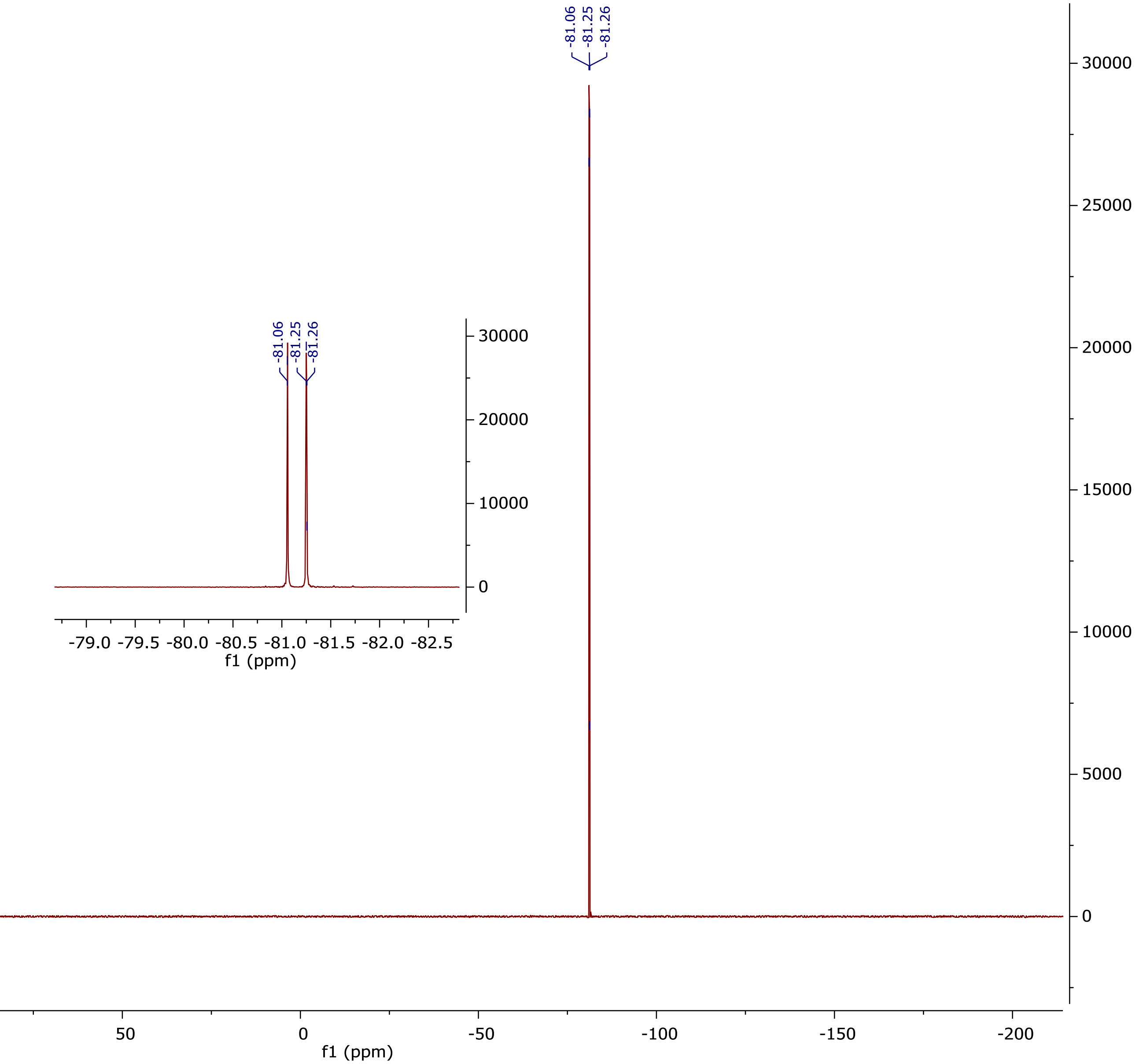

[1]H NMR (500 MHz, CDCl₃) (nementin-12-9)

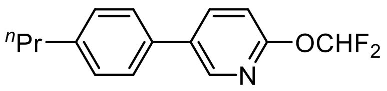


**nementin-12-9**

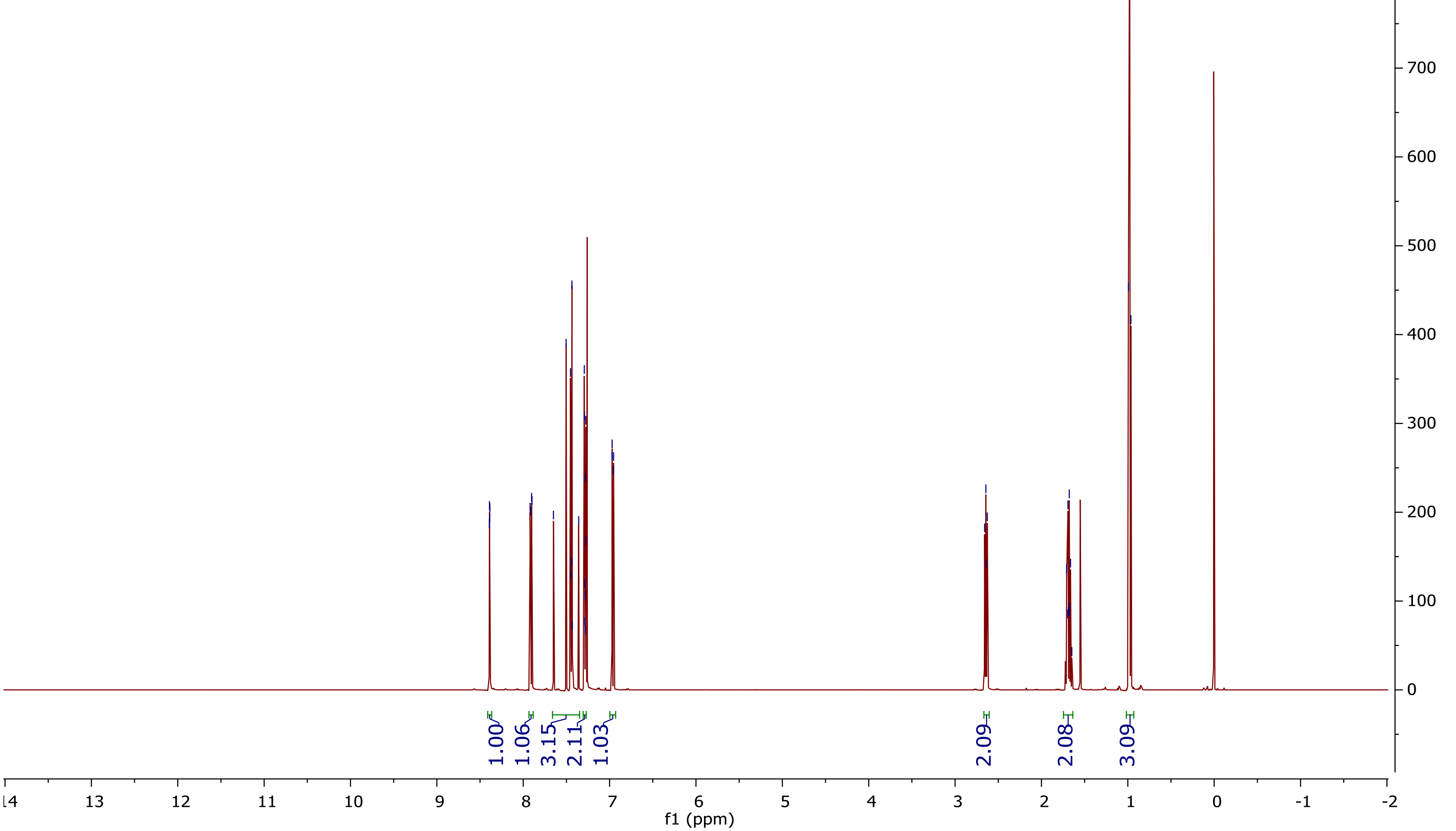

**$^{13}$C NMR (125 MHz, CDCl$_3$)**

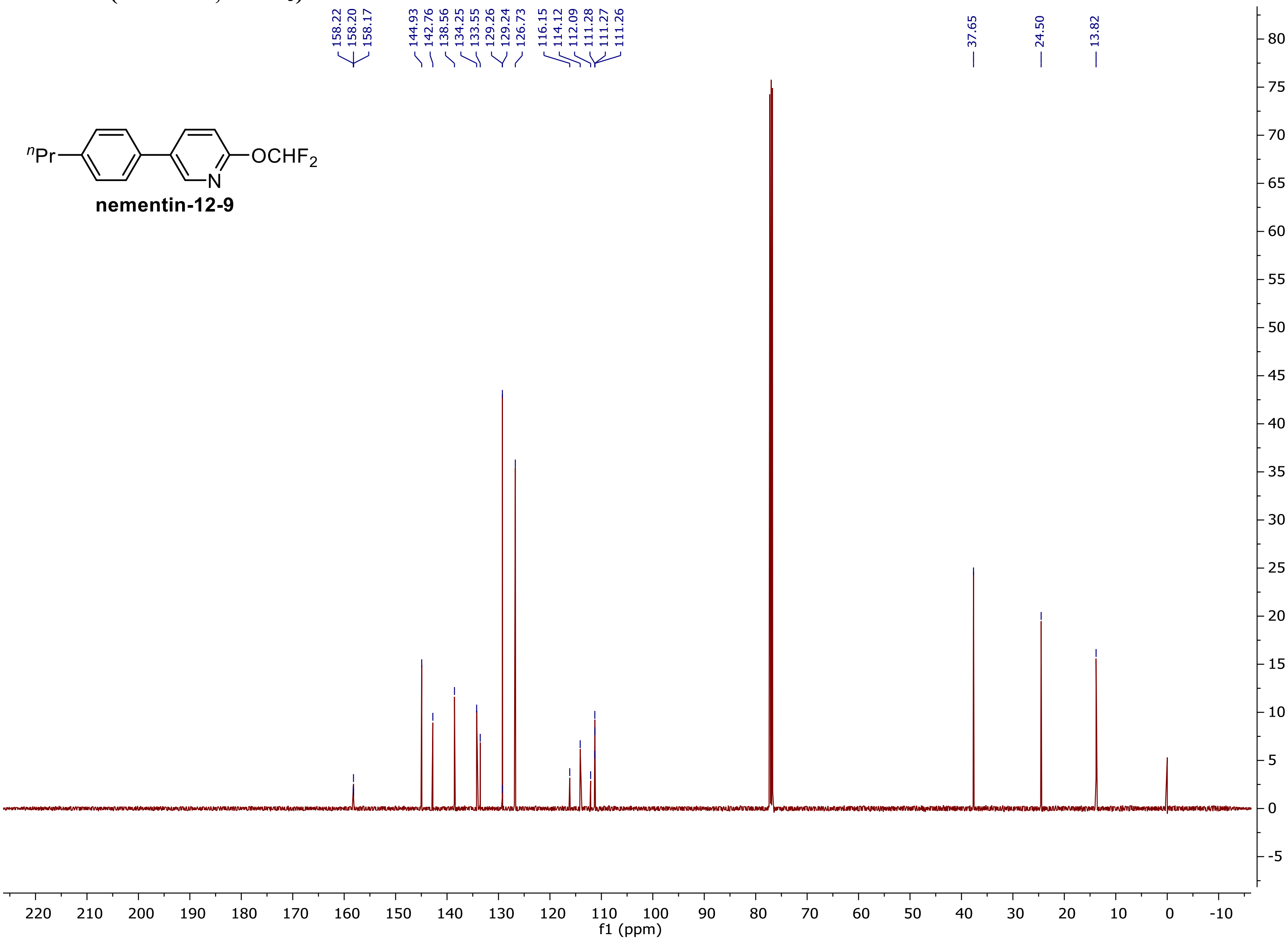

$^{19}$F NMR (375 MHz, $CDCl_3$)

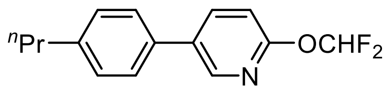


**nementin-12-9**

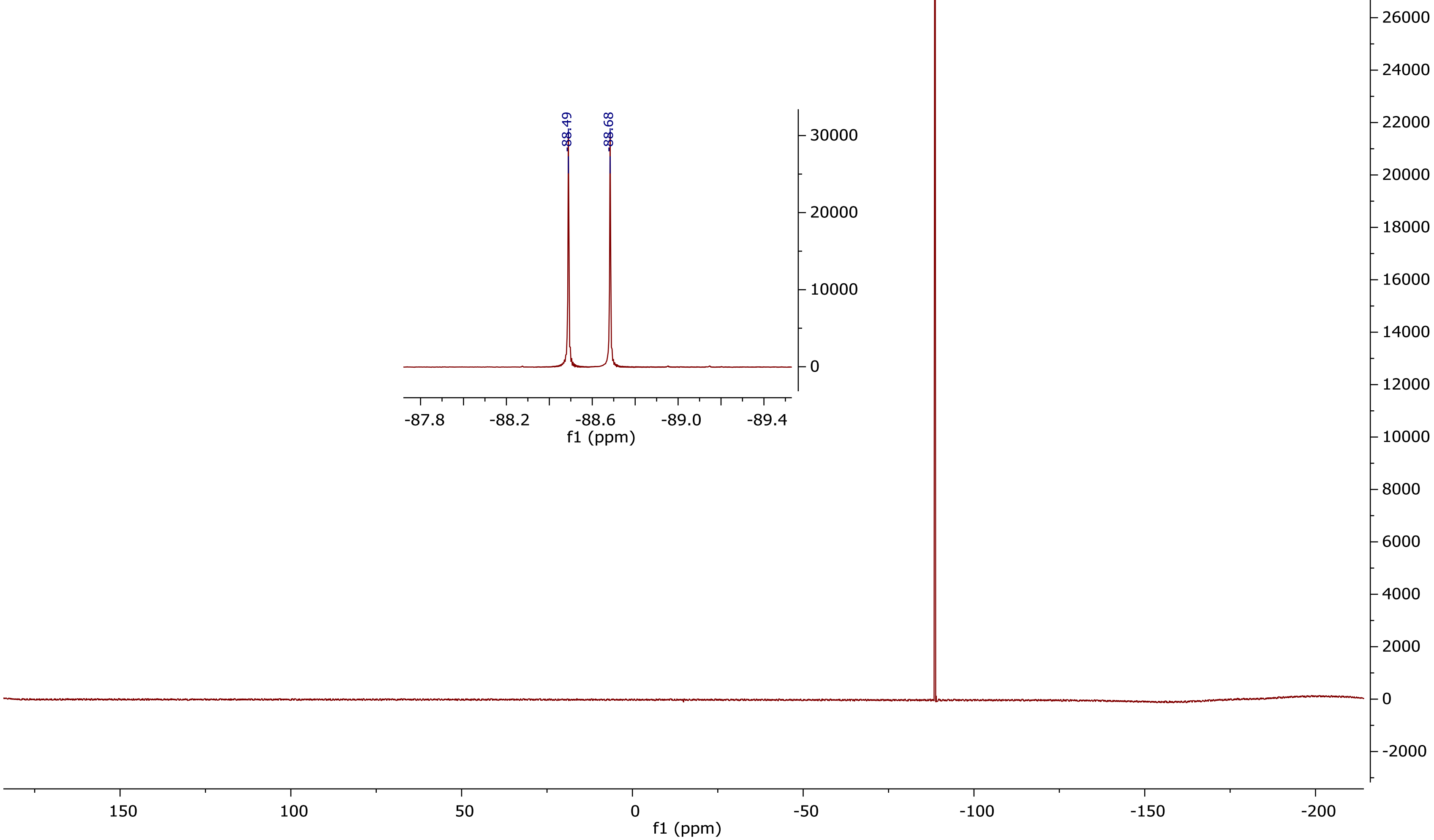

¹H NMR (500 MHz, $CDCl_3$) (nementin-12-10)

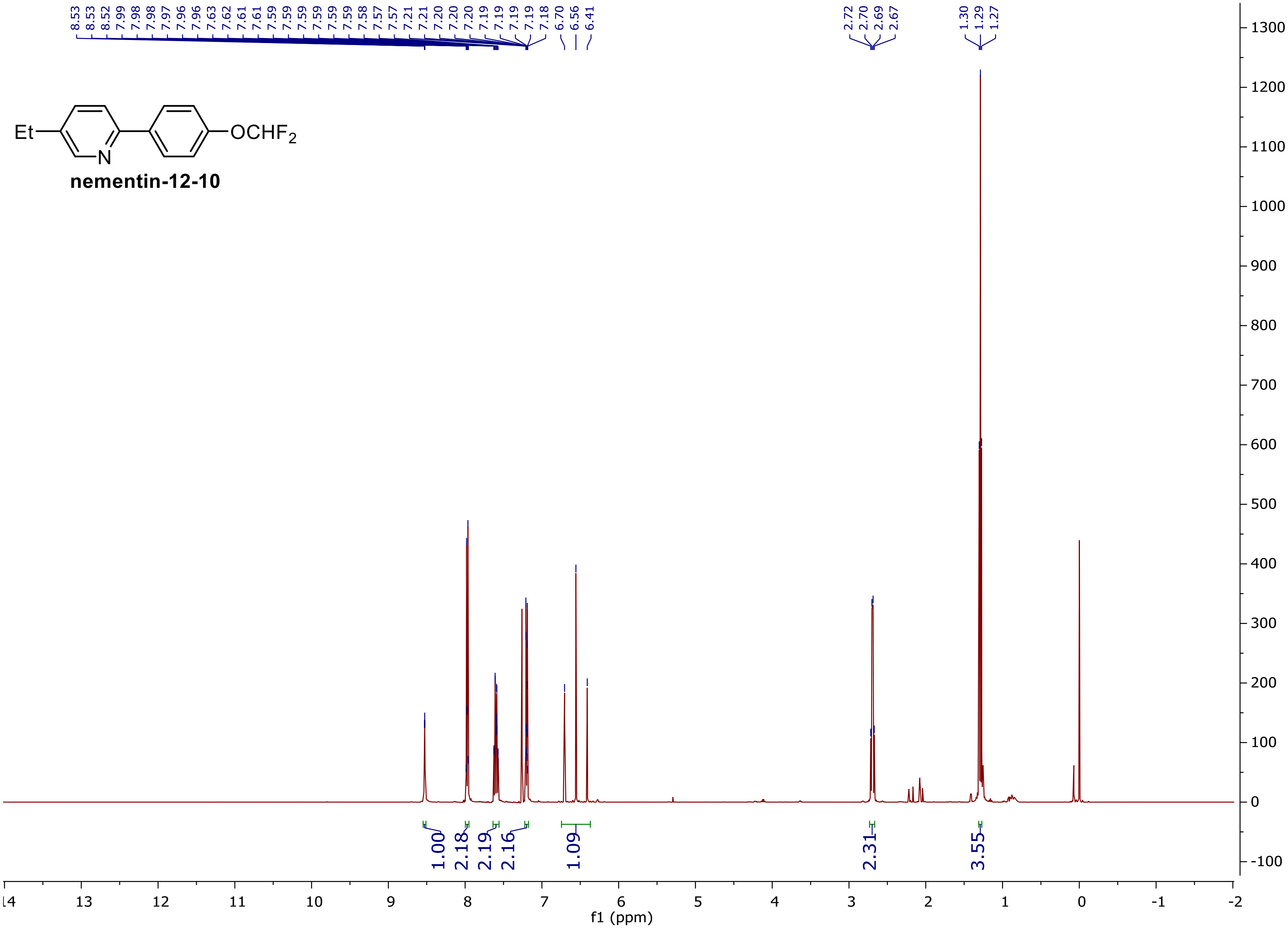

### $^{13}C$ NMR (125 MHz, $CDCl_3$)

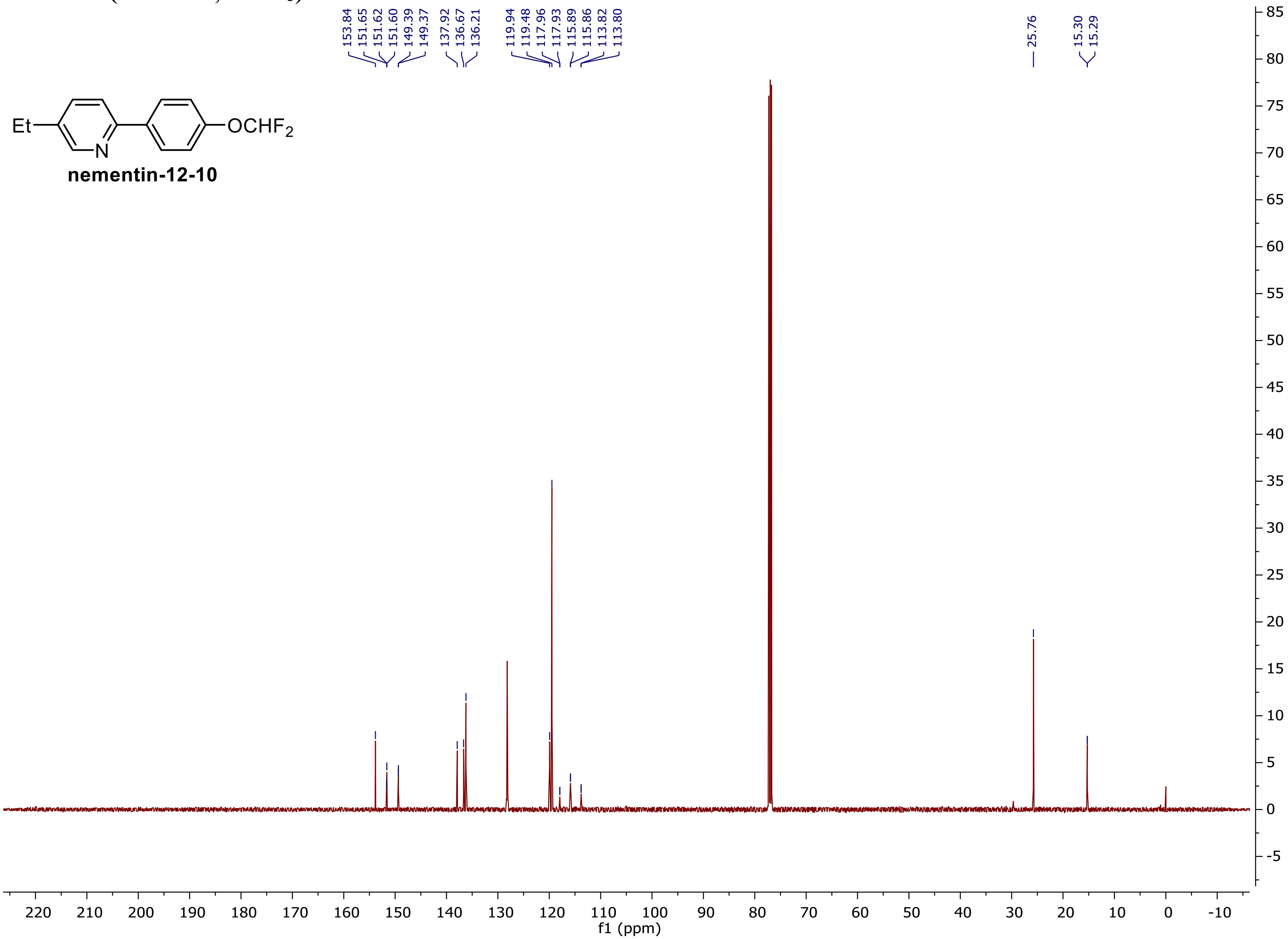

$^{19}F$ NMR (375 MHz, $CDCl_3$)

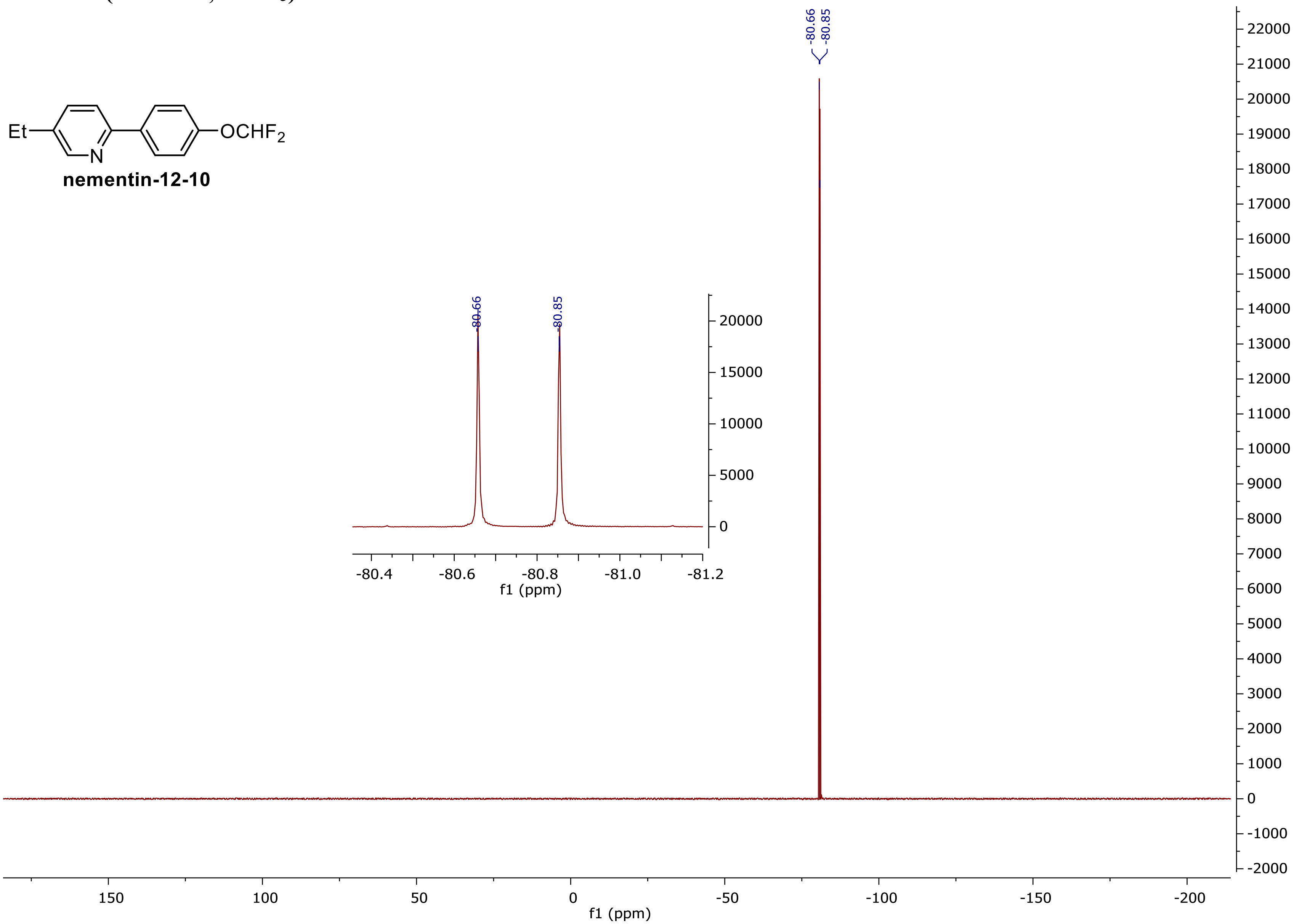

### $^{1}H$ NMR (500 MHz, $CDCl_3$) (nementin-12-11)

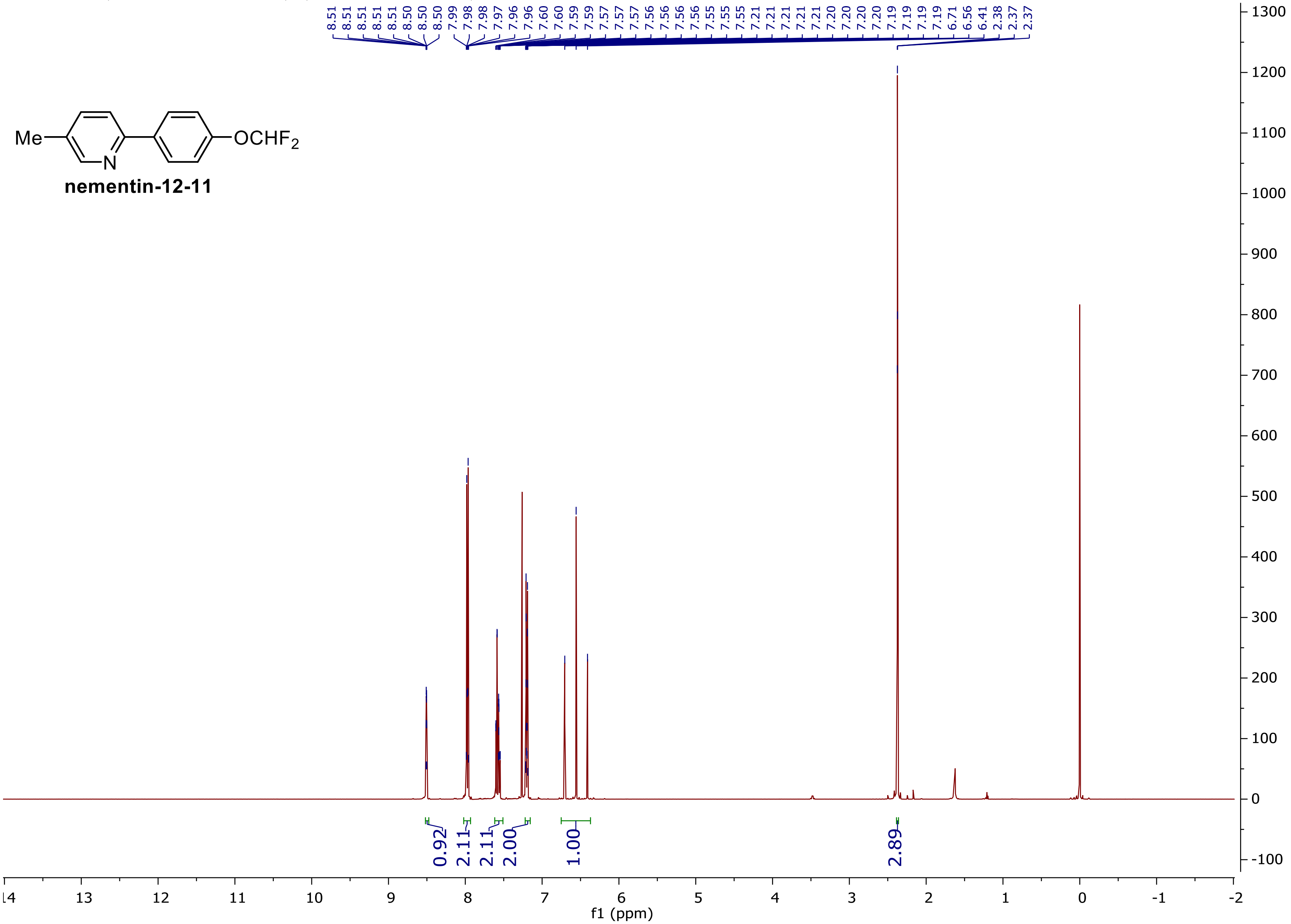

$^{13}C$ NMR (125 MHz, $CDCl_3$)

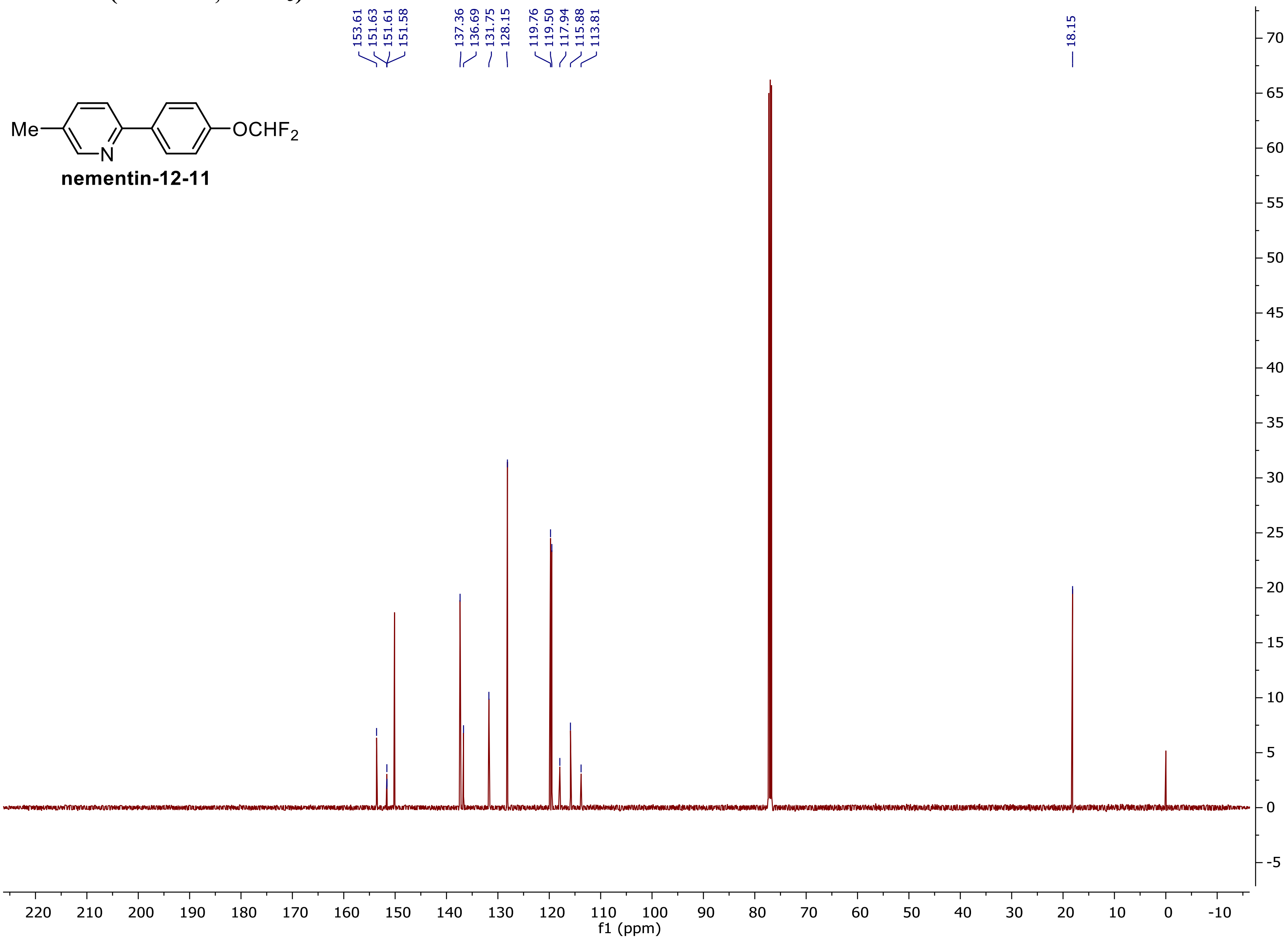

$^{19}$F NMR (375 MHz, CDCl$_3$)

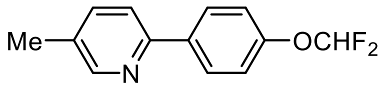


**nementin-12-11**

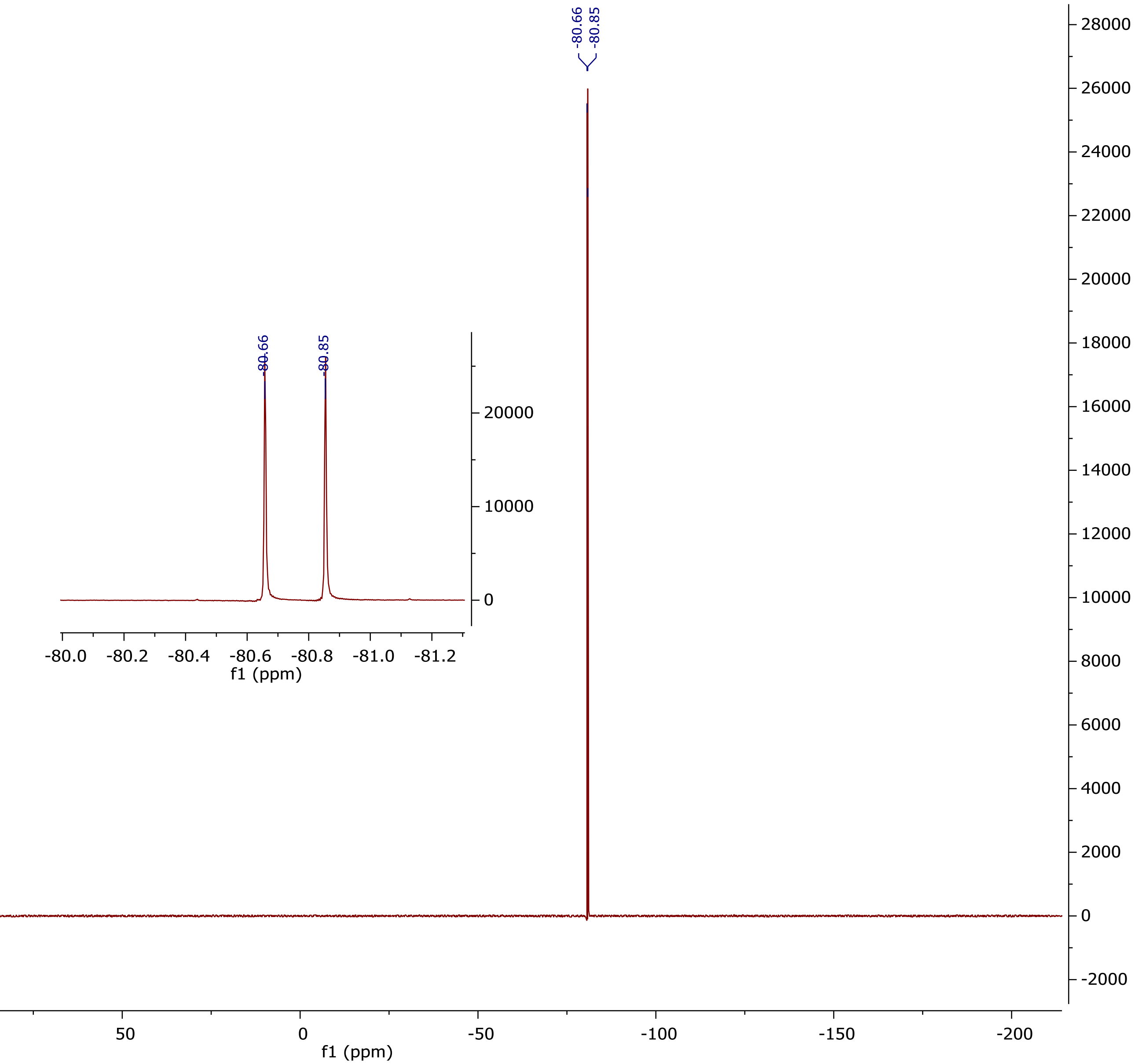

### $^1$H NMR (500 MHz, $CDCl_3$) (nementin-12-12)

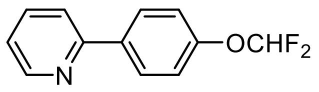


nementin-12-12

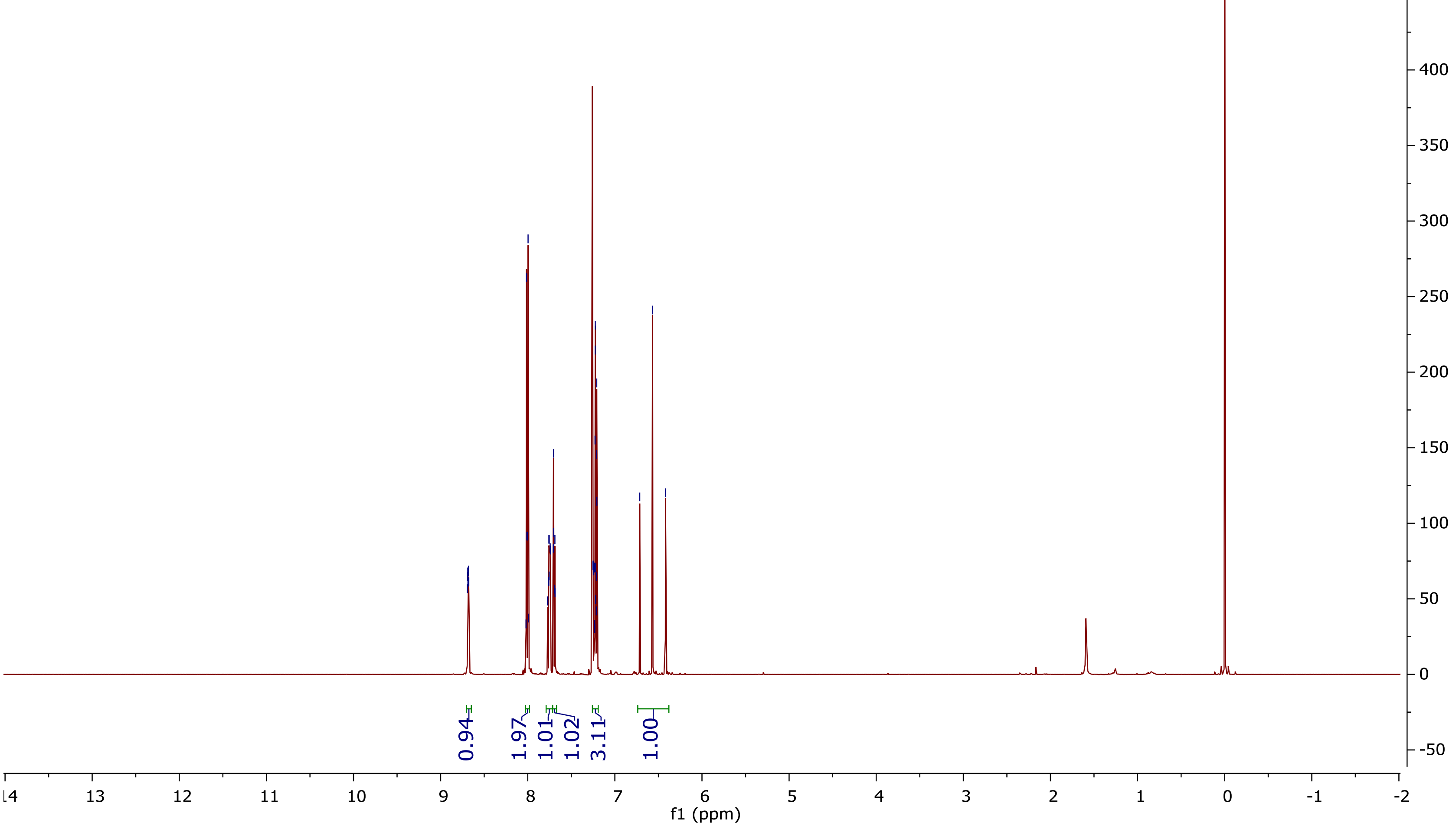

**$^{13}$C NMR (125 MHz, CDCl$_3$)**

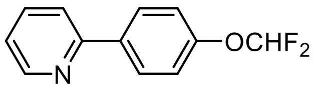


**nementin-12-12**

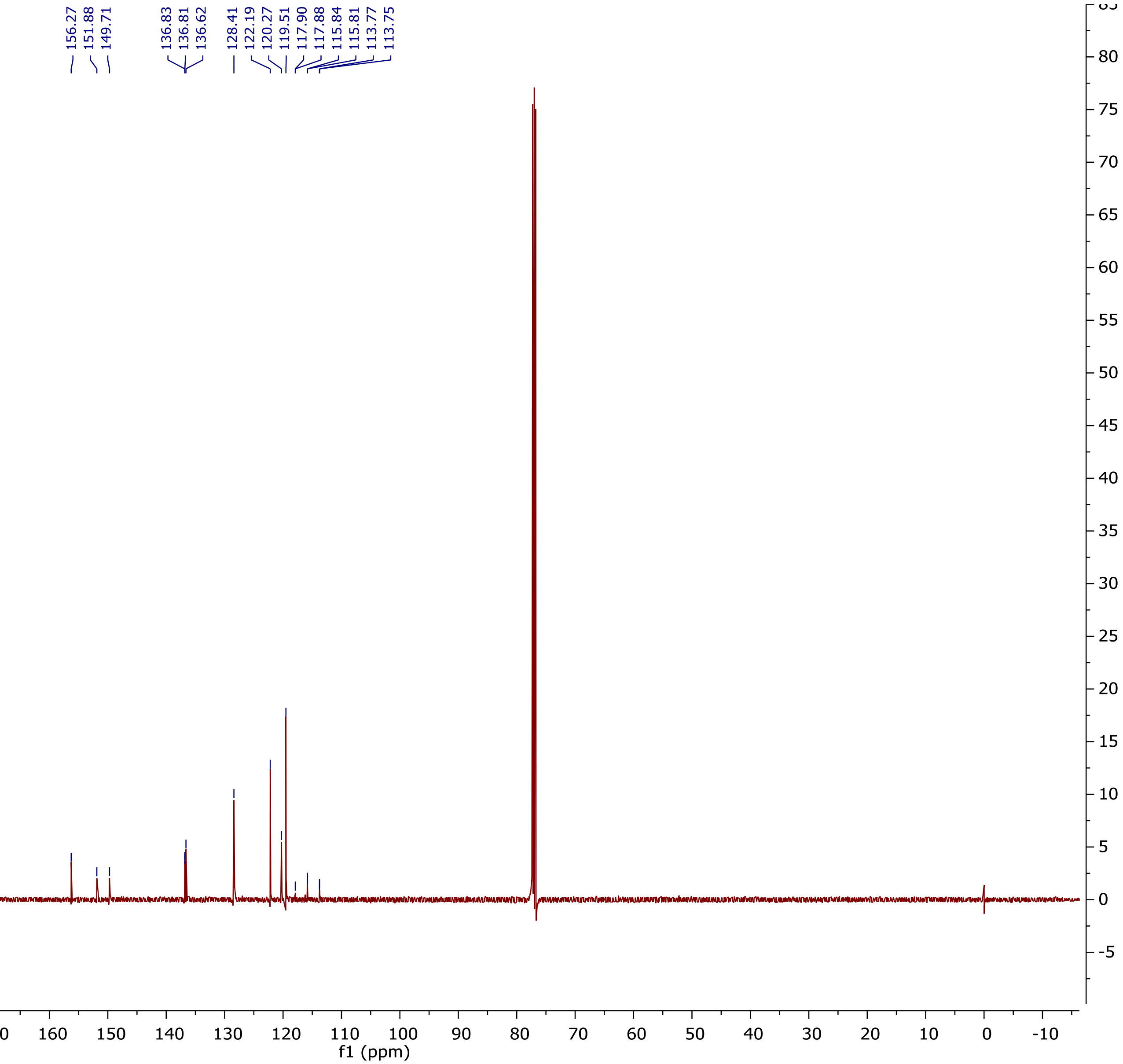

$^{19}$F NMR (375 MHz, $CDCl_3$)

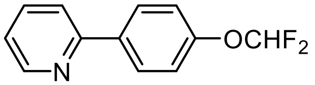


**nementin-12-12**

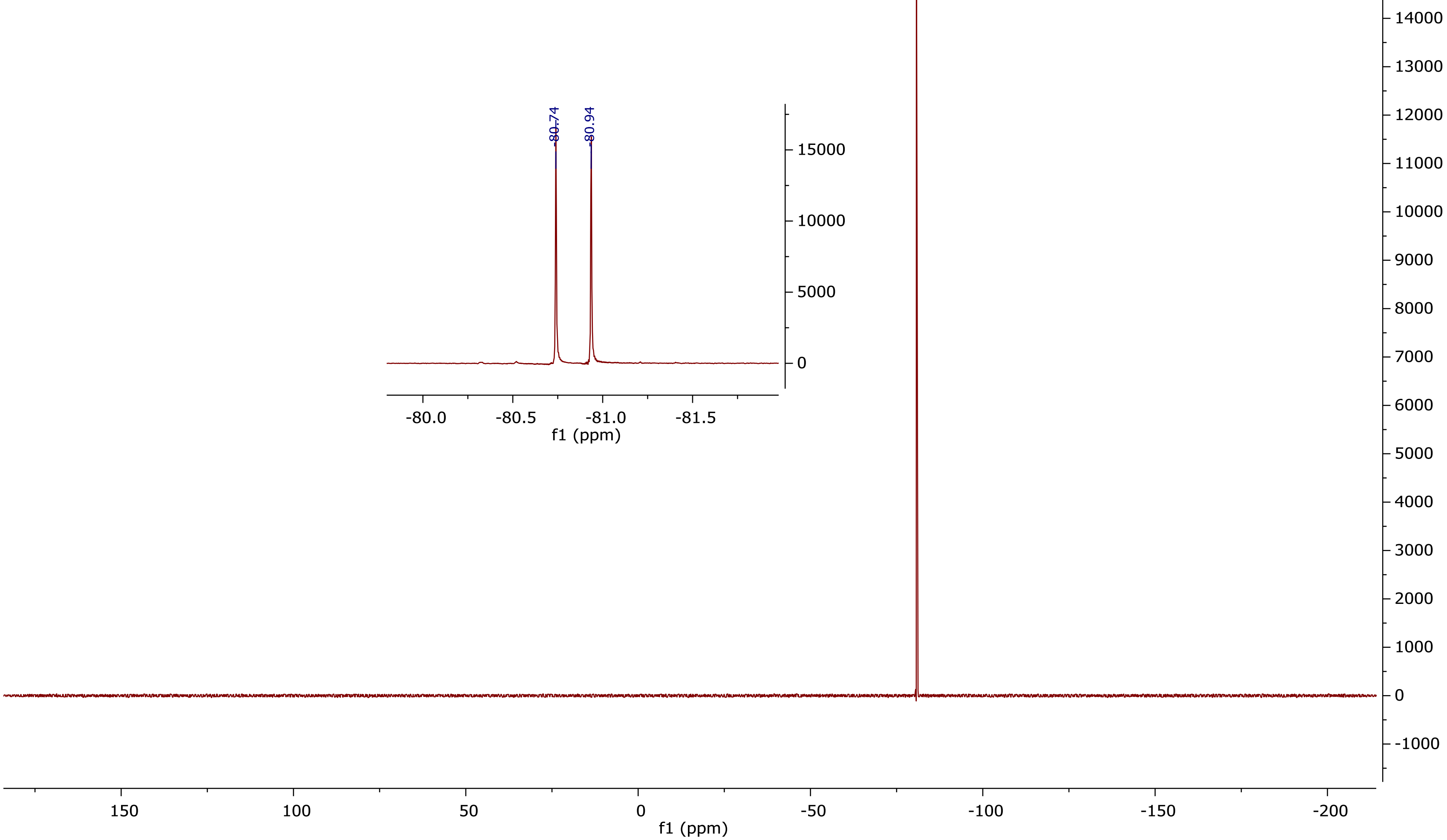

**$^{1}$H NMR (500 MHz, $CDCl_3$) (nementin-12-14)**

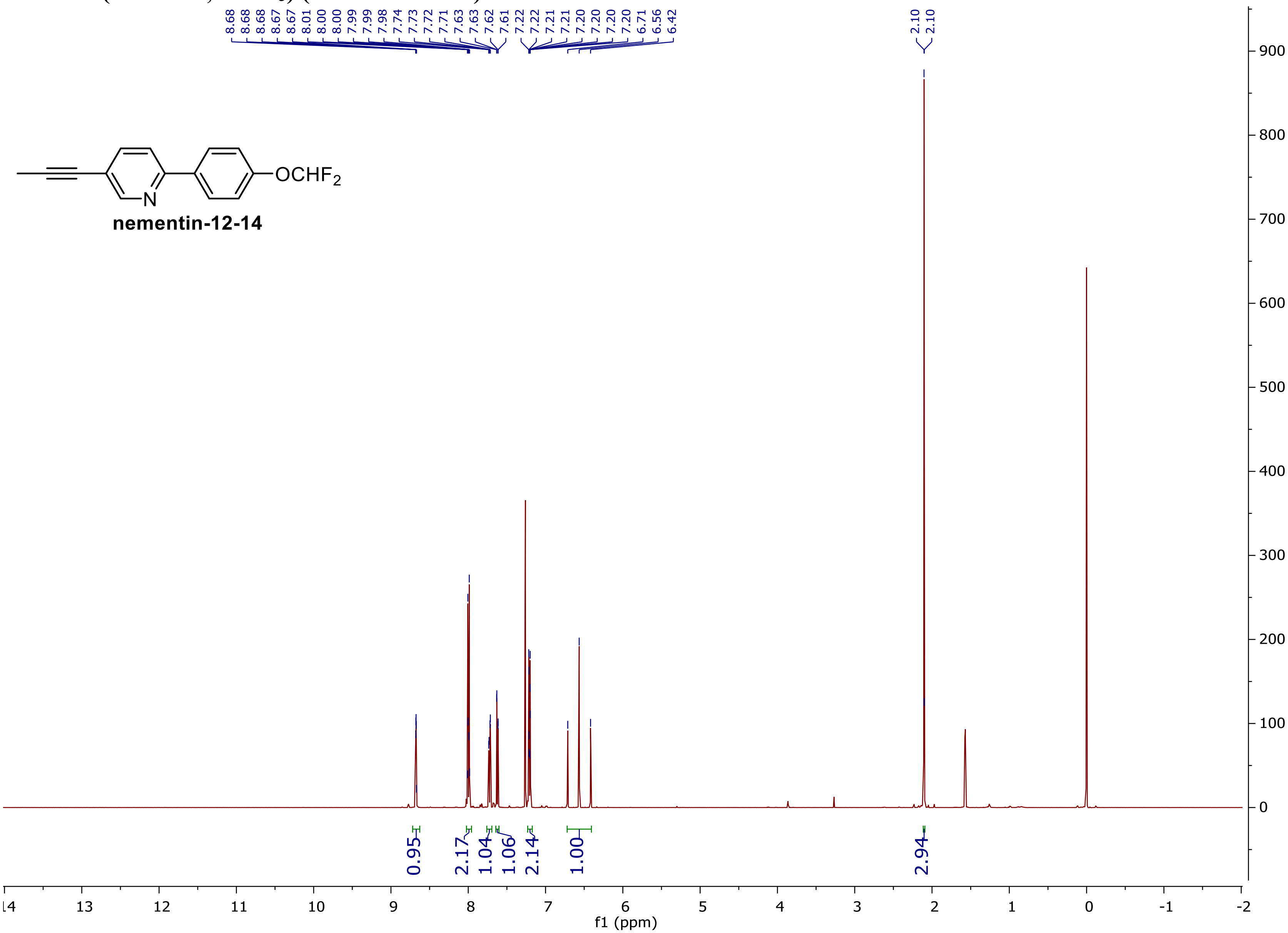

**$^{13}$C NMR (125 MHz, CDCl$_3$)**

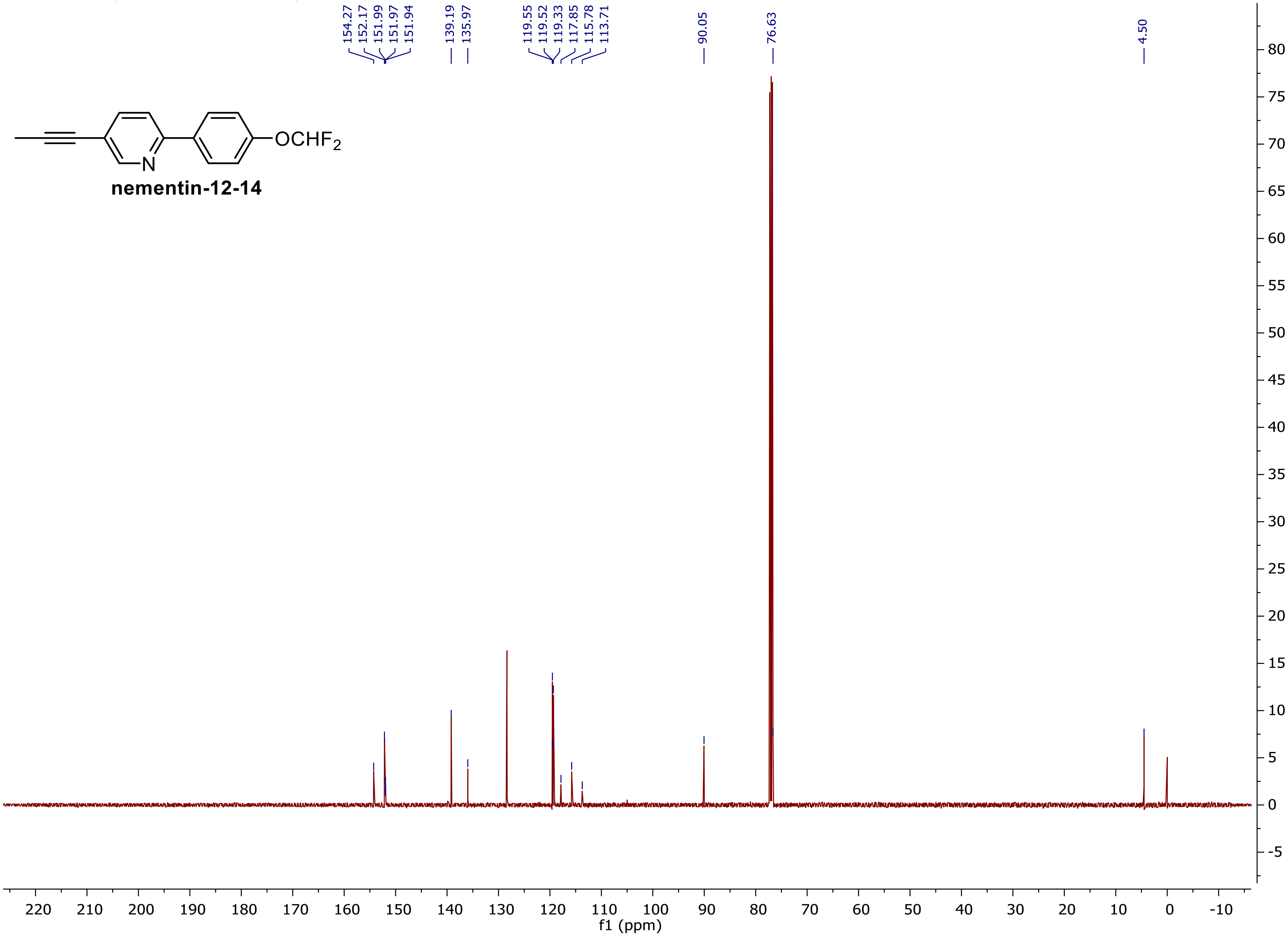

$^{19}F$ NMR (375 MHz, $CDCl_3$)

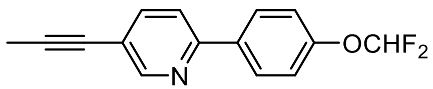


**nementin-12-14**

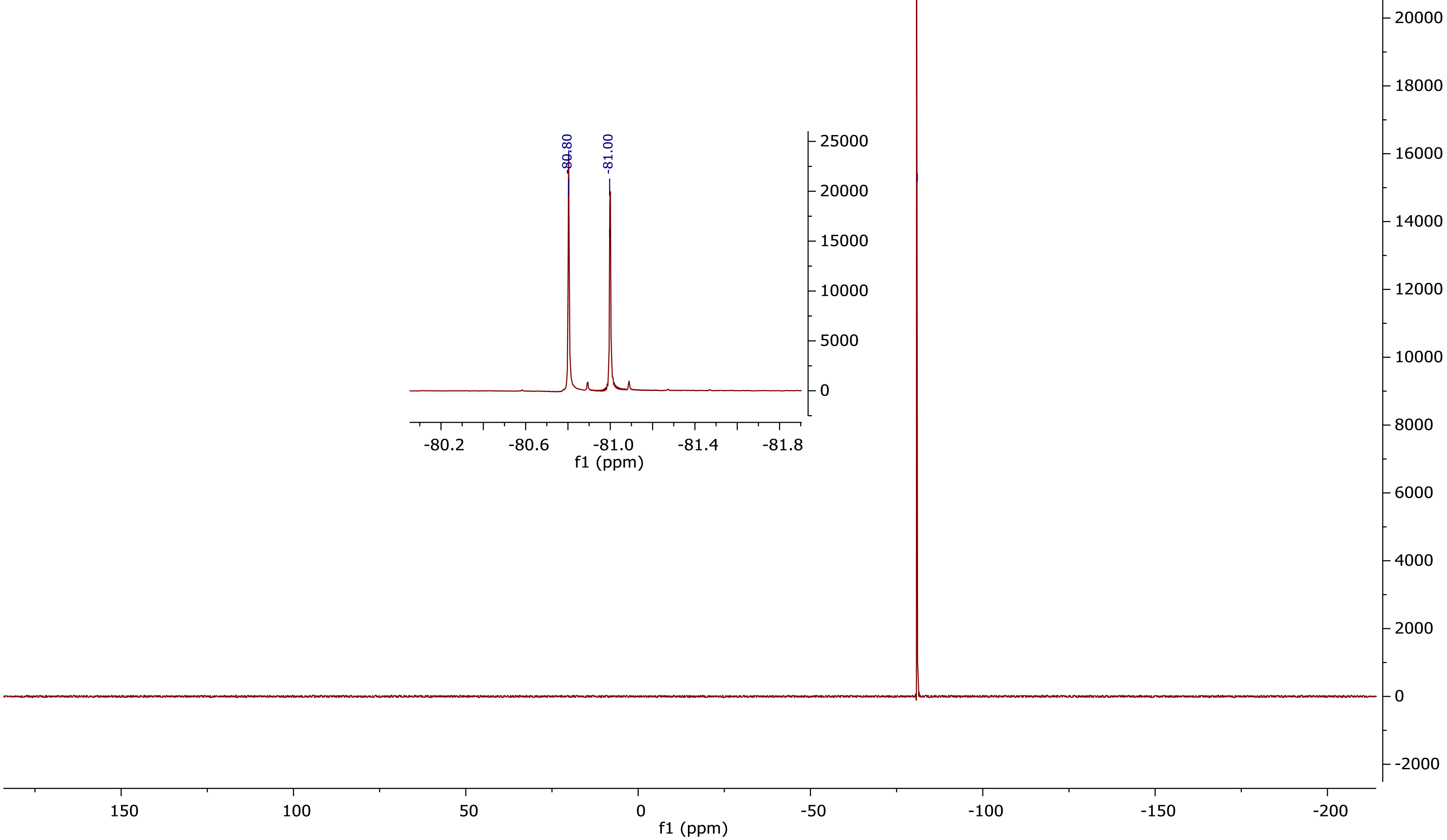

$^1$H NMR (500 MHz, CDCl$_3$) (nementin-12-15)

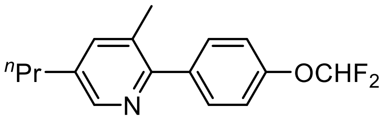


**nementin-12-15**

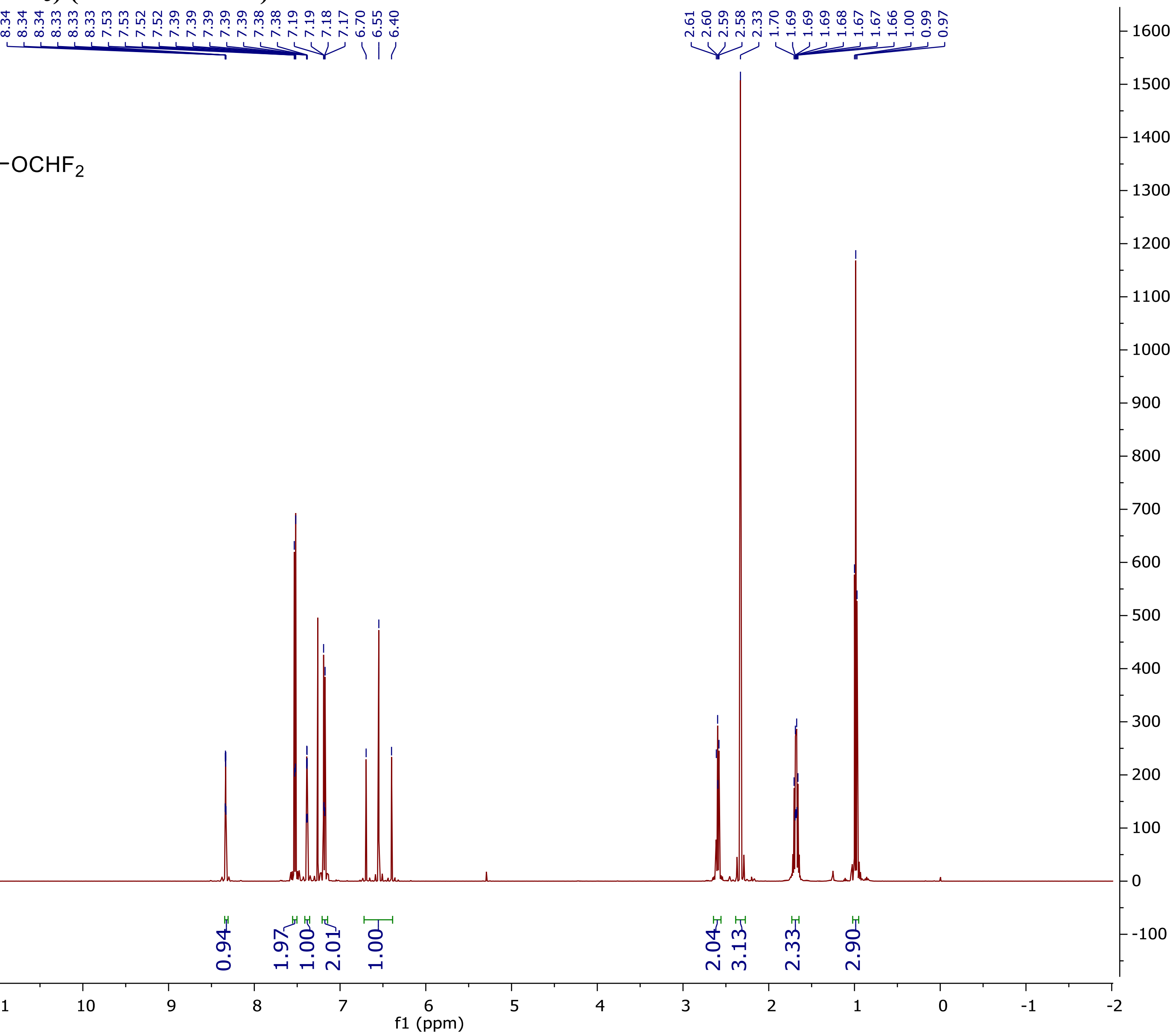

$^{13}$C NMR (125 MHz, $CDCl_3$)

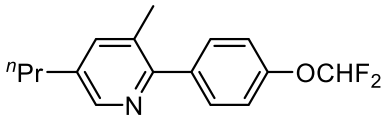


**nementin-12-15**

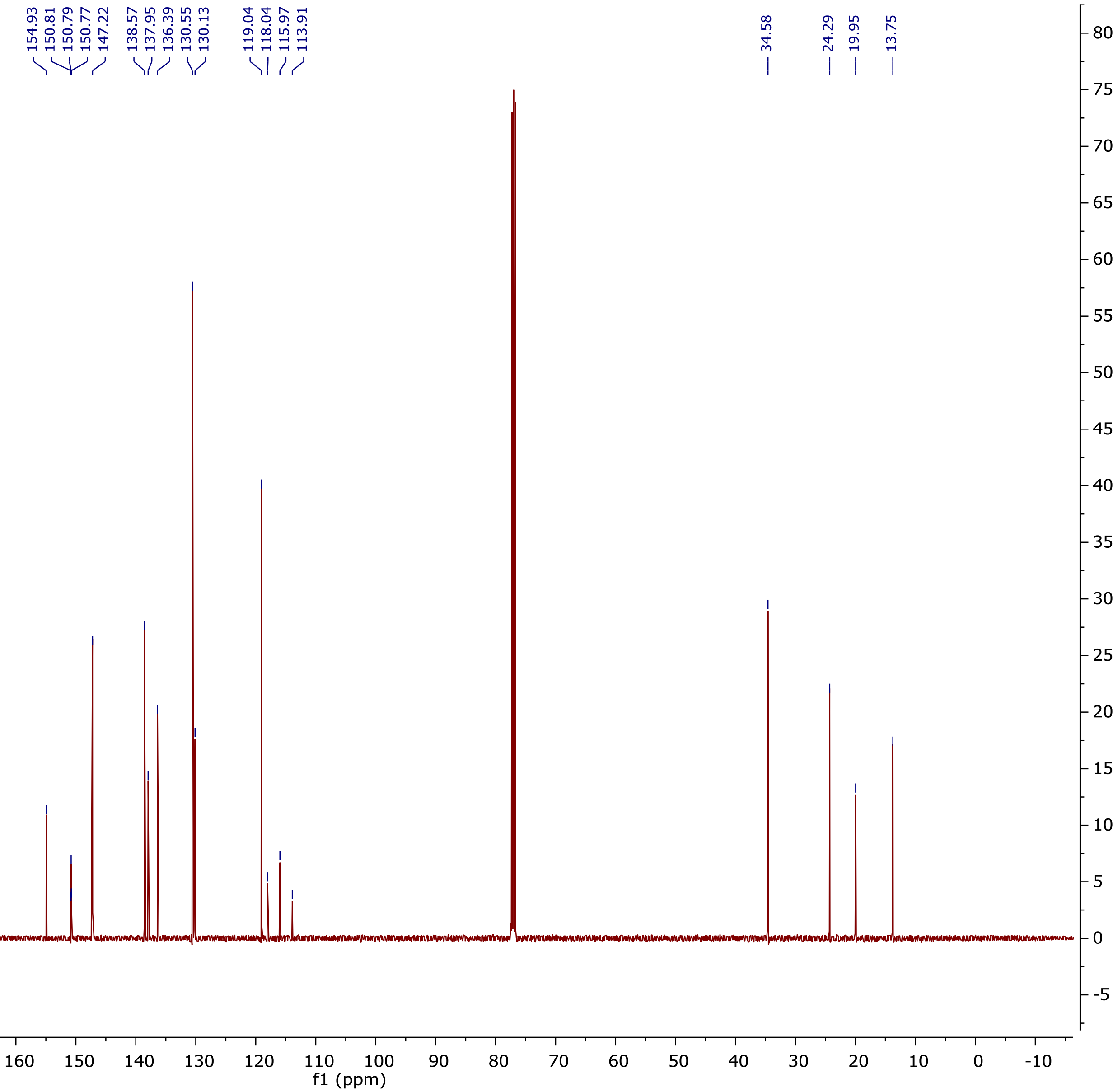

$^{19}$F NMR (375 MHz, $CDCl_3$)

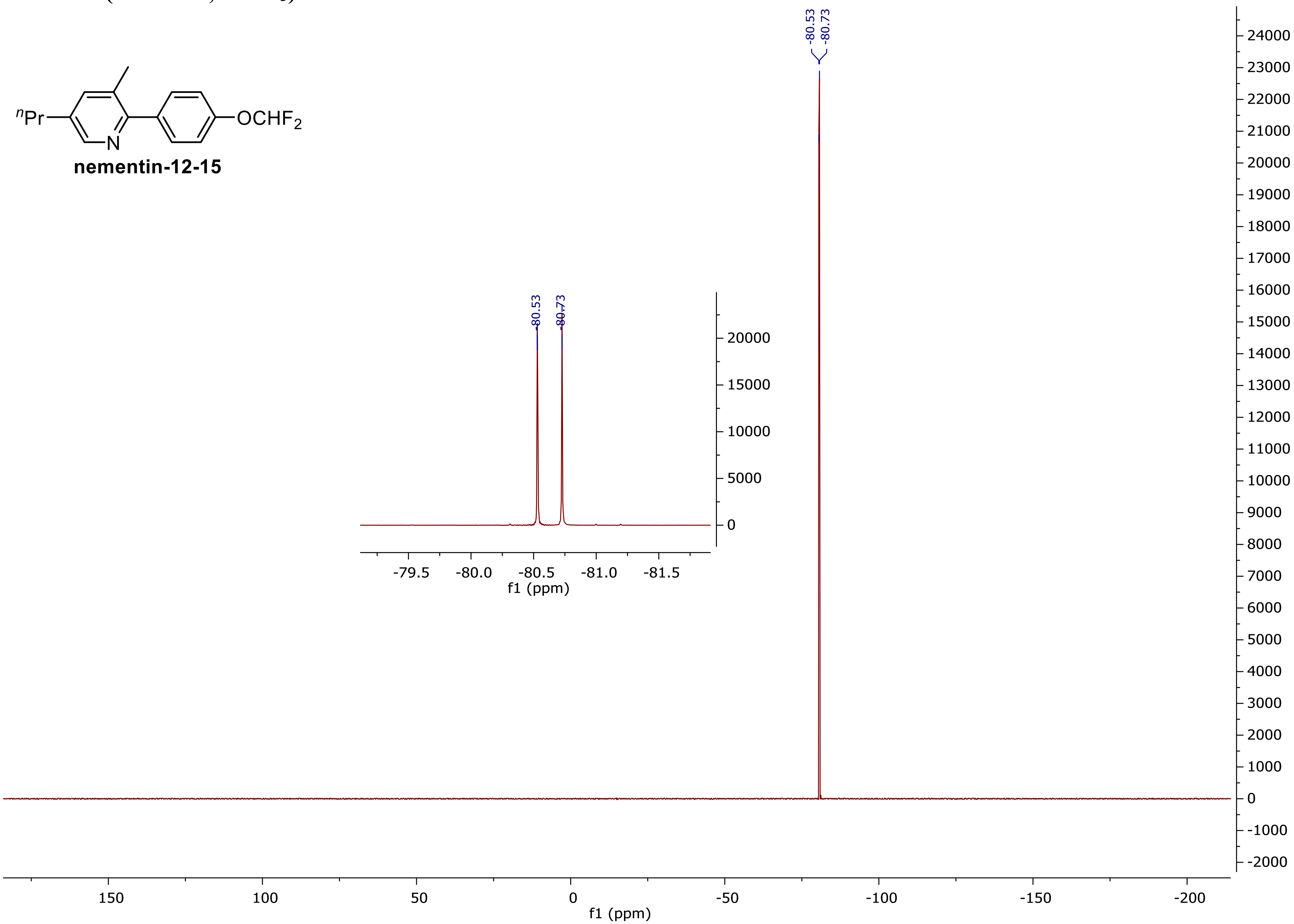

### $^{1}$H NMR (500 MHz, $CDCl_3$) (nementin-12-17)

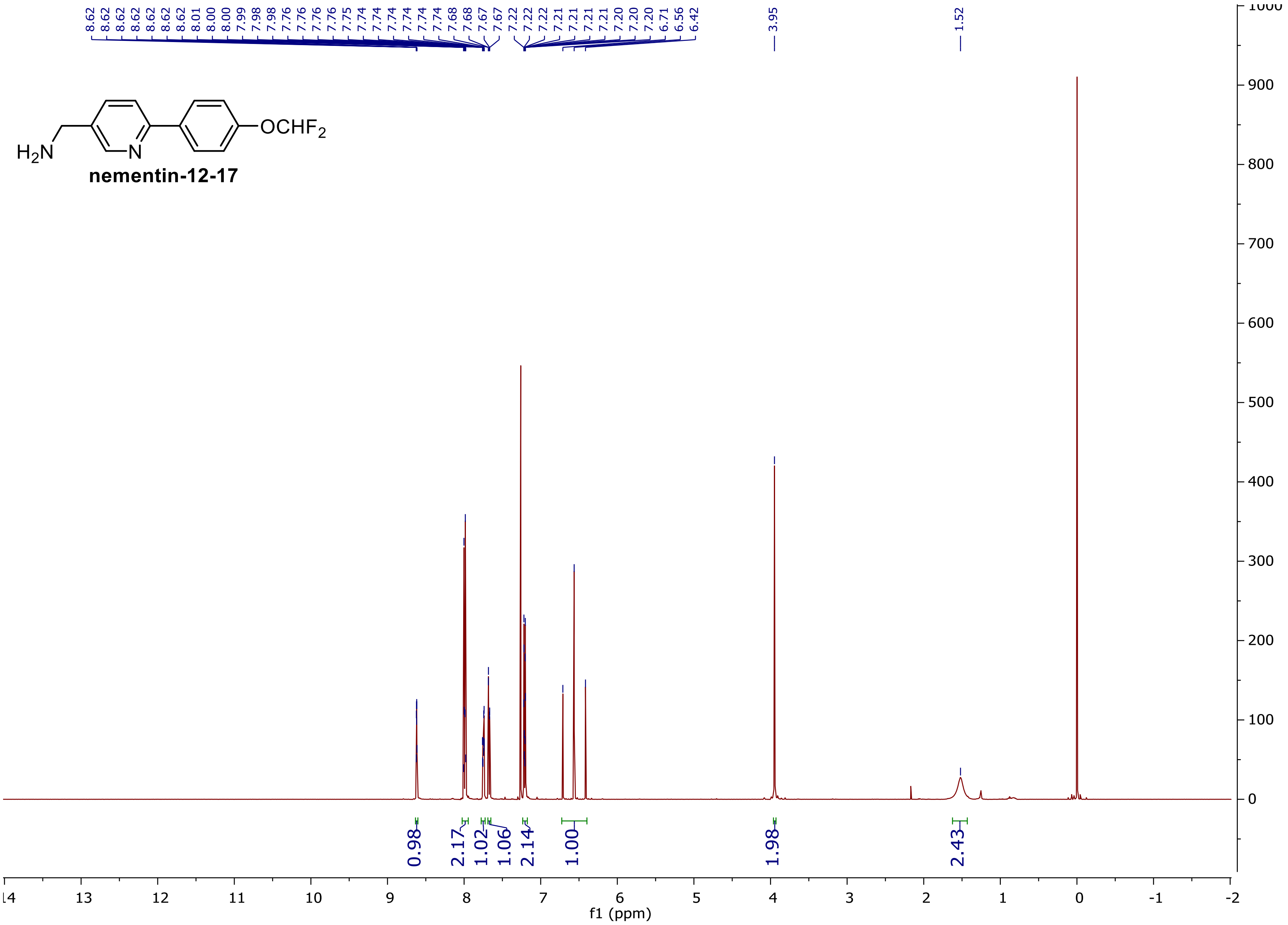

### $^{13}C$ NMR (125 MHz, $CDCl_3$)

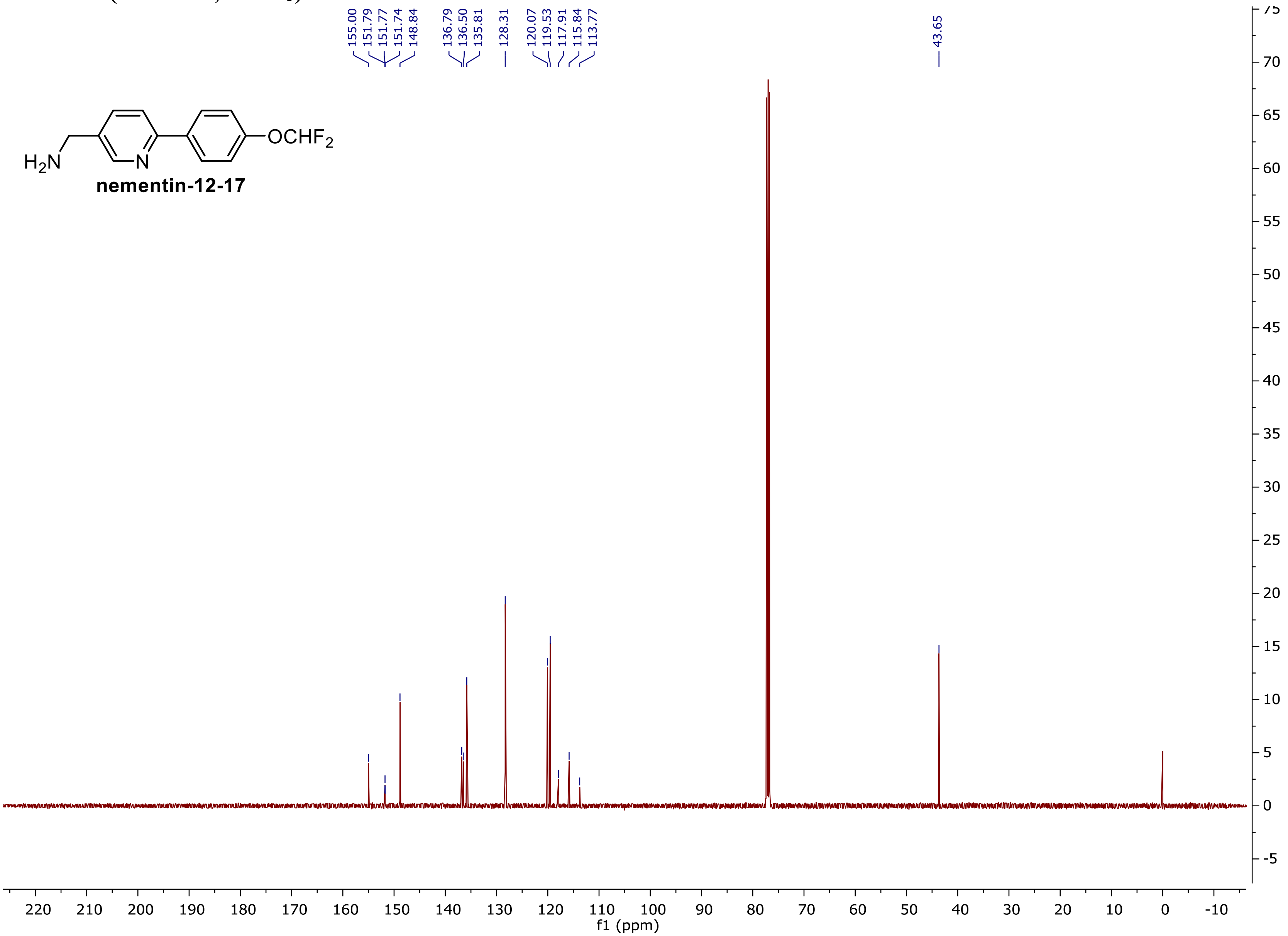

$^{19}F$ NMR (375 MHz, $CDCl_3$)

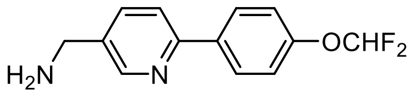


**nementin-12-17**

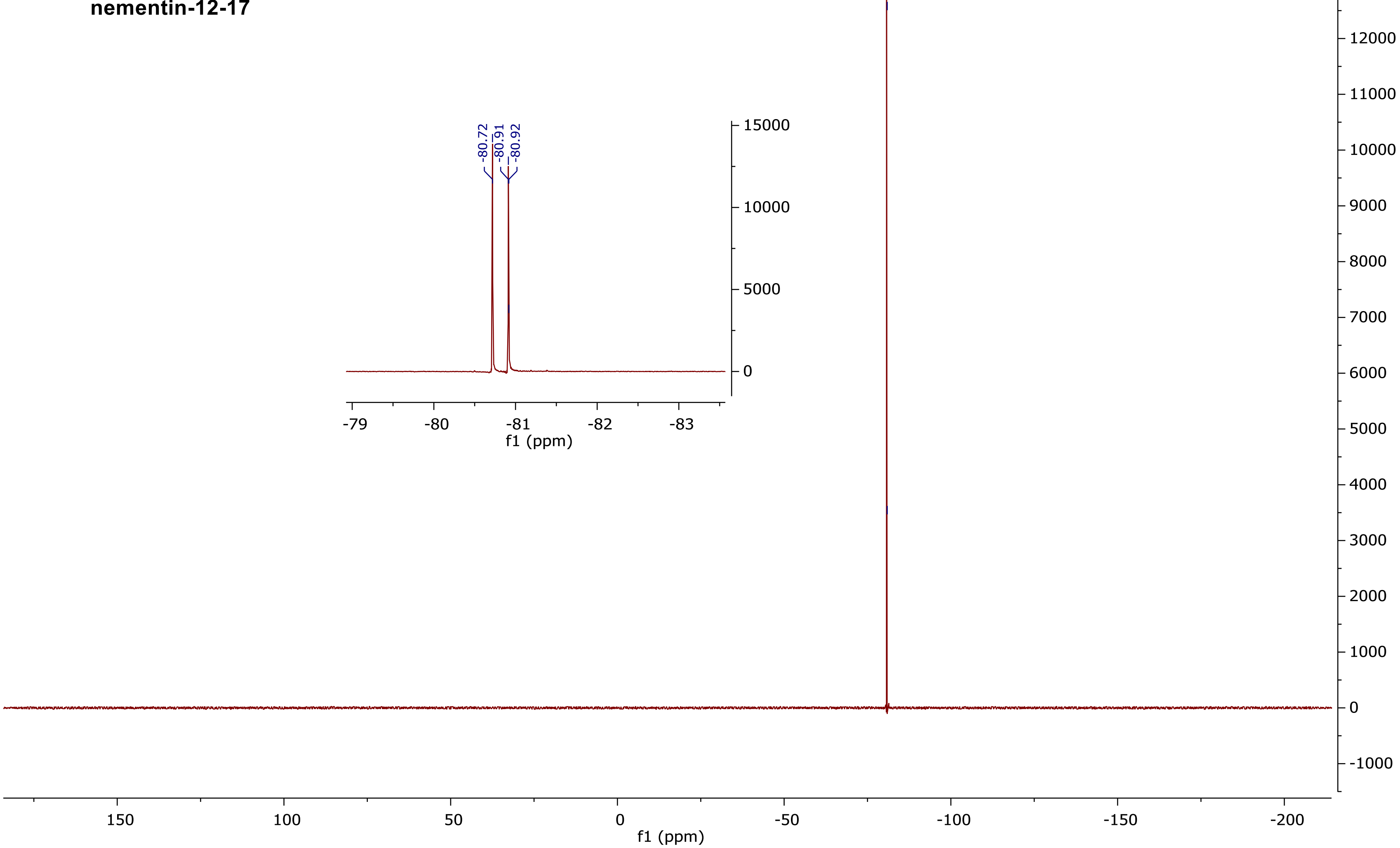

### $^1$H NMR (500 MHz, $CDCl_3$) (nementin-12-18)

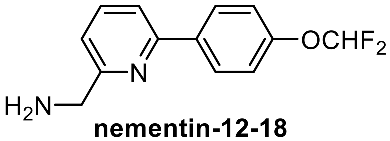

**nementin-12-18**

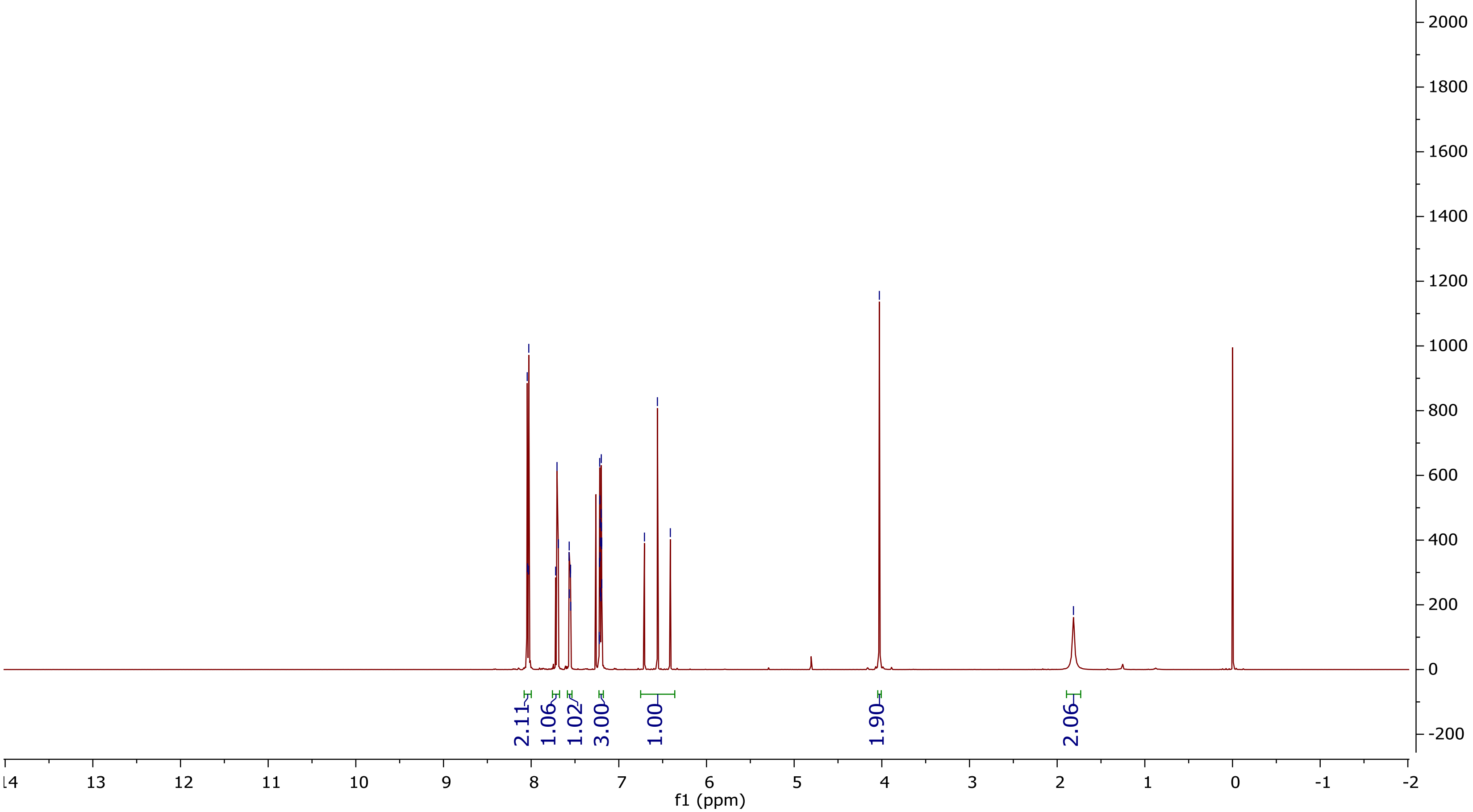

**$^{13}$C NMR (125 MHz, $CDCl_3$)**

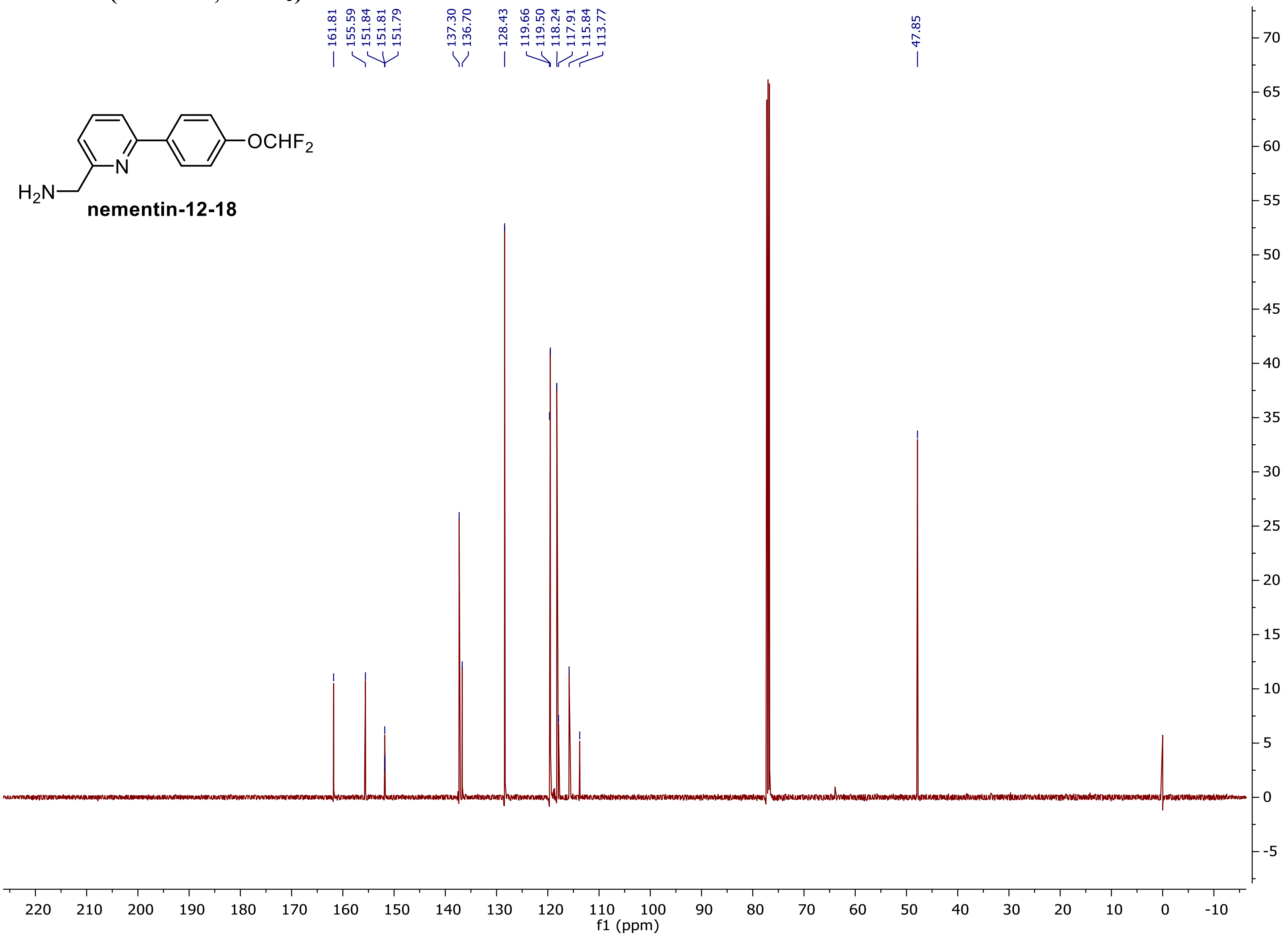

$^{19}F$ NMR (375 MHz, $CDCl_3$)

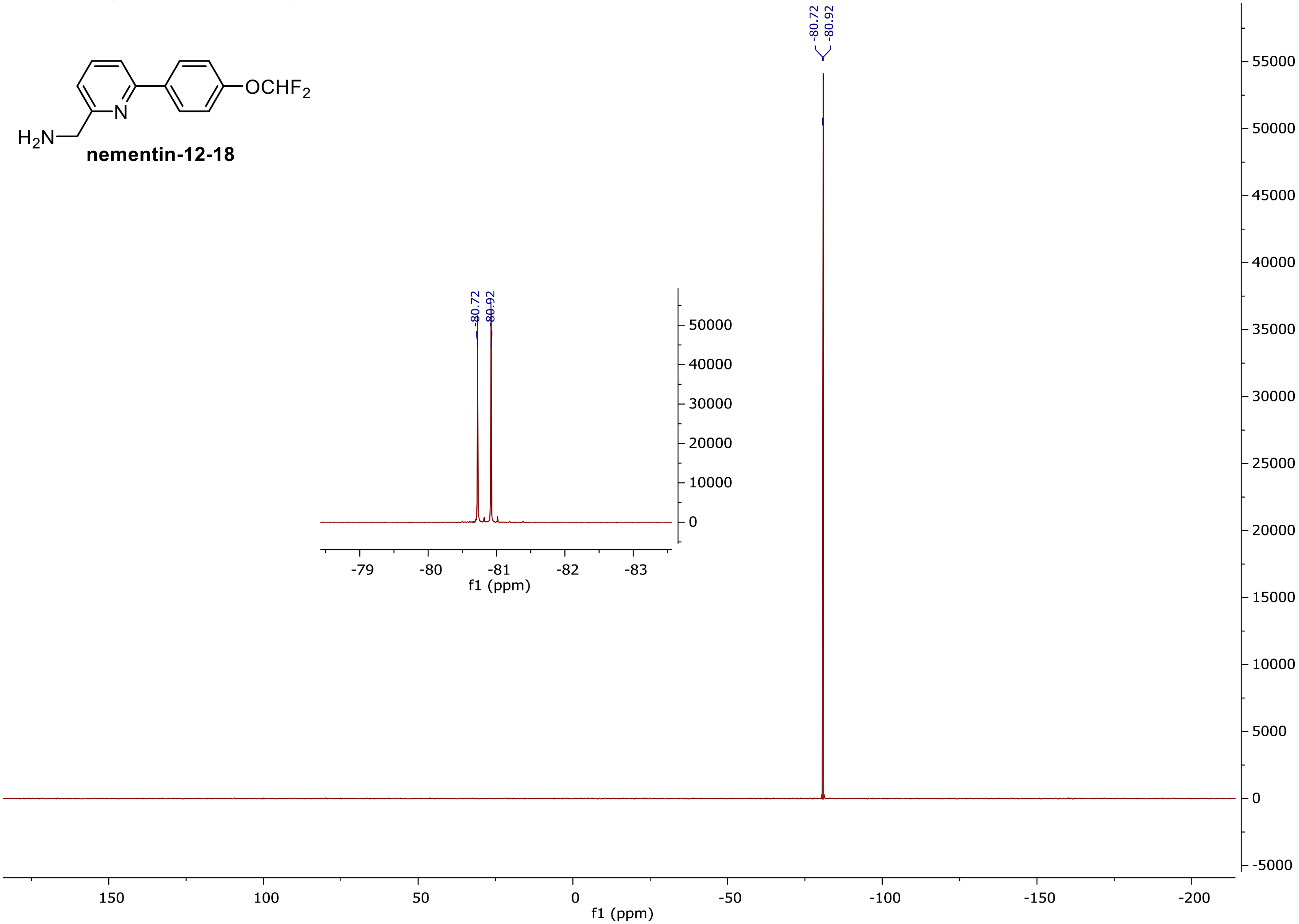


**nementin-12-18**

$^1$H NMR (500 MHz, $CDCl_3$) (nementin-12-20)

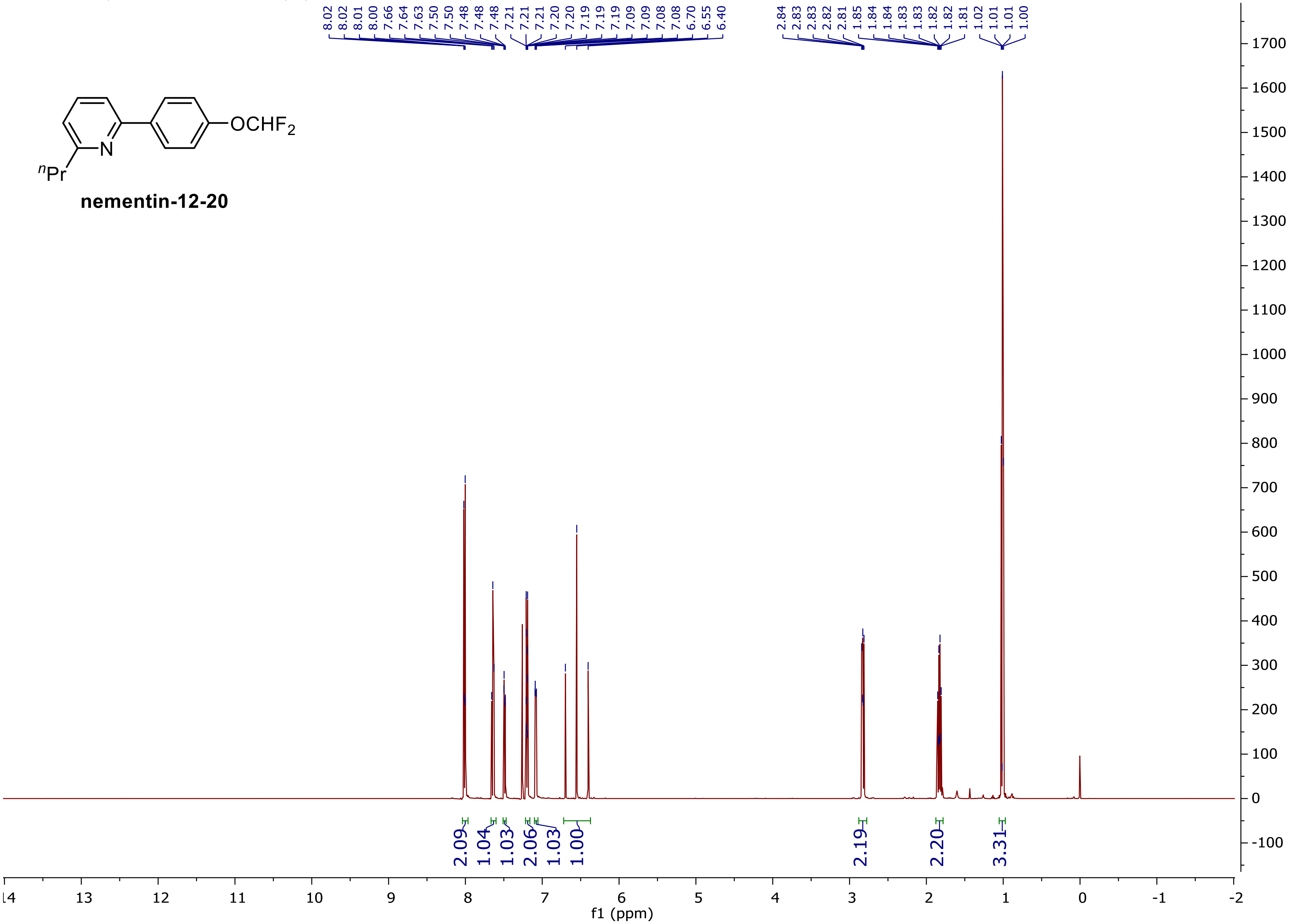

### $^{13}$C NMR (125 MHz, CDCl$_3$)

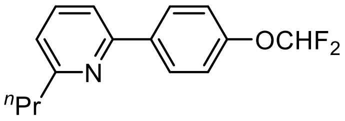


**nementin-12-20**

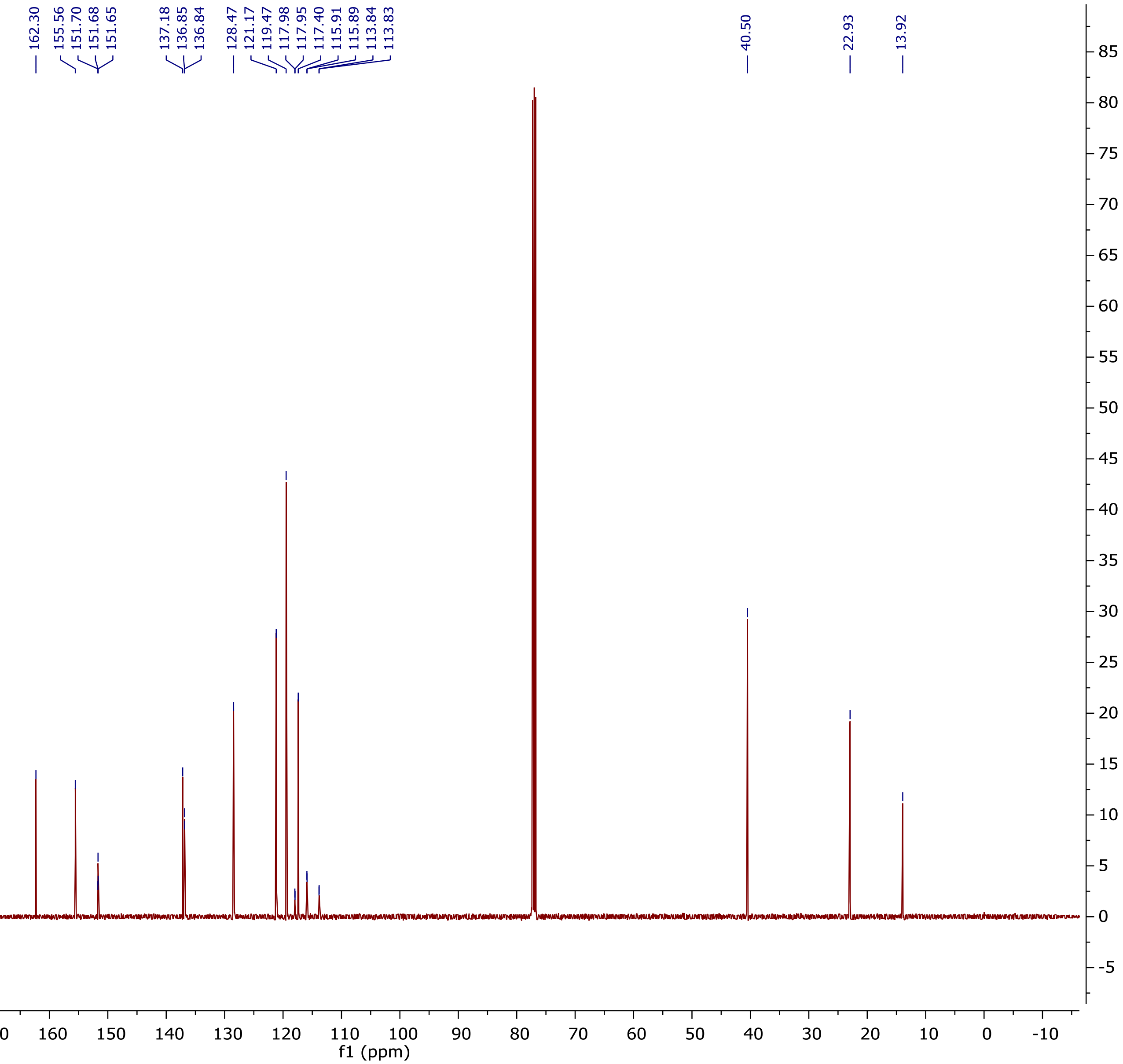

**$^{19}$F NMR (375 MHz, $CDCl_3$)**

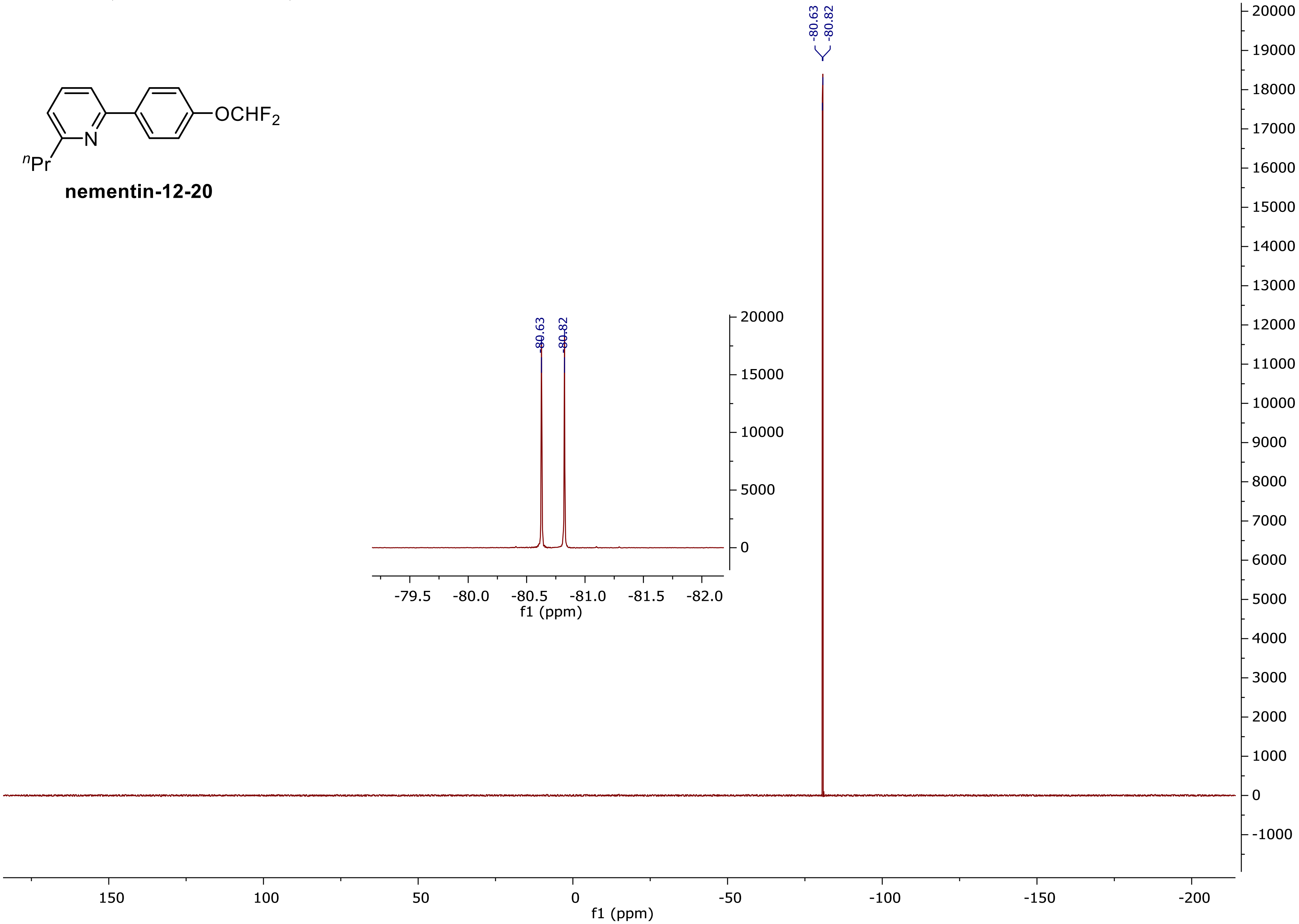


**nementin-12-20**

**$^{1}H$ NMR (500 MHz, $CDCl_3$) (nementin-12-21)**

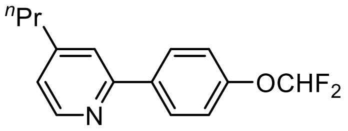


**nementin-12-21**

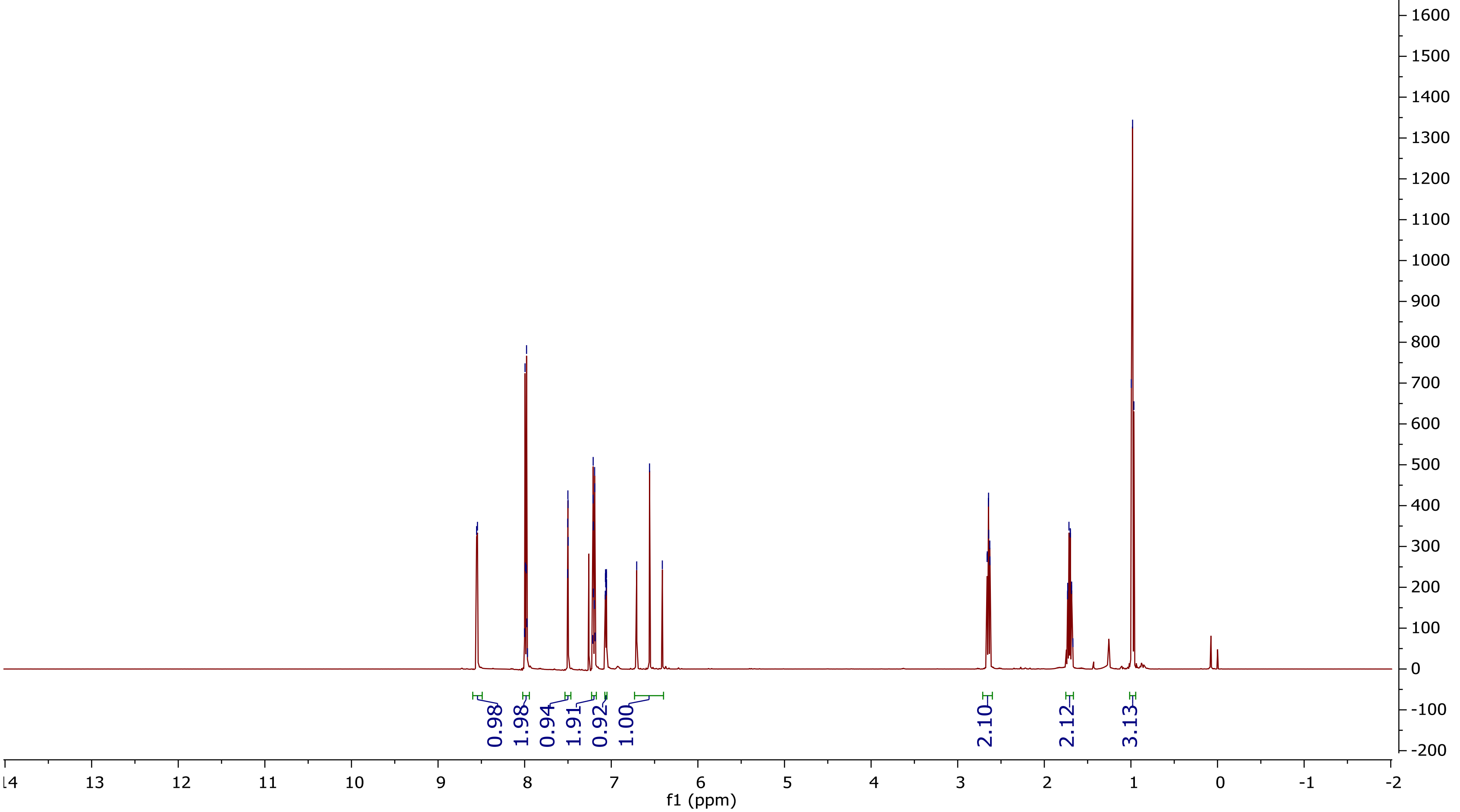

$^{13}$C NMR (125 MHz, $CDCl_3$)

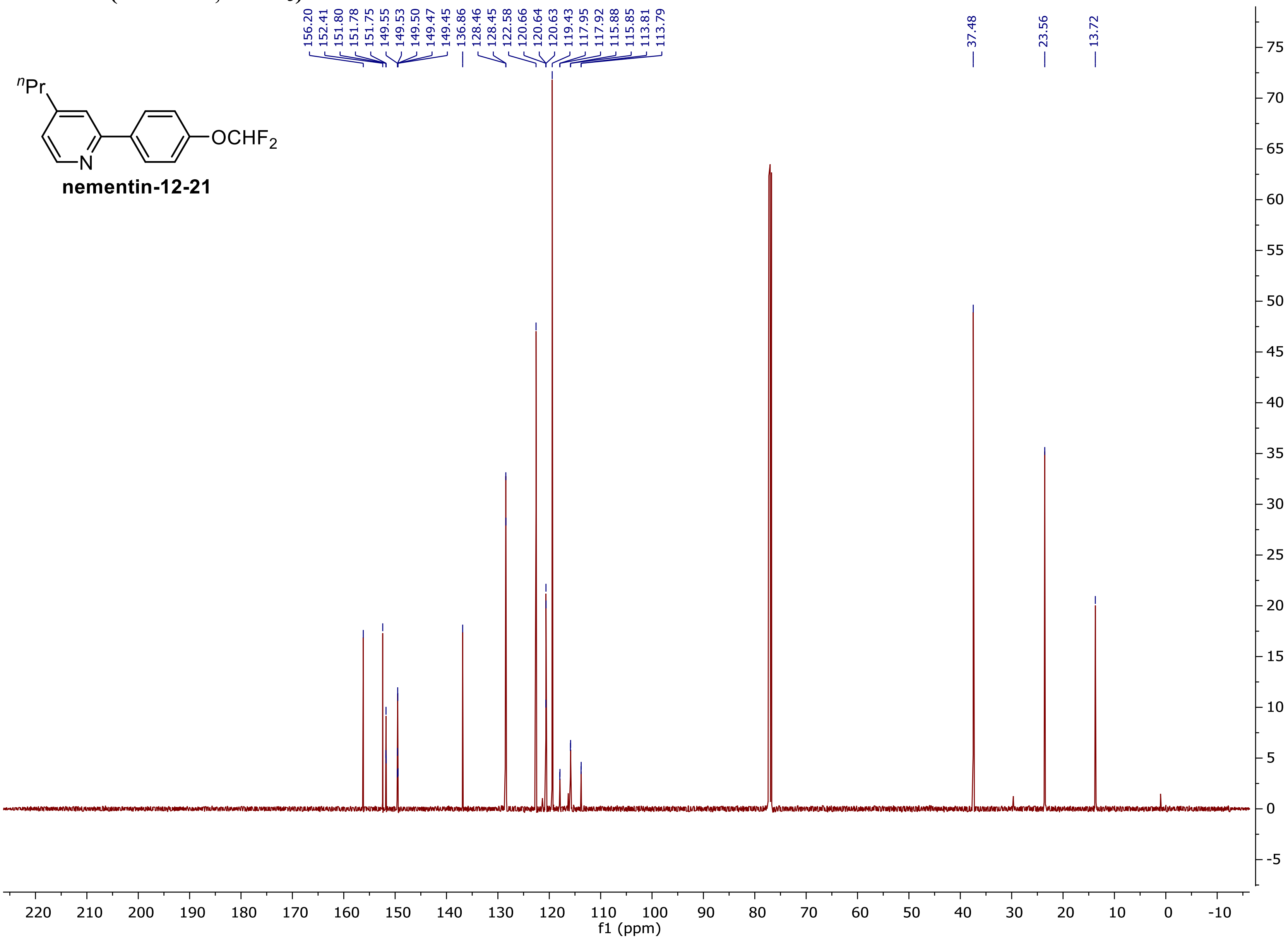

### [19]F NMR (375 MHz, $CDCl_3$)

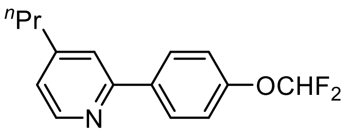


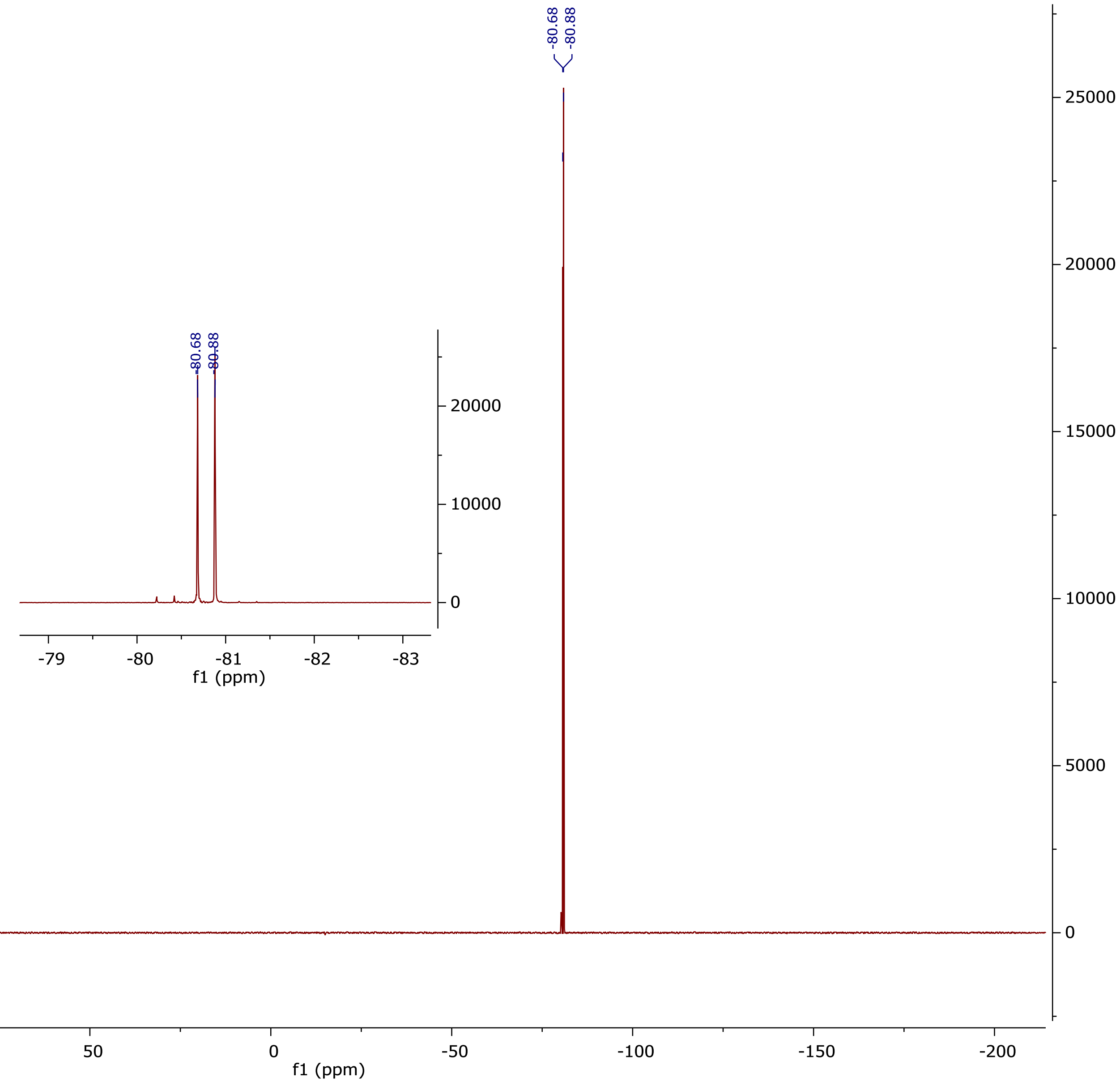

$^{1}H$ NMR (500 MHz, $CDCl_3$) (nementin-12-22)

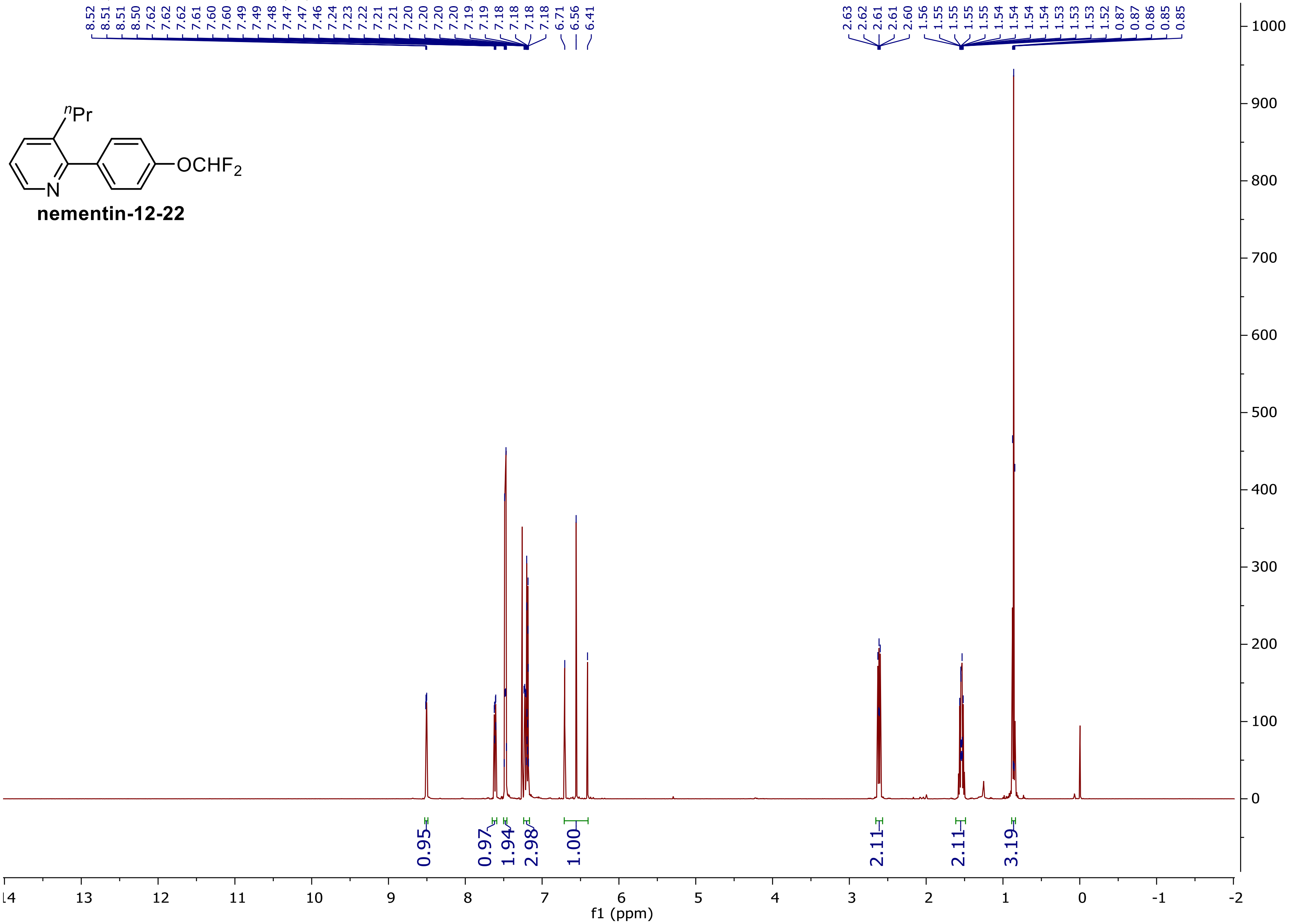

$^{13}$C NMR (125 MHz, CDCl$_3$)

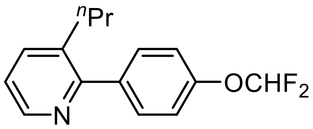


nementin-12-22

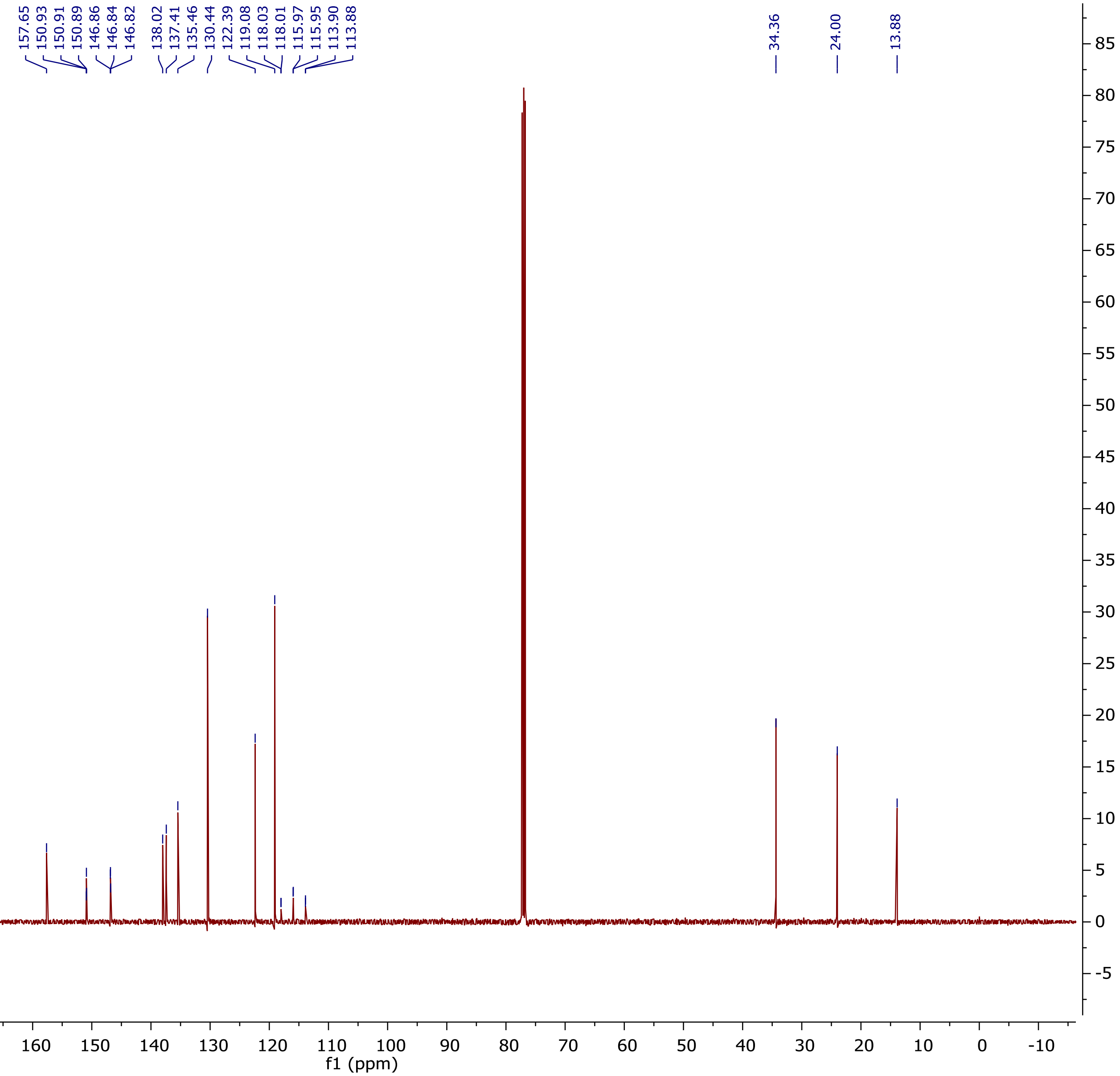

**$^{19}$F NMR (375 MHz, $CDCl_3$)**

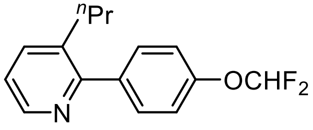


**nementin-12-22**

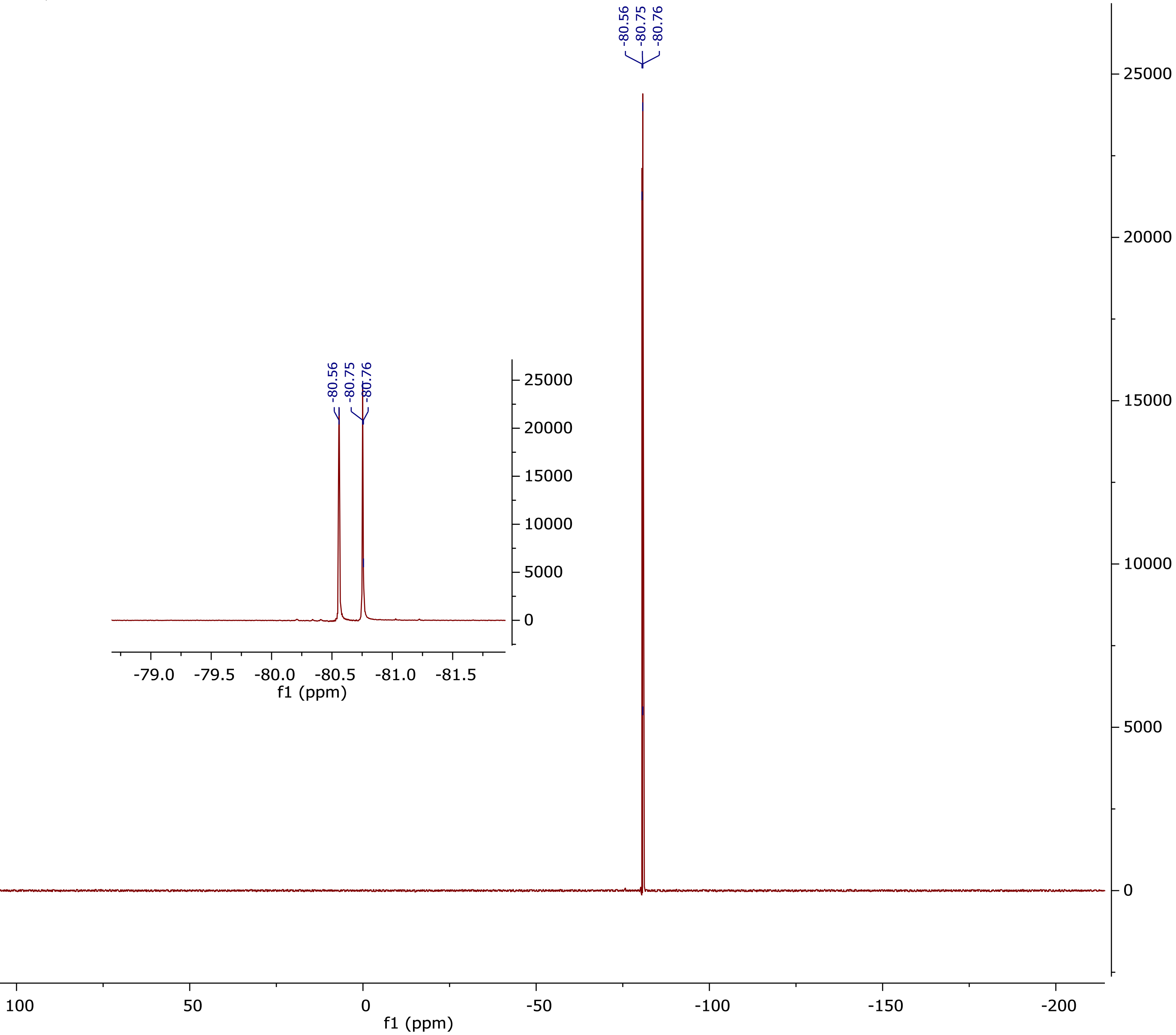

### $^1$H NMR (500 MHz, $CDCl_3$) (nementin-12-23)

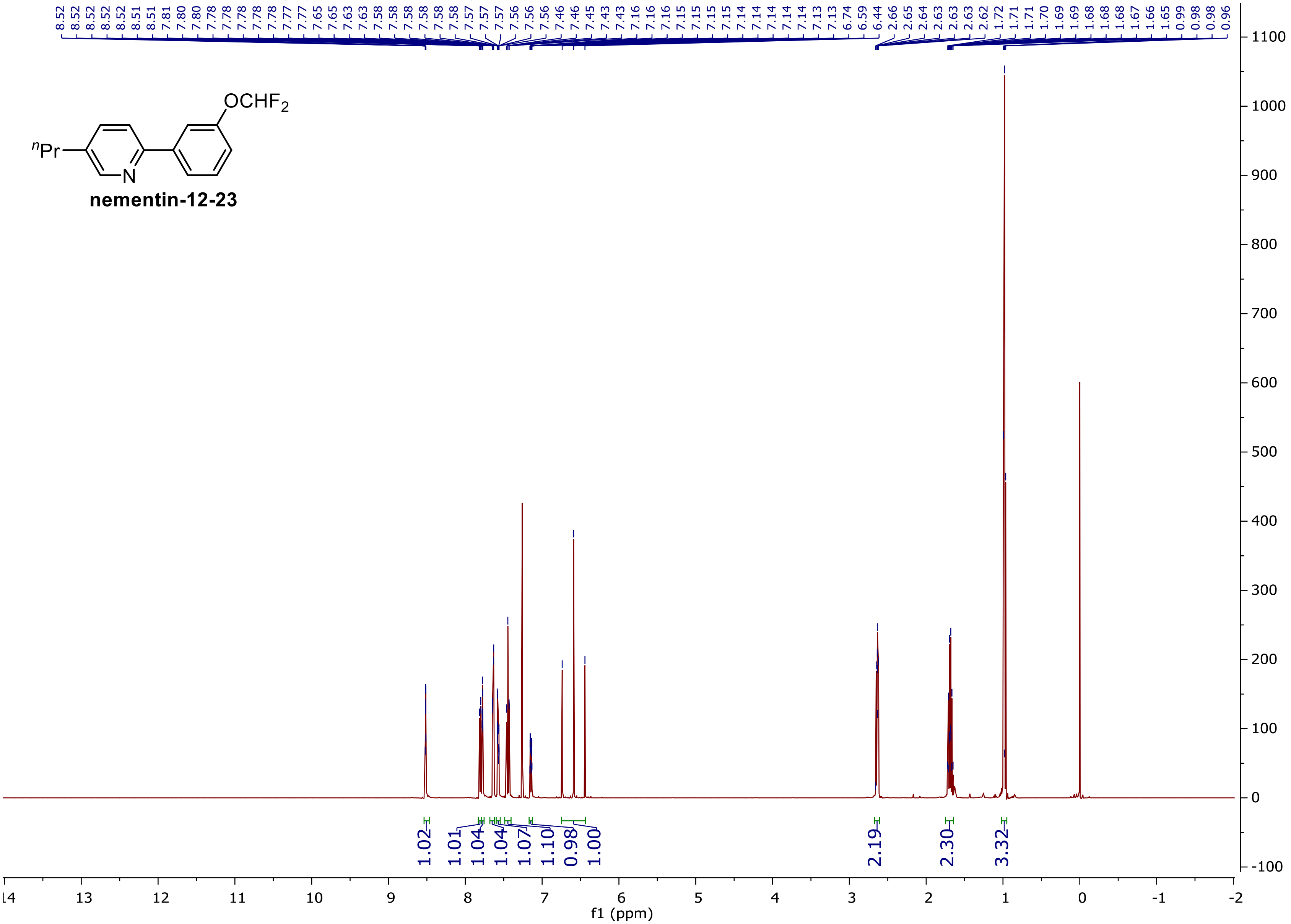

### $^{13}$C NMR (125 MHz, CDCl$_3$)

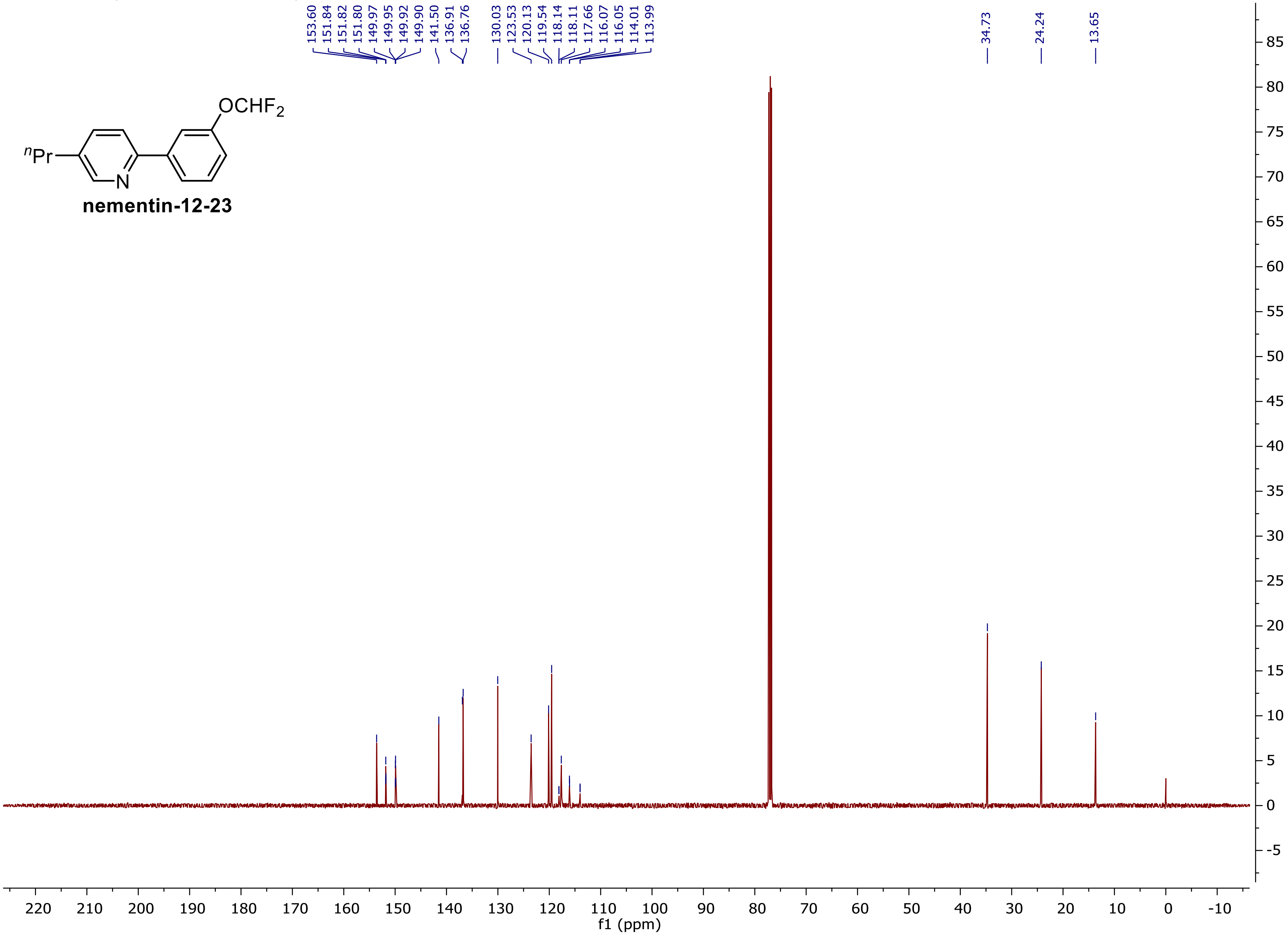


**nementin-12-23**

**$^{19}$F NMR (375 MHz, CDCl$_3$)**

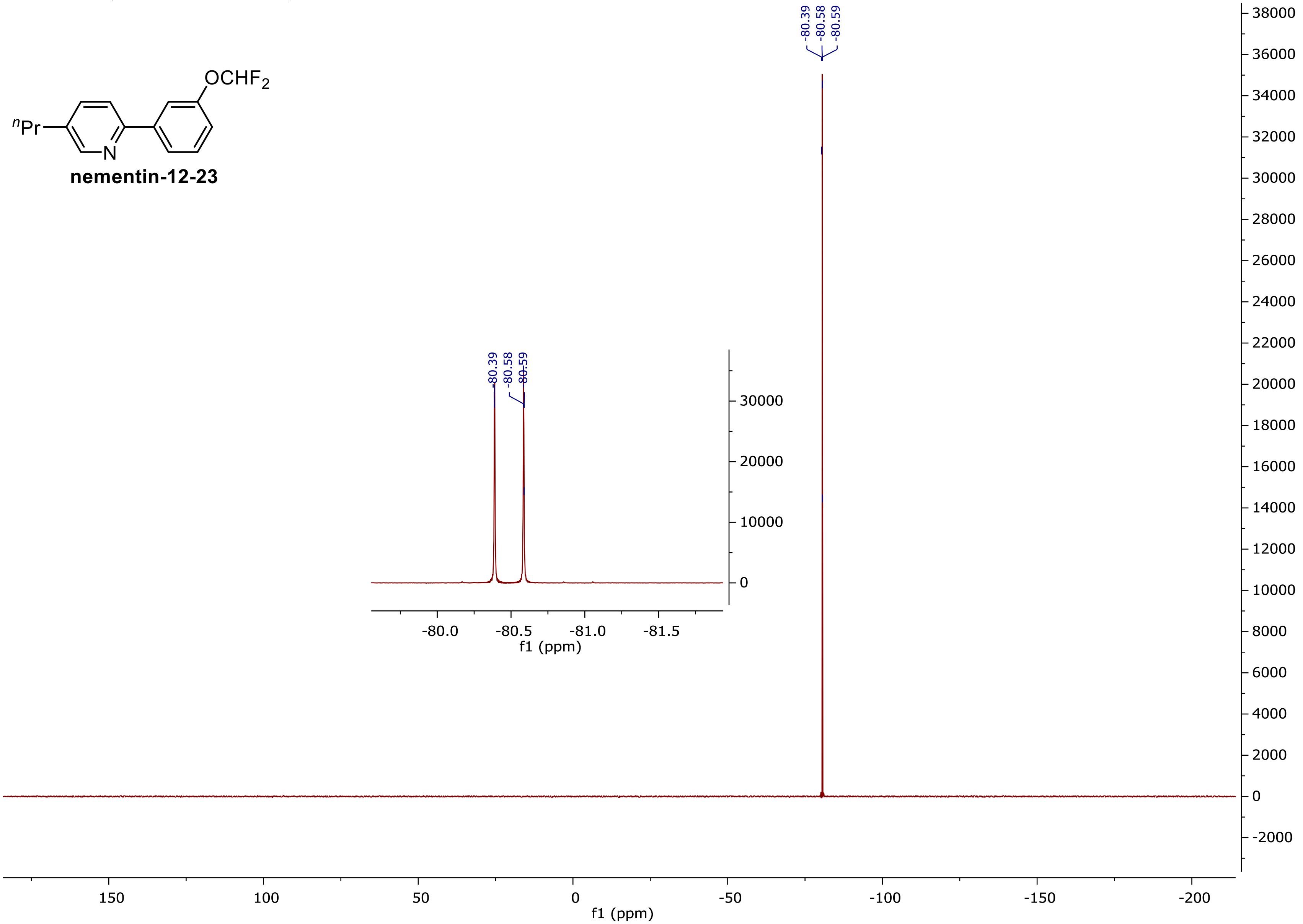

**$^1$H NMR (500 MHz, CDCl$_3$) (nementin-12-24)**

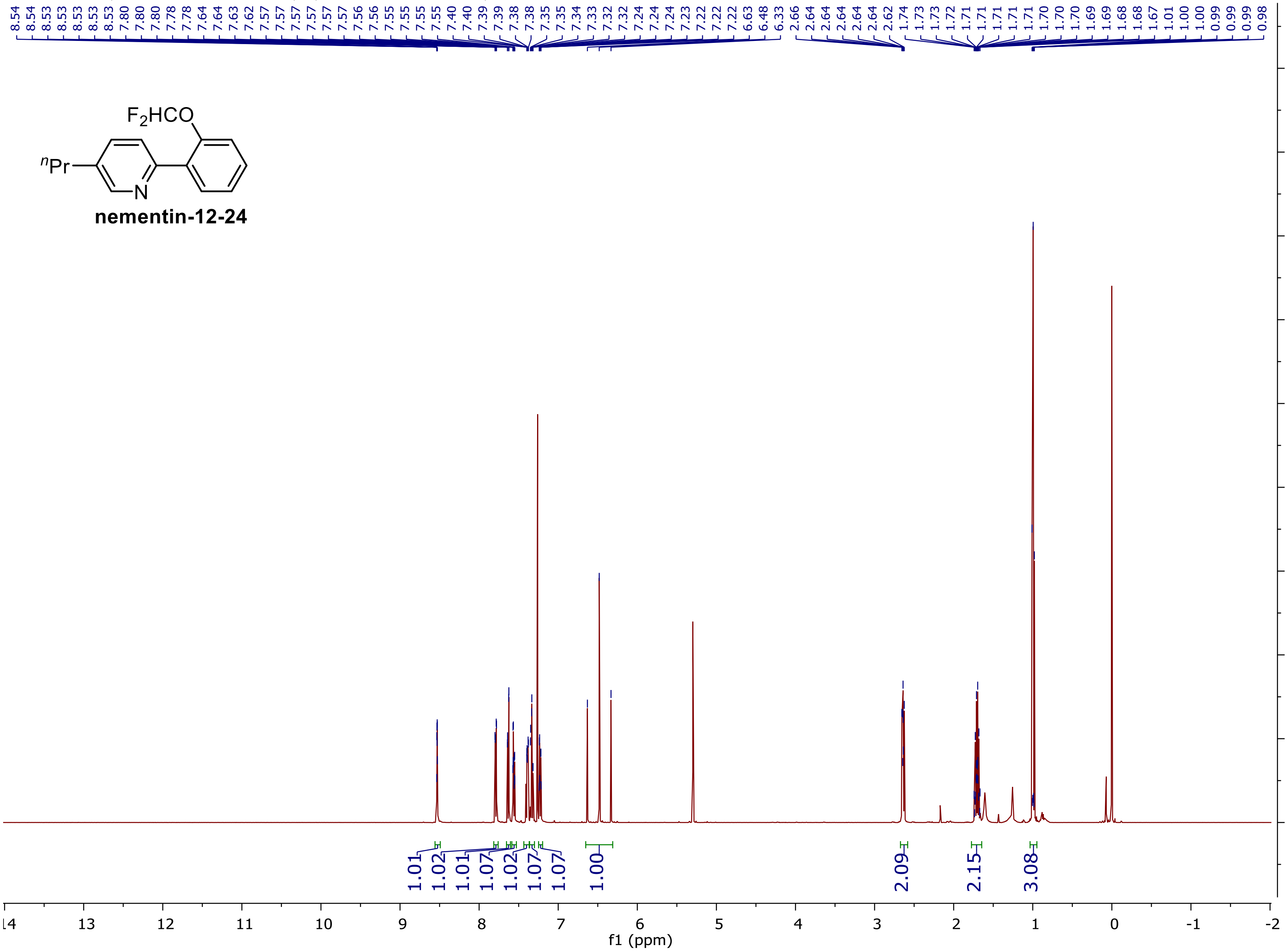

**$^{13}$C NMR (125 MHz, CDCl$_3$)**

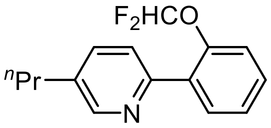


**nementin-12-24**

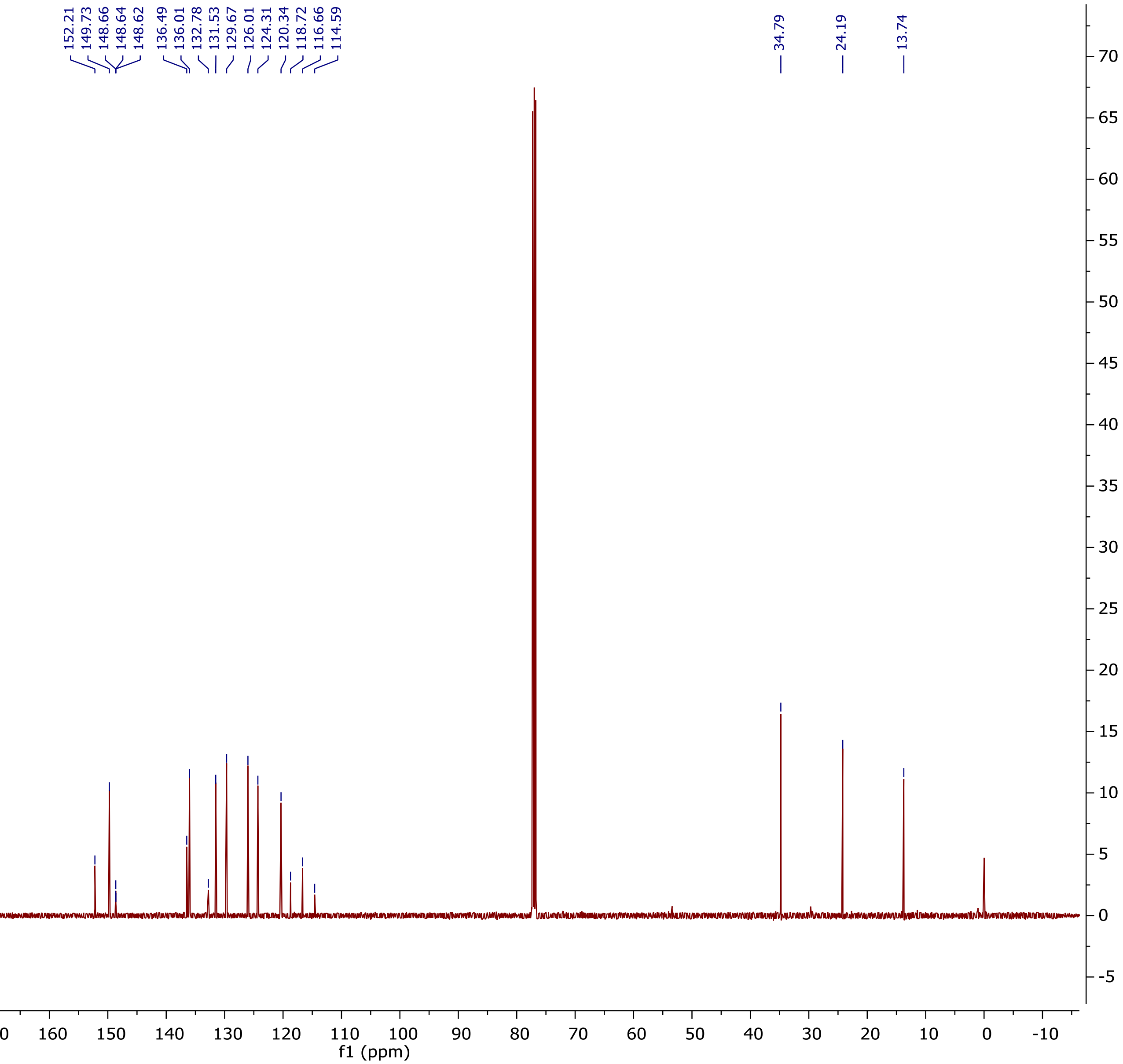

$^{19}$F NMR (375 MHz, $CDCl_3$)

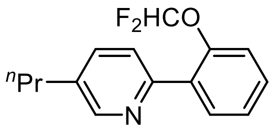


**nementin-12-24**

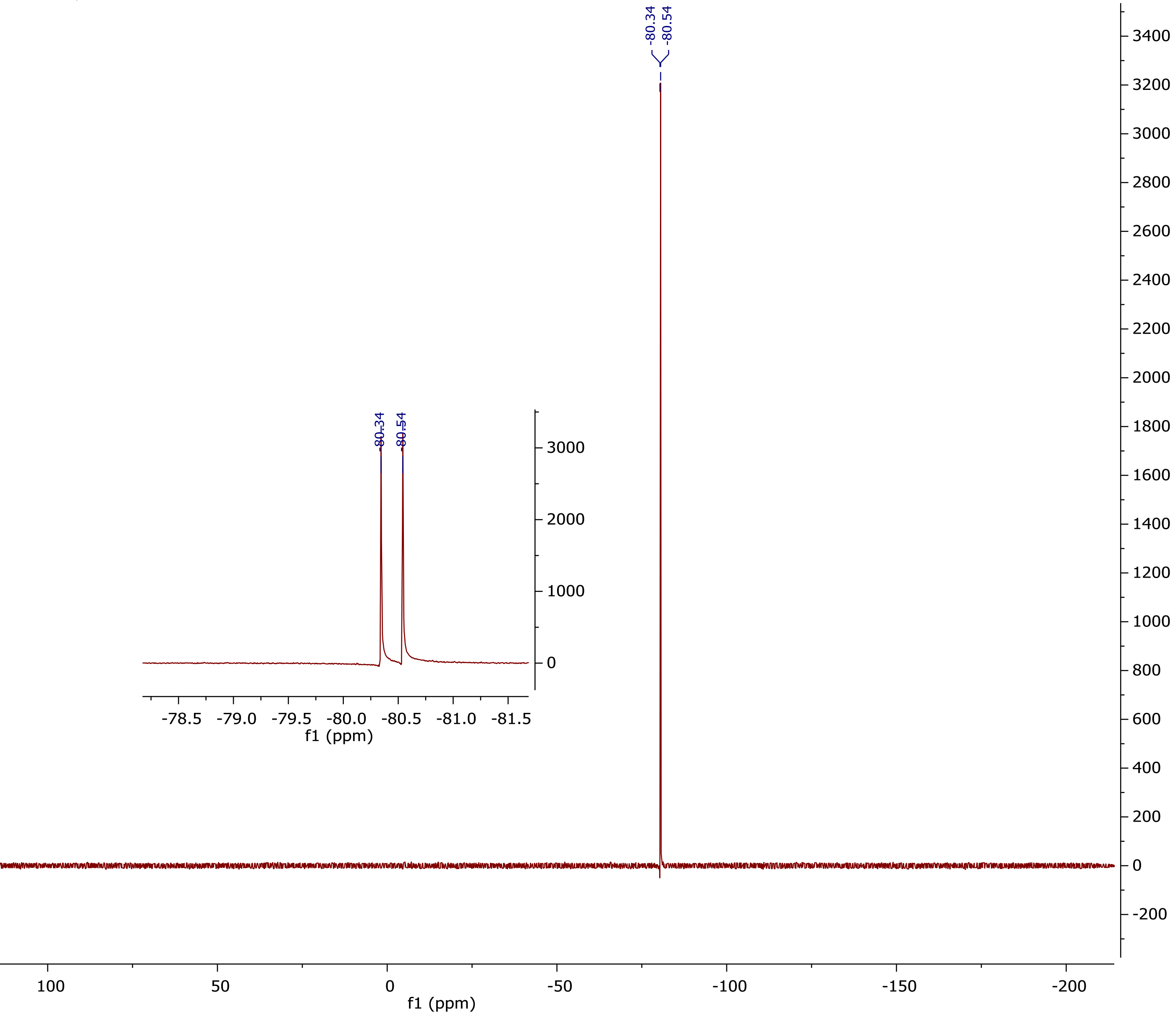

**$^1$H NMR (500 MHz, CDCl$_3$) (nementin-12-25)**

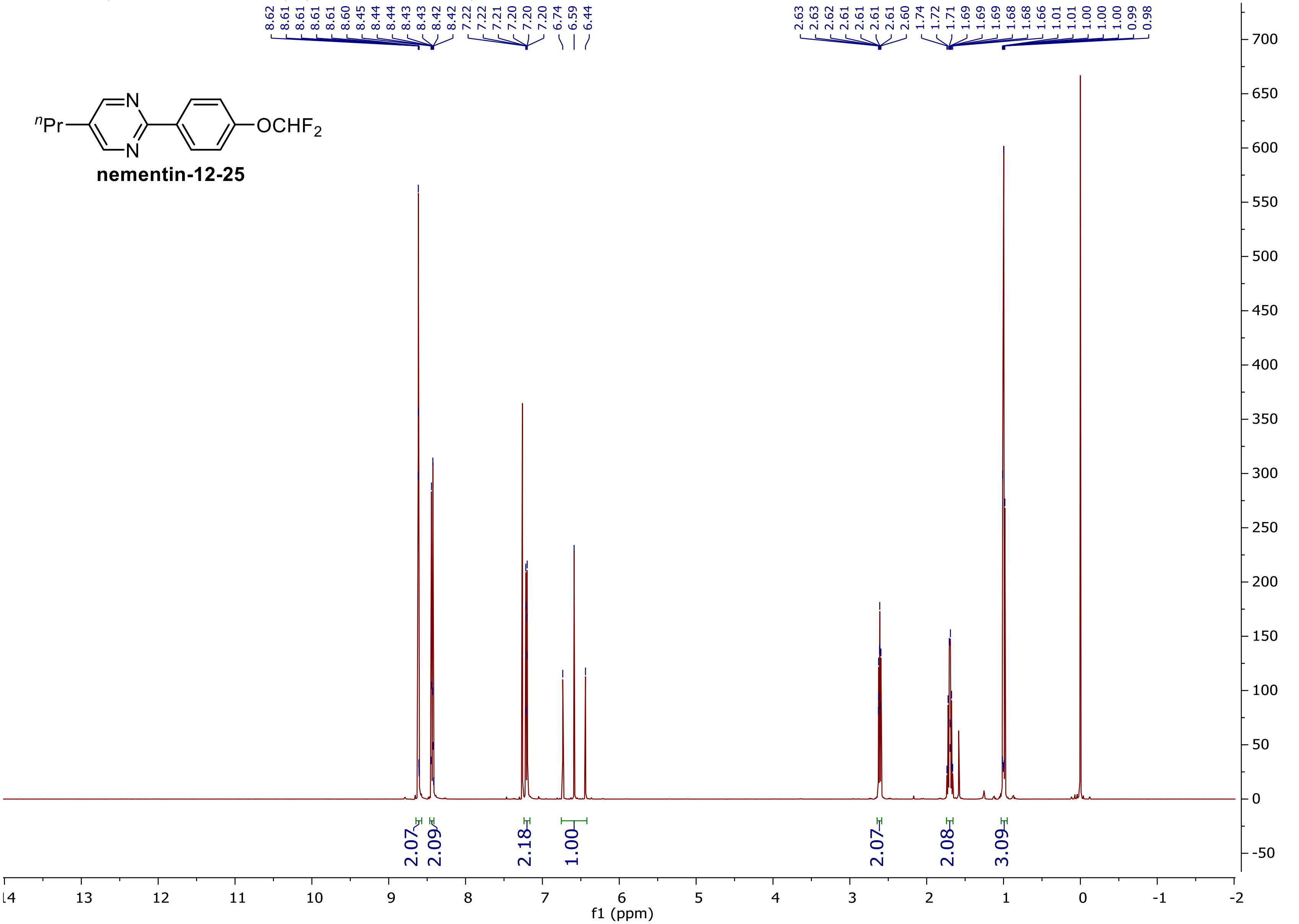

**$^{13}$C NMR (125 MHz, CDCl$_3$)**

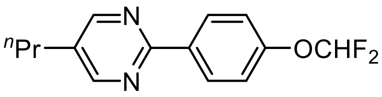


**nementin-12-25**

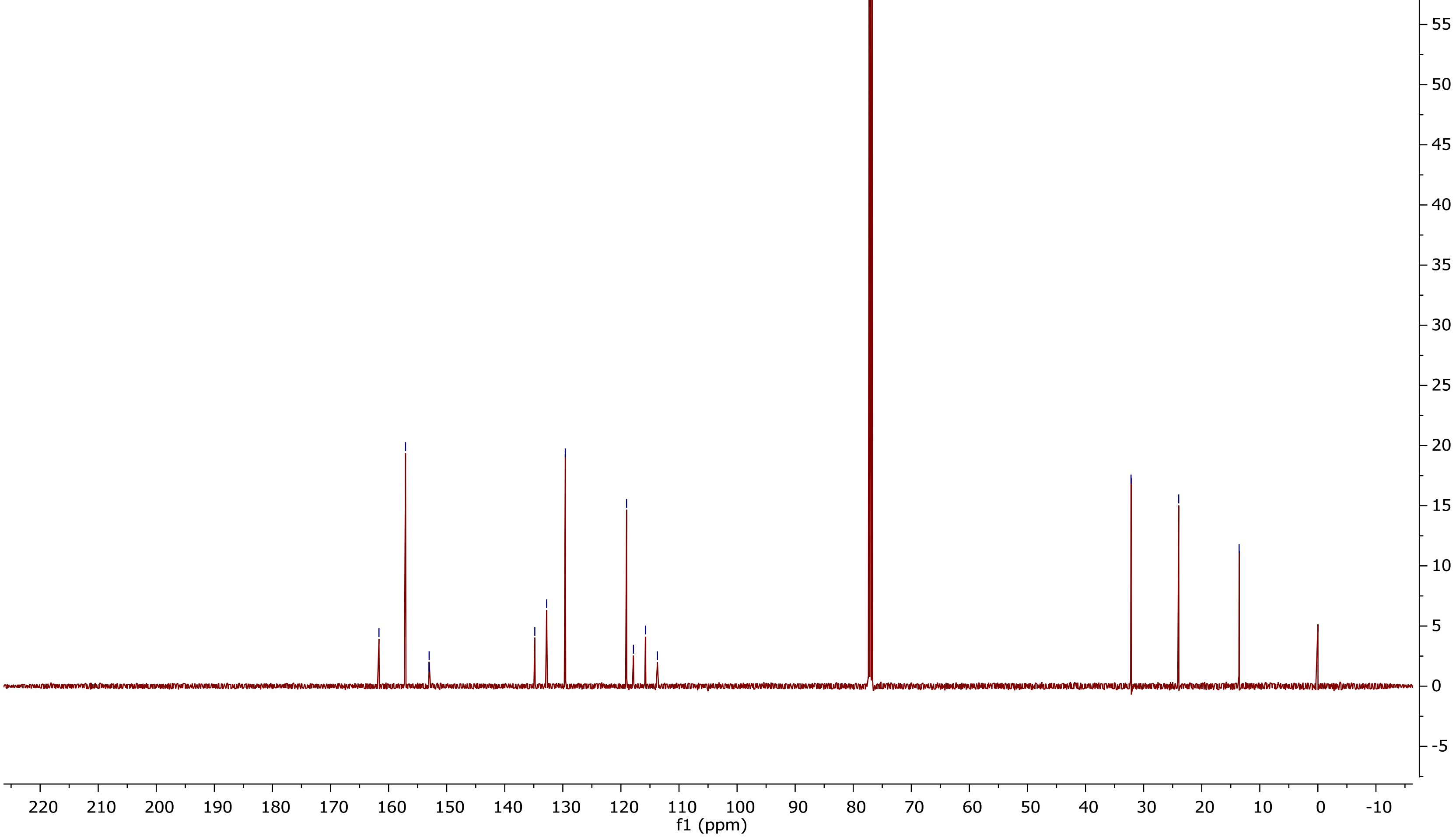

**$^{19}$F NMR (375 MHz, $CDCl_3$)**

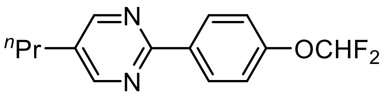

**nementin-12-25**

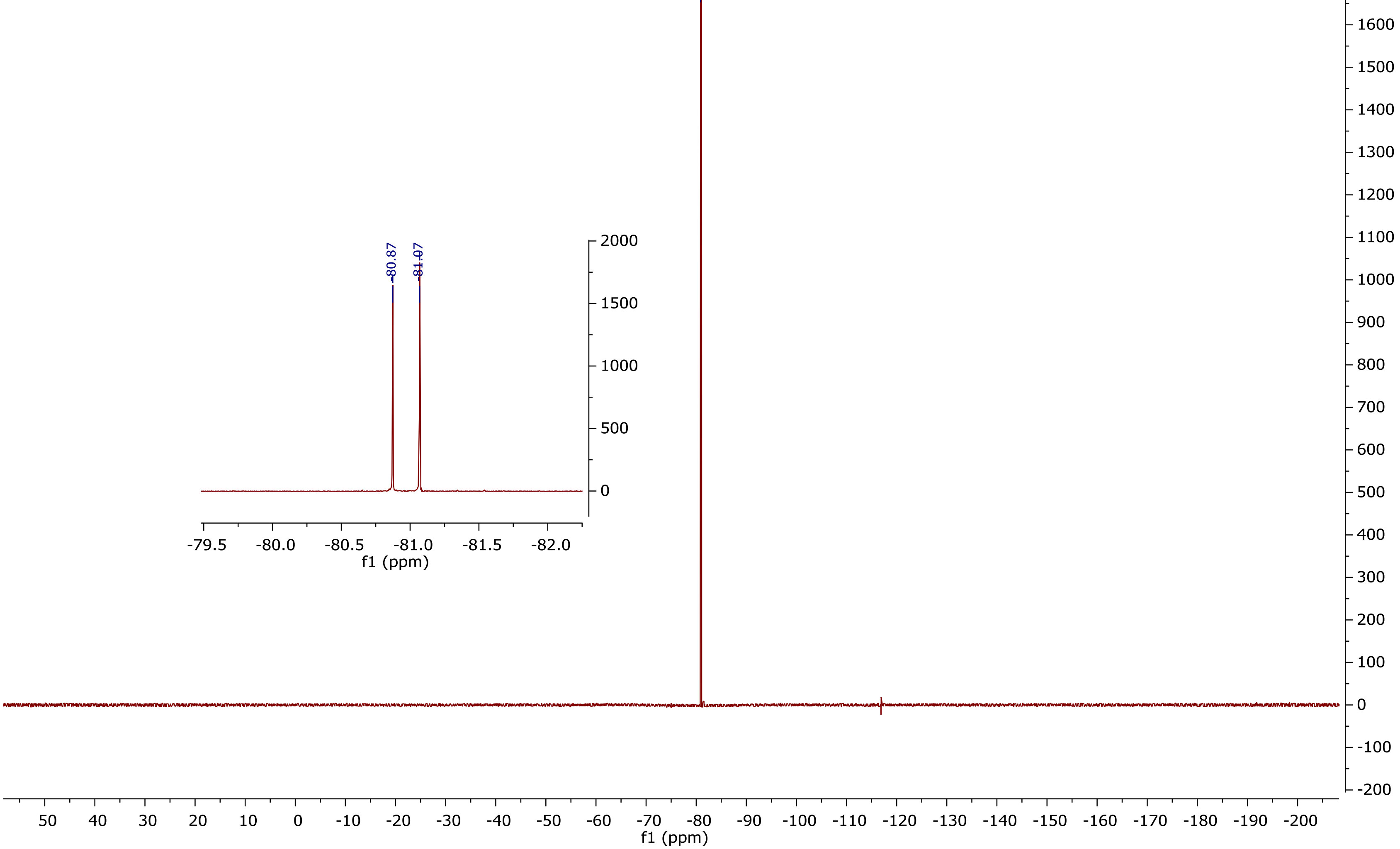

**[1]H NMR (500 MHz, CDCl₃) (nementin-12-26)**

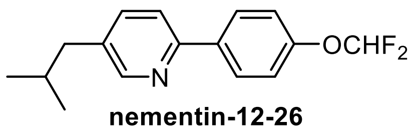


**nementin-12-26**

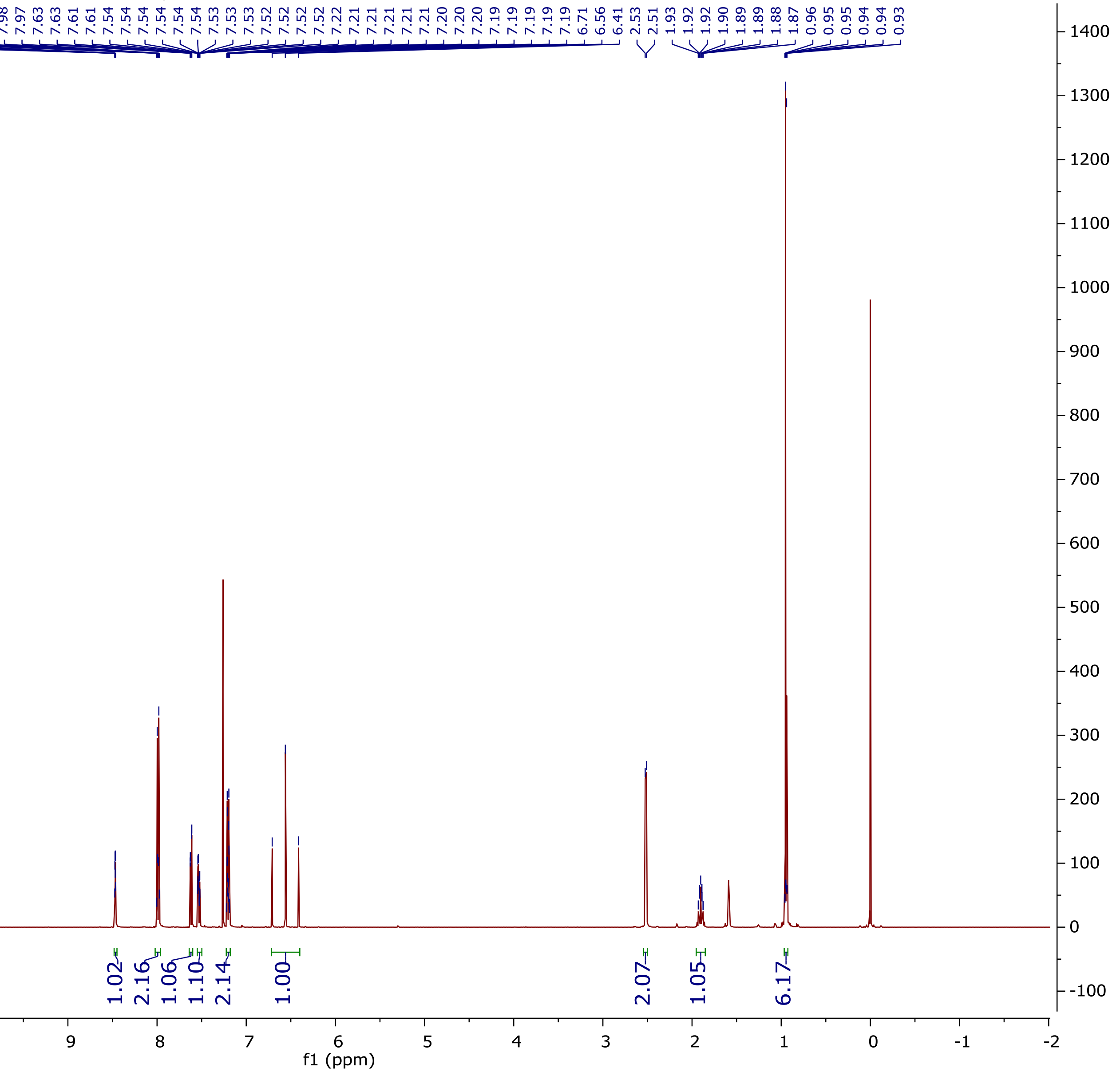

$^{13}$C NMR (125 MHz, $CDCl_3$)

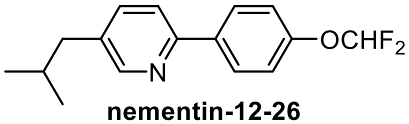


**nementin-12-26**

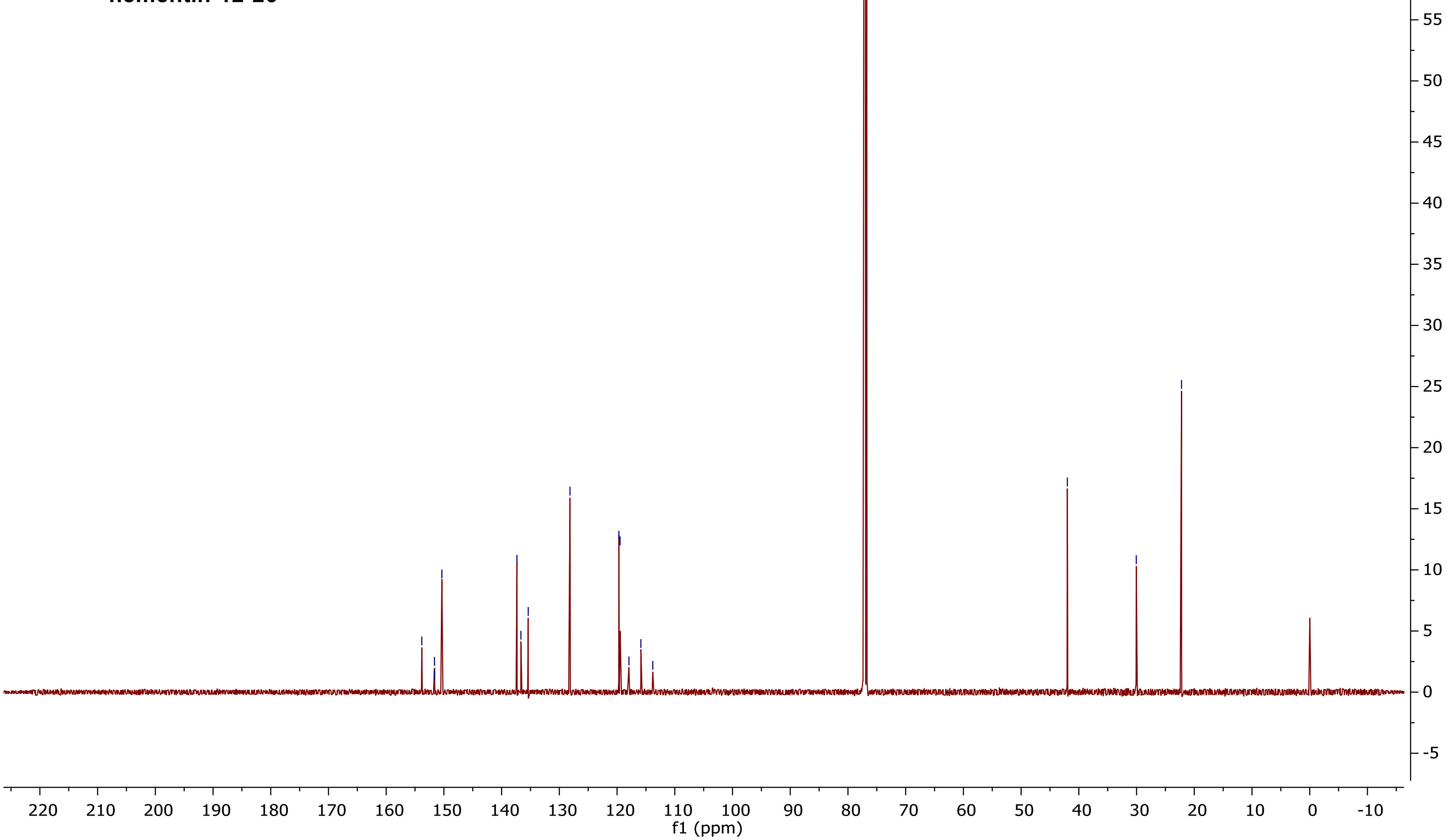

$^{19}F$ NMR (375 MHz, $CDCl_3$)

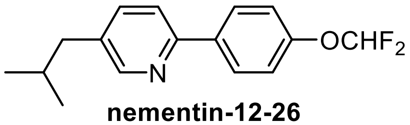


**nementin-12-26**

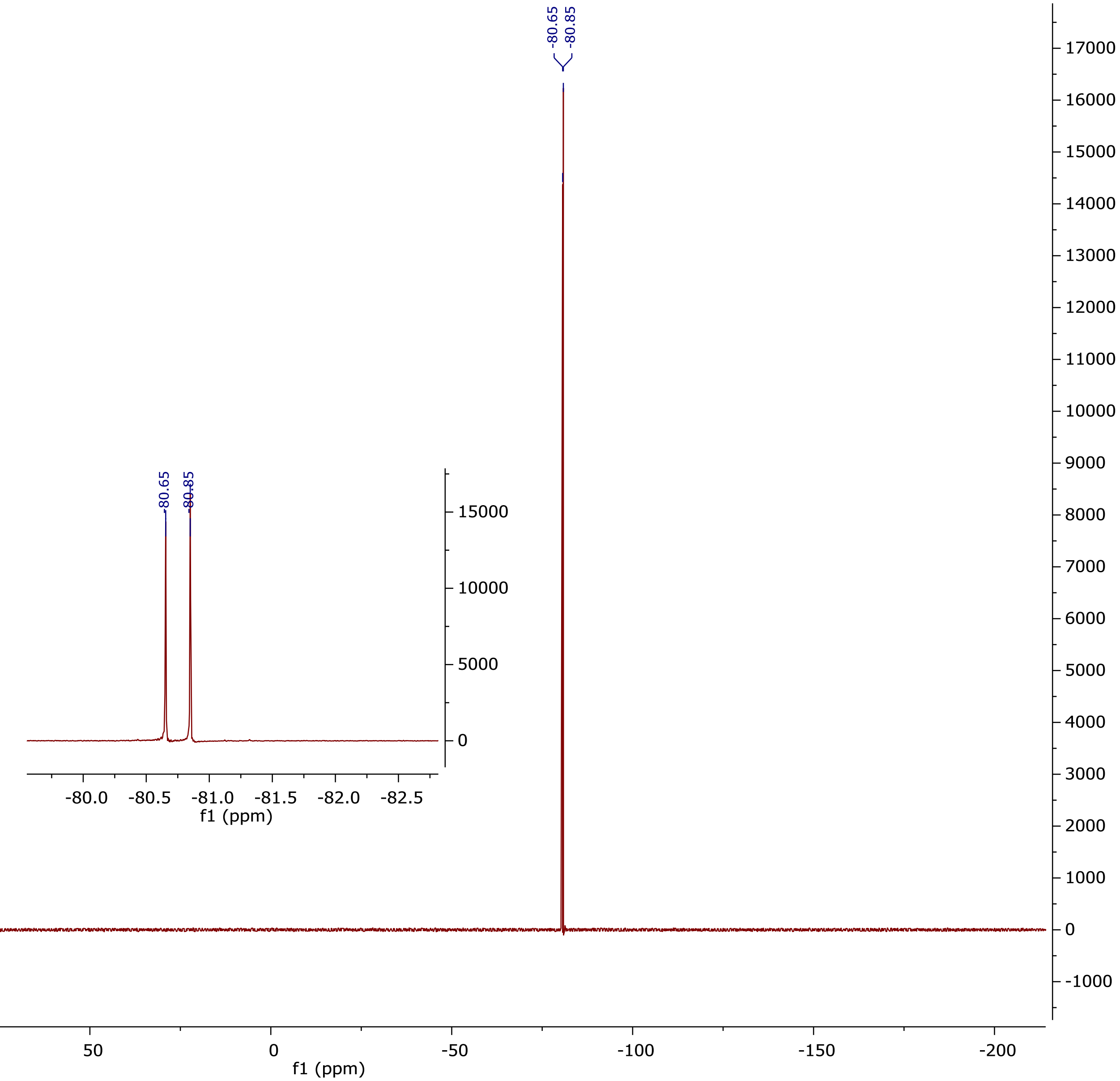

**[1]H NMR (500 MHz, $CDCl_3$) (nementin-12-27)**

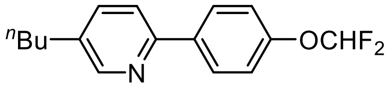


**nementin-12-27**

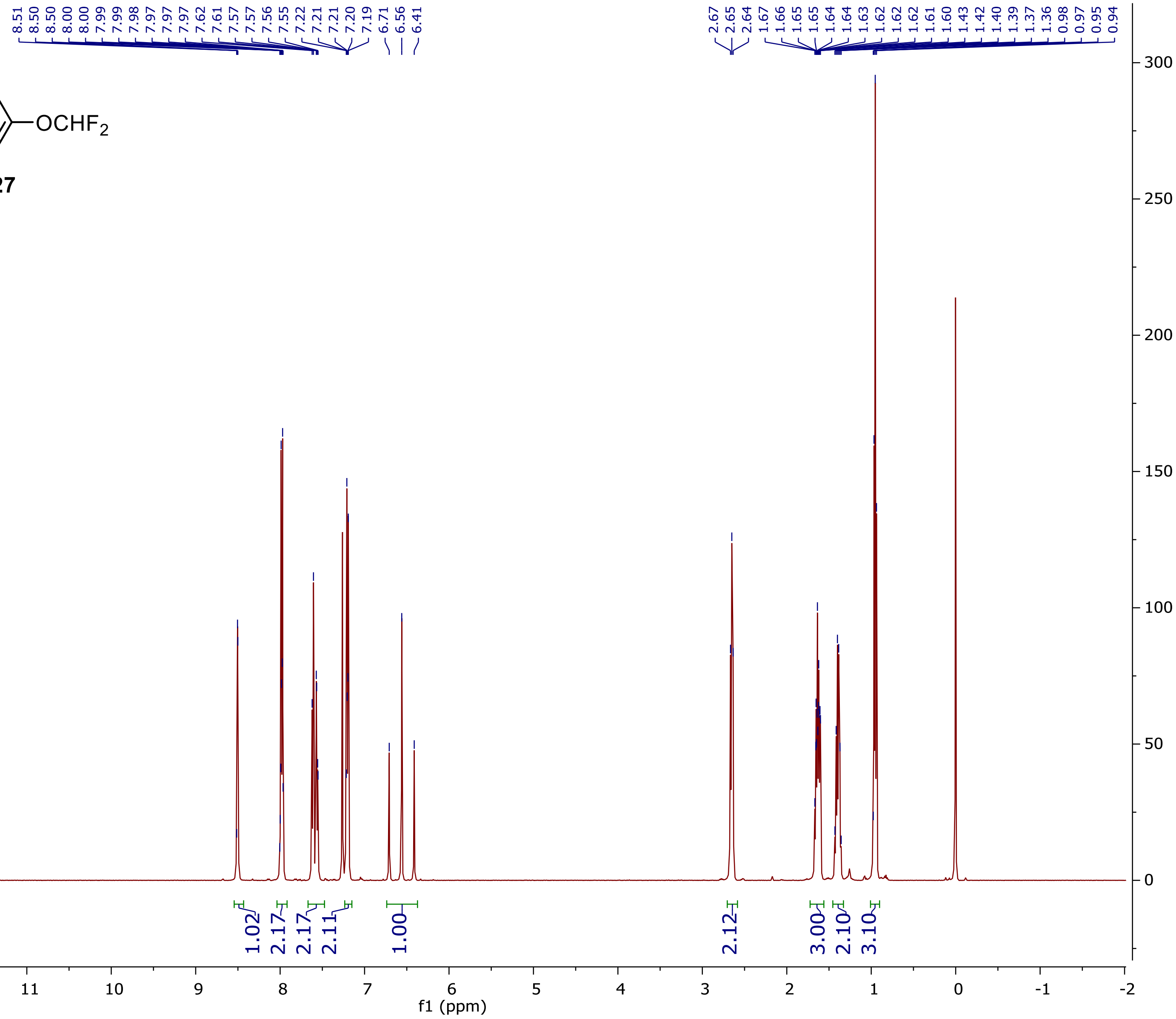

**$^{13}$C NMR (125 MHz, $CDCl_3$)**

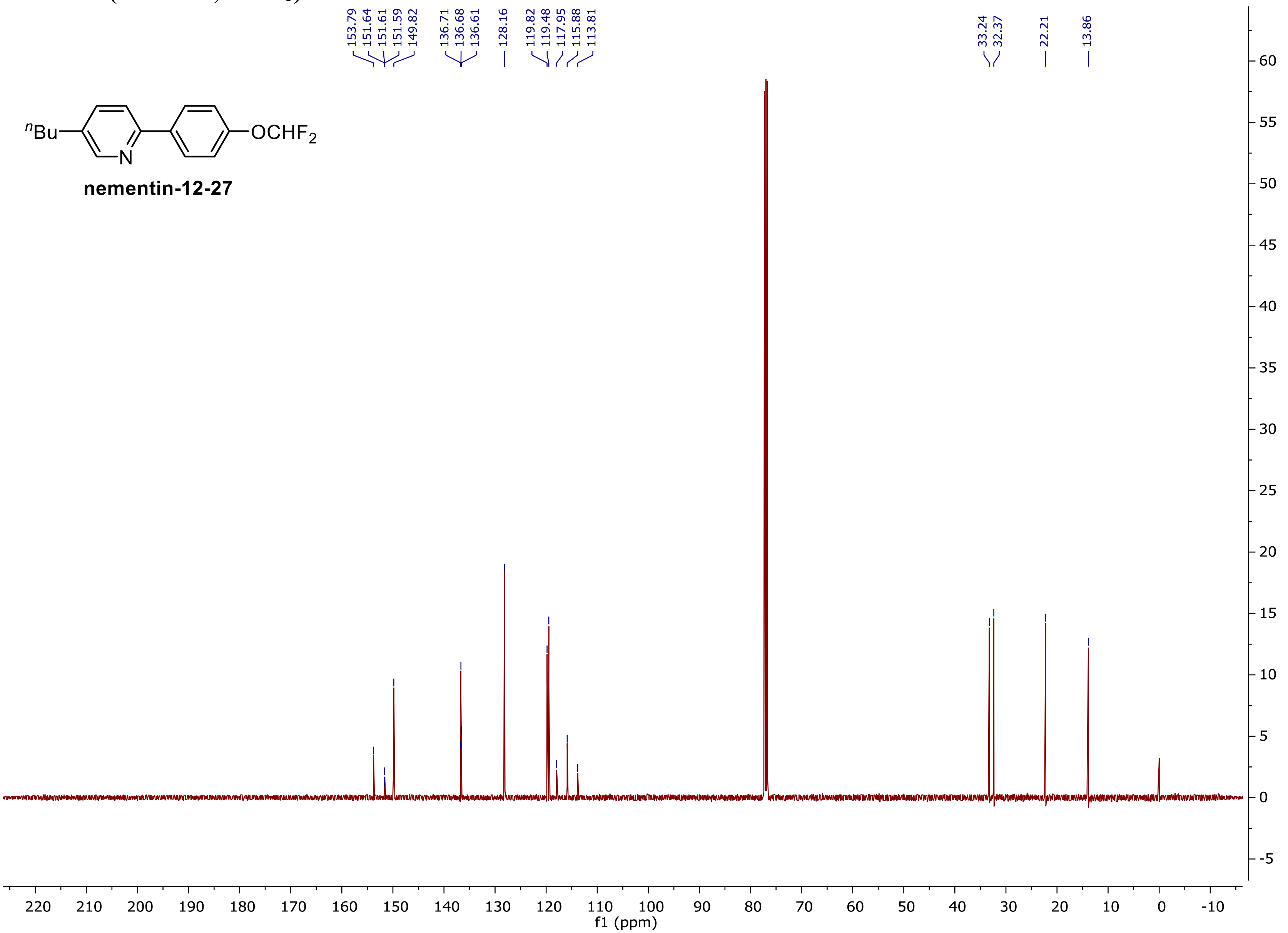

**$^{19}$F NMR (375 MHz, CDCl$_3$)**

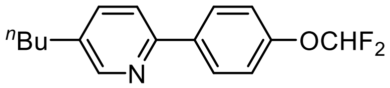


**nementin-12-27**

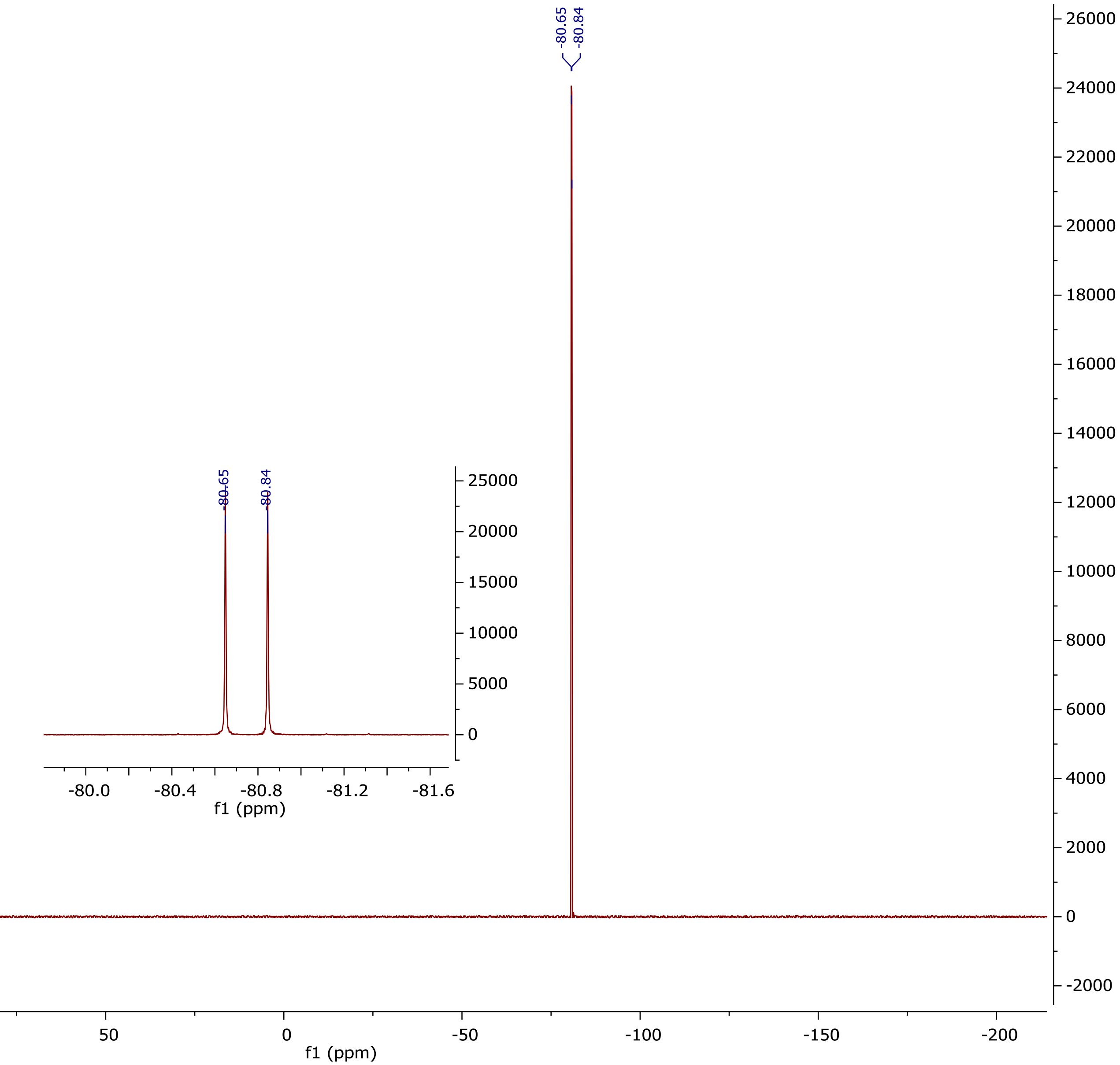

$^1$H NMR (500 MHz, $CDCl_3$) (nementin-12-28)

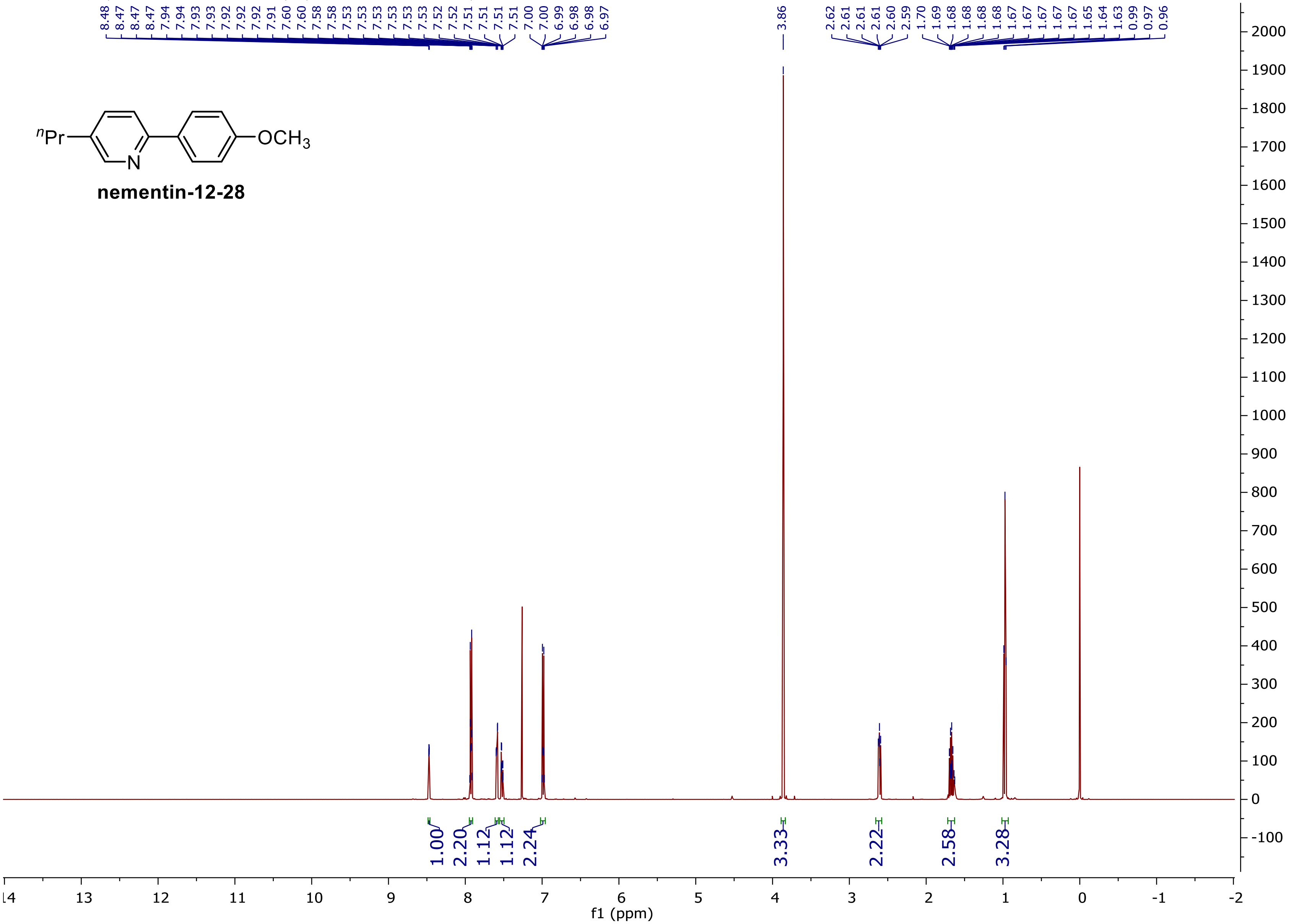


nementin-12-28

### $^{13}C$ NMR (125 MHz, $CDCl_3$)

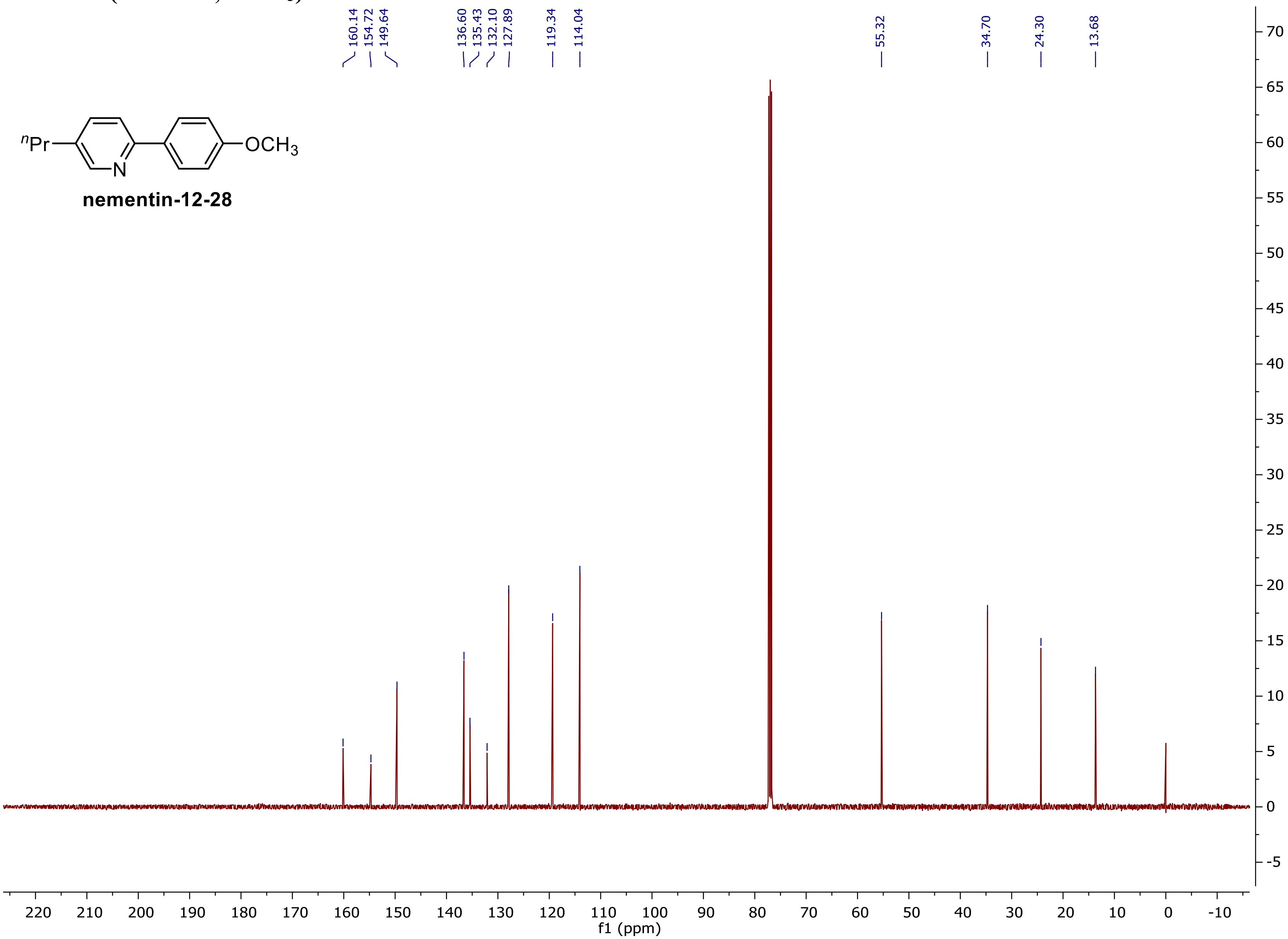

**¹H NMR (500 MHz, $CDCl_3$) (nementin-12-29)**

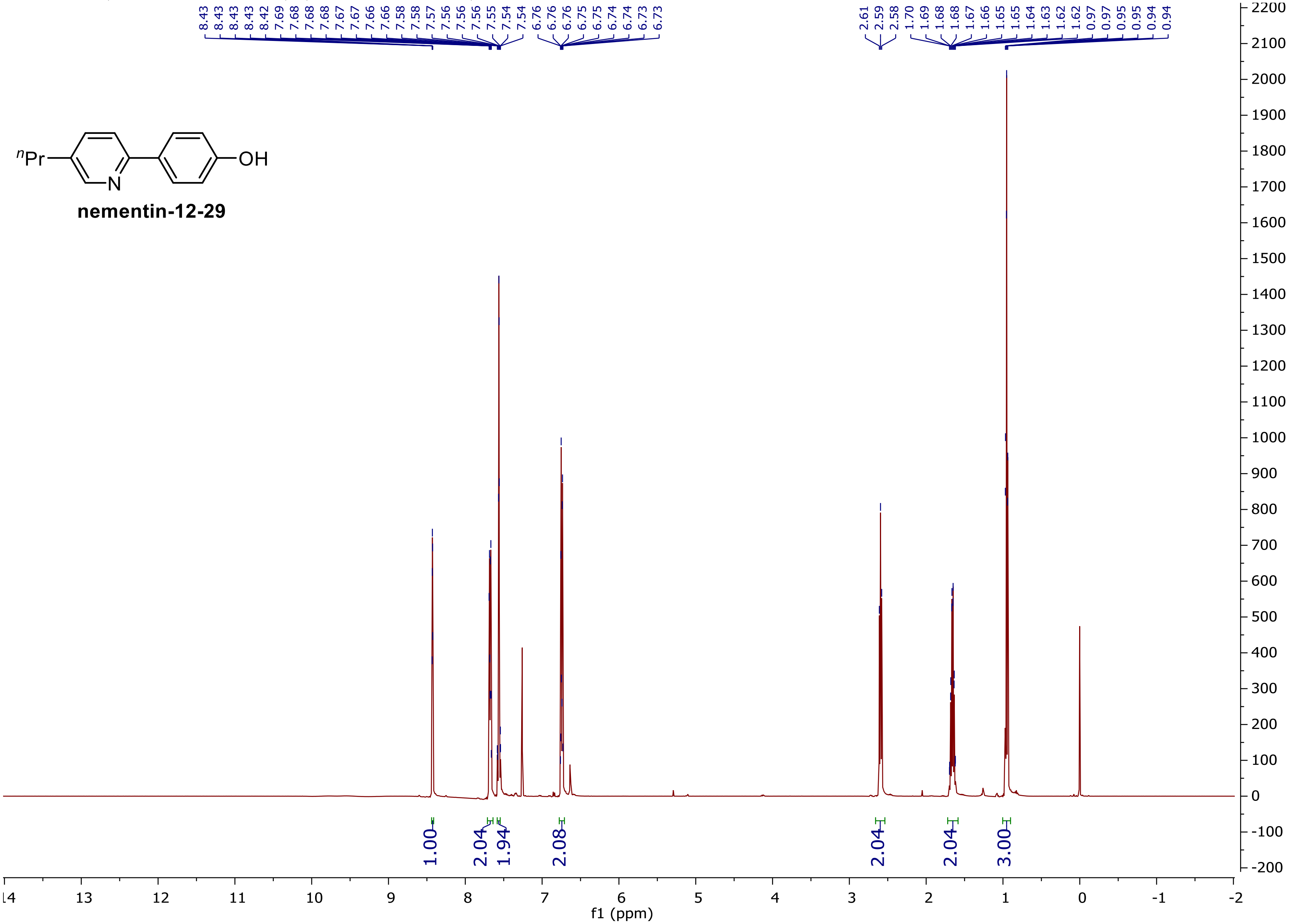

**$^{13}$C NMR (125 MHz, CDCl$_3$)**

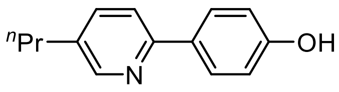


**nementin-12-29**

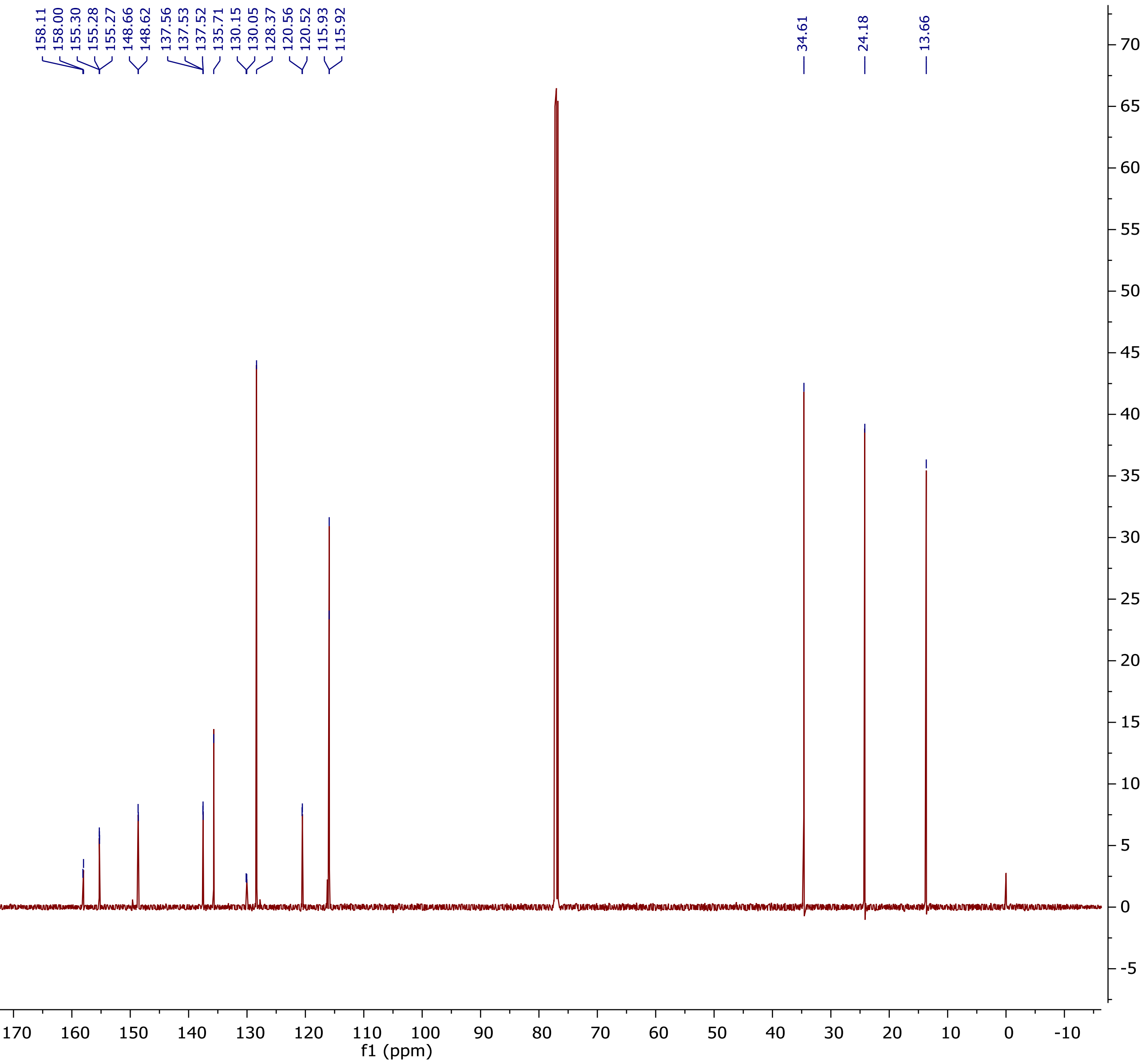

**$^1$H NMR (500 MHz, $CDCl_3$) (nementin-12-30)**

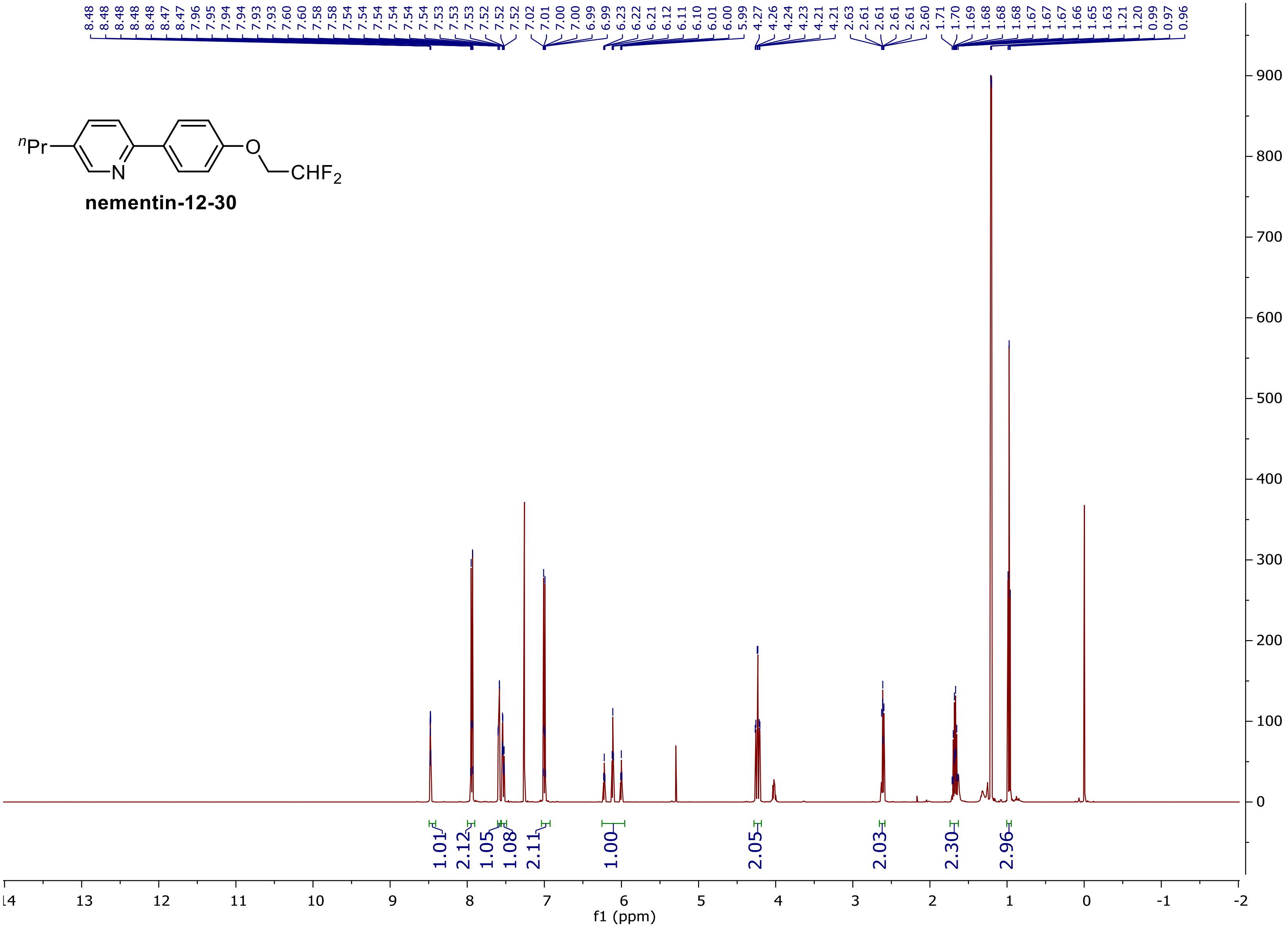


**nementin-12-30**

**$^{13}$C NMR (125 MHz, CDCl$_3$)**

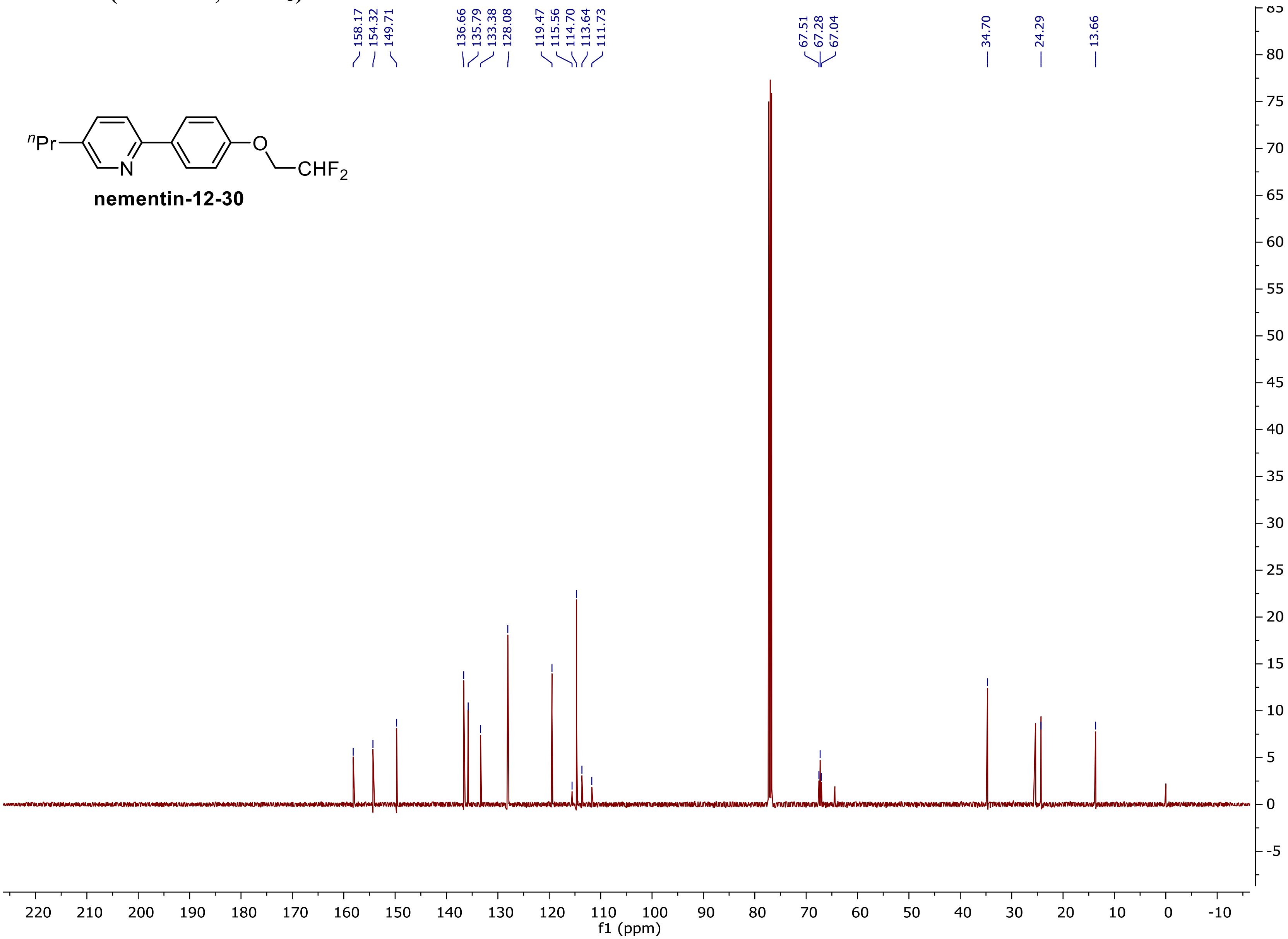

$^{19}$F NMR (375 MHz, $CDCl_3$)

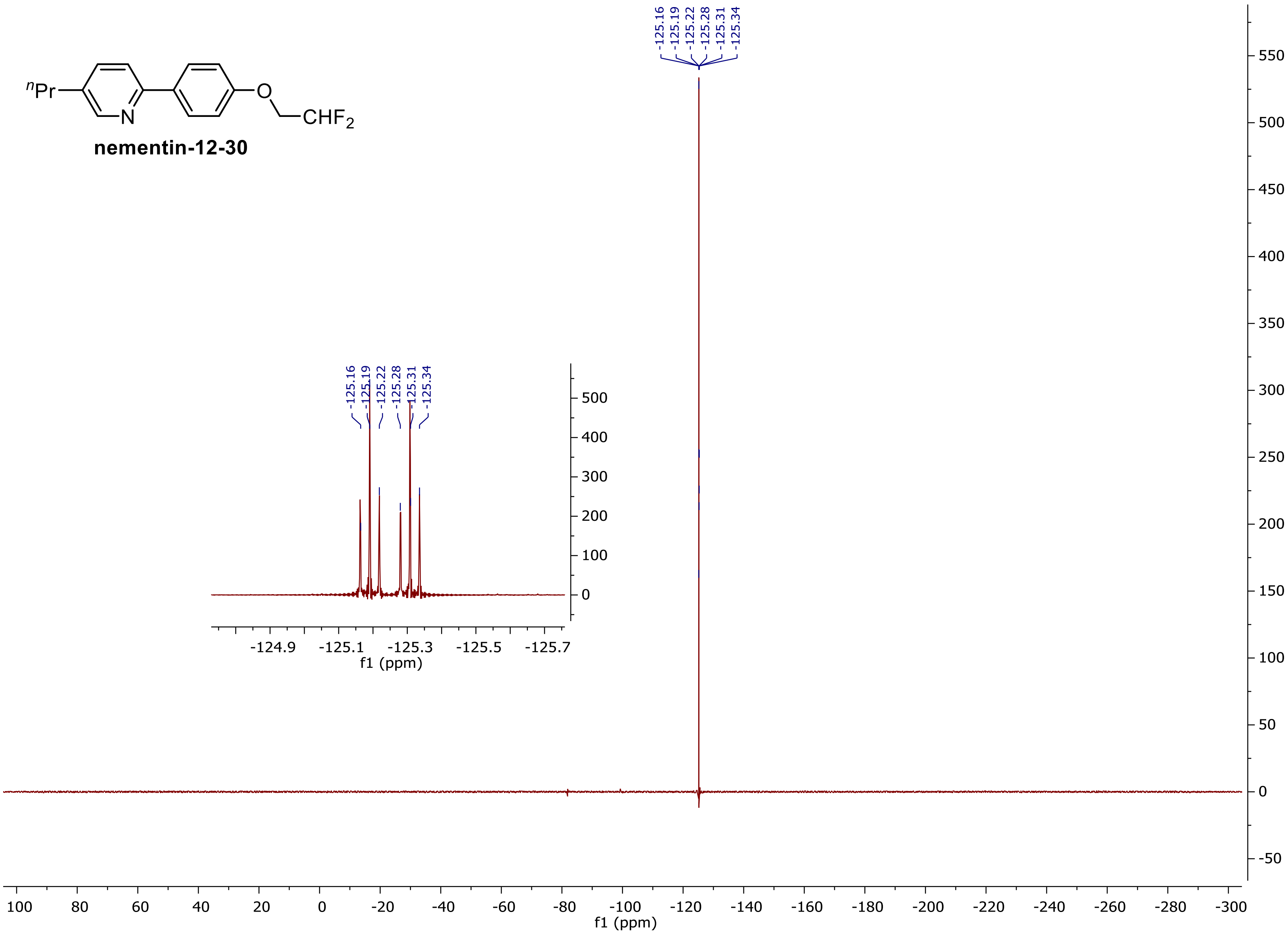


**nementin-12-30**

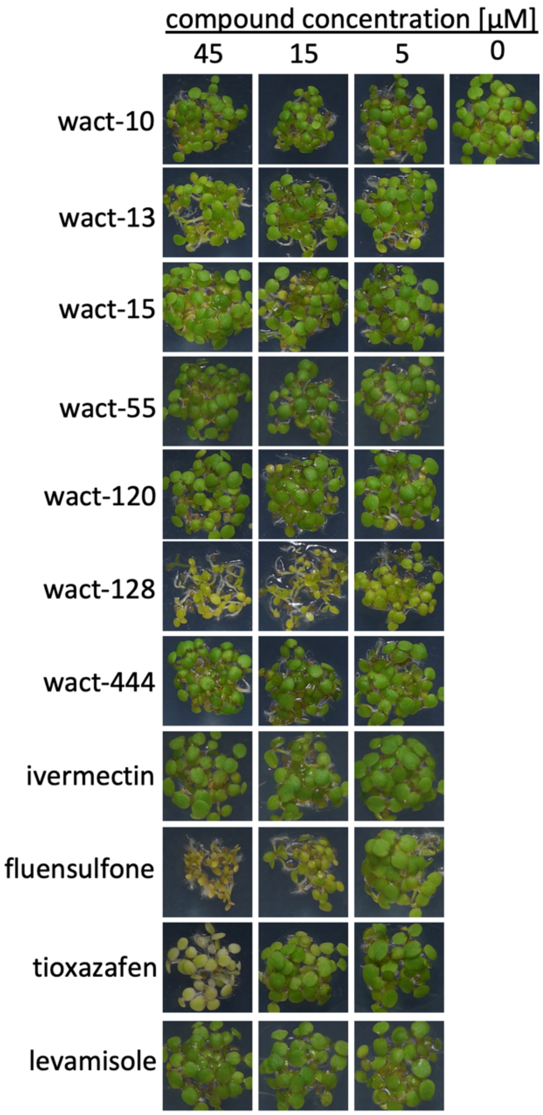


**Supplementary Figure 1.** Effect of prioritized wactives and established anthelmintics on *Arabidopsis* growth and greening. Assays were performed in 12 well polystyrene plates using 20-25 seeds per well (see Methods). Experiments were performed in triplicate. Any yellowing or obvious growth impairment was reported as activity.

| Strain | Resistance | [Nementin-1] µM 0 | 0.94 | 1.88 | 3.75 | 7.5 | 15 | 30 | 60 |
|---|---|---|---|---|---|---|---|---|---|
| N2 | - | 1 | 1 | 1.1 | 1.1 | 1 | 0 | 0 | 0 |
| VC731 *unc-63(ok1075)* | LEV, APAs, THPs | 1 | 1.1 | 1.1 | 1 | 0.8 | 0 | 0 | 0 |
| CB1072 *unc-29(e1072)* | LEV, THPs | 1 | 1 | 1.1 | 1 | 0.8 | 0 | 0 | 0 |
| DA1316 *avr-14;avr-15;glc-1* | MCL | 1 | 1.1 | 1 | 1 | 0.3 | 0 | 0 | 0 |
| CB3474 *ben-1(e1880)* | Benzimidazoles | 1 | 1 | 1.1 | 1.1 | 0.9 | 0 | 0 | 0 |
| RB2119 *acr-23(ok2804)* | AADs | 1 | 0.9 | 1.1 | 1 | 0.7 | 0 | 0 | 0 |
| NM1968 *slo-1(js379)* | Emodepside | 1 | 1.1 | 1.1 | 1 | 0.5 | 0 | 0 | 0 |
| CF1038 *daf-16(mu86)* | Apigenin | 1 | 1 | 1 | 1 | 0.4 | 0 | 0 | 0 |
| RP2674 *mev-1(tr393)* | Fluopyram | 1 | 1 | 1.1 | 0.9 | 0.6 | 0 | 0 | 0 |
| PR1152 *cha-1(p1152)* | AChE inhibitor | 1 | 1 | 1 | 0.9 | 0.2 | 0 | 0 | 0 |

| Strain | Resistance | [levamisole] µM 0 | 9.38 | 18.8 | 37.5 | 75 | 150 | 300 | 600 |
|---|---|---|---|---|---|---|---|---|---|
| N2 | - | 1 | 1 | 1 | 1.1 | 1.1 | 0.5 | 0.2 | 0 |
| VC731 *unc-63(ok1075)* | LEV, APAs, THPs | 1 | 1 | 1 | 1 | 1.1 | 1.1 | 1 | 1 |
| CB1072 *unc-29(e1072)* | LEV, THPs | 1 | 1.1 | 1.1 | 1.1 | 1 | 1.1 | 1 | 1 |
| DA1316 *avr-14;avr-15;glc-1* | MCL | 1 | 0.8 | 1 | 1 | 0.9 | 0.5 | 0.1 | 0 |
| CB3474 *ben-1(e1880)* | Benzimidazoles | 1 | 0.9 | 1 | 1.1 | 0.9 | 0.6 | 0.4 | 0.1 |
| RB2119 *acr-23(ok2804)* | AADs | 1 | 0.9 | 0.8 | 0.6 | 0.5 | 0.1 | 0 | 0 |
| NM1968 *slo-1(js379)* | Emodepside | 1 | 0.9 | 1 | 0.9 | 0.9 | 0.6 | 0.1 | 0 |
| CF1038 *daf-16(mu86)* | Apigenin | 1 | 0.6 | 0.9 | 1 | 0.8 | 0.7 | 0.3 | 0.1 |
| RP2674 *mev-1(tr393)* | Fluopyram | 1 | 0.8 | 0.9 | 1 | 0.9 | 0.4 | 0.1 | 0 |
| PR1152 *cha-1(p1152)* | AChE inhibitor | 1 | 0.3 | 0.2 | 0 | 0 | 0 | 0 | 0 |

| Strain | Resistance | [fluopyram] µM 0 | 0.09 | 0.19 | 0.38 | 0.75 | 1.5 | 3 | 6 |
|---|---|---|---|---|---|---|---|---|---|
| N2 | - | 1 | 1.1 | 1 | 0.7 | 0 | 0 | 0 | 0 |
| VC731 *unc-63(ok1075)* | LEV, APAs, THPs | 1 | 1 | 0.7 | 0.3 | 0 | 0 | 0 | 0 |
| CB1072 *unc-29(e1072)* | LEV, THPs | 1 | 0.9 | 0.7 | 0.7 | 0 | 0 | 0 | 0 |
| DA1316 *avr-14;avr-15;glc-1* | MCL | 1 | 1 | 0.9 | 0.2 | 0 | 0 | 0 | 0 |
| CB3474 *ben-1(e1880)* | Benzimidazoles | 1 | 1 | 0.8 | 0.7 | 0 | 0 | 0 | 0 |
| RB2119 *acr-23(ok2804)* | AADs | 1 | 1 | 1 | 0.4 | 0 | 0 | 0 | 0 |
| NM1968 *slo-1(js379)* | Emodepside | 1 | 1 | 0.6 | 0 | 0 | 0 | 0 | 0 |
| CF1038 *daf-16(mu86)* | Apigenin | 1 | 1 | 1 | 0.5 | 0 | 0 | 0 | 0 |
| RP2674 *mev-1(tr393)* | Fluopyram | 1 | 1.1 | 1.1 | 1 | 1 | 1.1 | 1 | 0.9 |
| PR1152 *cha-1(p1152)* | AChE inhibitor | 1 | 1 | 0.3 | 0 | 0 | 0 | 0 | 0 |

| Strain | Resistance | [ivermectin] nM 0 | 1.88 | 3.75 | 7.5 | 15 | 30 | 60 | 120 |
|---|---|---|---|---|---|---|---|---|---|
| N2 | - | 1 | 1 | 1.1 | 0.8 | 0.2 | 0 | 0 | 0 |
| VC731 *unc-63(ok1075)* | LEV, APAs, THPs | 1 | 1 | 1.1 | 1 | 0 | 0 | 0 | 0 |
| CB1072 *unc-29(e1072)* | LEV, THPs | 1 | 0.9 | 1.1 | 1 | 0 | 0 | 0 | 0 |
| DA1316 *avr-14;avr-15;glc-1* | MCL | 1 | 1 | 1.1 | 1 | 1 | 1 | 0.8 | 0.4 |
| CB3474 *ben-1(e1880)* | Benzimidazoles | 1 | 1 | 1 | 0.7 | 0.2 | 0 | 0 | 0 |
| RB2119 *acr-23(ok2804)* | AADs | 1 | 1 | 0.9 | 1 | 0.9 | 0.2 | 0 | 0 |
| NM1968 *slo-1(js379)* | Emodepside | 1 | 1 | 0.8 | 0.1 | 0 | 0 | 0 | 0 |
| CF1038 *daf-16(mu86)* | Apigenin | 1 | 1 | 1 | 0.9 | 0 | 0 | 0 | 0 |
| RP2674 *mev-1(tr393)* | Fluopyram | 1 | 0.9 | 1 | 1 | 1 | 0.2 | 0 | 0 |
| PR1152 *cha-1(p1152)* | AChE inhibitor | 1 | 0.9 | 0.2 | 0 | 0 | 0 | 0 | 0 |

≥ 1 — 0
number of living worms relative to DMSO control

**Supplementary Figure 2.** *C. elegans* strains with mutant alleles conferring anthelmintic resistance are sensitive to the effects of Nementin-1. 3-day larval development assay monitoring the development of larval stage 1 (L1) to L4/adulthood. Reporting the fraction of animals that develop to L4/adulthood relative to the solvent-only controls. Data are the mean of 3 biological replicates measured in technical triplicate normalized to parallel N2 controls. Showing the strain name and genetic background of parasiticide resistant mutants. Resistance to marketed nematode parasiticides is demonstrated for levamisole (strain VC731 & CB1072), fluopyram (strain RB2674), and ivermectin (DA1316).

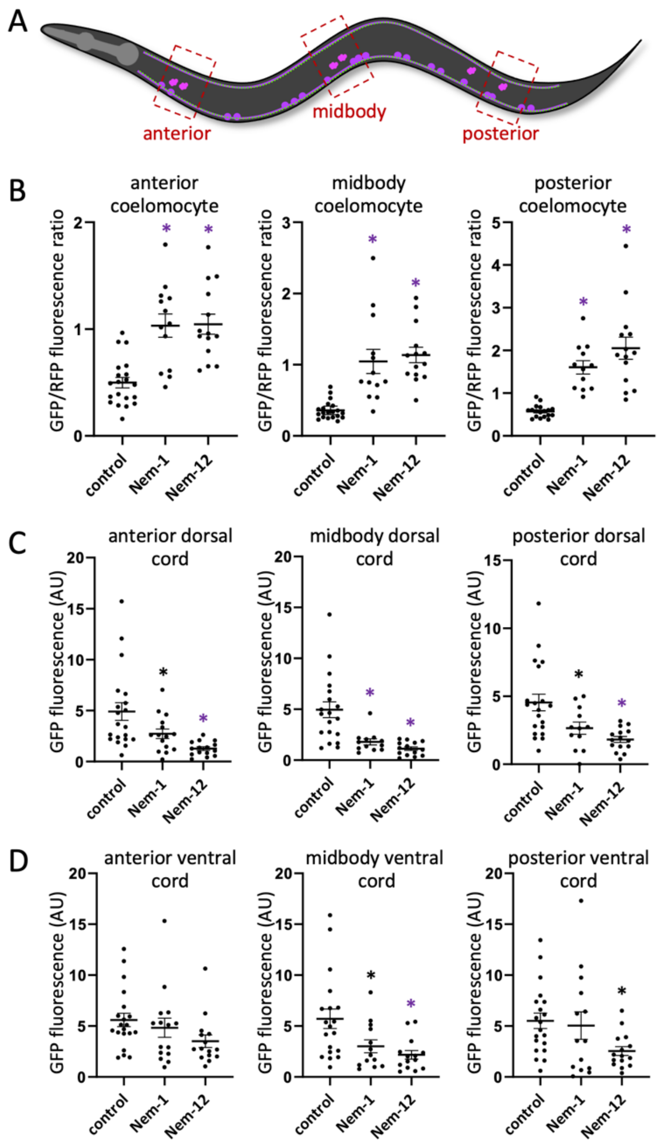


**Supplementary Figure 3.** Nementin treatment depletes *C. elegans* cholinergic motor neurons of neuropeptides that are released in DCVs. (**a**) Schematic of KG4247 *C. elegans* witih the *ceIs201* integrated transgene expressing INS-22::GFP in cholinergic motor neurons. INS-22::GFP gets packaged into DCVs serving as a marker for DCVs. The cholinergic motor neurons of KG4247 animals also constitutively secrete mCherry, which is taken up by the coelomocytes, and serves as a coelomocyte marker (10).

Imaged regions are indicated with dashed red boxes. (**b-d**) Quantification of signal from the indicated cell or tissue in the indicated area of the animal. Signal is measured from coelomocytes with a ratio of GFP to RFP fluorescence (because measuring GFP signal alone is too variable because of the variable size of coelomocyte endosomes). Signal is reported from the dorsal and ventral cords as the mean GFP signal with the mean surrounding background signal subtracted. Animals are treated either with DMSO-only (control) or 60 $\mu$M of Nementin for 4 hours. For all images, * $p<0.05$;** $p<0.01$; *** $p<0.001$, 1-way ANOVA with Dunnett correction for multiple comparisons; means with SEM are shown.

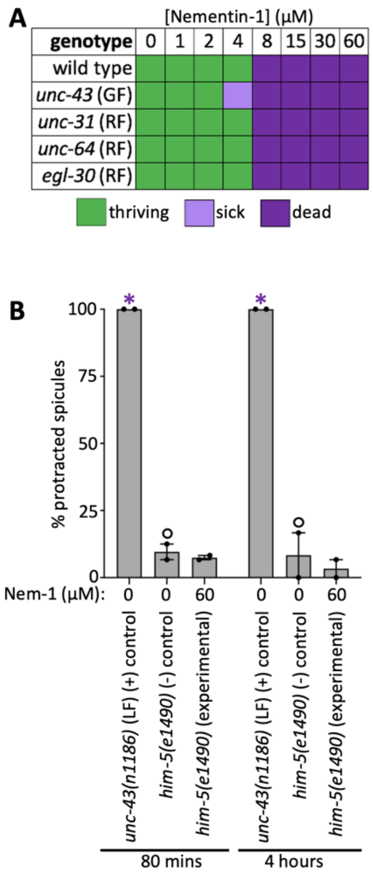


**Supplementary Figure 4.** UNC-43/CaMKII is unlikely to be the target of Nementin. (**a**) Six-day larval reproduction assay measuring the effects of Nementin-1 on generational development. ~20 L1s were plated per well containing 60 *μ*M Nementin-1 in NGM media supplemented with HB101 *E. coli* as a food source with 0.6% DMSO. Thriving indicates wells are overgrown with progeny after 6-days. Sick indicates <50 L1 larvae are present per well. Dead indicates <10 live worms are present in test wells. Data are the median response of 3 biological replicates. (**b**) A test to determine whether Nementin-1 inhibits UNC-43 in its role in *C. elegans* male tail spicule protraction. The *unc-43* null mutant *(n1186)* is the positive control. The asterisk signifies Chi-Square *p*<0.001 comparing the experimental and positive control to untreated *him-5(e1490)*.

| | | | | locomotor phenotype[c] | | | | | | | larval lethal assay[d] | | | | | | |
|---|---|---|---|---|---|---|---|---|---|---|---|---|---|---|---|---|---|
| | wactive | Chembridge ID[a] | egg-laying phenotype[b] | shrinker | shaker | coiler | jerky-Unc | reversal-defective | paralyzed | slow | *C. elegans* | *C. briggsae* | *P. pacificus* | *C. oncophora* | *H. contortus* | HEK293T[e] | *Danio rerio*[f] |
| **Convulsive (primarily)** | | | | | | | | | | | | | | | | | |
| 1 | wact-10 | 5905384 | Egl-S | 30 | 60 | n | n | n | n | n | 8 | 30 | 30 | 4 | 60 | n | n |
| 2 | wact-13 | 7632721 | Egl-S | 30 | n | n | 30 | 60 | n | n | 8 | 30 | 30 | 8 | 60 | n | n |
| 3 | wact-15 | 7687977 | Egl-I | 30 | n | n | 30 | 30 | n | n | 8 | 60 | 60 | 8 | 60 | n | n |
| 4 | wact-55 | 5688274 | Egl-S | 60 | 60 | n | n | n | 30 | n | 8 | 30 | 30 | 8 | 8 | n | n |
| 5 | wact-128 | 7971816 | Egl-S | 30 | n | n | n | n | n | n | 30 | 30 | 30 | 4 | 60 | n | n |
| 6 | wact-181 | 9032631 | Egl-S | 30 | >60 | n | n | n | n | n | 30 | 30 | 8 | 15 | 60 | 30 | dead |
| 7 | wact-209 | 7153658 | Egl-S | 30 | 60 | n | n | n | n | n | 30 | 30 | 30 | 15 | 60 | 60 | n |
| 8 | wact-223 | 7638283 | Egl-I | 60 | n | n | n | n | n | n | n | n | n | 60 | 60 | 60 | n |
| 9 | wact-542 | 7781713 | Egl-I | 30 | n | n | n | n | n | n | n | n | 60 | 15 | 60 | n | dev |
| **Shaker (primarily)** | | | | | | | | | | | | | | | | | |
| 10 | wact-203 | 9062286 | Egl-S | >60 | 60 | n | 60 | n | n | n | 30 | 30 | 30 | 15 | 60 | n | dev |
| 11 | wact-444 | 5910902 | Egl-S | n | 30 | n | n | n | n | n | 30 | 30 | 30 | 15 | 60 | n | n |
| 12 | wact-577 | 9039023 | Egl-S | n | 60 | n | n | n | 60 | n | 60 | 60 | 60 | 60 | 60 | n | n |
| **Coiler (primarily)** | | | | | | | | | | | | | | | | | |
| 13 | wact-6 | 5419367 | Egl-S | n | n | 30 | n | n | 30 | n | 30 | 30 | 30 | >60 | >60 | 30 | card |
| 14 | wact-45 | 5426270 | Egl-S | n | n | 30 | 60 | n | 60 | n | n | n | n | 60 | >60 | n | n |
| 15 | wact-47 | 5427063 | Egl-S | n | n | 60 | >60 | n | >60 | n | 30 | n | n | 60 | >60 | 30 | card |
| **Reversal Defective (primarily)** | | | | | | | | | | | | | | | | | |
| 16 | wact-38 | 5356411 | Egl-S | n | n | n | n | 30 | n | n | 30 | 30 | 30 | 8 | 60 | n | n |
| 17 | wact-558 | 9008591 | Egl-S | n | n | n | n | 60 | n | 30 | 8 | 30 | 30 | 60 | 60 | n | n |
| **Jerky-Unc (primarily)** | | | | | | | | | | | | | | | | | |
| 18 | wact-46 | 5426998 | Egl-S | n | n | n | 30 | n | >60 | n | n | 60 | n | 8 | >60 | n | card |
| 19 | wact-120 | 7889289 | Egl-I | n | n | n | 60 | n | n | n | 30 | 30 | 30 | 4 | 60 | n | n |
| **Paralysis** | | | | | | | | | | | | | | | | | |
| 20 | wact-11 | 6222549 | Egl-I | n | n | n | n | n | 30 | n | 8 | 8 | 8 | 8 | 60 | n | n |
| 21 | wact-12 | 7003409 | Egl-I | n | n | n | n | n | n | n | 8 | 30 | 8 | 8 | 60 | n | n |
| 22 | wact-220 | 5784085 | Egl-I | n | n | n | n | n | 30 | n | 8 | n | 30 | >60 | 60 | n | dev |
| **Slow** | | | | | | | | | | | | | | | | | |
| 23 | wact-423 | 5547023 | Egl-I | n | n | n | n | n | n | 30 | 60 | n | 60 | 8 | 60 | 30 | n |
| 24 | wact-503 | 7568929 | Egl-S | n | n | n | n | n | n | 30 | 60 | 60 | 60 | 60 | 60 | 30 | n |
| 25 | wact-614 | 9039813 | Egl-I | n | n | n | n | n | n | 30 | 8 | 30 | 60 | >60 | >60 | 30 | dead |
| 26 | wact-622 | 5373894 | Egl-I | n | n | n | n | n | n | 30 | n | n | n | 60 | >60 | 30 | n |
| **Weak or No Locomotory Phenotype** | | | | | | | | | | | | | | | | | |
| 27 | wact-2 | 5185411 | Egl-S | n | n | n | n | n | n | 60 | 8 | 8 | 8 | 8 | >60 | 60 | dev |
| 28 | wact-4 | 5352487 | Egl-S | >60 | n | n | n | n | n | n | 30 | 30 | 30 | 30 | 60 | n | n |
| 29 | wact-8 | 5652977 | Egl-S | n | n | n | n | >60 | n | n | 30 | 8 | 8 | 60 | >60 | 30 | dead |
| 30 | wact-21 | 5129511 | Egl-S | n | n | n | >60 | n | n | n | 30 | n | n | 8 | 60 | n | n |
| 31 | wact-23 | 5156707 | Egl-S | n | n | n | n | >60 | n | n | 30 | 30 | 8 | 8 | 60 | 30 | dead |
| 32 | wact-33 | 5308651 | Egl-I | n | n | n | n | n | n | n | 30 | 30 | 30 | 8 | >60 | 30 | n |
| 33 | wact-35 | 5344384 | Egl-I | n | n | n | n | 60 | n | n | 30 | 60 | 60 | 8 | 60 | 30 | n |
| 34 | wact-36 | 5347942 | Egl-I | n | n | n | n | n | n | n | n | n | n | 8 | 60 | n | n |
| 35 | wact-39 | 5357418 | Egl-S | n | n | n | n | n | n | n | 30 | n | 30 | 8 | 60 | n | n |
| 36 | wact-62 | 6269333 | Egl-I | depleted | | | | | | | 30 | 30 | 60 | 8 | 60 | n | n |
| 37 | wact-124 | 7943845 | Egl-I | n | n | n | n | n | n | >60 | 30 | 30 | 30 | 8 | 60 | 60 | card |
| 38 | wact-125 | 7949759 | Egl-I | n | n | n | n | n | n | n | 30 | 60 | 8 | 60 | >60 | n | n |
| 39 | wact-169 | 9024879 | Egl-S | n | n | n | n | n | n | n | 8 | 30 | 8 | >60 | >60 | n | n |
| 40 | wact-182 | 9033974 | Egl-I | n | n | n | n | 60 | n | n | 30 | 60 | n | n | n | 60 | n |
| 41 | wact-187 | 9035461 | Egl-I | n | n | n | n | n | n | n | <<30 | 60 | n | n | n | 60 | n |
| 42 | wact-204 | 9063096 | Egl-I | n | n | n | n | >60 | n | n | 30 | 60 | 30 | >>60 | >60 | 30 | dead |
| 43 | wact-213 | 5107544 | Egl-I | n | n | n | n | n | n | n | 60 | n | n | >60 | >60 | n | n |
| 44 | wact-218 | 5379978 | Egl-S | n | n | n | >60 | n | n | n | 8 | 30 | 30 | 15 | 60 | 30 | n |
| 45 | wact-371 | 5100784 | Egl-S | n | n | n | n | n | n | n | 8 | 60 | 60 | 8 | 60 | 30 | n |
| 46 | wact-376 | 5192203 | Egl-I | depleted | | | | | | | 60 | 30 | 30 | 8 | 60 | n | dead |
| 47 | wact-378 | 5192696 | Egl-S | n | n | n | n | n | n | 60 | 30 | 60 | 60 | 8 | 60 | n | n |
| 48 | wact-380 | 5218876 | Egl-I | n | n | n | n | n | n | n | n | n | 30 | >60 | n | n | card |
| 49 | wact-385 | 5238652 | Egl-I | n | n | n | n | >60 | n | n | 30 | 60 | 60 | 8 | 60 | 30 | dead |
| 50 | wact-396 | 5322542 | Egl-S | n | n | n | n | n | n | n | 60 | 60 | 60 | 8 | 60 | n | n |
| 51 | wact-419 | 5469460 | Egl-S | >60 | >60 | n | n | n | n | n | 60 | 60 | 60 | 60 | 60 | n | n |
| 52 | wact-432 | 5689365 | Egl-S | n | n | n | n | n | n | n | 60 | 60 | 60 | 8 | 60 | n | n |
| 53 | wact-515 | 7653692 | Egl-I | n | n | n | n | >60 | n | n | n | n | 60 | 60 | 60 | 30 | n |
| 54 | wact-516 | 7664300 | Egl-I | >60 | n | n | n | n | n | n | >60 | 60 | 30 | 4 | 60 | n | card |
| 55 | wact-525 | 7705966 | Egl-I | n | n | n | n | n | n | n | n | n | n | 60 | 60 | 60 | n |
| 56 | wact-547 | 7877602 | Egl-I | n | n | n | n | n | n | n | n | n | 60 | >60 | >60 | 30 | dead |
| 57 | wact-578 | 9040453 | Egl-I | n | n | n | n | n | n | n | 30 | n | n | 8 | 8 | n | dead |
| 58 | wact-596 | 5649594 | Egl-I | n | n | n | n | n | n | n | 30 | 30 | 30 | >60 | 60 | n | dead |

**Supplementary Table 1.** *C. elegans* locomotor phenotype and phylogenetic activity profile of identified egg-laying modulators.

[a]The Chembridge Inc. identification (ID) numbers are indicated.

[b]Egg-laying phenotype; Egl-S = egg-laying stimulators; Egl-I = egg-laying inhibitors.

[c]The observed acute motor phenotypes are indicated. The concentration at which strong and moderate phenotypes appear are indicated in bright green. The concentration at which weaker phenotypes appear are indicated in light gray. 'n' indicates no phenotype was observed.

[d]Larval lethality phenotypes; the previously reported results (11) of liquid-based larval lethal assays (for *C. elegans*, *C. briggsae*, and *P. pacificus*) are shown. The lowest concentration at which 100% of the larvae die/arrest is shown. Colours highlight relatively potent activity.

[e]HEK293T cell proliferation summary; compounds that reduce HEK293T proliferation below one standard of deviation from the mean at 30 or 60 $\mu$M as previously reported are indicated as purple '30' or '60', respectively (see (11) for details).

[f]*Danio rerio* (zebrafish) developmental defect summary; molecules that induce cardiac defects (card), developmental defects (dev) or death (dead) at a concentration of 10 $\mu$M as previously reported(11) are summarized here. Throughout the table, 'n' reports no lethality or phenotype observed.

| gene | ortholog | exemplar phenocopying mutation and phenotypes[a] | potential suppressing mutation and phenotypes[b] |
|---|---|---|---|
| *acr-2* | CHRNA1 (cholinergic receptor nicotinic subunit) (12) | *n2420* (GF) convulsions, Egl-c (13, 14) | *acr-2(ok1887)* (LF) non-Unc (14) |
| *unc-2* | CaV2α (voltage gated calcium channel)(15) | *zf35* (GF) hyperactive, convulsions (16) | *unc-2(e55)* (LF) sluggish (17) |
| *unc-43* | CaMKII(18) | *e408* (RF) convulsions, Egl-c (18, 19) | *unc-43(n498)* (GF) paralysis (20) |
| *unc-58* | KCNK3, KCNK9 and KCNK18 (two-pore $K^+$ channel)(21) | *e665* (GF) convulsive, Egl-c (21) | *unc-58(e665n273)* (LF) weak Unc(21) |
| *unc-93* | UNC93A (SUP-9 $K^+$ channel complex member)(22) | *e1500* (GF) rubberband (23) | *sup-9(n1012)* (LF) non-Unc (22) |
| *sup-9* | two-pore $K^+$ channel(22) | *n200* (GF) convulsive (23, 24) | *sup-9(n1012)* (LF) non-Unc (22) |
| *sup-10* | novel (SUP-9 $K^+$ channel complex member)(22) | *n983* (GF) rubber band (23) | *sup-9(n1012)* (LF) non-Unc (22) |
| *twk-18* | TWiK $K^+$ channel(25) | *e1913* (GF) lethal, rubberband, Egl-d (GF) (26) | *twk-18(gk5009)* (LF) non-Unc (wormbase version WS280) |

**Supplementary Table 2.** Genes that can be mutated to induce convulsions or 'rubber-band' phenotypes. [a]**GF, gain-of-function; RF, reduction-of-function; Egl-c, constitutive egg-laying**

[b]LF, loss-of-function; Unc, uncoordination.

A

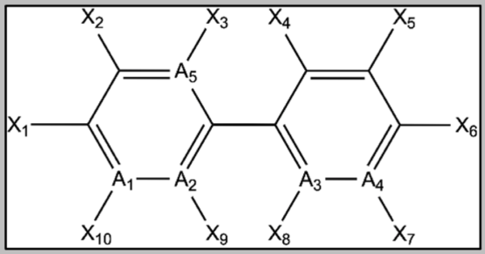


| A | | | | | | | | | | B free living nematodes | | | | | | C nematode parasites of animals | | | | | | | D plant parasites | | | | E non target models | | | |
|---|---|---|---|---|---|---|---|---|---|---|---|---|---|---|---|---|---|---|---|---|---|---|---|---|---|---|---|---|---|---|
| nementin analog | $A_1$ | $A_2$ | $A_3$ | $A_4$ | $A_5$ | $X_1$ | $X_6$ | R | R Position | C. elegans (motor phenotype) EC50 | P. pacificus (motor phenotype) EC50 | Rhabditophanes sp. (motor phenotype) EC50 | C. elegans (3-day larval development) EC50 | P. pacificus (3-day larval development) EC50 | Rhabditophanes sp. (3-day larval development) EC50 | Necator americanis (L3 viability) | Trichuris muris (L1 viability) | Trichuris muris (adult viability) | Strongyloides ratti (L3 viability) | Strongyloides ratti (adult viability) | Heligmosomoides polygyrus (L3 lethality) | Heligmosomoides polygyrus (adult lethality) | M. incognita (J2 viability in vitro) | M. incognita (egg hatch rate) | M. incognita (hatchling mobility) | M. incognita (50-day soil test ) | Drosophila melanogaster (45 µM highest tested) | Danio rerio (45 µM highest tested) | Arabidopsis thaliana greening (45 µM highest tested) | HEK293 EC50 (µM) |
| 1 | C | N | C | C | C | $(CH_2)_4CH_3$ | $OCH_3$ | - | - | 31 | 25 | 5 | 9 | 10 | 2 | 25 | nt | 25 | np | 25 | np | 25 | 45 | 45 | 45 | np | np | np | np | np |
| 2 | C | N | C | C | C | $(CH_2)_4CH_3$ | $OCHF_2$ | - | - | 19 | 15 | nt | 15 | nt | nt | 10 | nt | 25 | np | 25 | np | 25 | np | 45 | np | nt | nt | 45 | nt | nt |
| 3 | C | N | C | C | C | $(CH_2)_6CH_3$ | OH | - | - | np | 36 | 22 | 15 | 17 | 2 | 10 | nt | 25 | np | 25 | 25 | 25 | 45 | 45 | 45 | 45 | np | np | np | 43 |
| 4 | C | N | C | C | C | $(CH_2)_7CH_3$ | OH | - | - | np | >60 | 45 | 18 | 23 | 3 | 10 | nt | 25 | 25 | 25 | 25 | 25 | 45 | 45 | 45 | np | np | np | np | 35 |
| 5 | C | N | C | C | C | $(CH_2)_4CH_3$ | $(CH)_2CN$ | - | - | >60 | >60 | 31 | 3 | 4 | 2 | 10 | nt | 25 | 25 | 10 | np | 25 | np | 45 | 45 | np | nt | 15 | nt | np |
| 6 | C | N | C | C | C | $(CH_2)_6CH_3$ | $O(CH_2)_5CH_3$ | F | $X_7$ | np | >60 | >60 | np | np | np | 25 | nt | 25 | 25 | np | 25 | 25 | nt | nt | nt | nt | nt | nt | nt | np |
| 7 | C | N | C | C | C | $(CH_2)_4CH_3$ | $(CH)_2CONH_2$ | - | - | np | >60 | np | >60 | nt | nt | 25 | nt | 10 | 25 | 25 | 25 | 25 | nt | nt | nt | nt | nt | nt | nt | np |
| 8 | C | N | C | C | C | $(CH_2)_4CH_3$ | $OCH_2CH_3$ | - | - | np | >60 | >60 | 4 | 8 | 2 | 10 | nt | 25 | 25 | 10 | np | 25 | np | 45 | np | nt | nt | np | nt | np |
| 9 | C | N | C | C | C | $(CH_2)_2CH_3$ | $OCF_3$ | - | - | np | np | nt | 38 | >60 | 15 | 10 | nt | 25 | 25 | 25 | 25 | 25 | nt | nt | nt | nt | nt | nt | nt | nt |
| 10 | C | N | C | C | C | $(CH_2)_6CH_3$ | $OCH_3$ | - | - | >60 | np | np | >60 | >60 | >60 | 10 | nt | 25 | 25 | 10 | 25 | 25 | nt | nt | nt | nt | nt | nt | nt | np |
| 11 | C | N | C | C | C | $(CH_2)_5CH_3$ | $OCH_3$ | - | - | np | nt | nt | 21 | nt | nt | 10 | nt | 25 | np | 25 | np | 25 | np | 45 | np | nt | nt | nt | nt | nt |
| 12-1 | C | N | C | C | C | $(CH_2)_2CH_3$ | $OCHF_2$ | - | - | 5 | 13 | 7 | 12 | 21 | 5 | 25 | nt | 25 | np | 10 | 25 | 25 | np | 45 | 45 | np | 45 | np | 45 | 60 |
| 12-5 | C | N | C | C | C | $(CH_2)_2CH_3$ | $SCHF_2$ | - | - | 12 | 13 | 6 | 5 | 8 | 7 | nt | 25 | nt | 10 | 25 | np | nt | np | 45 | 45 | np | 45 | np | np | 60 |
| 12-6 | C | C | C | C | C | $(CH_2)_2CH_3$ | $OCHF_2$ | - | - | np | np | 47 | 28 | 22 | 4 | 25 | 25 | nt | 25 | nt | 25 | nt | np | np | np | nt | nt | nt | nt | nt |
| 12-7 | N | C | C | C | C | $(CH_2)_2CH_3$ | $OCHF_2$ | - | - | 9 | 14 | 5 | 7 | 8 | 4 | nt | 25 | nt | 25 | 25 | np | nt | 45 | 45 | 45 | np | np | 15 | 45 | 60 |
| 12-8 | C | C | N | C | C | $(CH_2)_2CH_3$ | $OCHF_2$ | - | - | 9 | 15 | 7 | 21 | 19 | 4 | np | np | nt | 25 | 25 | np | nt | np | 45 | np | nt | nt | nt | nt | np |
| 12-9 | C | C | C | N | C | $(CH_2)_2CH_3$ | $OCHF_2$ | - | - | 42 | >60 | 15 | 31 | 30 | 8 | 25 | 25 | nt | 25 | nt | 25 | 25 | np | np | np | nt | nt | nt | nt | np |
| 12-10 | C | N | C | C | C | $CH_2CH_3$ | $OCHF_2$ | - | - | 10 | 18 | 7 | 12 | 27 | 4 | np | 25 | nt | np | nt | np | nt | nt | nt | nt | nt | nt | nt | nt | np |
| 12-11 | C | N | C | C | C | $CH_3$ | $OCHF_2$ | - | - | 30 | 29 | 17 | 29 | 32 | 18 | np | 25 | nt | np | nt | np | nt | np | 45 | np | nt | nt | nt | nt | np |
| 12-12 | C | N | C | C | C | H | $OCHF_2$ | - | - | 33 | >60 | 32 | 35 | 31 | 9 | np | 25 | nt | 25 | nt | np | nt | 45 | 45 | np | np | nt | nt | nt | np |
| 12-14 | C | N | C | C | C | $CH_3(C)_2$ | $OCHF_2$ | - | - | 14 | 12 | 5 | >60 | >60 | 6 | 25 | np | nt | np | nt | np | nt | np | 45 | np | nt | nt | nt | nt | np |
| 12-17 | C | N | C | C | C | $CH_2NH_2$ | $OCHF_2$ | - | - | np | np | >60 | np | np | >60 | 25 | 25 | nt | np | nt | np | nt | np | np | np | nt | nt | nt | nt | np |
| 12-20 | C | N | C | C | C | H | $OCHF_2$ | $(CH_2)_2CH_3$ | $X_{10}$ | np | >60 | 27 | np | np | 50 | np | 10 | nt | np | nt | np | nt | nt | nt | nt | nt | nt | np | nt | 60 |
| 12-21 | C | N | C | C | C | H | $OCHF_2$ | $(CH_2)_2CH_3$ | $X_2$ | 53 | np | 17 | 38 | np | 6 | 25 | 10 | nt | np | nt | np | nt | np | np | np | np | nt | nt | nt | nt |
| 12-22 | C | N | C | C | C | H | $OCHF_2$ | $(CH_2)_2CH_3$ | $X_3$ | >60 | np | 36 | np | >60 | 17 | np | 25 | nt | np | nt | np | nt | nt | nt | nt | nt | nt | nt | nt | nt |
| 12-23 | C | N | C | C | C | $(CH_2)_2CH_3$ | H | $OCHF_2$ | $X_5$ | 13 | np | 18 | 28 | np | 4 | np | 10 | nt | np | nt | np | nt | np | np | np | nt | nt | np | nt | 60 |
| 12-24 | C | N | C | C | C | $(CH_2)_2CH_3$ | H | $OCHF_2$ | $X_4$ | np | np | >60 | >60 | np | >60 | nt | nt | nt | nt | nt | nt | nt | nt | nt | nt | 45 | np | np | 45 | 60 |
| 12-25 | C | N | C | C | N | $(CH_2)_2CH_3$ | $OCHF_2$ | - | - | 20 | 30 | 10 | 8 | 18 | 6 | 25 | 25 | nt | np | nt | np | nt | np | 45 | 45 | 45 | np | 45 | 45 | 60 |
| 12-26 | C | N | C | C | C | $CH_2CH(CH_3)_2$ | $OCHF_2$ | - | - | 22 | 35 | 2 | 15 | 21 | 19 | np | 25 | nt | 25 | 25 | np | nt | np | 45 | np | np | np | 45 | np | np |
| 12-27 | C | N | C | C | C | $(CH_2)_3CH_3$ | $OCHF_2$ | - | - | 11 | 15 | 4 | 15 | 11 | 4 | 25 | 25 | nt | np | nt | np | nt | np | 45 | 45 | 45 | np | np | np | 60 |
| 12-28 | C | N | C | C | C | $(CH_2)_2CH_3$ | $OCH_3$ | - | - | 19 | 22 | 6 | 11 | 12 | 2 | np | 25 | nt | 25 | nt | 25 | nt | 45 | 45 | 45 | np | np | 15 | 45 | 60 |
| 12-29 | C | N | C | C | C | $(CH_2)_2CH_3$ | OH | - | - | np | np | 52 | np | >60 | 59 | 25 | np | nt | np | nt | np | nt | np | np | 45 | 45 | 45 | 45 | np | np |
| 12-30 | C | N | C | C | C | $(CH_2)_2CH_3$ | $OHCH_2CHF_2$ | - | - | np | np | 6 | 4 | 8 | 2 | nt | nt | nt | nt | nt | np | nt | nt | nt | nt | 45 | np | np | np | np |
| 13 | C | N | C | C | C | $(CH_2)_6CH_3$ | $OCHF_2$ | - | - | >60 | >60 | 18 | 48 | 44 | 2 | 25 | nt | 25 | 25 | 25 | 25 | 25 | 45 | np | np | 45 | np | np | np | np |
| 14 | C | N | C | C | C | $(CH_2)_4CH_3$ | I | - | - | >60 | np | 56 | >60 | >60 | 2 | 10 | nt | 25 | nt | 25 | np | 25 | np | np | 45 | np | np | np | np | np |
| 15 | C | N | C | C | C | $(CH_2)_4CH_3$ | Br | - | - | >60 | >60 | 31 | >60 | >60 | 2 | 10 | nt | 25 | 25 | 25 | np | 25 | np | 45 | np | nt | nt | nt | nt | np |
| 16 | C | N | C | C | N | $(CH_2)_7CH_3$ | OH | - | - | np | >60 | 44 | 17 | 25 | 5 | 25 | 25 | nt | np | nt | np | nt | 45 | 45 | 45 | np | nt | nt | nt | 60 |
| 17 | C | N | C | C | N | $(CH_2)_5CH_3$ | $H_2N$ | - | - | 58 | 22 | 20 | 32 | 28 | 2 | nt | 25 | nt | 10 | 25 | 25 | nt | 45 | 45 | 45 | np | nt | 45 | nt | 55 |
| 18 | C | N | C | C | N | $(CH_2)_3CH_3$ | Br | - | - | >60 | np | >60 | 7 | 9 | 5 | 25 | 25 | nt | np | np | 25 | nt | 45 | 45 | 45 | np | np | 45 | np | np |
| levamisole | | | | | | | | | | 48 | 32 | 44 | 32 | 48 | 44 | 10 | 10 | 10 | 10 | 10 | 10 | 10 | nt | nt | nt | nt | np | np | np | np |
| ivermectin | | | | | | | | | | <2 | <2 | 2 | <2 | <2 | 2 | 10 | 10 | 10 | 10 | 10 | 10 | 10 | nt | nt | nt | nt | 5 | <4 | np | nt |
| fluopyram | | | | | | | | | | <2 | <2 | 3 | <2 | 1 | 3 | nt | nt | nt | nt | nt | nt | nt | 45 | 45 | 45 | 45 | np | np | 15 | np |
| tioxazafen | | | | | | | | | | 5 | 4 | <2 | 4 | 5 | <2 | nt | nt | nt | nt | nt | nt | nt | 45 | 45 | 45 | 45 | np | np | 45 | np |
| oxamyl | | | | | | | | | | nt | nt | nt | nt | nt | nt | nt | nt | nt | nt | nt | nt | nt | np | 45 | 45 | 45 | 5 | 25 | np | np |
| iprodione | | | | | | | | | | np | >60 | 60 | >60 | np | 60 | nt | nt | nt | nt | nt | nt | nt | nt | nt | nt | np | 15 | 45 | nt | np |
| solvent | | | | | | | | | | np | np | np | np | np | np | np | np | np | np | np | np | np | np | np | np | np | np | np | np | np |

free-living nematode potency (µM): 0 – ≥60

nematode parasite activity scale (colour represents % of animals with phenotype): ≤20 | >20<50 | ≥50

non-target model activity scale (%): ≤20 | >20<50 | ≥50

**Supplementary Table 3.** Nementin structure-activity relationship analysis.

[A]The analog structures are represented by the Markush structure at the top and includes atom and substituent identifiers.

[B]Data for the free-living nematodes are EC50s ($\mu$M) and color-coded based on the potency scale at the bottom of this section.

[C-D]Nematode parasites of mammals or plants were tested with a limited number of compound concentrations. In each cell in these sections, the lowest concentration ($\mu$M) tested that exhibited a phenotype is shown, and the percentage of animals that exhibit the phenotype is indicated by the colour scale at the bottom of this section.

[E]Data for models of non-targeted systems is reported either as an EC50 ($\mu$M) (for the HEK293 cells) or as the lowest concentration tested ($\mu$M) that exhibited a discernible phenotype compared to wild-type at the indicated concentration; the percentage of animals that exhibit the phenotype is indicated by the colour scale at the bottom of this section. np, no phenotype (< 20% effect); nt, not tested. See the Methods for a description of each assay.

**Supplemental Movie 1.**

**Example of typical wild type *Caenorhabditis elegans* locomotion.** Wild type *C. elegans* (strain N2) are swimming on solid media containing only 1% DMSO solvent control that is incorporated into the agar. Animals have been swimming on the plate for 80 minutes at the time the movie was made.

**Supplemental Movie 2.**

**Example of compound-induced convulsions.** Wild type *C. elegans* (strain N2) are shown after swimming on solid media containing 60 $\mu$M Nementin-1 for 80 minutes.

**Supplemental Movie 3.**

**Example of compound-induced coiling.** Wild type *C. elegans* (strain N2) are shown after swimming on solid media containing 60 $\mu$M wact-45 for 80 minutes.

**Supplemental Movie 4.**

**Example of compound-induced shaking.** Wild type *C. elegans* (strain N2) are shown after swimming on solid media containing 60 $\mu$M Nementin-1 for 80 minutes. At the 18 second mark of the movie, *C. elegans* (strain N2) are shown after swimming on solid media containing 60 $\mu$M wact-203 for 80 minutes. At the 21 second mark of the movie, wild type *Rhabditophanes diutinus* animals are shown after swimming on solid media containing 60 $\mu$M Nementin-1 for 170 minutes.

**Supplemental Movie 5.**

**Example of compound-induced jerky-unc phenotype.** Wild-type *C. elegans* (strain N2) are shown after swimming on solid media containing 60 $\mu$M wact-45 for 80 minutes.

**Supplemental Movie 6.**

**Example of compound-induced reversal-defective phenotype.** Wild type *C. elegans* (strain N2) are shown after swimming on solid media containing 60 $\mu$M wact-38 for 80 minutes.

**Supplementary Data 1. (Separate file)**

Egg-laying rates with and without a stimulatory cocktail.

**Supplementary Data 2. (Separate file)**

The Wactive Library egg-laying screen data.

**Supplementary Data 3. (Separate file)**

The locomotory survey of the egl-modulators.

**Supplementary Data 4. (Separate file)**

Nematode and Counter-Screen Bioassay Data.
